# Upstream open reading frames buffer translational variability during *Drosophila* evolution and development

**DOI:** 10.1101/2024.11.13.623404

**Authors:** Yuanqiang Sun, Yuange Duan, Peixiang Gao, Chenlu Liu, Kaichun Jin, Shengqian Dou, Wenxiong Tang, Hong Zhang, Jian Lu

**Affiliations:** State Key Laboratory of Gene Function and Modulation Research, Center for Bioinformatics, School of Life Sciences, Peking University, Beijing 100871, China; Eye Institute of Shandong First Medical University, State Key Laboratory Cultivation Base, Shandong Provincial Key Laboratory of Ophthalmology, Qingdao 266071, China; College of Ecology, Lanzhou University, Lanzhou 730000, China; Beijing Advanced Center of RNA Biology (BEACON), Peking University, Beijing 100871, China; Southwest United Graduate School, Kunming 650092, China

**Keywords:** Upstream open reading frames (uORFs), TASEP modeling, stabilizing selection, translational buffer, interspecific gene expression, canalization, *bicoid*

## Abstract

Protein abundance tends to be more evolutionarily conserved than mRNA levels both within and between species, yet the mechanisms underlying this phenomenon remain largely unknown. Upstream open reading frames (uORFs) are widespread *cis*-regulatory elements in eukaryotic genomes that regulate translation, but it remains unclear whether and how uORFs contribute to stabilizing protein levels. In this study, we performed ribosome translation simulations on mRNA to quantitatively assess the extent to which uORF translation influences the translational variability of downstream coding sequences (CDSs) across varying contexts. Our simulations revealed that uORF translation dampens CDS translational variability, with buffering capacity increasing in proportion to uORF efficiency, length, and number. We then compared the translatomes at different developmental stages of two *Drosophila* species, demonstrating that uORFs buffer mRNA translation fluctuations during both evolution and development. Experimentally, deleting a uORF in the *bicoid* (*bcd*) gene—a prominent example of translational buffering—resulted in extensive changes in gene expression and phenotypes in *Drosophila melanogaster*. Additionally, we observed uORF-mediated buffering between primates and within human populations. Together, our results reveal a novel regulatory mechanism by which uORFs stabilize gene translation during development and across evolutionary time.

## Introduction

Organisms have evolved various strategies for the spatiotemporal regulation of gene expression ^1–3^. This is important because aberrant gene expression can result in phenotypic defects or diseases, while the variation and evolution of gene expression patterns frequently promote phenotypic diversification and adaptation ^4^. Although variations in mRNA abundance are widely observed within or between species, protein abundance tends to show stronger evolutionary constraint ^5,6^, as observed in yeasts ^7–9^, primates ^10,11^, and other organisms ^12–14^. Nevertheless, the molecular mechanisms by which the conservation of protein abundance across species is achieved are largely unknown ^5,6^.

Eukaryotic mRNA translation is a crucial step in gene expression and is highly regulated by multilayered mechanisms ^15–17^. Upstream open reading frames (uORFs), which are short open reading frames in the 5-terminal untranslated regions (5’ UTRs) of eukaryotic mRNAs, play crucial roles in regulating mRNA translation. Approximately 50% of eukaryotic genes contain uORFs ^18^, and their evolution has been tightly shaped by natural selection ^19–23^. The functions of uORFs have been explored in various contexts, including development ^23–30^, disease ^31–35^, and stress responses ^36–43^. The prevailing consensus is that uORFs typically repress downstream coding sequence (CDS) translation by sequestering ribosomes, a process influenced by factors such as uORF length, position, and sequence context ^34,43–48^. However, under stress conditions, certain uORFs can facilitate CDS translation by promoting ribosome reinitiation, illustrating their context-dependent functions ^42,43,48–52^.

Gene expression noise, which arises from the inherent stochasticity of biological processes such as transcription and translation, is generally detrimental to organismal fitness ^53^ and is primarily determined at the translational level ^54^. Recent studies suggest uORFs might play essential roles in buffering translational noise and stabilizing protein expression. For example, Wu et al. (2022) demonstrated that uORFs reduce protein production rates to stabilize TOC1 protein levels, ensuring precise circadian clock function in plants ^55^. Similarly, Bottorff et al. (2022) used a human cell reporter system to show that a ribosome stall in cytomegaloviral *UL4* uORFs buffers against CDS translation reductions ^56^. Under stress, translation initiation is typically downregulated, yet most human mRNAs resistant to this inhibition contain translated uORFs, with a single uORF often being sufficient for resistance ^42^. The computational model of Initiation Complexes Interference with Elongating Ribosomes (ICIER) suggests that derepression of downstream translation is a general mechanism of uORF-mediated stress resistance ^43^. Despite these findings, the current understanding of uORFs in stabilizing translation is limited to single-gene cases or stressed conditions. It remains unclear whether and how uORFs affect gene translation variability on a genome-wide scale during evolution and development, and whether the identified mechanisms are universal or vary substantially among different taxa. To address these questions, a combination of modeling, genome-wide analyses, and comparative studies across species is required.

In this study, we first adapted the ICIER framework ^43^ to simulate the translating ribosome on a mRNA to quantitatively measure the extent to which uORF translation reduces the translational variability of the downstream CDS under different translation contexts. We then compared the translatomes of two closely related *Drosophila* species, *D. melanogaster* and *D. simulans*, and further supported the notion that uORFs could buffer the fluctuations of CDS translation during the development and evolution of *Drosophila*. The patterns also reappeared among primates and human populations. We next knocked out the *bicoid* (*bcd*) uORF, a case showing the significant buffering effect in our data, and observed broad changes in the embryonic transcriptome and phenotypic defects in *D. melanogaster*. Together, our results demonstrate a novel role for uORFs by maintaining translation stabilization during *Drosophila* evolution and development.

## Results

### An extended ICIER model for quantifying uORF buffering in CDS translation

To quantitatively assess how uORF translation modulates the variability of downstream CDS translation, we adapted the ICIER model ^43^, originally grounded in the totally asymmetric simple exclusion process (TASEP). TASEP has been extensively utilized to model the stochastic nature of ribosome movement along mRNA, capturing the effects of ribosome traffic jams, where ribosomes may slow down or stall when a site ahead is occupied ^57–60^. The ICIER model extends this by simulating the interplay between scanning (40S) and elongating (80S) ribosomes, particularly focusing on how uORFs impact the overall translation rate of the main CDS ^43^.

We extended the original ICIER model ^43^ with several major modifications (Figure 1A). First, while the original ICIER model only considered the scenario where the elongating ribosome (80S) causes downstream scanning ribosomes (40S) to dissociate from the mRNA when they move along the mRNA and collide, recent findings have shown that upstream dissociation can also play a critical role in uORF-mediated regulation ^56^. To incorporate this, we accounted for more complex ribosome interactions, including three possible scenarios where the 80S collides with 40S: i) 80S only causes the downstream 40S to dissociate from the mRNA with a probability of *K_down_*(“downstream dissociation”, *K_down_* ranging from 0 to 1), following the original ICIER model; ii) the 80S only causes the upstream 40S to dissociate from the mRNA with a probability of *K_up_* (“upstream dissociation”, *K_up_* > 0 and *K_down_* = 0); and iii) a combination of the downstream and upstream dissociation models (“double dissociation”, *K_up_* > 0 and *K_down_* > 0).

**Figure 1.**
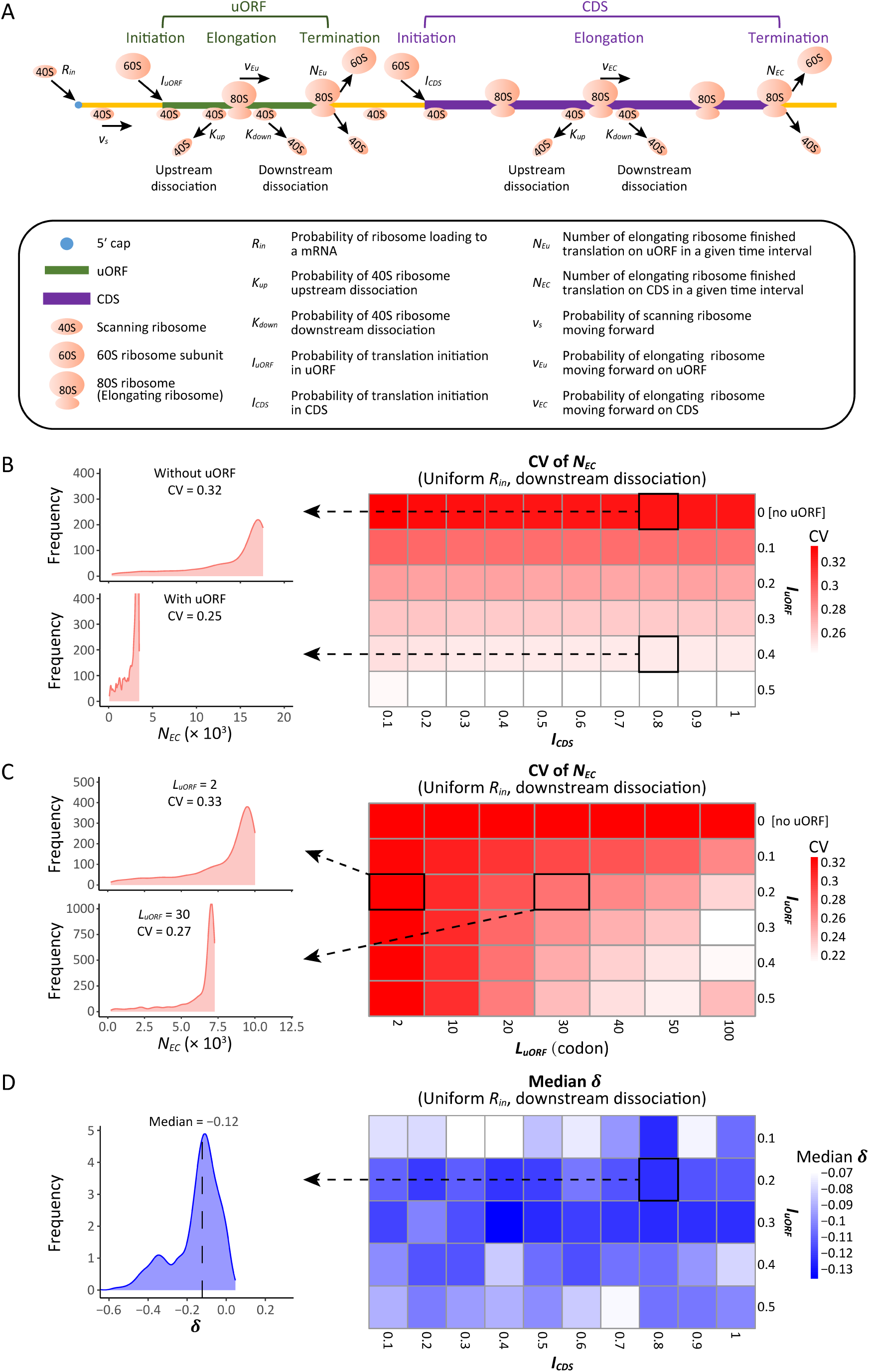
Modeling simulation of uORF-mediated translation buffering. (A) Model schema of the modified ICIER model (on the top); the parameters are listed in the box below the schema. (B) Heatmap showing the CVs of CDS translation rate (*N_EC_*) under different *I_CDS_* (x-axis) and *I_uORF_* (y-axis) combinations with a uniform distribution of *R_in_* input and the downstream dissociation model. The left panels elicited by the dotted lines from specific squares of right heatmap were two examples showing the distribution of *N_EC_* under *I_CDS_* = 0.8 & *I_uORF_* = 0 (top panel, without uORF) and *I_CDS_* = 0.8 & *I_uORF_* = 0.4 (bottom panel, with uORF). (C) Heatmap showing CVs of CDS translation rate (*N_EC_*) under different *L*_*uORF*_ (x-axis) and *I_uORF_* (y-axis) combinations with a uniform distribution of *R_in_* input and the downstream dissociation model. The left panels elicited by the dotted lines from specific squares of right heatmap were two examples showing the distribution of *N_EC_*under *L_uORF_* = 2 & *I_uORF_* = 0.2 (top panel) and *L_uORF_* = 30 & *I_uORF_* = 0.2 (bottom panel). (D) Heatmap showing median *δ* [*log*_2_(Δ*N*_*EC*_/Δ*N*_*EU*1_)] under different *I_CDS_* (x-axis) and *I_uORF_* (y-axis) combinations with a uniform distribution of *R_in_* input and the downstream dissociation model. The left panel elicited by the dotted line from a specific square of right heatmap was an example showing the distribution of *δ* under *I_CDS_* = 0.8 & *I_uORF_* = 0.2. The vertical dashed line indicated the median value of *δ*.

Second, the original ICIER model only considered the ribosome collision and dissociation in the uORF and counted the 40S scanning ribosome escaping from the uORF as a proxy of the CDS translation rate. In our extended model, we also considered the ribosome collision and dissociation events in the CDS downstream of the uORF. We recorded the number of 80S ribosomes that completed translation at the stop codon of a CDS (*N*_*EC*_) or uORF (*N*_*Eu*_) during a given time interval, using these counts as proxies to quantify the translation rate of CDS or uORF. These indices allowed us to directly and quantitively measure the impact of uORFs on CDS translation.

Third, while the original ICIER model accounted for only a single uORF within an mRNA, we extended it to consider two uORFs coexisting on an mRNA molecule, allowing us to explore the possible combinatorial effects of uORF-mediated translation regulation in a more complex yet more common scenario, as previous studies have shown that uORFs tend to be clustered within genes ^18^.

These extensions allow for a more comprehensive exploration of uORF-mediated translational buffering, offering deeper insights into how these regulatory elements might stabilize protein synthesis across varying translation contexts.

### uORF-mediated buffering of CDS translation across different parameter settings

To systematically investigate the extent to which uORF translation modulates the variability of downstream CDS translation, we conducted simulations across various parameter settings using three dissociation models: upstream, downstream, and double dissociation, each reflecting different possible interactions between scanning and elongating ribosomes on the mRNA. We considered a range of parameters crucial to the translation process (Table S1), including the length of the 5’ leader before the uORF (fixed at 150 nucleotides), the length of the uORF itself (ranging from 2 to 100 codons), the distance between the uORF stop codon and the CDS start codon (150 nucleotides), the length of the CDS (500 codons), and the length of the 3’ UTR (150 nucleotides). Additionally, we modeled the probabilities associated with ribosome movement and initiation, such as the probability of a 40S ribosome moving to the next nucleotide (*v*_*s*_ = 0.3), the probability of an 80S ribosome moving to the next position within the uORF (*v*_*Eu*_ = 0.3) and within the CDS (*v*_*EC*_ = 0.5), and the probability of loading a new 40S ribosome at the 5’ end of the mRNA (*R_in_*). We also explored different probabilities of translation initiation at both the uORF start codon (*I*_*uORF*_) and the CDS start codon (*I*_*CCS*_). Key parameters were adapted from the original ICIER model ^43^, ensuring a robust basis for comparison while allowing exploration of additional variables that influence uORF-mediated translational buffering.

In our simulations, *R*_*in*_ values were varied to simulate fluctuations in translational resources, such as ribosome availability, that could arise from genetic differences or environmental changes during evolution or development. By generating 1,000 *R*_*in*_ values following either uniform or exponential distributions (ranging from 0 to 0.1, Figure S1A), we aimed to capture the natural variability in ribosome loading rates that might occur across different species, individuals, or developmental stages.

These values were then fed into the simulation models to evaluate their impact on both the level and variability of translation rate for CDSs, with the number of ribosomes completing translation on CDSs (*N*_*EC*_) in a given time interval used as a proxy for translational rate (Figure S1B).

Across all model settings, uORF translation (*I*_*uORF*_ > 0) consistently reduced CDS translation rate (*N*_*EC*_) by about 30% to 80% as *I*_*uORF*_ increased from 0.1 to 0.5, compared to scenarios where the uORF was absent or untranslated (*I*_*uORF*_ = 0) (Figure S2 and S3). This confirms the inhibitory effect of uORFs on downstream CDS translation. Notably, the coefficient of variation (CV), a measurement of variability for translation rate (*N*_*EC*_), was lower when the uORF was translated (Figure 1B). These CV values further decreased by approximately 10% to 25% as *I*_*uORF*_ increased from 0.1 to 0.5 (Figure 1B), and this buffering effect persisted across different parameter settings (Figure S4 and S5). Moreover, the CV of translation rate (*N*_*EC*_) further decreased by about 6% to 30% as uORF length (*L*_*uORF*_) increased from 2 to 100 codons (Figure 1C and S6-7). These simulation results indicate that uORF translation can reduce variability in downstream CDS translation, with the buffering capacity positively correlated with both uORF translation initiation efficiency and length under the applied simulation conditions.

To more quantitatively investigate the relationship between changes in translation rate in uORF (*N*_*EU*_) and CDS (*N*_*EC*_), for each parameter setting, we used the median value (*R*_*inm*_) of the 1,000 *R*_*in*_ inputs as the baseline, calculating the corresponding *N*_*ECm*_ and *N*_*EUm*_values. We then calculated the changes of *N*_*EC*_ (*ΔN*_*EC*_) and *N*_*EU*_ (*ΔN*_*EU*_) relative to *N*_*ECm*_ and *N*_*EUm*_ for each *R*_*in*_ value, respectively. Across different parameter settings, *ΔN*_*EU*_ was consistently and significantly positively correlated with *ΔN*_*EC*_ (*P* < 0.001, Spearman’s correlation) (Figure S8-9), indicating that fluctuations of the ribosome loading rate influence the translation of both uORFs and CDSs in the same direction. Nevertheless, our simulations showed that variations in *R*_*in*_ led to a larger change in *N*_*EU*_ than in *N*_*EC*_, as the median value of *δ* [defined by log_2_(*ΔN*_*EC*_/*ΔN*_*EU*_)] was consistently less than 0 across various *I*_*uORF*_ and *I*_*CCS*_ combinations (Figures 1D and S10-11). This finding suggests that translational fluctuation in uORFs is greater than that in downstream CDSs, indicating that uORFs can buffer against upstream fluctuations.

Simulations of mRNAs with two uORFs revealed patterns consistent with a buffering role for uORFs (Figure S12). To compare the buffering effects of a single uORF versus two uORFs, we calculated the ratio of the CV of *N*_*EC*_ with two uORFs to that with a single uORF. A ratio less than 1 suggests that two uORFs provide greater buffering than a single uORF. For comparability, we examined the CV of *N*_*EC*_ where the *I*_*uORF*_ in the single-uORF model equals *I*_*uORF*1_ (*I*_*uORF*_ of the first uORF) in the two-uORF model, both ranging from 0 to 0.5. The CV ratio consistently remained below 1 across a range of *I*_*uORF*2_ (*I*_*uORF*_ of the second uORF) (Figure S13), indicating that two uORFs offer stronger buffering than a single uORF.

Collectively, these simulations collectively suggest: (1) uORF-mediated translational control buffers against CDS translation variability, (2) uORFs exhibit greater translational fluctuations than downstream CDSs, and (3) the buffering capacity of uORFs positively correlates with their translation initiation efficiency and length. Subsequently, we sought to validate these simulation results in a biological context by confirming the uORF-mediated buffering effect during organismal evolution and development, as both processes face environmental and/or genetic changes that frequently disturb mRNA translation.

### Generating matched translatome data from two *Drosophila* species for comparative analysis

To validate the uORF-mediated translational buffering during *Drosophila* evolution, we performed a comparative analysis of the translatomes of two closely related *Drosophila* species, *D. melanogaster* and *D. simulans*, which diverged approximately 5.4 million years ago ^61^. We generated high-throughput sequencing data for *D. simulans*, including transcriptome (mRNA-Seq) and translatome (Ribo-Seq) profiles from various developmental stages and tissues, including embryos at 0–2 h, 2–6 h, 6–12 h, and 12–24 h, third-instar larvae, P7–8 pupae, female and male bodies, and female and male heads. In total, we obtained approximately 786 million high-quality reads for *D. simulans* (Table S2). These datasets were designed to be directly comparable to the previously published *D. melanogaster* data ^23^, with identical embryonic stages and tissue types. For each sample, we followed established procedures ^62–66^ to calculate the translational efficiency (TE) for each feature (CDS or uORF). TE serves as a proxy for the translation rate at which ribosomes translate mRNA into proteins, typically quantified by comparing the density of ribosome-protected mRNA fragment (RPF) to the mRNA abundance for that feature (see **Materials and Methods**). This comprehensive comparative translatome analysis, utilizing matched developmental stages and tissues between the two species, allowed us to assess the uORF-mediated translational buffering effects during evolution.

### Translational conservation and dominance of uORFs between *Drosophila* species

Given that the translation of uORFs is a crucial determinant of their functional impact, we first characterized the translational profiles of uORFs in *D. melanogaster* and *D. simulans* to explore the evolutionary roles of uORFs in translation regulation. We focused on canonical uORFs that initiate with an ATG start codon in the 5′ UTR and terminate with a stop codon (TAA, TAG, or TGA). Because the ATG start codon is the defining feature of a canonical uORF and tends to be more conserved than its downstream sequence ^67^, we defined uORF conservation based on the presence of the ATG start codon in the 5′ UTR of *D. melanogaster* and its orthologous positions in *D. simulans*, regardless of differences in the stop codon. Using this criterion, we identified 18,412 canonical uORFs with conserved start codons between the two species. Additionally, we identified 2,789 canonical uORFs specific to *D. melanogaster* and 2,440 canonical uORFs specific to *D. simulans*. The TE values of the conserved uORFs were highly correlated between the two species across all developmental stages and tissues examined, with Spearman correlation coefficients (*rho*) ranging from 0.478 to 0.573 (Figure 2A). In contrast, TE of CDSs exhibited a significantly higher correlation between the two species in the corresponding samples compared to that of uORFs, with Spearman’s *rho* ranging from 0.588 to 0.806 (*P* = 0.002, Wilcoxon signed-rank test; Figure 2A). This observation is consistent with our simulation results, which indicate that uORFs experience greater translational fluctuations than their downstream CDSs. Notably, conserved uORFs exhibited significantly higher TEs compared to species-specific uORFs in both species. The median TE of conserved uORFs was 1.62 times that of non-conserved uORFs in *D. simulans*, while the corresponding ratio in *D. melanogaster* was 1.52 (Figure 2B).

**Figure 2.**
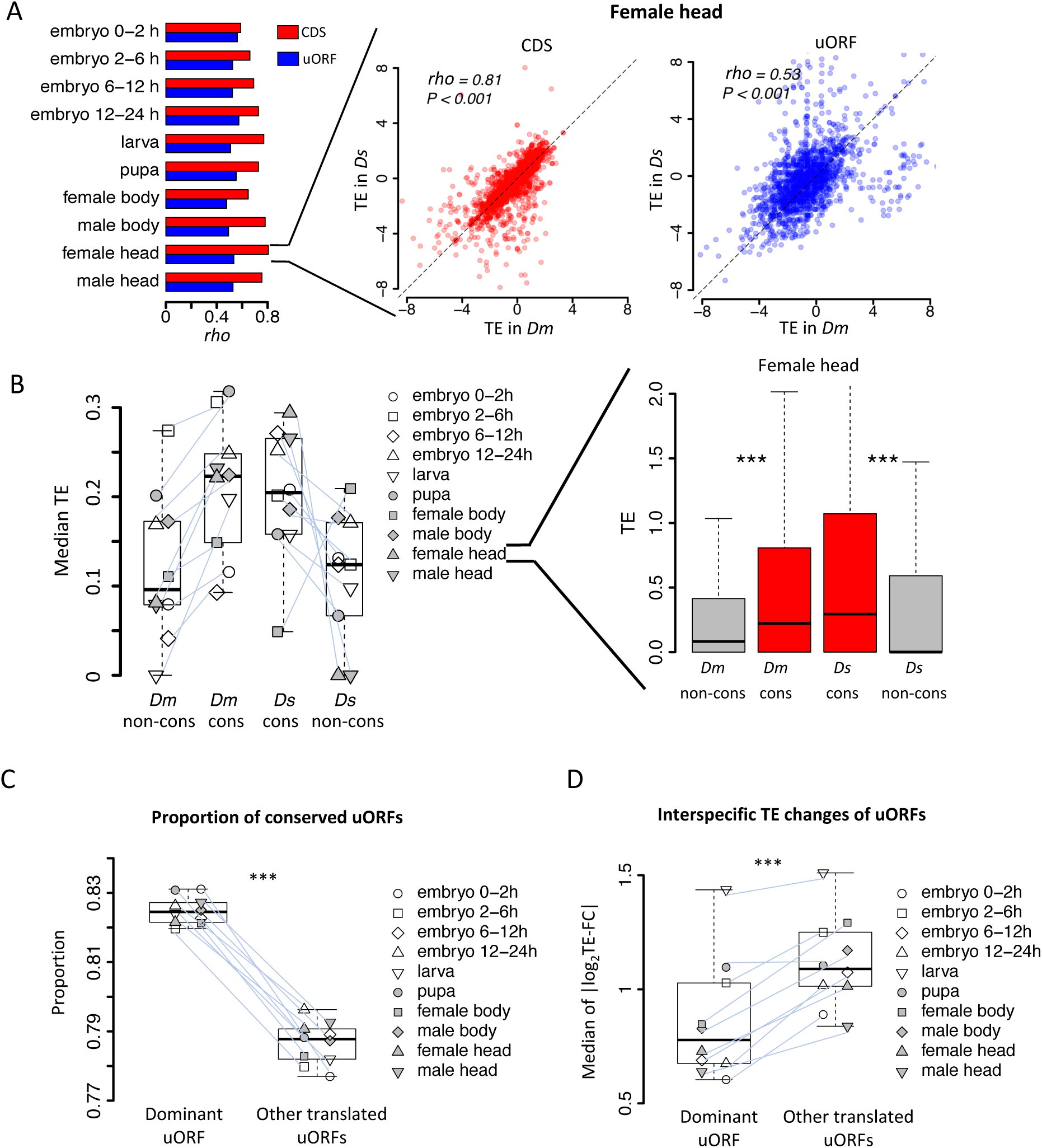
Conservation and translation of uORFs between *D. melanogaster* and *D. simulans*. (A) Spearman’s correlation coefficients (*rho*, represented by the bars) of TEs between *Dm* (*D. melanogaster*) and *Ds* (*D. simulans*) for CDS (red) and uORFs (blue). All *P*-values for Spearman’s correlation are less than 0.001. The *P* value for the comparison between *rho* values of CDSs TE and uORFs TE is 0.002 (Wilcoxon signed-rank test). Data for the female head sample is shown as an example in the right panel. The x- and y-axes represent the TEs in *Dm* and *Ds*. (B) The median of TE of conserved and species-specific uORFs in each sample. Each dot represents the median TE of a sample for a specific uORF class. Data from the female head sample is shown as an example in the right panel. *P* values were obtained from Wilcoxon rank-sum tests. ***, *P* < 0.001. (C) Fraction of conserved uORFs among dominant uORFs and other translated uORFs in each sample. The paired samples in *Dm* and *Ds* were linked together. The *P* value was obtained by the paired Wilcoxon signed-rank test. ***, *P* < 0.001. (D) Absolute values of the interspecific TE fold changes (log_2_TE-FC) of dominant uORFs and the other translated uORFs in each sample. The paired samples in *Dm* and *Ds* were linked together. The median value of each sample is shown. The *P* value was obtained via the paired Wilcoxon signed-rank test. ***, *P* < 0.001. Data from the female head sample were used as an example in the right panel.

In *D. melanogaster*, 7,259 (52.2%) genes had no uORFs, 2,687 (19.3%) had a single uORF, and 3,961 (28.5%) contained multiple uORFs. Among genes with multiple uORFs, we defined the uORF with the highest TE as the dominant uORF for that gene, as TE is one of the most relevant metrics for assessing uORF function ^45,67^. The median TE of the dominant uORF was 4.84 times that of the second-highest uORF within the same gene in *D. melanogaster*, and the corresponding ratio was 5.21 times in *D. simulans* (Figure S14). To assess the consistency of this dominance across different tissues and developmental stages, we identified 3,072 multiple-uORF genes in *D. melanogaster* with at least one translated uORF (TE > 0.1 in at least five stages/tissues). Of these, 569 genes consistently used the same dominant uORF across the measured samples, significantly higher than the number expected under randomness (5 genes, 95% confidence interval: 1–10) based on shuffling the TEs of uORFs 1,000 times (Figure S15). This trend was also observed in *D. simulans* and persisted under different thresholds for defining “translated uORFs” (Figure S15). These results suggest that genes with multiple uORFs tend to retain the same dominant uORF across developmental stages, indicating that the dominant uORF may serve as the key translational regulator of the downstream CDS. Moreover, we found that the dominant uORFs showed a higher proportion of conserved uATGs than the other translated uORFs (median proportion 82.5% versus 78.8%, *P* < 0.001) (Figure 2C). Additionally, the absolute values of TE fold-change (|log_2_TE-FC|) between the two species were, on average, 23.2% smaller for dominant uORFs than for other conserved uORFs (Figure 2D), suggesting that dominant uORFs are more likely under stronger stabilizing selection. These findings suggest that, in genes with multiple uORFs, the dominantly translated uORF may play a more important role in regulating CDS translation than the other uORFs.

### uORFs buffer interspecific translational divergence of CDSs

To investigate the relationship between translational changes in uORFs and their downstream CDSs across species, we analyzed the translation efficiency (TE) of uORFs and corresponding downstream CDSs in *D. melanogaster* and *D. simulans*. Consistent with our simulations that the translation of a uORF is tightly linked to that of downstream CDS, uORFs exhibited a modest, yet statistically significant, positive correlation with the TE of their downstream CDSs across all samples analyzed (*P* < 0.001, Spearman’s correlation) (Figure 3A). We then compared the interspecific TE change of a uORF (*β*_*u*_ = *TE*_*uORF*,*sim*_/*TE*_*uORF*,*mel*_) with that of its corresponding CDS (*β*_*C*_ = *TE*_*CCS*,*sim*_/*TE*_*CCS*,*mel*_) between *D. melanogaster* and *D. simulans*. We found *β*_*u*_ is significantly positively correlated with *β*_*C*_ across all samples (*P* values < 0.001, Spearman’s correlation) (Figure S16). These results align well with our simulations, which showed that fluctuations in translational factors (such as ribosomes) influence both uORF and CDS translation in the same direction (Figure S8–S9).

**Figure 3.**
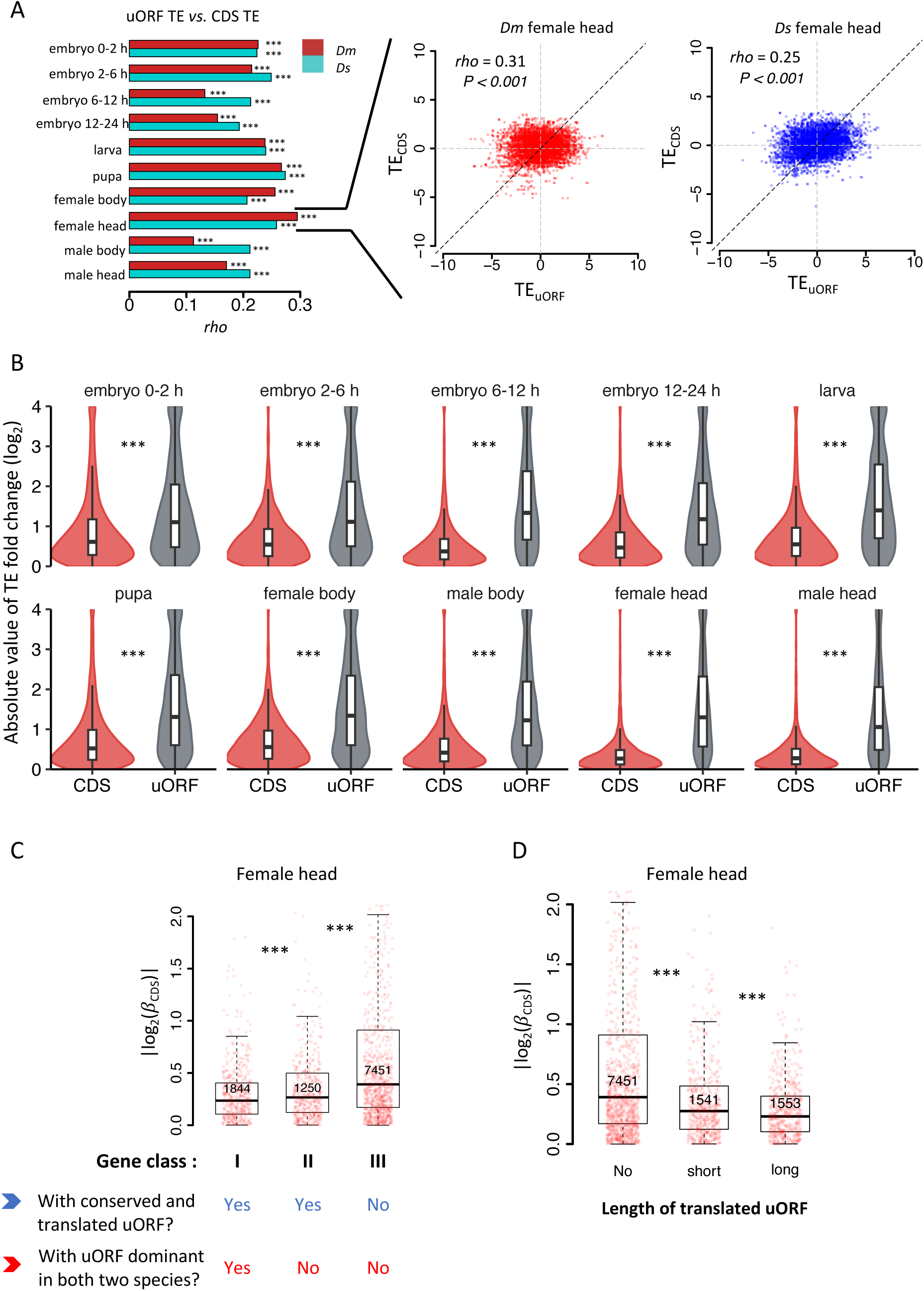
uORFs reduce CDS translational divergence between *D. melanogaster* and *D. simulans*. (A) The correlation of uORF TEs and the corresponding CDS TEs in 10 samples of *Dm* (*D. melanogaster*) and *Ds* (*D. simulans*). The bars represent Spearman’s correlation coefficient (*rho*). In all samples, we obtained both *P* values < 0.001. Data for the female head sample of *Dm* and *Ds* are shown as examples in the right panel. (B) The absolute values of interspecific TE changes for CDS and uORF in each sample between two species. For visualization purposes, all values greater than 4 were assigned a value of 4. ***, *P* < 0.001, Wilcoxon rank-sum test. (C) Genes expressed in female heads (mRNA RPKM > 0.1 in both species) were classified into three classes according to whether a gene had a conserved and dominantly translated uORF or not. Boxplots showing interspecific CDS TE variability |*log*_2_(*β*_*c*_)| of different gene classes. *P* values were calculated using Wilcoxon rank-sum tests between the neighboring groups. ***, *P* < 0.001. (D) Genes expressed in female heads were classified into three classes according to the length of translated uORFs. Boxplots showing interspecific CDS TE variability |*log*_2_(*β*_*c*_)| of different gene classes. *P* values were calculated using Wilcoxon rank-sum tests between the neighboring groups. ***, *P* < 0.001.

While the direction of TE changes for uORFs and CDSs tends to be consistent, our simulations suggest that the magnitude of TE changes in CDSs is generally smaller than that in uORFs, due to the buffering effect of uORF (Figure S10–S11). To validate this, we first identified uORFs and CDSs with significant interspecific TE differences by assessing whether *β*_*u*_ or *β*_*C*_ significantly deviated from 1, using an established statistical framework ^23^. This analysis uncovered 1,151 to 4,189 CDSs with significant interspecific TE changes (FDR < 0.05) (Table 1), with genes involved in development, morphogenesis, and differentiation being significantly enriched during embryonic stages (Figure S17). Conversely, genes related to metabolism, response to stimuli, and signaling were enriched in larval, pupal, and adult stages (Figure S17). Additionally, we identified 144 to 1,193 uORFs with significant TE differences between species, accounting for approximately 1–15% of expressed uORFs (Table 1). Note that due to their shorter length and generally lower TE, uORFs had considerably lower read counts than CDSs, limiting the statistical power to detect significant interspecific TE differences for uORFs. This trend consistently holds whether analyzing all expressed uORFs (Figure S18A) or only highly expressed genes (Figure S18B). Thus, the fewer uORFs showing significant TE divergence likely reflects lower read counts and statistical sensitivity rather than reduced translational variability relative to CDSs. In fact, the absolute values of log_2_(fold change) of TE for uORFs between *D. melanogaster* and *D. simulans* were significantly greater than those observed for corresponding CDSs across all samples (*P* < 0.001, Wilcoxon signed-rank test; Figure 3B), suggesting that the magnitude of TE changes in CDSs is generally smaller than that in uORFs, due to the buffering effect of uORF.

**Table 1.**
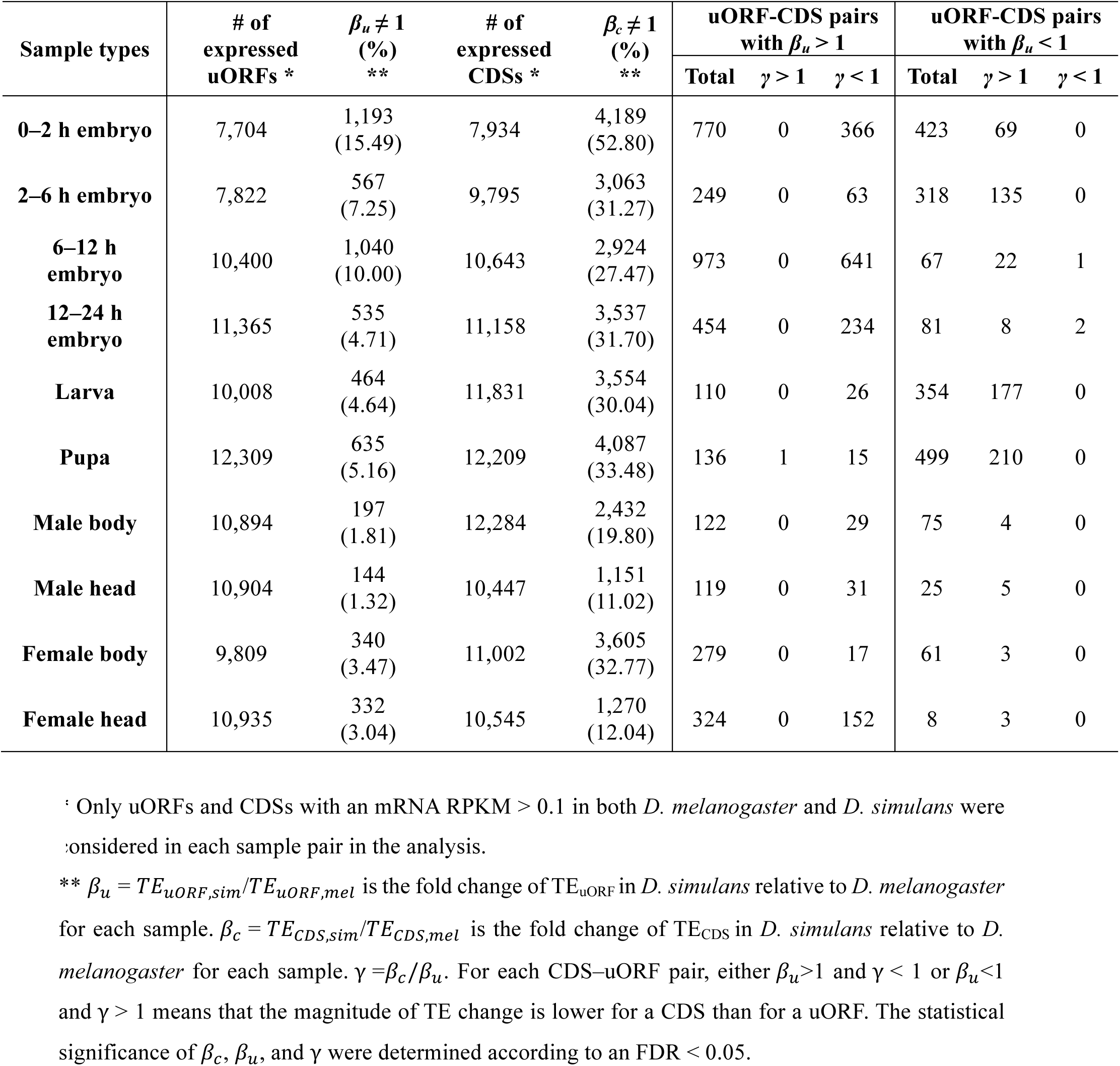
Numbers of genes showing different magnitudes of TE changes between uORFs and CDS at the interspecific level.

| Sample types | # of<br>expressed<br>uORFs * | $\beta_u \neq 1$<br>(%)<br>** | # of<br>expressed<br>CDSs * | $\beta_c \neq 1$<br>(%)<br>** | uORF-CDS pairs<br>with $\beta_u > 1$ | | | uORF-CDS pairs<br>with $\beta_u < 1$ | | |
| --- | --- | --- | --- | --- | --- | --- | --- | --- | --- | --- |
| | | | | | Total | $\gamma > 1$ | $\gamma < 1$ | Total | $\gamma > 1$ | $\gamma < 1$ |
| 0–2 h embryo | 7,704 | 1,193<br>(15.49) | 7,934 | 4,189<br>(52.80) | 770 | 0 | 366 | 423 | 69 | 0 |
| 2–6 h embryo | 7,822 | 567<br>(7.25) | 9,795 | 3,063<br>(31.27) | 249 | 0 | 63 | 318 | 135 | 0 |
| 6–12 h embryo | 10,400 | 1,040<br>(10.00) | 10,643 | 2,924<br>(27.47) | 973 | 0 | 641 | 67 | 22 | 1 |
| 12–24 h embryo | 11,365 | 535<br>(4.71) | 11,158 | 3,537<br>(31.70) | 454 | 0 | 234 | 81 | 8 | 2 |
| Larva | 10,008 | 464<br>(4.64) | 11,831 | 3,554<br>(30.04) | 110 | 0 | 26 | 354 | 177 | 0 |
| Pupa | 12,309 | 635<br>(5.16) | 12,209 | 4,087<br>(33.48) | 136 | 1 | 15 | 499 | 210 | 0 |
| Male body | 10,894 | 197<br>(1.81) | 12,284 | 2,432<br>(19.80) | 122 | 0 | 29 | 75 | 4 | 0 |
| Male head | 10,904 | 144<br>(1.32) | 10,447 | 1,151<br>(11.02) | 119 | 0 | 31 | 25 | 5 | 0 |
| Female body | 9,809 | 340<br>(3.47) | 11,002 | 3,605<br>(32.77) | 279 | 0 | 17 | 61 | 3 | 0 |
| Female head | 10,935 | 332<br>(3.04) | 10,545 | 1,270<br>(12.04) | 324 | 0 | 152 | 8 | 3 | 0 |

\* Only uORFs and CDSs with an mRNA RPKM > 0.1 in both *D. melanogaster* and *D. simulans* were considered in each sample pair in the analysis.
\*\* $\beta_u = TE_{uORF,sim}/TE_{uORF,mel}$ is the fold change of $TE_{uORF}$ in *D. simulans* relative to *D. melanogaster* for each sample. $\beta_c = TE_{CDS,sim}/TE_{CDS,mel}$ is the fold change of $TE_{CDS}$ in *D. simulans* relative to *D. melanogaster* for each sample. $\gamma = \beta_c/\beta_u$ . For each CDS–uORF pair, either $\beta_u > 1$ and $\gamma < 1$ or $\beta_u < 1$ and $\gamma > 1$ means that the magnitude of TE change is lower for a CDS than for a uORF. The statistical significance of $\beta_c$ , $\beta_u$ , and $\gamma$ were determined according to an FDR < 0.05.

We further quantitatively compared the magnitude of interspecific TE changes between uORFs and their corresponding CDSs using a previous method ^23^. We defined γ =*β*_*c*_/*β*_*u*_ for a uORF-CDS pair within the same mRNA and tested whether γ was significantly different from 1 to identify pairs with differential TE changes between uORFs and CDSs (Figure S19). When γ < 1 and *β*_*u*_ >1, or γ > 1 and *β*_*u*_ <1, it indicates that the TE change for the CDS is smaller than that for the uORF (Figure S19). Among CDS-uORF pairs where *β*_*u*_ > 1, nearly all (8–487) showed a significant γ < 1 in each sample, except for one pair from the pupal stage where γ > 1 (Table 1). This suggests that the magnitude of TE changes in CDSs was generally smaller than in uORFs when uORF TE increased, and vice versa when uORF TE decreased (*β*_*u*_ < 1) (Table 1). These comparative translatome analyses indicate that uORFs buffered downstream CDS translation changes during the evolutionary divergence of *D. melanogaster* and *D. simulans*.

### uORF buffering is influenced by its conservation, dominance, and length

To investigate how the conservation level and translation patterns of uORFs influence their buffering capacity on CDS translation, we categorized genes expressed in each pair of samples into three classes: Class I, genes with conserved uORFs that are dominantly translated (i.e., exhibiting the highest TE among all uORFs within the same gene) in both *Drosophila* species; Class II, genes with conserved uORFs that are translated in both species but not dominantly translated in at least one; and Class III, the remaining expressed genes. We then compared the absolute values of interspecific TE changes of the CDS (|*β*_*c*_ |) across these three categories. Significant differences in |*β*_*c*_ | were observed, with a consistent hierarchy of Class I < II < III across all pairs of samples (Figure 3C and S20). On average, Class I genes exhibited an average of 8.18% and 23.8% lower |*β*_*c*_| values compared to Class II and Class III, respectively. This indicates that conserved and dominantly translated uORFs exert a stronger buffering effect on CDS translation.

To further validate the simulation results suggesting that longer uORFs have a stronger buffering effect (Figure 1C and S6–7), we divided genes expressed in each pair of samples into three groups: those without translated uORFs (No), those with short uORFs (short, total length below the median), and those with long uORFs (long, total length above the median). Consistently, longer uORFs were associated with stronger buffering effects on CDS translation across all pairs of samples (Figure 3D and S21). Specifically, genes with longer uORFs showed 12.7% and 26.5% lower |*β*_*c*_| values compared to genes with short uORFs or no uORFs, respectively.

Overall, these findings underscore that the buffering capability of a uORF is positively correlated with its conservation level, translation dominance, and length.

### uORFs buffer translational fluctuations during *Drosophila* development

Gene expression undergoes dynamic changes during *Drosophila* development, with significant alterations in the translation program to meet developmental demands, including shifts in ribosome loading rates ^24,68,69^. Therefore, we extended our analysis to investigate the role of uORFs in buffering these translational fluctuations, hypothesizing that uORFs could mitigate them during development. According to our hypothesis, if a gene has a translated uORF in *D. melanogaster* but not in its orthologous gene in *D. simulans*, then the translation of this gene is likely more stable across developmental stages in *D. melanogaster* than its ortholog in *D. simulans*, and vice versa.

To test this hypothesis, we compared the CV of CDS TE across 10 developmental stages in *D. melanogaster* and in *D. simulans*, respectively. Genes with translated uORFs (TE>0.1 in at least one sample) in *D. melanogaster* exhibited significantly 22.5% smaller CVs in *D. melanogaster* than their orthologs lacking these uORFs in *D. simulans*, indicating more stable translation (Figure 4A). Consistently, for genes with translated uORFs in *D. simulans* but not in *D. melanogaster*, the CVs were 13.3% lower in *D. simulans* compared to their orthologs lacking these uORFs in *D. melanogaster* (Figure 4B). Consistent results were observed when a uORF is required to be translated (TE > 0.1) in all 10 samples (Figure S22A), in 4 embryonic stages (Figure S22B), or in 6 stages including embryos, larva, and pupa (Figure S22C). Moreover, within each species, genes with translated uORFs also showed less variability in CDS TE across developmental stages compared to those without translated uORFs, with a 31.8% reduction in *D. melanogaster* and a 28.9% reduction in *D. simulans* (Figure 4C). This effect was consistent across different thresholds for defining “translated uORFs” (Figure S23).

**Figure 4.**
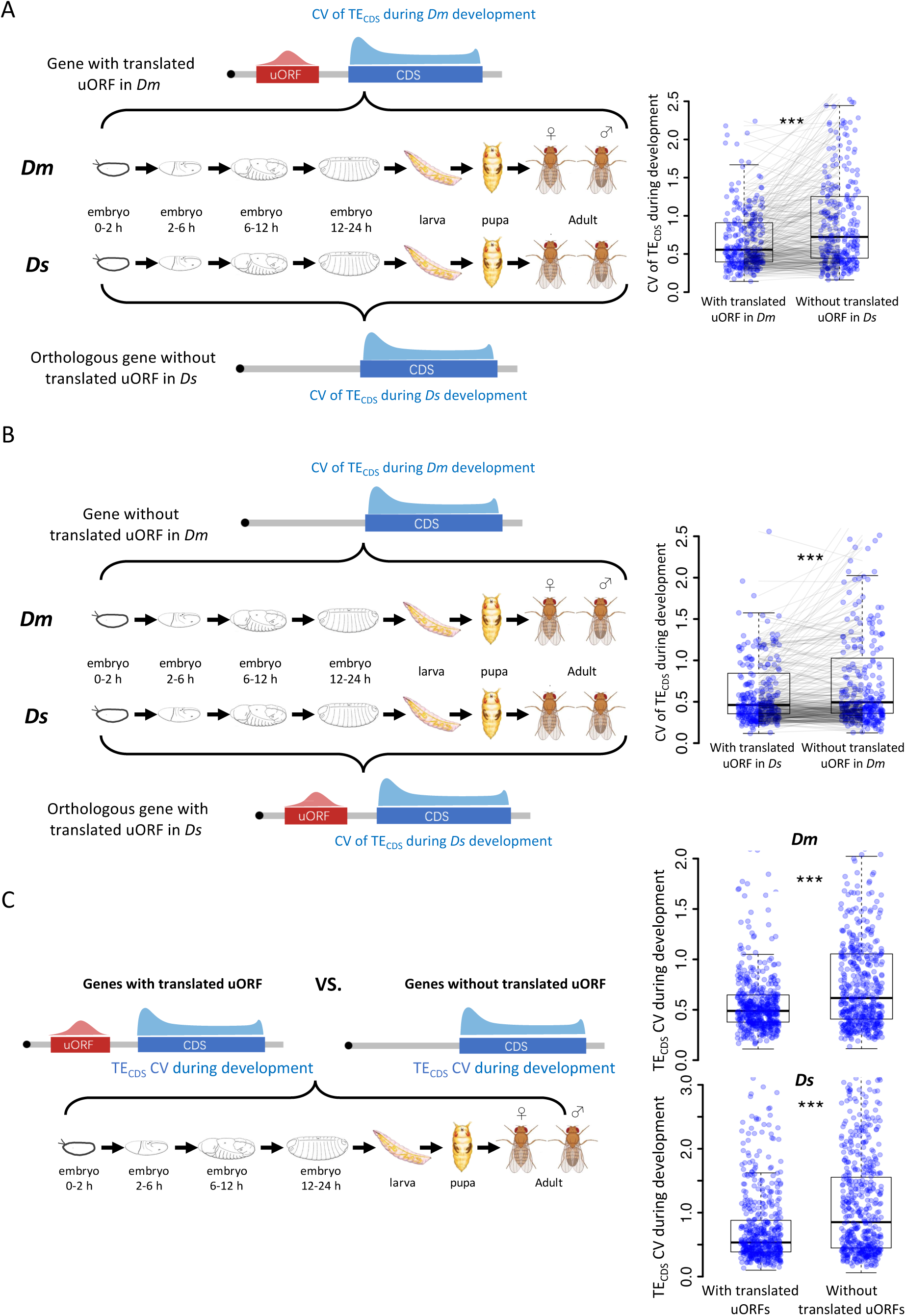
uORFs could reduce CDS translational fluctuation during *Drosophila* development. (A) The CV of TE_CDS_ across 10 *Dm* (*D. melanogaster*) samples and 10 *Ds* (*D. simulans*) samples. The selected gene with uORFs translated (TE > 0.1) in at least one *Dm* sample but its homologous gene without translated uORF in *Ds* samples. Each pair of dots linked by a gray line represents a pair of homologous genes in *Dm* and *Ds*. ***, *P* < 0.001, Wilcoxon signed-rank test. (B) The CV of TE_CDS_ across 10 *Dm* samples and 10 *Ds* samples. The selected gene with uORFs translated (TE > 0.1) in at least one *Ds* sample but its homologous gene without translated uORF in *Dm* samples. Each pair of dots linked by a gray line represents a pair of homologous genes in *Dm* and *Ds*. ***, *P* < 0.001, Wilcoxon signed-rank test. (C) Within each *Drosophila* species, the CV of TE_CDS_ of genes with translated uORFs compared to genes without the translated uORFs. The *P* values are obtained by the Wilcoxon rank-sum test. ***, *P* < 0.001.

These results suggest that uORFs function as translational buffers, reducing gene translation fluctuations during *Drosophila* development.

### Knocking out the uORF of *bcd* increased *bcd* CDS translation in *D. melanogaster*

After verifying the uORFs-mediated translational buffering during *Drosophila* evolution and development, we next aimed to directly explore the biological function of these buffering-capable uORFs in vivo. We first applied stringent criteria to identify uORFs with significant buffering effects on CDS translation between *D. melanogaster* and *D. simulans*. Specifically, we looked for 1) uORFs with significant TE changes (| *log*_2_ (*β*_*u*_)| > 1.5, adjusted *P* < 0.05), 2) negligible changes in its corresponding CDS translation (|*log*_2_(*β*_*C*_)| < 0.05, adjusted *P* > 0.05), and 3) a significant difference between the magnitude of these changes (|log₂(γ)| > 1.5, adjusted *P* < 0.05). We identified 131 uORF-CDS pairs in 103 genes that meet these criteria in at least one stage/tissue (Table S3), with a majority of the genes (67%, 69 out of 103) from embryonic stages (Figure S24A), suggesting a crucial role for uORFs in maintaining translational stability during early development. Among these genes, one notable case is the *bicoid* (*bcd*) gene, a master regulator of anterior-posterior axis patterning during early embryogenesis ^70–72^. The *bicoid* gene contains a 4-codon uORF (excluding the stop codon) in its 5’ UTR, and branch length score (BLS) analysis ^18^ showed that the start codon (uATG) of the *bcd* uORF is highly conserved across the *Drosophila* phylogeny, with a BLS of 0.90 (on a scale from 0 to 1, where higher values indicate greater conservation) (Figure 5A and S24B). Ribo-Seq data revealed that in 0–2 h embryos, the TE of the *bcd* uORF varied more than threefold, while the TE of the CDS was virtually the same between *D. melanogaster* and *D. simulans* (Figure 5B and Table S3). This suggests that the *bcd* uORF significantly buffers translation during early development.

**Figure 5.**
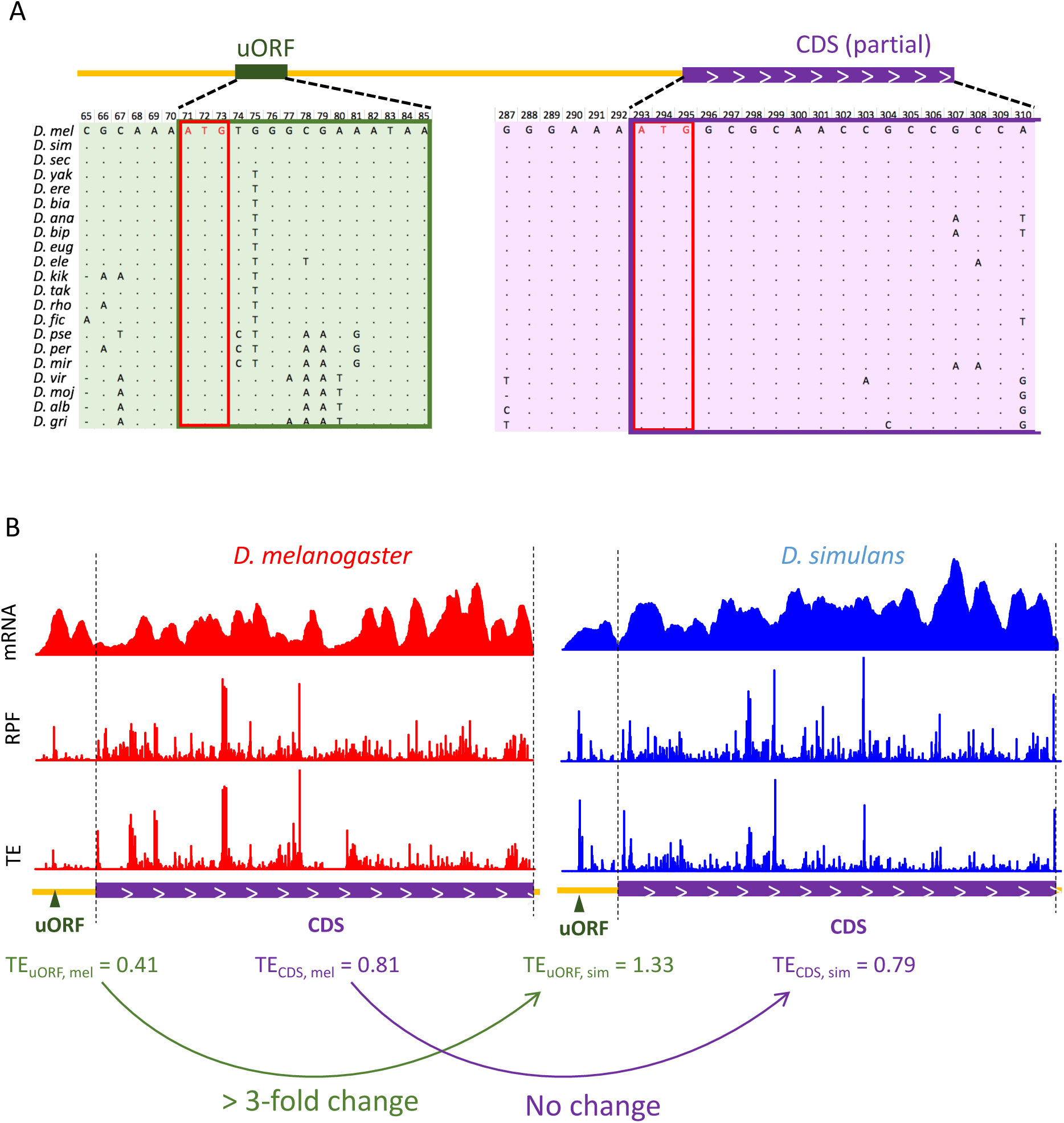
The strong buffering effect of the *bcd* uORF on CDS translation between the two *Drosophila* species. (A) Multiple sequence alignment of the *bcd* uORF and partial CDS in *D. melanogaster* and 20 other *Drosophila* species. The uORF and CDS are boxed in green and purple, respectively. The start codons of the uORF and CDS are boxed in red. (B) The coverage of mRNA-Seq (top), Ribo-Seq (middle), and TEs (bottom) of the *bcd* uORF and CDS in 0-2 h embryos of *D. melanogaster* (red) and *D. simulans* (blue). The uORF and CDS are denoted at the lower panel with dark green triangles and purple boxes, respectively. The 2 dashed lines mark the CDS region. The uORF TE, CDS TE and their interspecific changes were labeled at the bottom.

To investigate the regulatory role of the *bcd* uORF, we used CRISPR-Cas9 to knock out its start codon in *D. melanogaster*, generating two mutant homozygotes (uKO1/uKO1 and uKO2/uKO2) with a genetic background matched to that of the wild-type (WT) (Figures. 6A and S25). To determine whether these mutations enhanced *bcd* CDS translation, we performed ribosome fractionation followed by qPCR ^73–75^, comparing *bcd* mRNA levels in polysome and monosome fractions (P-to-M ratio) from 0–2 h embryos of uKO1/uKO1, uKO2/uKO2, and WT flies (Figure 6B). A larger P-to-M ratio means more mRNAs are enriched in the polysome fractions and bound by more ribosomes, thus indicative of higher translation efficiency. P-to-M ratios were higher in the mutants compared to WT at 29°C (Figure 6C), with a similar but less pronounced trend at 25°C, where the difference between uKO2/uKO2 and WT was not statistically significant (Figure 6C). These findings, along with the known impact of temperature on gene expression and phenotypic plasticity ^76,77^, suggest that the regulatory function of the uORF and overall translation efficiency are temperature-sensitive, highlighting a complex interplay between environmental conditions and gene regulation. To further confirm the *bcd* uORF’s regulatory function, we conducted dual-luciferase reporter assays. The 5’ UTR from *bcd* mutant (uKO1 or uKO2) or WT was cloned into a *Renilla* luciferase reporter construct (Figure 6D). Luciferase activity was significantly higher in uKO1 and uKO2 compared to the WT 5’ UTR, confirming the repressive role of the *bcd* uORF.

**Figure 6.**
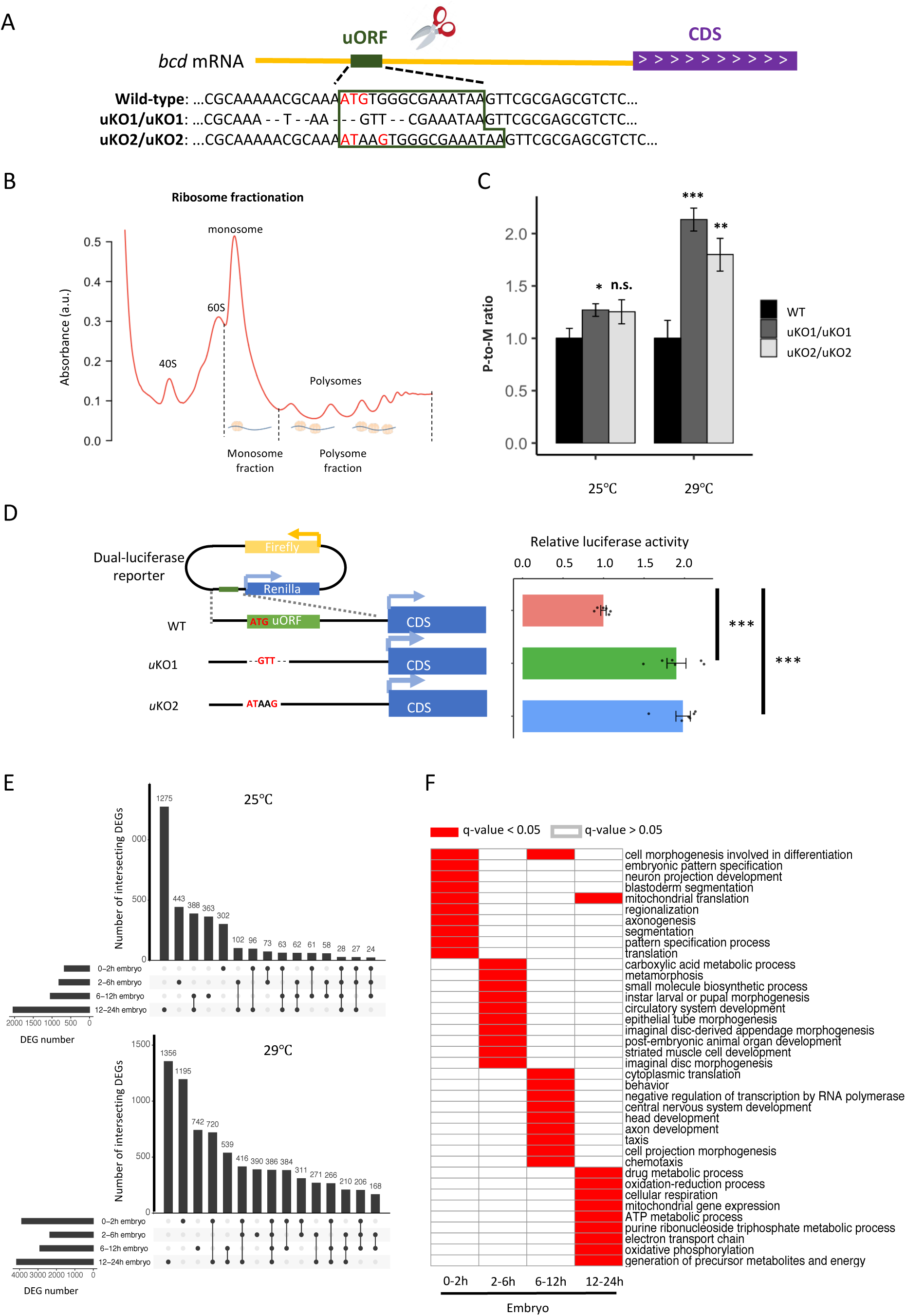
Knocking out the *bcd* uORF increases CDS translation and perturbs the transcriptome during *D. melanogaster* embryogenesis. (A) Genotypes of WT and two uORF knock-out strains (uKO1 and uKO2) generated by CRISPR-Cas9 technology. The uORF is boxed in dark green, and the red ATG represents the start codon of the uORF in the *D. melanogaster* genome. (B) Two ribosome fractions (monosome and polysome) of 0-2 h embryos were separated in a sucrose density gradient. Relative RNA abundance in the monosome and polysome fractions was quantified by real-time quantitative PCR. (C) P-to-M ratio of *bcd* mRNA (*bcd* mRNA abundance in polysome fraction/*bcd* mRNA abundance in monosome fraction) at 25°C (left) and 29°C (right). The P-to-M ratios of mutants were normalized to WT controls at 25°C and at 29°C, respectively. Error bars represent the S.E. of six biological replicates. Asterisks indicate statistical significance (*, *P* < 0.05; **, *P* < 0.01; ***, *P* < 0.001; n.s., *P* > 0.05). (D) Dual-luciferase assay for *bcd* WT uORF and mutated uORF. The reporter structures of the WT and uORF mutants are illustrated on the left. The uORF mutant sequence was the same as that in the fly mutant created with CRISPR-Cas9 technology. The relative activity of *Renilla* luciferase was normalized to that of firefly luciferase. Error bars represent the S.E. of six biological replicates. Asterisks indicate statistical significance (***, *P* < 0.001). (E) The number of DEGs in each stage and their intersection with each other at 25°C (top) and 29°C (bottom). (F) Gene ontology analysis of DEGs at 29°C in each stage. The biological process (BP) terms with q-values < 0.05 in each stage are indicated in red and others are indicated in white.

### *bcd* uORF mutants show wide transcriptomic alteration during *Drosophila* embryogenesis

Since Bcd regulates the expression of many zygotic genes ^70–72^, we anticipated that the increased translation of *bcd* resulting from disrupting the uORF would influence *Drosophila* transcriptomes and phenotypes. To verify this notion, we performed RNA sequencing on embryos from WT and uKO2/uKO2 flies at four developmental stages (0–2 h, 2–6 h, 6–12 h, and 12–24 h) under both 25°C and 29°C conditions, using two biological replicates (Table S4 and Figure S26). Differential expression analysis revealed widespread alterations in gene expression between WT and mutant embryos, with the number of differentially expressed genes (DEGs) increasing over developmental time and at higher temperature (Figure 6E). At 25°C, we identified 674, 817, 1047, and 2,041 DEGs in 0–2 h, 2–6 h, 6–12 h, and 12–24 h embryos, respectively; while at 29°C, we detected 3,884, 2,358, 2,901, and 4,164 DEGs in the corresponding stages (Figure 6E). The majority of DEGs were stage-specific, with only a small fraction consistently differentially expressed across all four stages. Functional enrichment analysis of the DEGs revealed distinct biological pathways affected at each stage, including cell morphogenesis and pattern specification in 0–2 h embryos, metabolic processes and tissue development in 2-6 h and 6-12 h embryos, and mitochondrial respiration in 12–24 h embryos (Figure 6F). Notably, direct targets of Bcd ^78^ were significantly enriched among the DEGs in three out of the four stages, with the exception of 2-6 h embryos (Figure S27). RT-qPCR validation of 20 target genes of Bcd confirmed the reliability of the RNA-seq differential expression analysis (Figure S28). Together, these findings demonstrate that disruption of the *bcd* uORF leads to widespread transcriptional changes during *Drosophila* development, affecting processes ranging from embryogenesis to postembryonic metabolism.

### *bcd* uORF mutants display decreased hatching rates and starvation resistance

Given the widespread transcriptome alterations, we anticipated phenotypic abnormalities in the *bcd* uORF mutants. As expected, both uKO1/uKO1 and uKO2/uKO2 mutants exhibited significantly lower hatching rates compared to WT [*P* = 1.4×10^-6^ and 7.9×10^-6^, respectively, Wilcoxon rank-sum test (WRST); Figure 7A]. At 25°C, uKO1/uKO1 mutants produced fewer offspring than WT flies (*P* < 0.001, WRST, Figure 7B). Given that *bcd* is a maternal gene, we expected reciprocal crosses between uKO1/uKO1 mutants and WT flies to produce different outcomes. Indeed, crossing uKO1/uKO1 males with WT females resulted in offspring numbers similar to those from WT crosses, while crossing uKO1/uKO1 females with WT males yielded offspring numbers comparable to crosses between uKO1/uKO1 mutants (Figure 7B). This verified that the reduction in offspring is due to maternal defects in the uKO1/uKO1 mutants. Similar patterns were observed for uKO2/uKO2 mutants at 25°C (Figure 7B). Notably, the fecundity reduction in uORF-KO mutants was more pronounced at 29°C (Figure 7C). Furthermore, crossing uKO1/uKO1 and uKO2/uKO2 mutants also produced significantly fewer progeny than WT flies (Figure 7C), ruling out genetic background or off-target effects. Collectively, these data demonstrate that disrupting the *bcd* uORF significantly impaired hatchability and fertility.

**Figure 7.**
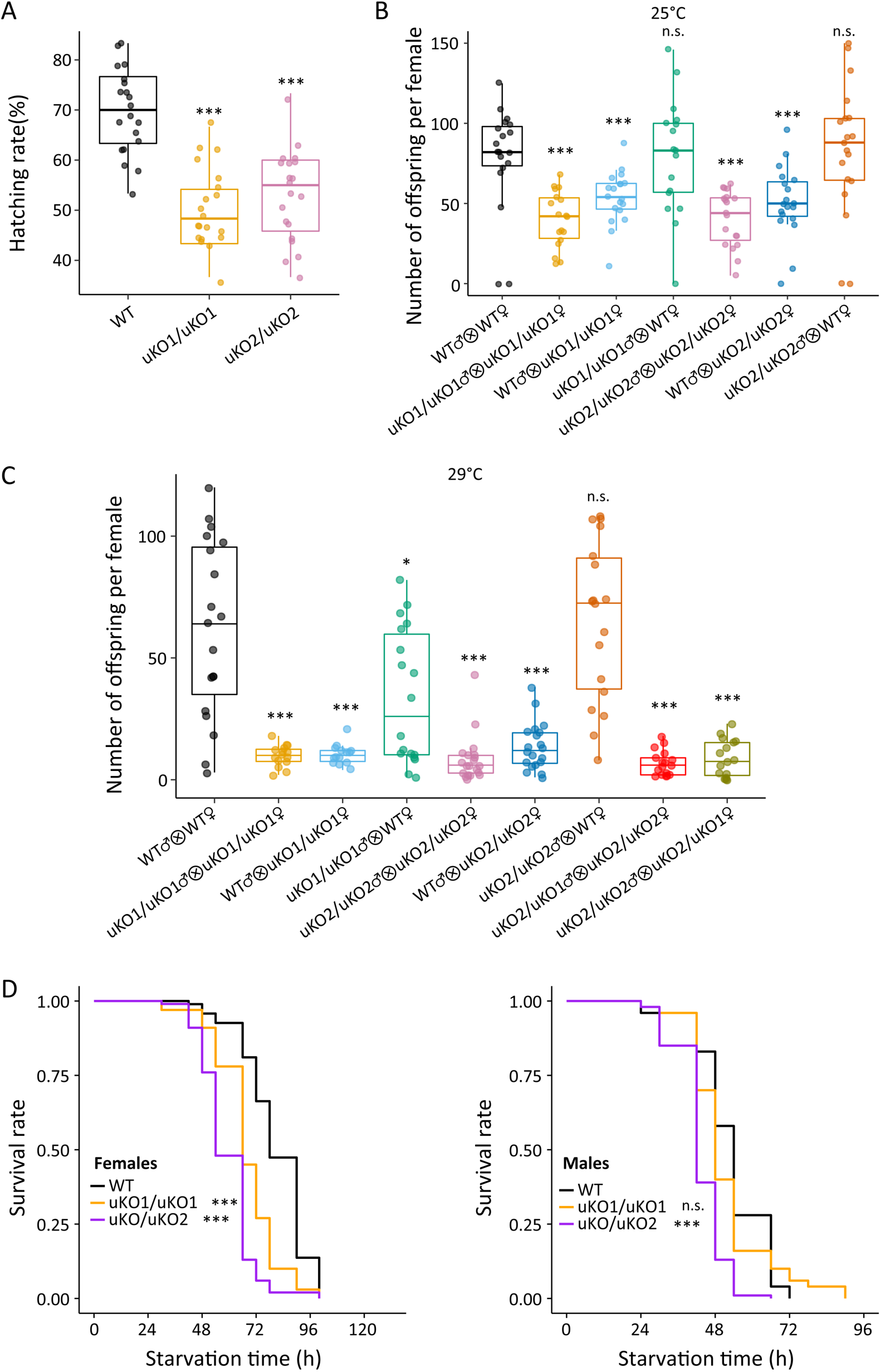
Knockout of the *bcd* uORF reduces offspring number and starvation resistance. (A) Comparison of the hatching rates (%) of mutant and WT offspring (*n* = 20, Wilcoxon rank-sum test; ***, *P* < 0.001). (B) The offspring number per maternal parent in different crosses over 10 days at 25°C. Asterisks indicate significant differences between various crosses and crosses of WT females with WT males (*n* = 20, Wilcoxon rank-sum test; *, *P* < 0.05; **, *P* < 0.01; ***, *P* < 0.001; n.s., *P* > 0.05). The different crosses were denoted as the x-axis labels. (C) The offspring number per maternal parent in different crosses over 10 days at 29°C. (D) Survival curves of WT and mutant adult flies of females (left) and males (right) under starvation conditions. The black line represents the WT, the red line represents the uKO1/uKO1 mutant, and the blue line represents the uKO2/uKO2 mutant. Asterisks indicate significant differences compared to the WT. (*n* = 200, log-rank test; ***, *P* < 0.001; n.s., *P* > 0.05).

We also found that both uKO1/uKO1 and uKO2/uKO2 female mutants perished significantly faster than WT flies under starvation conditions (Figure 7D). Males showed similar tendencies, although the difference was not statistically significant for uKO1/uKO1 mutants (Figure 7D). These data suggest that the knockout of *bcd* uORF diminished starvation resistance in adults, likely due to embryogenesis abnormalities induced by the *bcd* uORF deletion, even in those that successfully developed to adulthood.

### Conservation of uORF-mediated translational buffering in primates

To explore the generality of uORF-mediated translational buffering across evolutionary clades, we analyzed previously published transcriptome and translatome data from three tissues (brain, liver, and testis) in humans and macaques ^79^. We identified 33,680 canonical uORFs in humans and 29,516 in macaques, with 24,385 conserved between the two species. Despite the larger number of uORFs in primates compared to *Drosophila* due to differences in genome size and gene number, the median TE of conserved uORFs was 1.79 times that of non-conserved uORFs in humans, and the corresponding ratio was 3.43 in macaques (Figures 8A and S29). TEs of uORFs were positively correlated between humans and macaques across all tissues (*P* < 0.001, Figure 8B). Additionally, significant positive correlations were observed between the TEs of uORFs and their corresponding coding sequences (CDSs) in all tissues (Figure S30). Although interspecific TE divergence of uORFs (*β*_*u*_) and CDSs (*β*_*C*_) were positively correlated (*P* < 0.001, Figure 8C), uORFs generally exhibited larger divergence (Table S5). Notably, longer uORFs showed stronger buffering effects on CDS translation, reducing interspecific TE divergence by 12.9% and 38.0% compared to genes with short or no uORFs (Figures 8D and S31).

**Figure 8.**
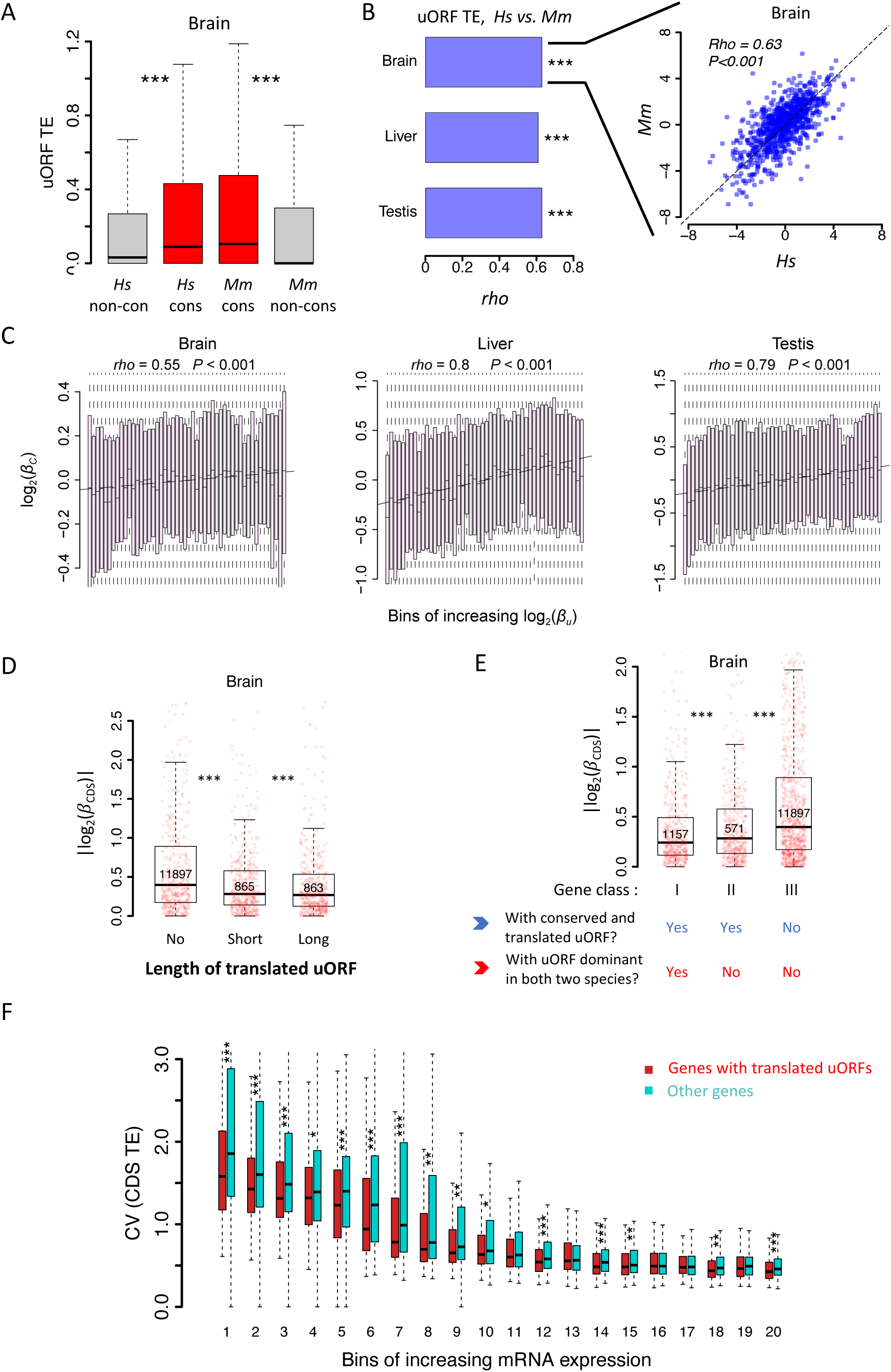
uORFs function as translational buffers in primates. (A) Boxplots showing the TEs of conserved and species-specific uORFs between *Hs* (*H. sapiens*) and *Mm* (*M. mulatta*). Data for the brain is shown as an example. Wilcoxon rank-sum tests. ***, *P* < 0.001. (B) Spearman’s correlation coefficient (*rho*) of uORFs’ TE between humans and macaques. The *rho* values in the brain, liver, and testis were shown as bar plots. ***, *P* < 0.001. Data for the brain is shown as an example in the right panel. (C) Correlation between interspecific uORF TE changes (log_2_*β*_*u*_) and corresponding CDS TE changes (log_2_*β*_*C*_) in three tissues. The x-axis was divided into 50 equal bins with increasing *β*_*u*_. (D) Genes expressed in brains were classified into three classes according to the total length of translated uORFs. Boxplots showing interspecific CDS TE variability |*log*_2_(*β*_*c*_)| of different gene classes. *P* values were calculated using Wilcoxon rank-sum tests between the neighboring groups. ***, *P* < 0.001. (E) Genes expressed in brains (mRNA RPKM > 0.1 in both species) were classified into three classes according to whether a gene had a conserved and dominantly translated uORF (TE > 0.1) in both species or not. Boxplots showing interspecific CDS TE variability |*log*_2_(*β*_*c*_)| of different gene classes. *P* values were calculated using Wilcoxon-rank sum tests between the neighboring groups. ***, *P* < 0.001. (F) Boxplot showing the coefficients of variation (CVs) of CDS TE among the 69 lymphoblastoid cell lines (LCLs). Expressed genes (mean mRNA RPKM > 0.1) were divided into 20 bins with increased mRNA expression levels. In each bin, the genes were divided into two fractions according to whether the gene had a translated uORF or not. Wilcoxon rank-sum tests. *, *P* < 0.05; **, *P* < 0.01; ***, *P* < 0.001.

As in *Drosophila*, we categorized expressed human genes into three classes based on uORF conservation and translation: Class I, genes with conserved and dominantly translated uORFs in both humans and macaques; Class II, genes with conserved uORFs translated in both species but not dominantly in at least one; and Class III, the remaining expressed genes (Figures 8E and S32). Consistent with findings in *Drosophila*, significant differences in |*β*_*c*_| between humans and macaques were observed in the order of Class I < II < III, with Class I genes showing 9.8% and 17.1% lower |*β*_*c*_| values compared to Class II and Class III genes, respectively (Figures 8E and S32). These similarities between primates and *Drosophila*—two clades that diverged over 700 million years ago ^80^ —suggest that uORF-mediated translational buffering is a widespread mechanism for stabilizing gene translation across evolutionary clades.

We also analyzed matched mRNA-Seq and Ribo-Seq data from 69 human lymphoblastoid cell lines ^81,82^ to test whether uORFs buffer against translational variability across different individuals within humans. Genes with translated uORFs exhibited, on average, 8.65% lower CVs in CDS TE across individuals than genes without translated uORFs (Figure 8F), with longer uORFs showing stronger buffering effects (Figure S33). Collectively, these findings suggest that uORFs play a crucial role in reducing translational variability both across evolutionary clades and within species.

## Discussion

Translational control is vital for maintaining protein homeostasis and cellular activities ^83,84^. However, the mechanisms underlying the conservation of protein abundance across species remain largely unknown ^5,6^. In this study, we extended the ICIER model and conducted simulations to explore uORFs’ regulatory roles in buffering translation during evolution. Our simulations demonstrated that uORF translation reduces variability in downstream CDS translation, with buffering capacity positively correlated with uORF translation efficiency, length, and number. Comparative translatome analyses across developmental stages of two *Drosophila* species provided evidence that uORFs mitigate interspecific differences in CDS translation. Similar patterns were observed between humans and macaques, and between human individuals, suggesting that uORF-mediated buffering is an evolutionarily conserved mechanism in animals. Additionally, in vivo experiments showed that knocking out the *bcd* uORF led to aberrant embryogenesis and altered starvation resistance, with more pronounced phenotypic effects at higher temperatures, supporting the role of uORFs in fine-tuning translation and phenotypic outcomes.

While the prevailing consensus on uORFs’ functions is that they repress the translation of downstream CDS by sequestering ribosomes ^34,43–48^, recent studies have suggested uORFs might play essential roles in stabilizing protein expression ^43,55,56^. While these studies have primarily focused on the role of uORFs in single-gene cases or under stress conditions, our work expands this knowledge by demonstrating that uORFs act as a general mechanism for buffering translational variability on a genome-wide scale across multiple species and developmental stages. Our findings also suggest that uORF-mediated translational buffering is not merely a response to environmental stress but an evolutionarily conserved mechanism integral to maintaining protein homeostasis across species. Organisms can buffer phenotypic variation against environmental or genotypic perturbations through “canalization” ^85^. The discovery that uORFs are crucial for *Drosophila* development links them to canalization, providing evidence that uORFs reduce interspecific CDS translation differences and enhance phenotypic stability. While previous studies primarily focused on the stabilization of transcriptional levels ^86–90^ or protein levels ^55,88,91–93^, our study opens new avenues for exploring how organisms maintain phenotypic robustness at the translational level through molecular mechanisms like uORF-mediated buffering. This study lays the groundwork for future research into the evolutionary and functional roles of uORFs, particularly in the context of canalization and adaptive evolution.

Previous studies have shown that a significant fraction of fixed uORFs in the populations of *D. melanogaster* and humans were driven by positive Darwinian selection ^63,67^, suggesting active maintenance through adaptive evolution rather than purely neutral or deleterious processes. While uORFs have traditionally been recognized for their capacity to attenuate translation of downstream CDSs, accumulating evidence now underscores their critical role in stabilizing gene expression under fluctuating cellular and environmental conditions ^43,55,56^. Whether the favored evolutionary selection of uORFs acts primarily through their role in translational repression or translational buffering remains a compelling yet unresolved question, as these two functions are inherently linked. Indeed, highly conserved uORFs tend to be translated at higher levels, resulting not only in stronger inhibition of CDS translation ^34,45,67^, but also in a more pronounced buffering effect, as demonstrated in this study. This buffering capacity of uORFs potentially provides selective advantages by reducing fluctuations in protein synthesis, thus minimizing gene-expression noise and enhancing cellular homeostasis. This suggests that selection may favor uORFs that contribute to translational robustness, a hypothesis supported by findings in yeast and mammals showing that uORFs are significantly enriched in stress-response genes and control the translation of certain master regulators of stress responses ^41,42,94,95^. Our study suggests that translational buffering, rather than translational repression alone, can also drive evolutionary selection favoring uORFs, although it remains challenging to empirically disentangle these functions. Future comparative genomic analyses, coupled with experimental approaches such as ribosome profiling and functional mutagenesis, will be crucial in elucidating the precise evolutionary forces driving uORF conservation and adaptation.

The original ICIER model was devised to elucidate how a single uORF can confer resistance to global translation inhibition on its host gene during stress conditions ^43^. It proposed that an elongating ribosome (80S) causes downstream 40S subunits to dissociate from the mRNA after collision. Our revised model expands this scope to include bidirectional dissociation, where an 80S ribosome can release both upstream and downstream 40S subunits, reflecting recent findings that 80S primarily causes upstream 40S dissociation during 40S/80S collisions ^56^. Mechanistically, increasing the loading rate of 40S ribosomes onto the mRNA (*R_in_*) elevates the density of 40S and 80S ribosomes scanning the uORF, thereby raising the likelihood of 40S-80S collisions and subsequent 40S dissociation. This leads to a reduced flow of 40S ribosomes reaching the CDS relative to the initial increase in 40S at the 5’ end of the mRNA and uORF. Conversely, a decrease in 40S ribosome loading rate reduces the density of scanning ribosomes, diminishing the probability of 40S-80S collisions and 40S dissociation, and partially restoring the flow of 40S ribosomes to the CDS. Overall, the 40S-80S collision mechanism and subsequent 40S dissociation at the uORF moderate downstream CDS translation less than changes in ribosome loading at the 5’ end of the mRNA or uORF. The revised model underscores the analogy of uORFs as “molecular dams”, adeptly modulating the stream of ribosomes and cushioning downstream CDS translation against fluctuations (Figure 9A). Additionally, our study broadens its perspective to account for the potential interactive effects of multiple uORFs within a gene, recognizing the prevalent clustering of these regulatory elements and their combinatorial influence on translational control. In essence, our study presents a nuanced and comprehensive expansion of the ICIER model, paving the way for a deeper understanding of uORFs as pivotal elements in the maintenance of protein synthesis stability under variable translational conditions. However, we realize the existence of alternative mechanisms governing uORF function, such as those found in traditional repression models ^55^ or ribosome queuing and reinitiation strategies ^56^. As such, further computational studies are needed to dissect and understand the full spectrum of uORF-mediated translational regulation.

**Figure 9.**
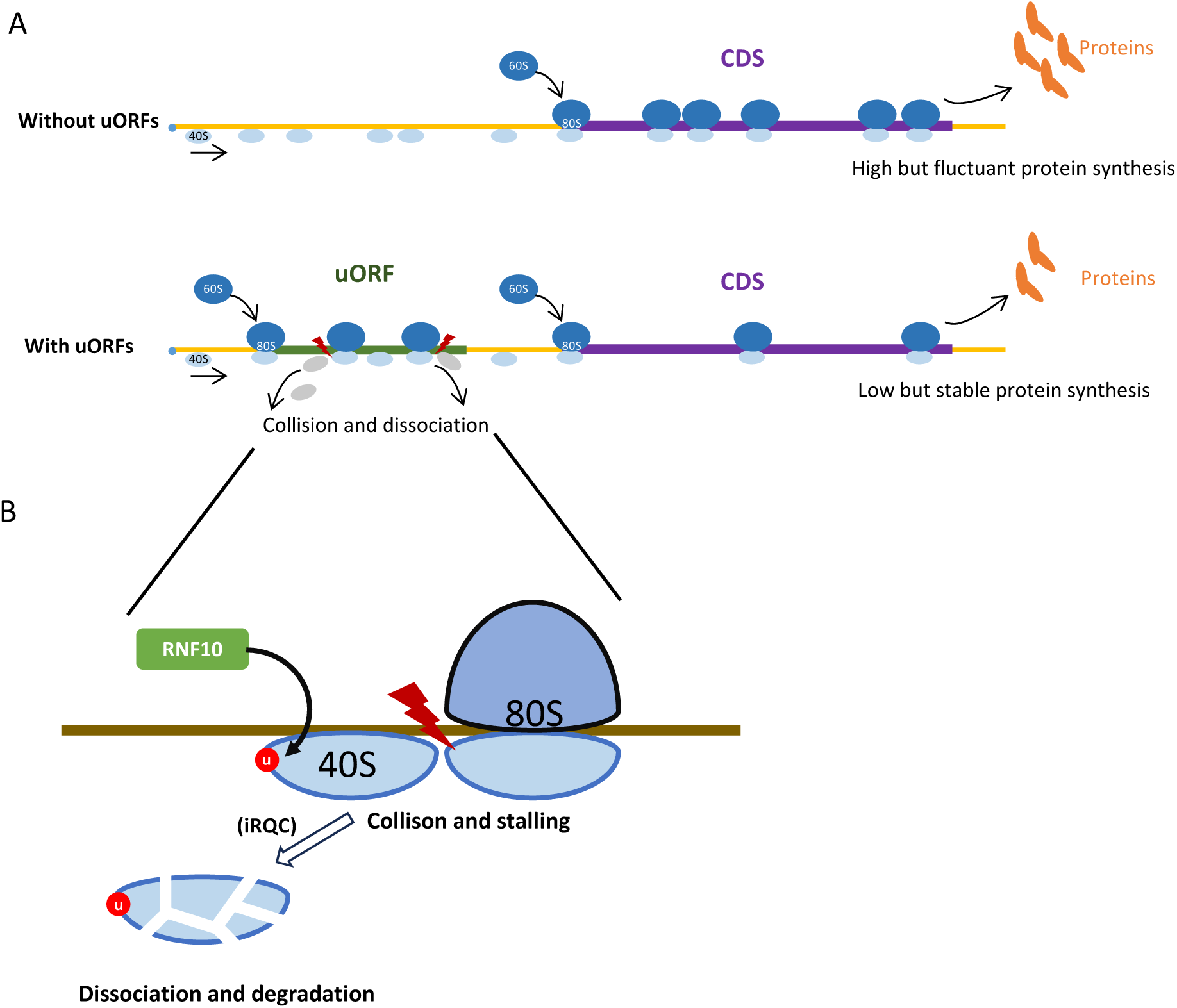
Model illustrating the uORF-mediated translational buffering. (A) uORFs can reduce and stabilize the downstream CDS translation through “ribosomes collision and dissociation” mechanism in an mRNA containing uORFs. In contrast, the CDS translation is higher and more fluctuant in an mRNA without uORFs. (B) The collided and stalled 40S subunit dissociates from mRNA, is ubiquitinated by the E3 ubiquitin ligase RNF10 and degraded by the initiation ribosome quality control (iRQC) pathway.

Ribosome slowdown or stalling on mRNA due to rare codons ^56,96–98^ or nascent blocking peptides ^99–102^ frequently triggers ribosome collisions genome-wide ^103–105^. Such collisions, especially among elongating 80S ribosomes, often activate ribosome quality control (RQC) pathways that recognize collision interfaces on the 40S subunit, leading to ribosomal subunit dissociation and degradation ^106–108^. In mammals, ZNF598 specifically identifies collided ribosomes to initiate ubiquitin-dependent protein and mRNA quality control pathways ^109–113^. Analogously, yeast employs Hel2-mediated ubiquitination of uS10, initiating dissociation via the RQC-trigger complex (RQT) ^114^. Furthermore, the human RQT (hRQT) complex recognizes ubiquitinated ribosomes and induces subunit dissociation similarly to yeast RQT ^115^. However, transient ribosome collisions can evade RQC by promoting resumed elongation through mechanical force provided by trailing ribosomes, thereby mitigating stalling ^116^. Beyond 80S collisions, evidence increasingly highlights a distinct collision type involving scanning 40S subunits or pre-initiation (43S) complexes. Recently, an initiation RQC pathway (iRQC) targeting the small ribosomal subunit (40S) has been described, particularly involving collisions between scanning 43S complexes or between stalled 43S and elongating 80S ribosomes (Figure 9B) ^117,118^. During iRQC, E3 ubiquitin ligase RNF10 ubiquitinates uS3 and uS5 proteins, resulting in 40S degradation ^118^. This mechanism aligns closely with our ICIER model, proposing collision-driven 43S dissociation in the 5’ UTRs. Future studies exploring these mechanisms in greater detail will clarify how uORFs modulate translational regulation through buffering effects.

The *bcd* gene, crucial for *Drosophila* embryogenesis, has been well studied ^71,72,119–123^, but its uORF remains underexplored. Our study showed that deleting the *bcd* uORF led to abnormal phenotypes and reduced fitness, underscoring the importance of uORF-mediated translation. These effects were more severe under heat stress (29°C) than at normal temperatures (25°C), suggesting uORFs play a vital role in buffering phenotypic variation. We demonstrated that the *bcd* uORF represses CDS translation using sucrose gradient fractionation followed by qPCR—an approach that directly measures translation efficiency while minimizing confounding from RNA/protein degradation. However, detecting Bcd protein levels with antibodies across developmental stages or conditions in the mutants and wild-type controls would provide an even stronger validation of our model and should be explored in future studies. Rescue experiments, typically used to confirm that phenotypic changes are due to specific genetic edits, faced challenges in our study and previous uORF-KO experiments in animals ^124,125^ and plants ^126–129^. We altered the start codon of the *bcd* uORF to minimize effects on the 5’ UTR, without impacting the coding sequence, making traditional rescue experiments impractical. To control for genetic background variations, we backcrossed two *bcd* uORF-KO mutant strains with *w*^1118^ flies for nine generations, using *w*^1118^ as a control. Both uORF-KO mutants (uKO1/uKO1 and uKO2/uKO2) consistently showed reduced progeny, ruling out background or off-target effects. Crosses between the two mutants produced fewer offspring than WT crosses, confirming that the reduced progeny was due to the uORF deletions, not genetic background. As *bcd* is maternally inherited, we expected defective phenotypes only in offspring from mutant females crossed with wild-type males. This was confirmed: uKO1/uKO1 males crossed with wild-type females had normal offspring, while uKO1/uKO1 females crossed with wild-type males had reduced offspring, similar to mutant crosses. Similar results were observed for uKO2/uKO2 mutants. These findings confirm that disrupting *bcd* uORFs significantly reduces hatchability and fertility, effects not attributable to genetic background. Our results highlight the critical role of *bcd* uORFs in development and fecundity, and provide a basis for future studies on uORF regulation in other genes across species.

Taken together, this study demonstrates the role of uORF-mediated translational buffering in mitigating variability in gene translation during species divergence and development, using large-scale comparative transcriptome and translatome data. Our work addresses gaps in understanding the stabilization of gene expression at the translational level and offers new insights into the evolutionary and functional significance of uORFs.

## Materials and Methods

### Modeling the uORF-mediated buffering effect on CDS translation

We adapted the stochastic ICIER framework developed by Andreev *et al.* ^43^ based on the TASEP model, with several major modifications. First, we concatenated a 500-codon CDS downstream of the uORF. A leaky scanning ribosome from the uORF would initiate translation at the CDS start codon with a probability of *I*_*CCS*_. The probability of elongating ribosomes moving along the CDS (*v*_*EC*_= 0.5) in each action was higher than that along uORFs (*v*_*Eu*_= 0.3) in the model settings, considering that uORFs usually encode blocking peptides ^99–102,130^ or contain stalling codons ^56,96,97^. Second, we recorded the number of elongating ribosomes that completed translation at the stop codon of a CDS (*N*_*EC*_) or uORF (*N*_*Eu*_) during a given time period and regarded them as proxies for quantifying the CDS or uORF TE. Third, we considered three models for simulating the consequences of scanning ribosomes colliding with elongating ribosomes mentioned above: a downstream dissociation model, an upstream dissociation model, and a double dissociation model.

In detail, the mRNA molecule structure in the simulations consisted of five parts: a 5’ leader before the uORF (150 nucleotides), the uORF (ranging from 2 to 100 codons), a segment between the uORF and CDS (150 nucleotides), the CDS (500 codons) and the 3’ UTR (150 nucleotides). The *R*_*in*_ determined the loading rate of the 40S scanning ribosomes on the mRNA 5’-terminus, and it roughly represented the availability of *trans* translational resources in the cell. We generated two sets of *R_in_* values (Figure S1A) to simulate the variation in the availability of translational resources (40S scanning ribosomes, etc.): i) 1000 values of *R_in_* were generated from a random generator of a uniform distribution, *U* (0, 0.1); ii) 1000 values were first generated from a random generator of an exponential distribution, *E* (1), and then each of the 1000 values was divided by 70 to obtain 1000 values of *R_in_*. *I*_*uORF*_ and *I*_*CCS*_ represent the probability of translation at the start codon of a uORF or CDS, respectively. We used different combinations of *I*_*uORF*_ and *I*_*CCS*_ to test how translational initiation strength influenced the buffering effect of uORFs. *v*_*s*_ determined the scanning rate of the 40S ribosome, and *v*_*Eu*_and *v*_*EC*_ determined the elongation rate of the 80S ribosome at a uORF or CDS, respectively. All the parameters used in our simulation are listed in Table S1.

The mRNA molecule was modeled as an array, and the value of each position in the array represented the occupation status of ribosomes along the mRNA molecule. The process of translation was simulated by a series of discrete actions referred to Andreev *et al.* ^43^. Upon each action, there was a probability that certain events would occur, including the addition of a 40S ribosome to the 5’-terminus, the transformation of a 40S ribosome into an 80S ribosome at a CDS or uORF start codon, the movement of a 40S ribosome or 80S ribosome to the next codon in the CDS or uORF, etc. More specific operations in each action were described in Supplementary Materials and Methods.

After 1,000,000 actions, we recorded the number of 80S ribosomes that completed translation at the uORF (*N*_*Eu*_) and CDS (*N*_*EC*_) for each *R*_*in*_ input, which was used to represent the protein production rates (i.e. translation rate) of the uORF and CDS, respectively. To test the effects of different factors, we used various combinations of the parameters in Table S1 in the simulations. For a given distribution of *R*_*in*_ input (1000 values) and the dissociation model, we obtained the corresponding 1000 *N*_*EC*_ values and their CV values by calculating the ratio of the standard deviation to the mean of *N*_*EC*_.

In the double-uORF simulation, we only adopted a uniform distribution of *R*_*in*_ input and the downstream dissociation model. Most parameters and simulation processes were similar to those in the single-uORF simulation, except that an additional uORF was introduced upstream of the CDS with an initiation probability of *I*_*uORF*2_.

### Annotation of uORFs in *D. melanogaster* and *D. simulans*

We downloaded the whole-genome sequence alignment (maf) of *D. melanogaster* (dm6) and 26 other insect species from the University of California Santa Cruz (UCSC) genome browser (<u>genome.ucsc.edu</u>)^131^. To improve the genome correspondence between *D. melanogaster* and *D. simulans*, we adopted a newly published reference-quality genome of *D. simulans* (NCBI, ASM438218v1) ^132^ to replace the original *D. simulans* genome in UCSC maf. We softly masked repetitive sequences in the new genome by RepeatMasker 4.1.1 (http://www.repeatmasker.org) and aligned it to dm6 with lastz ^133^ in runLastzChain.sh following UCSC guidelines. The lastz alignment parameters and the scoring matrix were the same as the parameters of dm6 and droSim1 in UCSC. Chained alignments were processed into nets by the chainNet and netSyntenic programs. The alignment was integrated into the multiple alignments of 27 species with multiz ^134^.

Based on the genome annotation of *D. melanogaster* (FlyBase r6.04, https://flybase.org/), we used the Galaxy platform to parse the multiple sequence alignments of 5’ UTRs in *Drosophila*. The 5’ UTR sequences of each annotated transcript from *D. melanogaster* and the corresponding sequences in *D. simulans* were extracted from the maf. The start codons of putative uORFs were identified by scanning all the ATG triplets (uATGs) within the 5’ UTRs of *D. melanogaster* and *D. simulans*. uATGs that overlapped with any annotated CDS region were removed. The presence or absence of each uATG of *D. melanogaster* was determined at orthologous sites in *D. simulans* based on multiple genome alignments, and vice versa. The conserved uORF was defined as a uORF where its uATG was present in both *D. melanogaster* and the corresponding orthologous positions of *D. simulans*. For each protein-coding gene, we only considered the canonical transcript. For transcripts containing multiple uORFs, we defined the uORF that showed the highest TE as the dominant uORF. The branch length score (BLS) of the uATGs was calculated as we previously described ^18^.

### Fly materials and general raising conditions

The *sim4* strain of *D. simulans* was used to generate all the libraries of *D. simulans* in this study. All flies were raised on standard corn medium and grown in 12 h light: 12 h dark cycles at 25 °C for general conditions or at 29 °C for specific experimental design. The samples of embryos at different stages, larva, pupa, bodies and heads were collected following a previous protocol ^23^.

### Processing *Drosophila* mRNA-Seq and Ribo-Seq data

Ribo-Seq and matched mRNA-Seq libraries for different developmental stages and tissues of *D. simulans* were constructed as we previously described ^23^ and sequenced on an Illumina HiSeq-2500 sequencer (run type: single-end; read length: 50 nucleotides) according to the manufacturer’s protocol. The 3’ adaptor sequences (TGGAATTCTCGGGTGCCAAGG) were trimmed using Cutadapt 3.0 with default parameters ^135^, and the NGS reads were mapped to the genomes of yeast, *Wolbachia*, *Drosophila* viruses and the sequences of tRNAs, ribosomal RNAs (rRNAs), small nuclear RNAs (snRNAs) or small nucleolar RNAs (snoRNAs) of *D. melanogaster* (FlyBase r6.04) and *D. simulans* (FlyBase r1.3) using Bowtie2 version 2.2.3 ^136^ with the parameters -p8 --local -k1. The mapped reads of these genomes/sequences were further removed in the downstream analysis.

After filtering, the mRNA-Seq and Ribo-Seq reads were mapped to the reference genomes of *D. melanogaster* (FlyBase, r6.04) and *D. simulans* (NCBI, ASM438218v1), respectively, using the Spliced Transcripts Alignment to a Reference (STAR) algorithm ^137^. For Ribo-Seq reads, we assigned a mapped RPF (27-34 nucleotides in length) to its P-site using the psite script from Plastid ^138^. The uniquely mapped reads were extracted and then mapped to the CDS of *D. melanogaster* and *D. simulans* respectively using STAR ^137^. The P-sites of RPF or mRNA-Seq reads that overlapped with a CDS were counted separately. The reads that were not mapped to CDSs were then mapped to the 5’ UTRs of *D. melanogaster* and *D. simulans*. The P-sites of RPF or mRNA-Seq reads that overlapped with uORFs were counted separately.

The TE of a given uORF or CDS was calculated as the RPKM_P-site_/RPKM_mRNA_ ratio. For a few uORFs with RPKM_mRNA_ = 0 but RPKM_P-site_ > 0, a pseudocount of 0.1 was added to both RPKM_mRNA_ and RPKM_P-site_ to avoid dividing a positive value by zero. In each sample, expressed genes were defined as the genes with a CDS RPKM_mRNA_ > 0.1 in both species, and translated uORFs were defined as uORFs with a TE > 0.1.

### Testing the statistical significance of the difference in the interspecific TE change between a uORF and its downstream CDS

For this analysis, we adopted the methods developed by Zhang *et al*. ^23^. Briefly, for the samples of female bodies or male bodies of *D. melanogaster* with biological replicates, we obtained the log_2_(TE) and SE of the log_2_(TE) of a uORF or CDS by contrasting the RPF counts against mRNA-Seq read counts using DESeq2. Then, we fitted the SE values against the normalized mRNA counts and log_2_(TE) values using the gam function in the R package mgcv, with a log link to obtain the SE ∼ mRNA counts + log_2_TE functions. For other samples without biological replicates, we estimated the SE of the log_2_(TE) for a feature (CDS or uORF) by applying the fitted functions obtained based on the biological replicates of female and male bodies to the observed mRNA counts and log_2_(TE) values. We identified uORFs whose TEs differed significantly between the paired samples of *D. melanogaster* and *D. simulans* by testing whether the value obtained for *log*_2_ (*β*_*u*_) = *log*_2_ (*TE*_*uORF*,*sim*_) – *log*_2_ (*TE*_*uORF*,*mel*_) was significantly different from 0. Based on the SE of the log_2_(TE) derived as described above, the SE of the log_2_(*β*_*u*_) can be derived as follows:

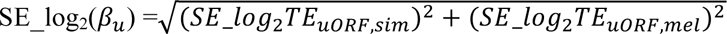

As the Wald statistic, 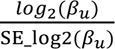, follows a standard normal distribution under the null hypothesis of log (*β*_*u*_) = 0, we calculated the *P* value as follows: 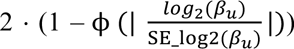. The identification of CDSs whose TEs differed significantly between the paired samples of *D. melanogaster* and *D. simulans* was similar to the procedures above.

Then, we defined 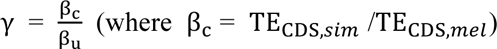 and tested whether log_2_ (γ) was significantly different from 0 to determine whether the magnitude of interspecific TE changes in CDSs and uORFs was significantly different. We obtained log_2_TE_uORF,*sim*_, log_2_TE_uORF,*mel*_, log_2_TE_CDS,*sim*_, and log_2_TE_CDS,*mel*_ as described above and estimated SE_log_2_TE_uORF,*sim*_, SE_log_2_TE_uORF,*mel*_, SE_log_2_TE_CDS,*sim*_, and SE_log_2_TE_CDS,*mel*_ based on the biological replicates of female and male bodies of *D. melanogaster*. Finally, log_2_(γ) also follows a normal distribution with SE denoted as SE_log_2_(*γ*) =

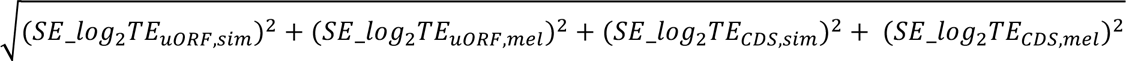

As the Wald statistic, 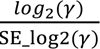, follows a standard normal distribution under the null hypothesis that log_2_ (*β*_*u*_) = 0, we calculated the *P* value as follows: 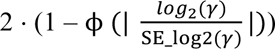.

uORFs are shorter than CDSs and have substantially lower read counts, reducing statistical power when comparing TE changes. To mitigate this bias, we focused on highly expressed genes, defined as those with both mRNA and RPF RPKM values above the 50th percentile in *D. melanogaster* and *D. simulans*. uORFs and CDSs of these genes were used in the calculation of significant TE change.

### Knocking out a uORF with CRISPR-Cas9 technology in *D. melanogaster*

We searched for possible sgRNA target sites near the uATG start codon of a uORF using the Benchling website (https://www.benchling.com/crispr/) to design optimal single guide RNA (sgRNA) sequences with high specificity and low off-target effects. We then synthesized single-stranded complementary DNAs (ssDNAs) and annealed them to obtain double-stranded DNA (dsDNA), which served as the template for sgRNA expression. The template sequences of the sgRNA used for *bcd* uATG-KO are listed in Table S6. The dsDNA was then ligated into the BbsI-digested pU6B vector. The pU6B-sgRNA plasmid was purified and injected into the embryos of transgenic Cas9 flies collected within one hour of laying at the Tsinghua Fly Center as described in Ni *et al.* ^139^. The injected embryos were kept at 25°C and 60% humidity until adulthood (G0). The G0 adult flies that hatched from injected embryos were individually crossed with other flies (*y sc v*) to increase the number of offspring. Then, the F1 progeny were crossed with flies carrying an appropriate balancer (*Dr, e/TM3, Sb*). After F2 spawning, the F1 individuals were screened for mutations of interest by genotyping. The primers used for genotyping are listed in Table S6. The F2 progeny whose parents showed positive genotyping results were then screened for the *yellow*^-^ gene to separate the chromosome carrying *nos-Cas9*. The screened F2 males were crossed with the flies containing the same balancer as above. After F2 genotyping, the progeny (F3) of positive F2 individuals showing the same mutation status were crossed individually to generate homozygous mutants in the F4 generation. The original homozygous mutants were sequentially outcrossed with *w*^1118^ flies for 9 generations to purify the genetic background (Figure S25B).

### Ribosome fraction analysis by sucrose gradient fractionation and RT‒qPCR

The 0-2 h embryos obtained from *w*^1118^, uKO1/uKO1, and uKO2/uKO2 mutants raised at 25°C and 29°C were collected and homogenized in a Dounce homogenizer with lysis buffer [50 mM Tris pH 7.5, 150 mM NaCl, 5 mM MgCl_2_, 1% Triton X-100, 2 mM dithiothreitol (DTT), 20 U/ml SuperaseIn (Ambion), 0.5 tablets of proteinase inhibitor (Roche), 100 µg/ml emetine (Sigma Aldrich), and 50 µM guanosine 5′-[β,γ-imido]triphosphate trisodium salt hydrate (GMP-PNP) (Sigma Aldrich)] at 4°C. The lysates were clarified by centrifugation at 4°C and 20,000×g for 8 min, and the supernatants were transferred to new 1.5 ml tubes. 10-45% sucrose gradients were prepared in buffer (250 mM NaCl, 50 mM Tris pH 7.5, 15 mM MgCl_2_, 0.5 mM DTT, 12 U/ml RNaseOUT, 0.5 tablets of protease inhibitor, and 20 µg/ml emetine) using a Gradient Master (Biocomp Instruments) in ULTRA-CLEAR Thinwall Tubes (Beckman Coulter). A sample volume of up to 500 µl was applied to the top of each gradient. After ultracentrifugation with a Hitachi P40ST rotor at 35,000 × rpm for 3 h at 4°C, the monosome and polysome fractions were collected, flash-frozen in liquid nitrogen, and stored at −80°C until further use.

The RNA in the monosome and polysome fractions was extracted separately using TRIzol reagent (Life Technologies, Inc.) and chloroform (Beijing Chemical Works) following the manufacturer’s instructions and were reverse transcribed into cDNA using the PrimeScript™ II 1st Strand cDNA Synthesis Kit (Takara). RT‒qPCR analysis of *bcd* cDNA and its targets was performed using PowerUp™ SYBR™ Green Master Mix (Thermo Fisher) following the manufacturer’s instructions with *rp49* as an internal control. The primer sequences employed for RT-qPCR are listed in Table S6. For each sample, the ratio of *bcd* mRNA abundance in the polysome fraction to that in the monosome fraction was calculated as the P-to-M ratio. Six biological replicates were performed for each sample.

### Dual-luciferase reporter assays

The wild-type (WT) 5’ UTR of *bcd* was cloned from cDNA by PCR, and uATG mutations were introduced into 5’ UTR using specific amplification primers (listed in Table S6). The WT and mutated 5’ UTR sequences were ligated into a linearized reporter plasmid (psiCHECK-2 vector, Promega). The whole sequence of all the plasmids was validated by Sanger sequencing.

*Drosophila* S2 cells were cultured in Schneider’s Insect Medium (Sigma) plus 10% (by volume) heat-inactivated fetal bovine serum, 100 U/ml penicillin and 100 µg/ml streptomycin (Thermo Fisher) at 25°C without CO_2_ for 24 h to reach 1–2×10^6^cells/ml before further treatments. Plasmid transfection was conducted with Lipofectamine 3000 (L3000001, Thermo Fisher) according to the supplier’s protocol.

The *Renilla* luciferase activity associated with WT or uORF-mutated 5’ UTRs was measured according to the manual of the Dual-Luciferase Reporter Assay System (Promega) 32 h after transfection and was normalized to the activity of firefly luciferase.

### mRNA-Seq in the embryos of mutant and WT flies

We collected 0-2 h, 2-6 h, 6-12 h, and 12-24 h embryos of *w*^1118^ and uKO2/uKO2 mutants raised at 25°C and 29°C and conducted mRNA-Seq. Library construction and sequencing with PE150 were conducted by Annoroad on the Illumina Nova6000 platform. Two biological replicates were sequenced for each sample. The clean data were mapped to the reference genome of *D. melanogaster* (FlyBase, r6.04) using STAR ^137^. Reads mapped to the exons of each gene were tabulated with htseq-count ^140^. The differentially expressed genes were identified using DESeq2 ^141^. The Gene Ontology (GO) analyses were conducted using the “clusterProfiler” package in R ^142^.

### Measurement of embryo hatchability

We collected embryos from WT and mutant flies and manually seeded them in vials containing standard corn medium at a density of 30 embryos per vial and 20 vials per strain. The embryos were cultivated under standard conditions (60% humidity, 12 h light: 12 h dark cycles at 25°C). We counted the number of pupae in each vial after the completion of pupation and calculated the corresponding hatching rate = *number*_*pupa*_ / 30

### Quantification of offspring number per female fly

Newly hatched virgins were picked out and allowed to mature for two days in separate vials. They were then mated by placing one virgin female with three male flies for two days. After that, each female parent was transferred to a new vial to count the offspring number produced in each 10-day period at 25°C and 29°C. All assays were performed with 20 females per genotype.

### Measurement of starvation resistance in adult flies

We selected 3- to 5-day-old adult males and females and placed them in the starvation medium (1.5% agar), with 10 flies per vial and 10 vials for both males and females from each strain. We made observations every 6 h or 12 h to count the number of deaths under starvation conditions until all flies had starved to death. The survival curves were plotted by the ggsurvplot package in R.

### mRNA-Seq and Ribo-Seq data analysis in primates

The mRNA-Seq and Ribo-Seq data from the brains, livers, and testes of humans and macaques were downloaded from reference ^79^ with accession number E-MTAB-7247 (ArrayExpress). The mRNA-Seq and Ribo-Seq data of human lymphoblastoid cell lines (LCL) from Yoruba individuals were downloaded from references ^81,82^ under accession numbers GSE61742 and E-GEUV-1. The matched mRNA-Seq and Ribo-Seq libraries of 69 individuals were used.

The uORF annotation and downstream analysis procedures for the human and macaque data were similar to those applied in *Drosophila* as described above. The differential analysis of translational efficiency in humans and macaques was conducted by Xtail ^143^. In each pair of human-macaque samples, expressed genes were defined as the genes with a CDS RPKM_mRNA_ > 0.1 in both species. The translated uORFs in a sample were defined as uORFs with a TE > 0.1. For the human cell line data, expressed genes were defined as genes with a mean CDS RPKM_mRNA_ > 0.1 across the cell lines, and translated uORFs were defined as uORFs with a mean TE > 0.1

## Acknowledgments

We thank Drs. Wei Xie, Weiwei Zhai, and Xionglei He for constructive comments and suggestions. This work was supported by grants from the Ministry of Science and Technology of the People’s Republic of China (2022YFE0132000), the Yunnan Provincial Science and Technology Project at Southwest United Graduate School (202302A0370006), the National Natural Science Foundation of China (32070597) and the Natural Science Foundation of Beijing (5212006). We thank the National Center for Protein Sciences at Peking University for their technical assistance. Some of the analyses were performed on the High-Performance Computing Platform of the Center for Life Sciences.

## Declaration of interests

The authors declare no competing interests.

## Data Availability Statement

All deep-sequencing data generated in this study, including single-ended mRNA-Seq and Ribo-Seq data of 10 developmental stages and tissues of *Drosophila simulans* and paired-end mRNA-Seq data of 0–2 h, 2–6 h, 6–12 h, and 12–24 h *Drosophila melanogaster* embryos, were deposited in the China National Genomics Data Center Genome Sequence Archive (GSA) under accession numbers CRA003198, CRA007425, and CRA007426. The mRNA-Seq and Ribo-Seq data for the different developmental stages and tissues of *Drosophila melanogaster* were published in our previous paper ^23^ and were deposited in the Sequence Read Archive (SRA) under accession number SRP067542. All original code has been deposited on GitHub: https://github.com/lujlab/uORF_buffer; https://github.com/lujlab/Buffer_eLife2025.

## Supplementary Materials and Methods

### The detailed operations per action in the ICIER simulation

The translation process of an mRNA molecule is modeled as a series of discrete actions. During each action, certain events may occur with certain probabilities for each entity (40S or 80S ribosome). The operations per action are described as follows:

1. If the first 10 triplets were vacant of the mRNA, a 40S ribosome would be added there with a probability of *R_in_*.
2. If a 40S ribosome was located at the start codon of the uORF, it would transform into an 80S ribosome with a probability of *I*_*uORF*_. If a 40S ribosome was located at the start codon of the CDS, it would transform into an 80S ribosome with a probability of *I*_*CCS*_.
3. If an 80S ribosome was located at the stop codon of the uORF or CDS, it would dissociate from the mRNA. The dissociation of an 80S ribosome from the uORF or CDS stop codon was counted as a completed translation event, and the values of *N_EC_* and *N_Eu_* were incremented by one, respectively.
4. If a 40S ribosome arrived at the 3’ end of the mRNA, it would dissociate from the mRNA.
5. For each 40S ribosome on the mRNA, if its next position was vacant (in the downstream-dissociation model) or not occupied by another 40S ribosome (in the upstream-dissociation model and double-dissociation model), it would move to the next position with a probability of *v*_*s*_.
6. For each 80S ribosome in the uORF, if its next position was vacant (in the upstream-dissociation model) or not occupied by another 80S ribosome (in the downstream-dissociation model and double-dissociation model), it would move to the next position with a probability of *v*_*Eu*_. Similarly, for each 80S ribosome in the CDS, if its next position was vacant (in the upstream-dissociation model) or not occupied by an 80S ribosome (in the downstream-dissociation model and double-dissociation model), it would move to the next position with a probability of *v*_*EC*_.
7. It was checked whether any 80S ribosome collided with a 40S ribosome. If such a collision occurred, the 40S ribosome involved would dissociate from the mRNA with a probability of *K_up_* (upstream dissociation or double-dissociation model) or *K_down_* (downstream dissociation or double-dissociation model).

These above operations are executed repeatedly until the action count reaches the limit given by the user (1,000,000 in this study).

## Supplementary Figures and Tables

**Figure S1.**
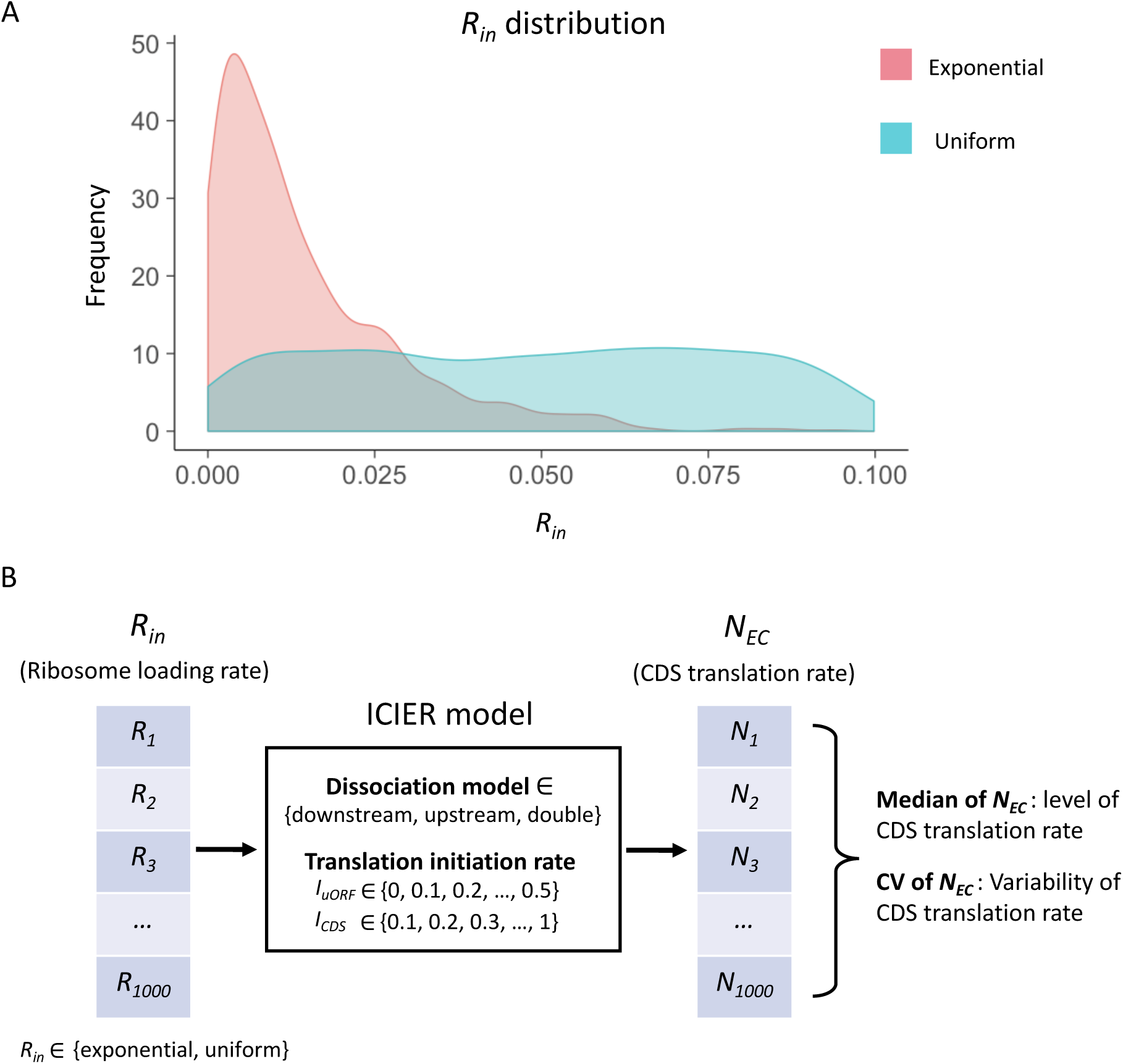
The input distribution and model simulation flow. (A) Two distributions of the *R_in_* input were used in our simulation: an exponential distribution and a uniform distribution. A total of 1,000 *R*_*in*_ values (ranging from 0 to 0.1) were randomly generated following either distribution. (B) Under a fixed combination of parameters (*R_in_*distribution, dissociation model, *I_uORF_* and *I_CDS_*), the model simulation produces 1000 *N_EC_* values from 1000 *R_in_* input. The median of these 1000 *N_EC_* represented the median of translation rate of CDS under the 1000 varying *R_in_* inputs and the other fixed parameters. The the coefficient of variation (CV) of these 1000 *N_EC_* reflected the variability of translation rate of CDS under the 1000 varying *R_in_* inputs and the other fixed parameters.

**Figure S2.**
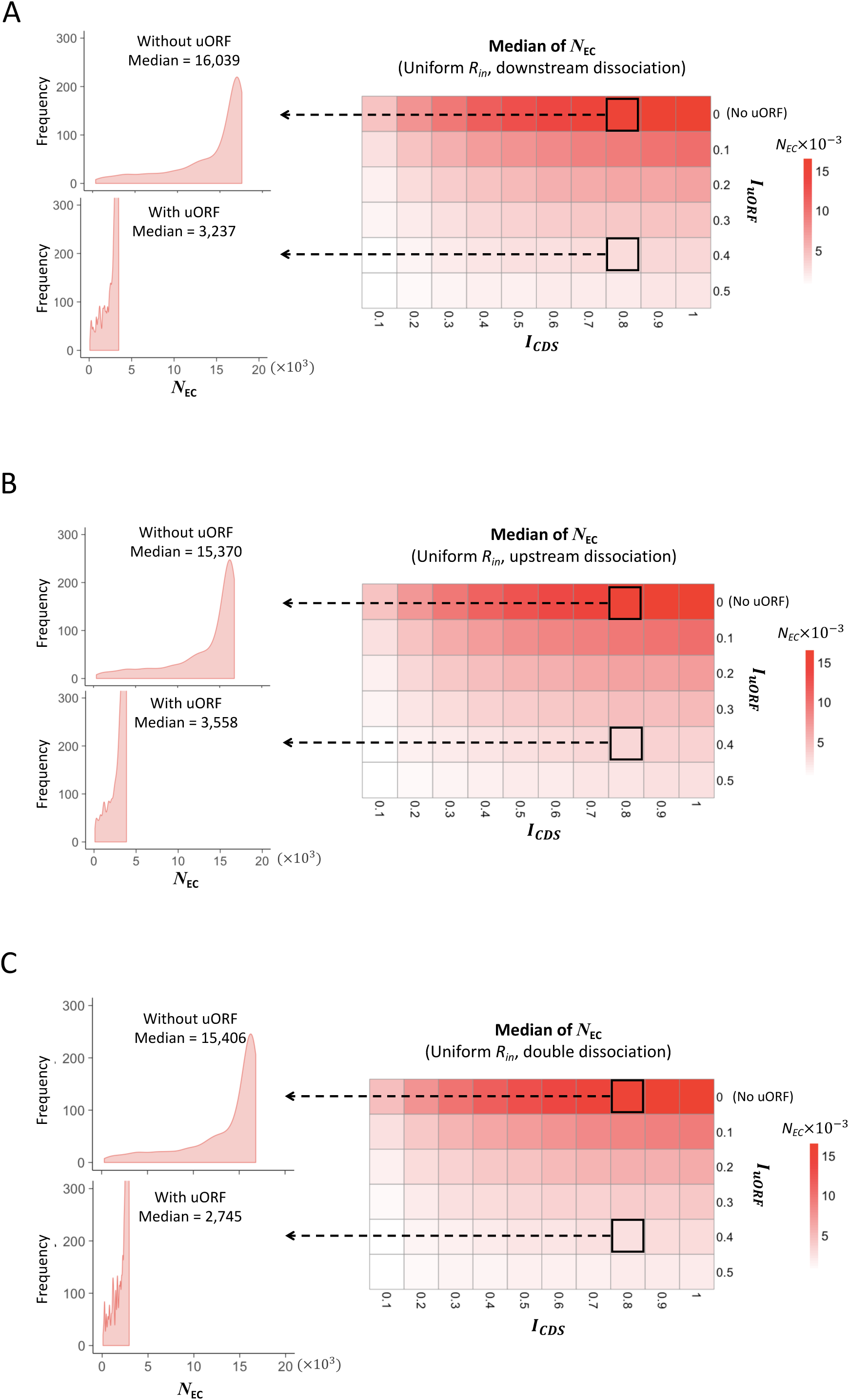
Heatmaps showing the median CDS translation rate (*N_EC_*) under different *I_CDS_* (x-axis) and *I_uORF_*(y-axis) combinations with a uniform distribution of *R_in_* input and the downstream dissociation model (A), a uniform distribution of *R_in_* input and the upstream dissociation model (B), a uniform distribution of *R_in_* input and the double dissociation model (C). For each heatmap, the left panels elicited by the dotted lines from specific squares of right heatmap were two examples showing the distribution of *N_EC_* under *I_CDS_* = 0.8 & *I_uORF_* = 0 (top panel, without uORF) and *I_CDS_*= 0.8 & *I_uORF_* = 0.4 (bottom panel, with uORF).

**Figure S3.**
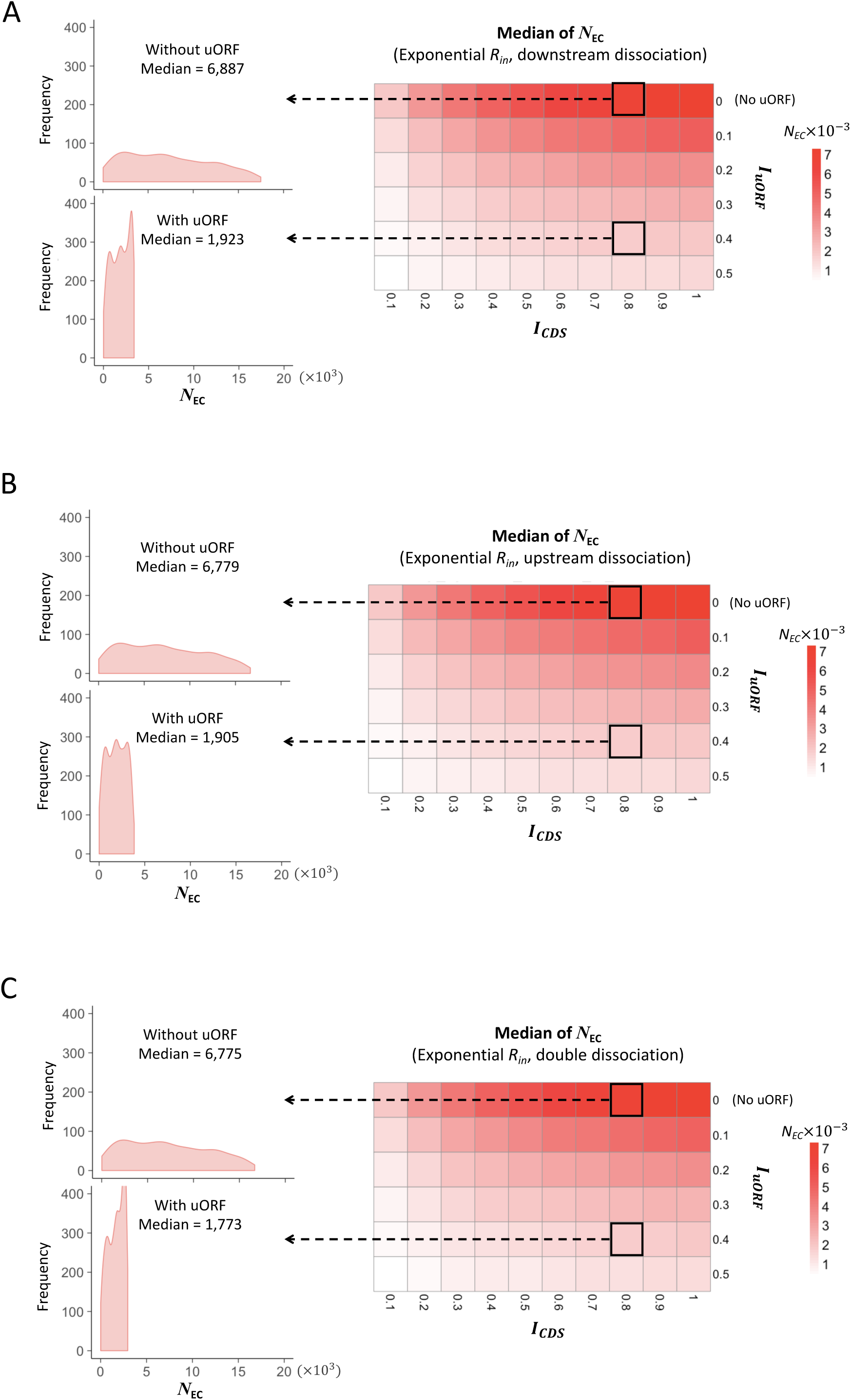
Heatmaps showing the median CDS translation rate (*N_EC_*) under different *I_CDS_* (x-axis) and *I_uORF_* (y-axis) combinations with an exponential distribution of *R_in_* input and the downstream dissociation model (A), an exponential distribution of *R_in_* input and the upstream dissociation model (B), an exponential distribution of *R_in_* input and the double dissociation model (C). For each heatmap, the left panels elicited by the dotted lines from specific squares of right heatmap were two examples showing the distribution of *N_EC_* under *I_CDS_* = 0.8 & *I_uORF_* = 0 (top panel, without uORF) and *I_CDS_* = 0.8 & *I_uORF_*= 0.4 (bottom panel, with uORF).

**Figure S4.**
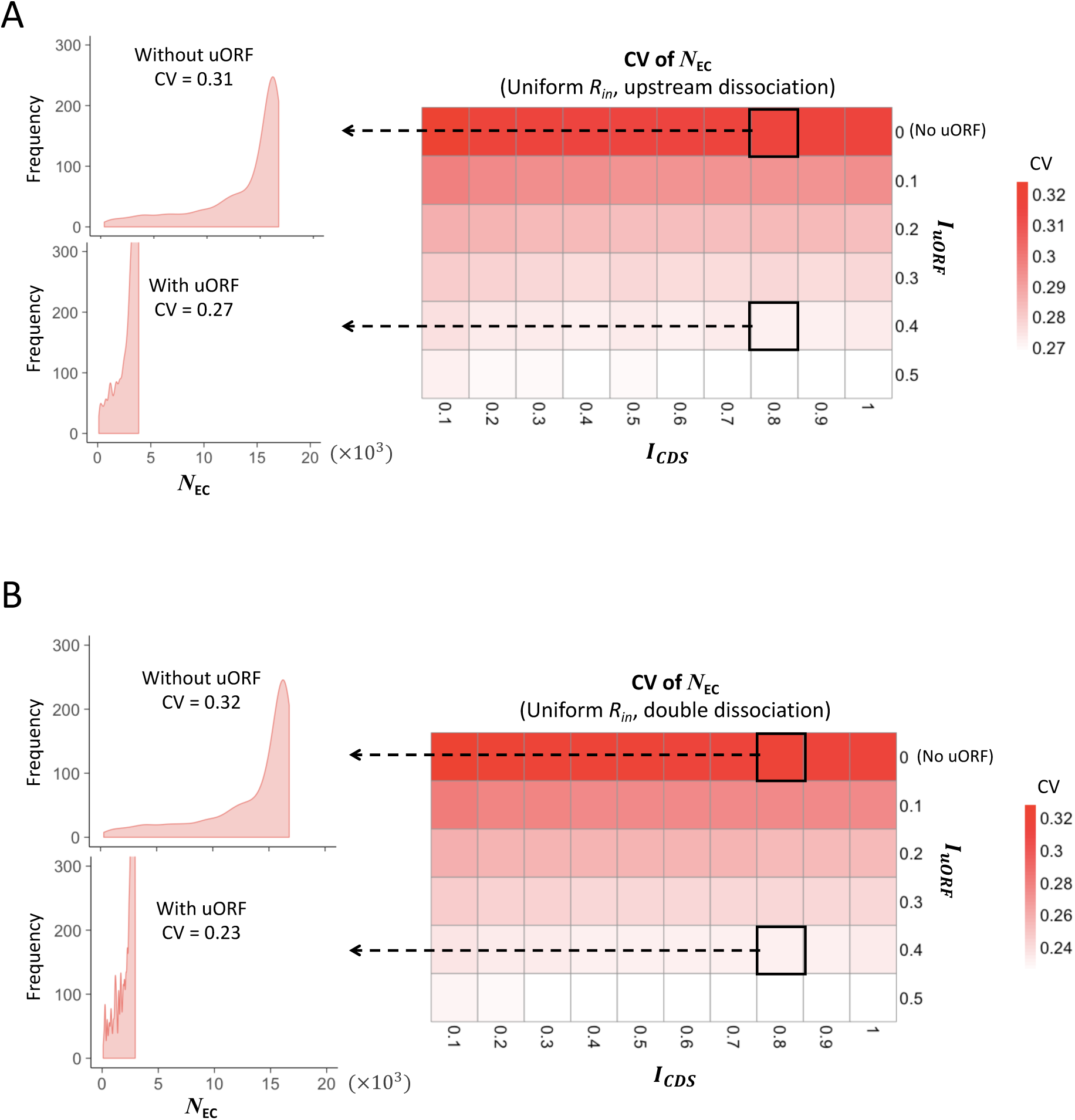
Heatmaps showing the CVs of CDS translation rate (*N_EC_*) under different *I_CDS_* (x-axis) and *I_uORF_*(y-axis) combinations with a uniform distribution of *R_in_* input and the upstream dissociation model (A), a uniform distribution of *R_in_* input and the double dissociation model (B). For each heatmap, the left panels elicited by the dotted lines from specific squares of right heatmap were two examples showing the distribution of *N_EC_* under *I_CDS_* = 0.8 & *I_uORF_* = 0 (top panel, without uORF) and *I_CDS_* = 0.8 & *I_uORF_*= 0.4 (bottom panel, with uORF).

**Figure S5.**
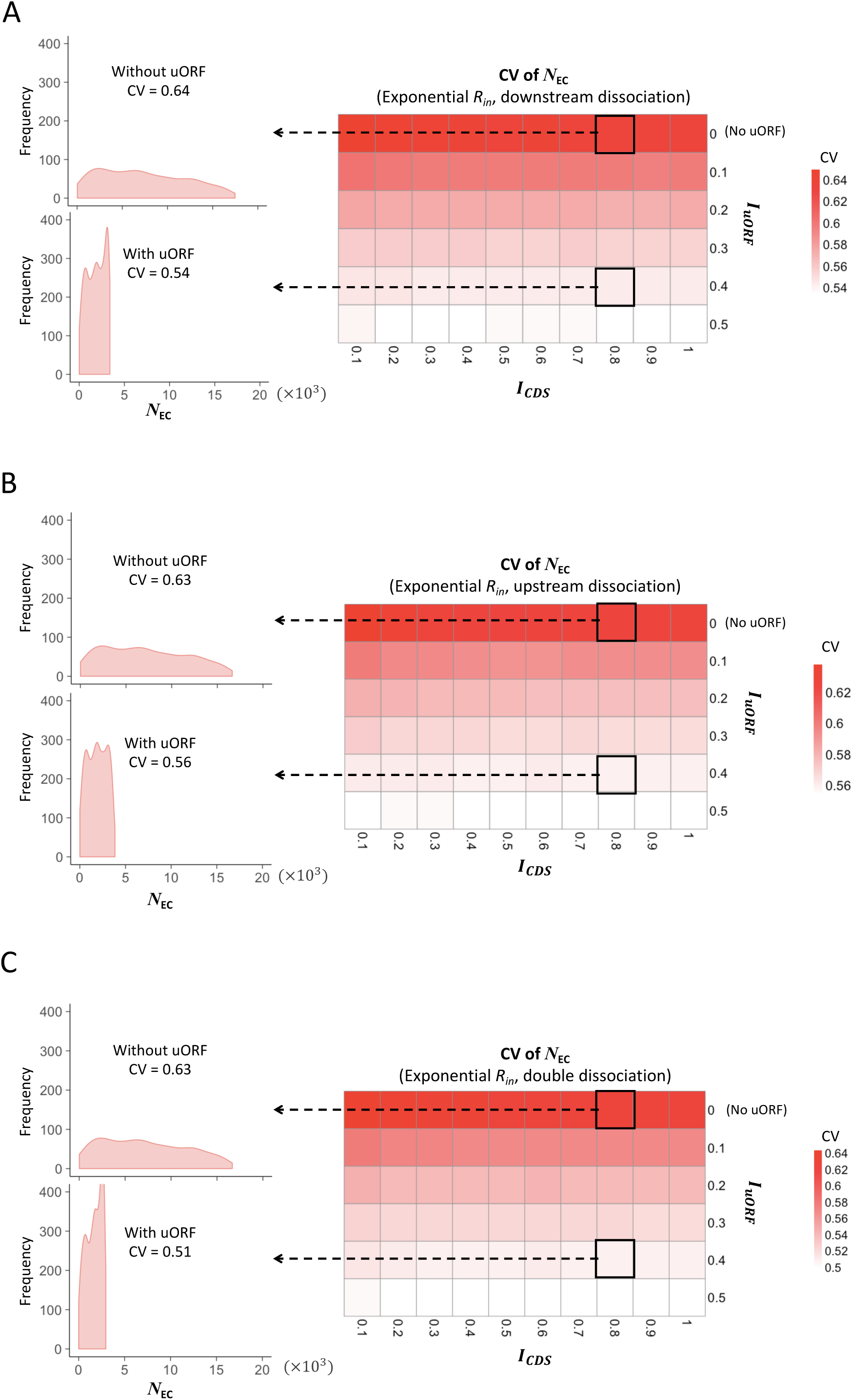
Heatmaps showing the CVs of CDS translation rate (*N_EC_*) under different *I_CDS_* (x-axis) and *I_uORF_* (y-axis) combinations with an exponential distribution of *R_in_* input and the downstream dissociation model (A), an exponential distribution of *R_in_* input and the upstream dissociation model (B), an exponential distribution of *R_in_* input and the double dissociation model (C). For each heatmap, the left panels elicited by the dotted lines from specific squares of right heatmap were two examples showing the distribution of *N_EC_* under *I_CDS_* = 0.8 & *I_uORF_* = 0 (top panel, without uORF) and *I_CDS_* = 0.8 & *I_uORF_*= 0.4 (bottom panel, with uORF).

**Figure S6.**
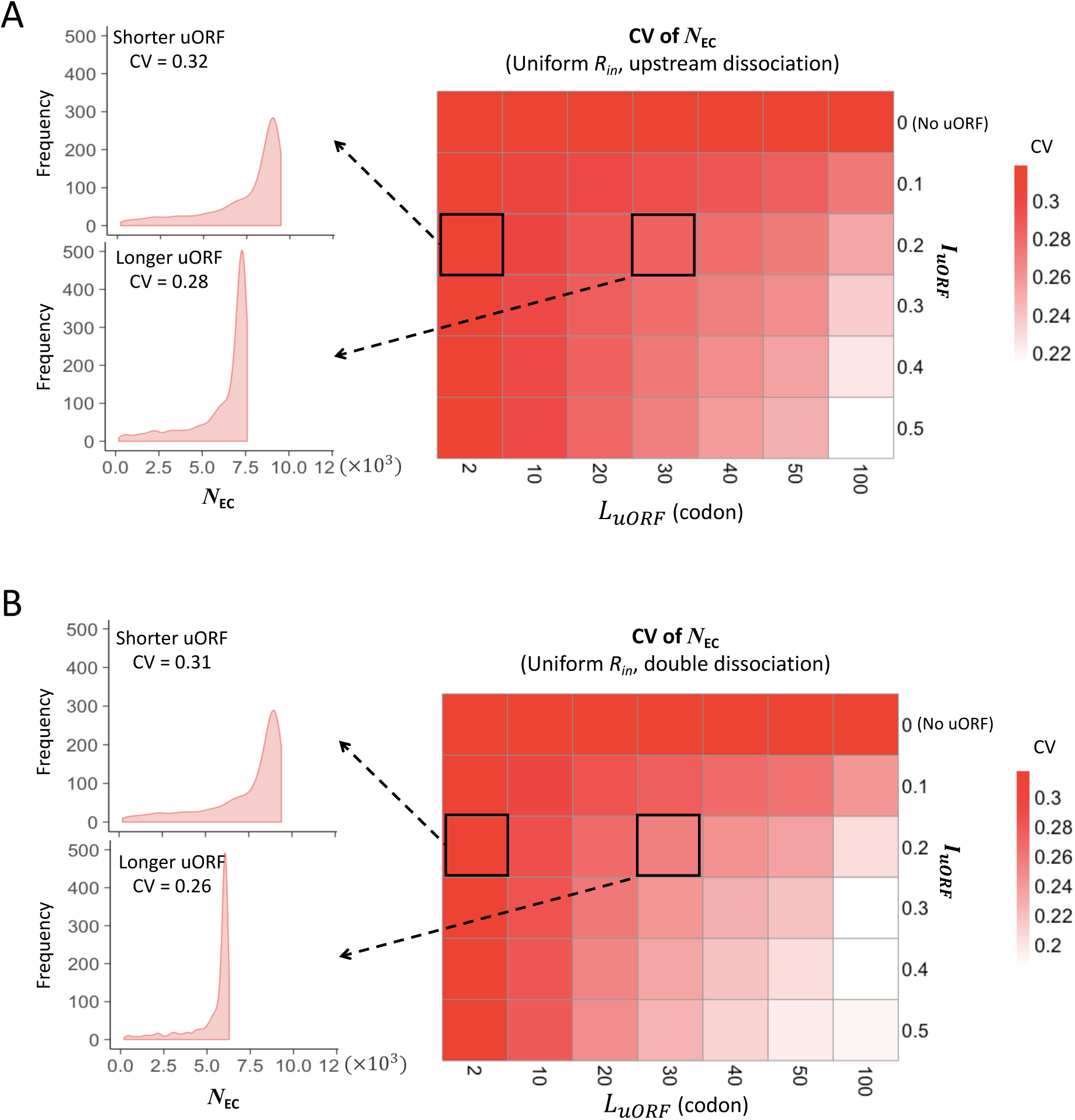
Heatmaps showing the CVs of CDS translation rate (*N_EC_*) under different *L_uORF_* (x-axis) and *I_uORF_* (y-axis) combinations with a uniform distribution of *R_in_* input and the upstream dissociation model (A), a uniform distribution of *R_in_* input and the double dissociation model (B). For each heatmap, the left panels elicited by the dotted lines from specific squares of right heatmap were two examples showing the distribution of *N_EC_* under *L_uORF_* = 2 & *I_uORF_* = 0.2 (top panel, shorter uORF) and *L_uORF_* = 30 & *I_uORF_* = 0.2 (bottom panel, longer uORF).

**Figure S7.**
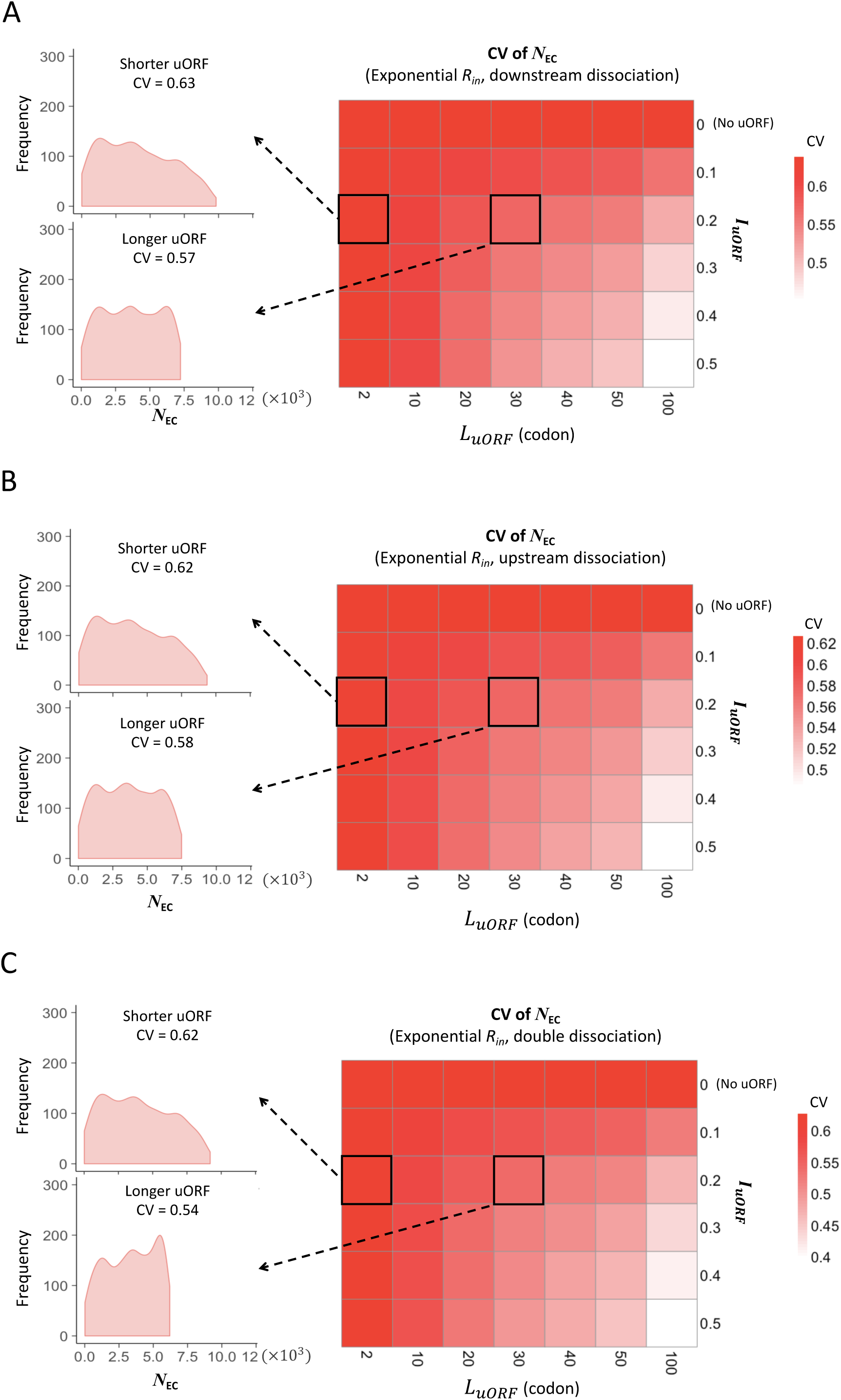
Heatmaps showing the CVs of CDS translation rate (*N_EC_*) under different *L_uORF_* (x-axis) and *I_uORF_* (y-axis) combinations with an exponential distribution of *R_in_* input and the downstream dissociation model (A), an exponential distribution of *R_in_* input and the upstream dissociation model (B), an exponential distribution of *R_in_* input and the double dissociation model (C). For each heatmap, the left panels elicited by the dotted lines from specific squares of right heatmap were two examples showing the distribution of *N_EC_* under *L_uORF_* = 2 & *I_uORF_* = 0.2 (top panel, shorter uORF) and *L_uORF_* = 30 & *I_uORF_* = 0.2 (bottom panel, longer uORF).

**Figure S8.**
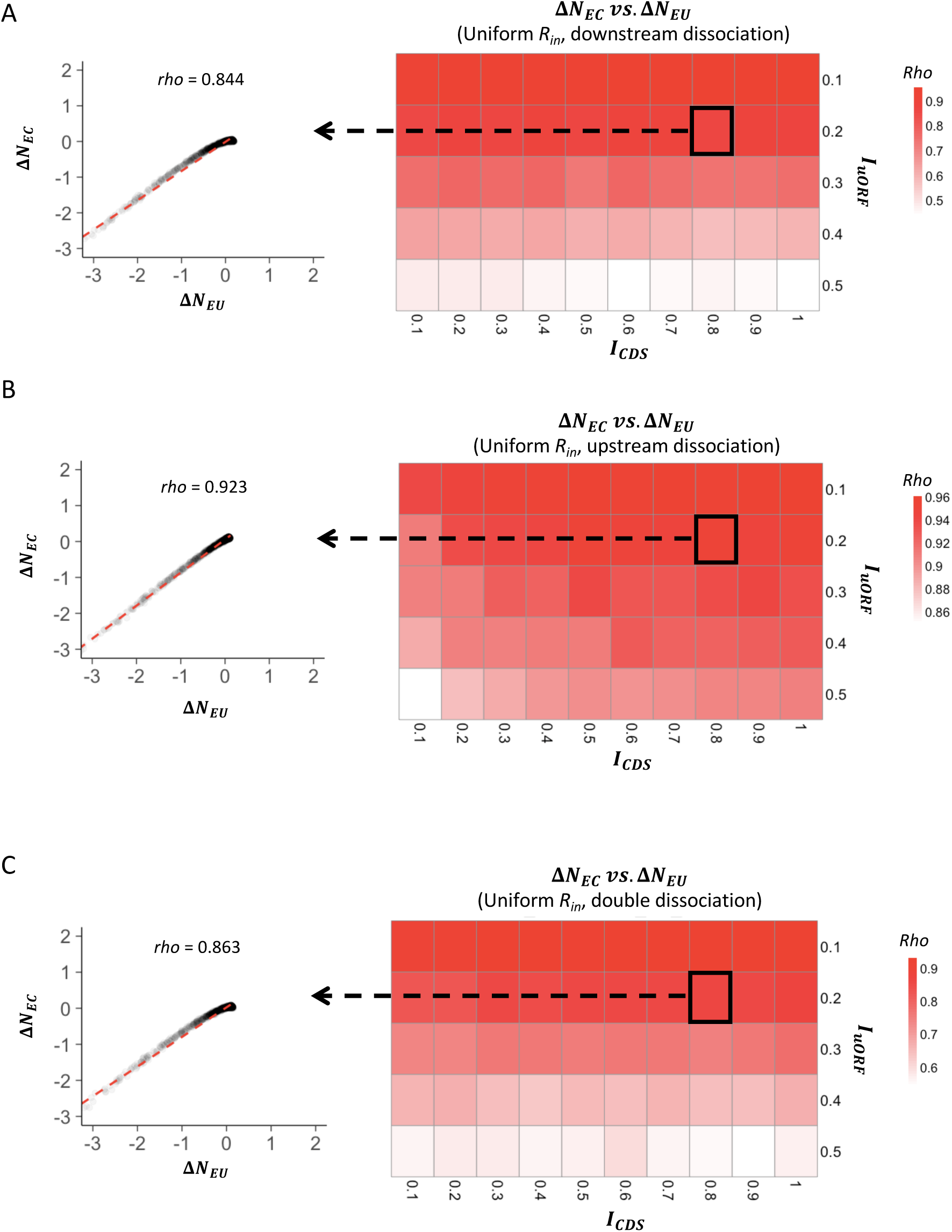
Heatmaps showing the Spearman’s correlations between changes of uORF translation rate (Δ*N*_*EU*_) and downstream CDS translation rate (Δ*N*_*EC*_) under different *I_CDS_* (x-axis) and *I_uORF_* (y-axis) combinations with a uniform distribution of *R_in_* input and the downstream dissociation model (A), a uniform distribution of *R_in_* input and the upstream dissociation model (B), a uniform distribution of *R_in_*input and the double dissociation model (C). For each heatmap, the left panel elicited by the dotted line from the specific square of right heatmap was an example showing the Spearman’s correlations between changes of uORF translation rate (Δ*N*_*EU*_) and downstream CDS translation rate (Δ*N*_*EC*_) under *I_CDS_* = 0.8 & *I_uORF_* = 0.2. All *P* values < 0.001.

**Figure S9.**
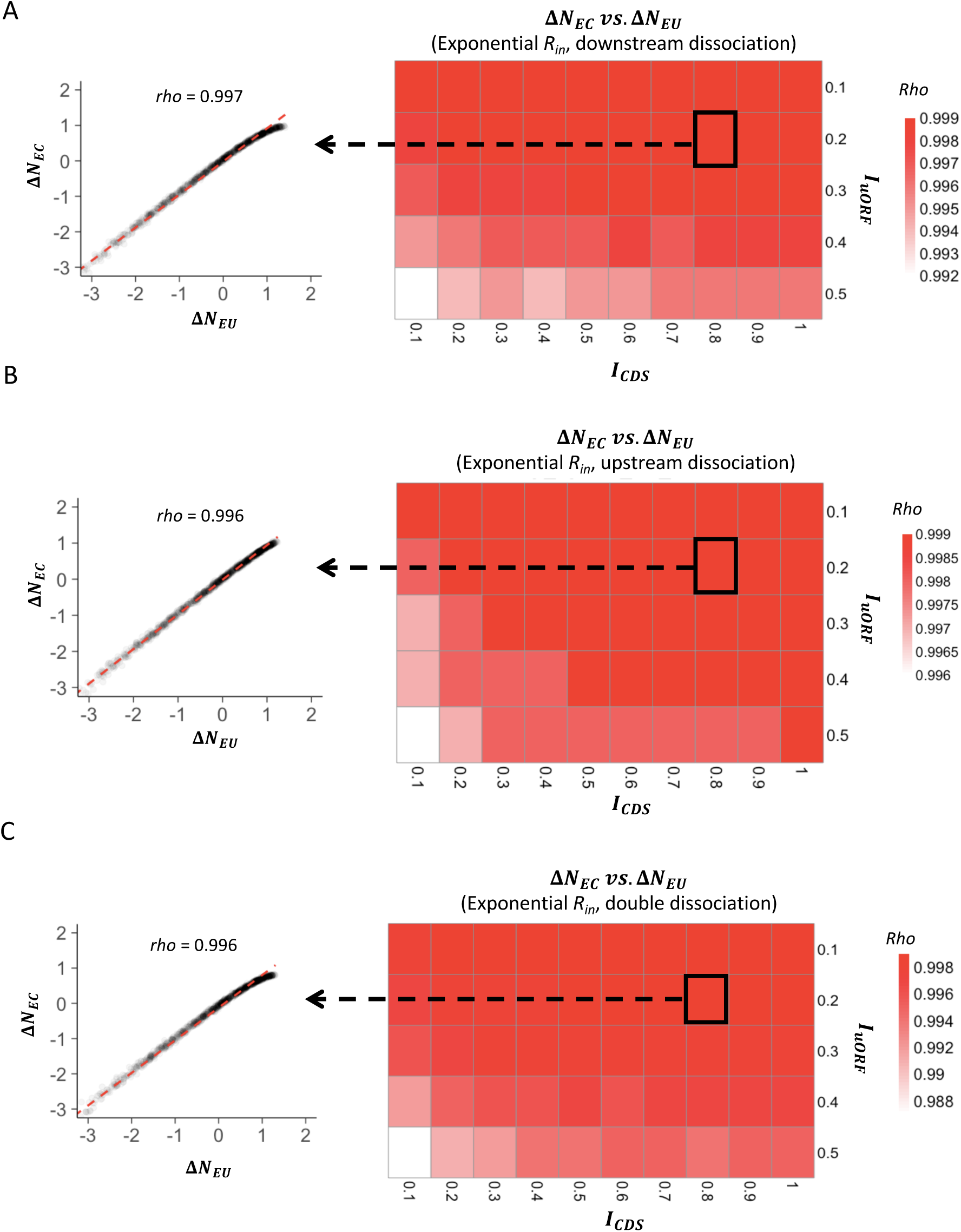
Heatmaps showing the Spearman’s correlations between changes of uORF translation rate (Δ*N*_*EU*_) and downstream CDS translation rate (Δ*N*_*EC*_) under different *I_CDS_* (x-axis) and *I_uORF_* (y-axis) combinations with an exponential distribution of *R_in_* input and the downstream dissociation model (A), an exponential distribution of *R_in_*input and the upstream dissociation model (B), an exponential distribution of *R_in_* input and the double dissociation model (C). For each heatmap, the left panel elicited by the dotted line from the specific square of right heatmap was an example showing the Spearman’s correlations between changes of uORF translation rate (Δ*N*_*EU*_) and downstream CDS translation rate (Δ*N*_*EC*_) under *I_CDS_* = 0.8 & *I_uORF_* = 0.2. All *P* values < 0.001.

**Figure S10.**
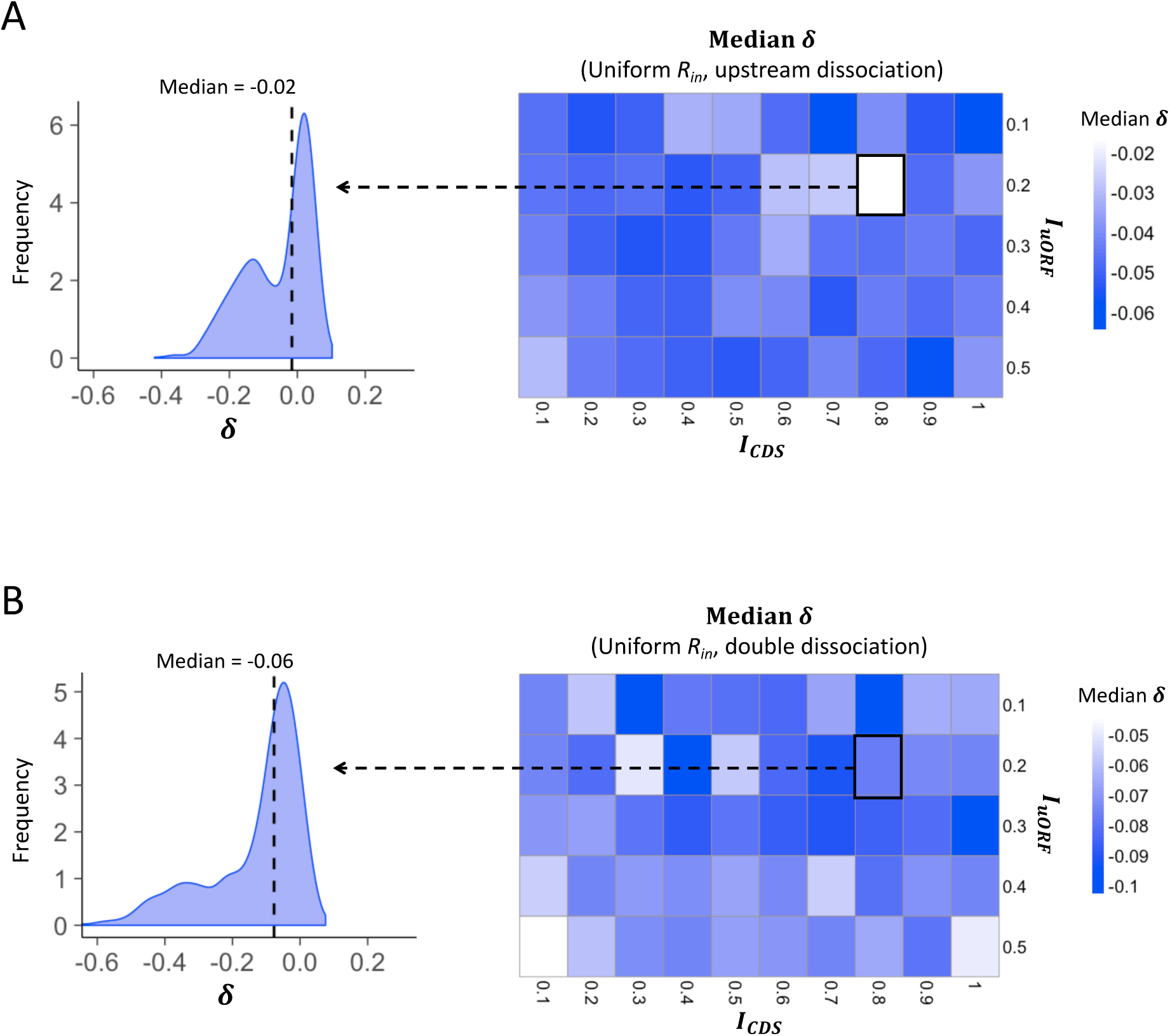
Heatmaps showing the median *δ* under different *I_CDS_* (x-axis) and *I_uORF_* (y-axis) combinations with a uniform distribution of *R_in_* input and the upstream dissociation model (A), a uniform distribution of *R_in_* input, and the double dissociation model (B). For each heatmap, the left panel elicited by the dotted line from the specific square of right heatmap was an example showing the distribution of *δ* under *I_CDS_* = 0.8 & *I_uORF_* = 0.2. The vertical dashed line indicated the median value of *δ*.

**Figure S11.**
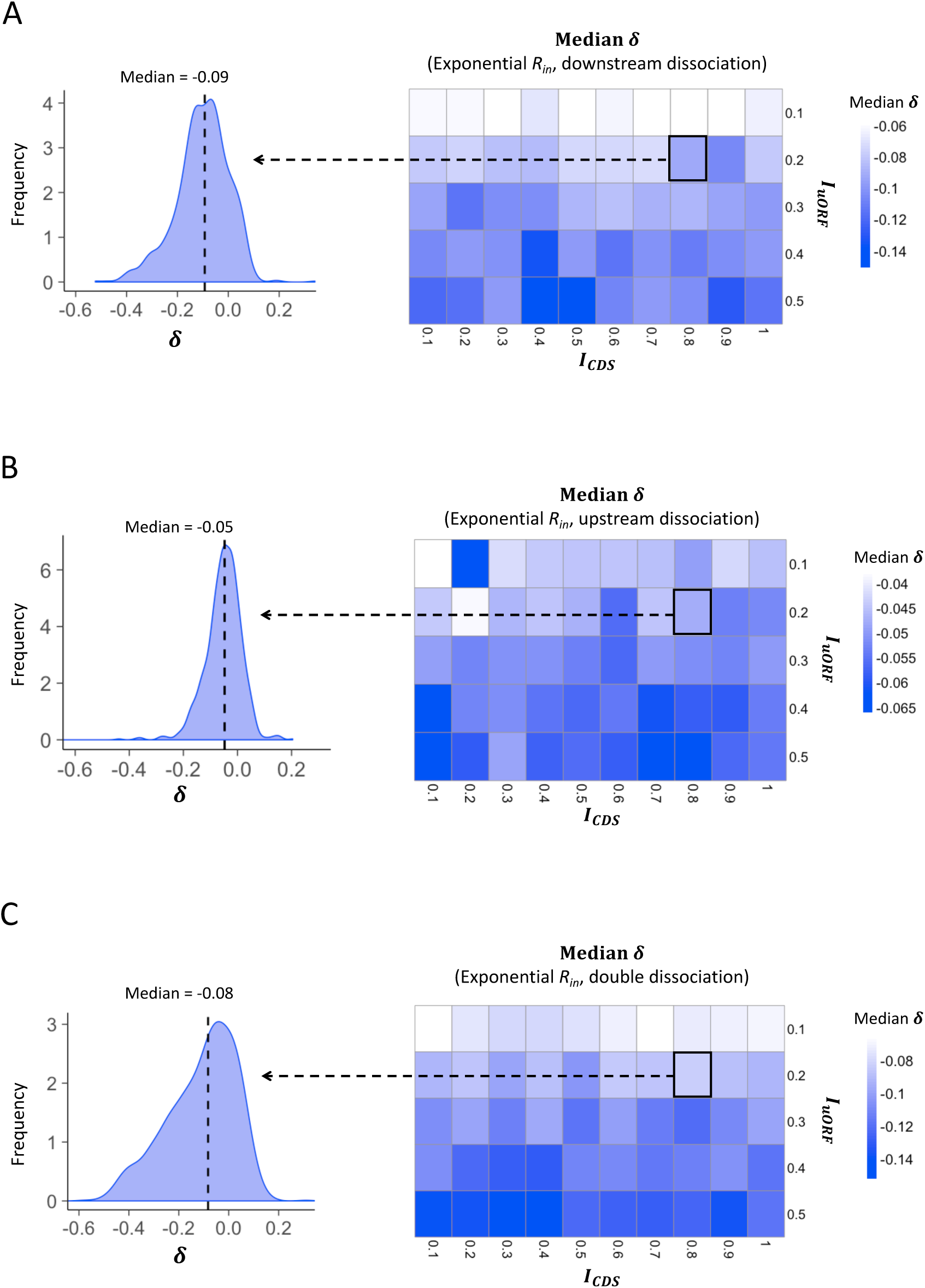
Heatmaps showing the median *δ* under different *I_CDS_* (x-axis) and *I_uORF_* (y-axis) combinations with an exponential distribution of *R_in_* input and the downstream dissociation model (A), an exponential distribution of *R_in_* input and the upstream dissociation model (B), an exponential distribution of *R_in_* input and the double dissociation model (C). For each heatmap, the left panel elicited by the dotted line from the specific square of right heatmap was an example showing the distribution of *δ* under *I_CDS_* = 0.8 & *I_uORF_* = 0.2. The vertical dashed line indicated the median value of *δ*.

**Figure S12.**
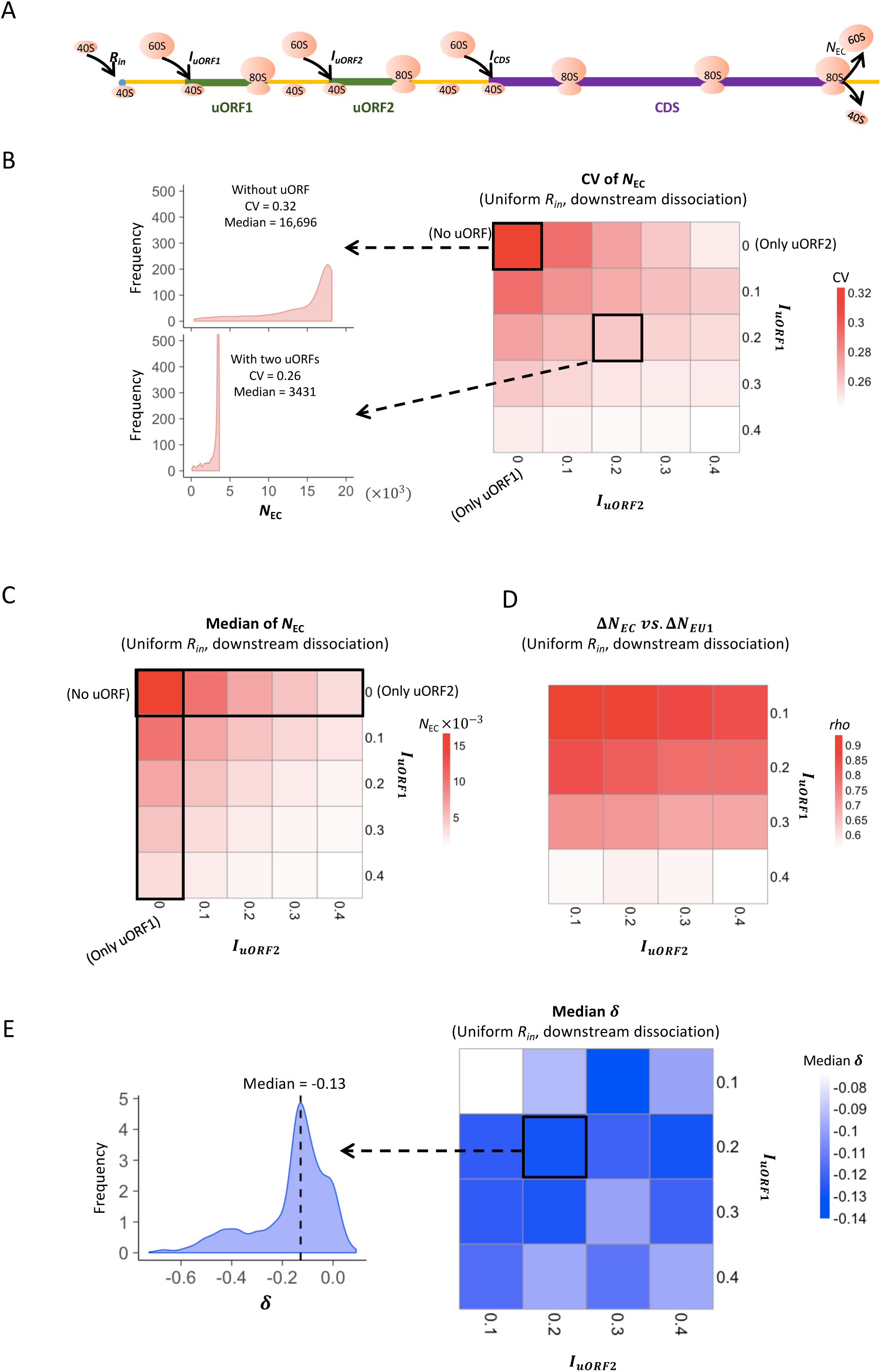
Two-uORF model simulation measurements. (A) The two-uORF model schema. The factors and parameters were the same as those illustrated in Fig. 1A except where specifically indicated. (B) Heatmap showing the CVs of CDS translation rate (*N_EC_*) under different *I_uORF2_* (x-axis) and *I_uORF1_*(y-axis) combinations with a uniform distribution of *R_in_* input and the downstream dissociation model. The left panels elicited by the dotted lines from specific squares of right heatmap were two examples showing the distribution of *N_EC_* under *I_uORF1_*= 0 & *I_uORF2_* = 0 (top panel, without uORF) and *I_uORF1_*= 0.2 & *I_uORF2_* = 0.2 (bottom panel, with two uORFs). (C) Heatmap showing the median CDS translation rate (*N_EC_*) under different *I_uORF2_* (x-axis) and *I_uORF1_* (y-axis) combinations with a uniform distribution of *R_in_* input and the downstream dissociation model. (D) Spearman’s correlations between changes of uORF1 translation rate (*ΔN*_*EU*1_) and downstream CDS translation rate (*ΔN*_*EC*_) under different *I_uORF2_* (x-axis) and *I_uORF1_* (y-axis) combinations with a uniform distribution of *R_in_* input and the downstream dissociation model. (E) Heatmap showing the median *δ* [*log*_2_(Δ*N*_*EC*_/Δ*N*_*EU*1_)] under different *I_uORF2_* (x-axis) and *I_uORF1_* (y-axis) combinations with a uniform distribution of *R_in_* input and the downstream dissociation model. The left panel elicited by the dotted line from the specific square of right heatmap was an example showing the distribution of *δ* under *I_uORF1_* = 0.2 & *I_uORF2_* = 0.2. The vertical dashed line indicated the median value of *δ*.

**Figure S13.**
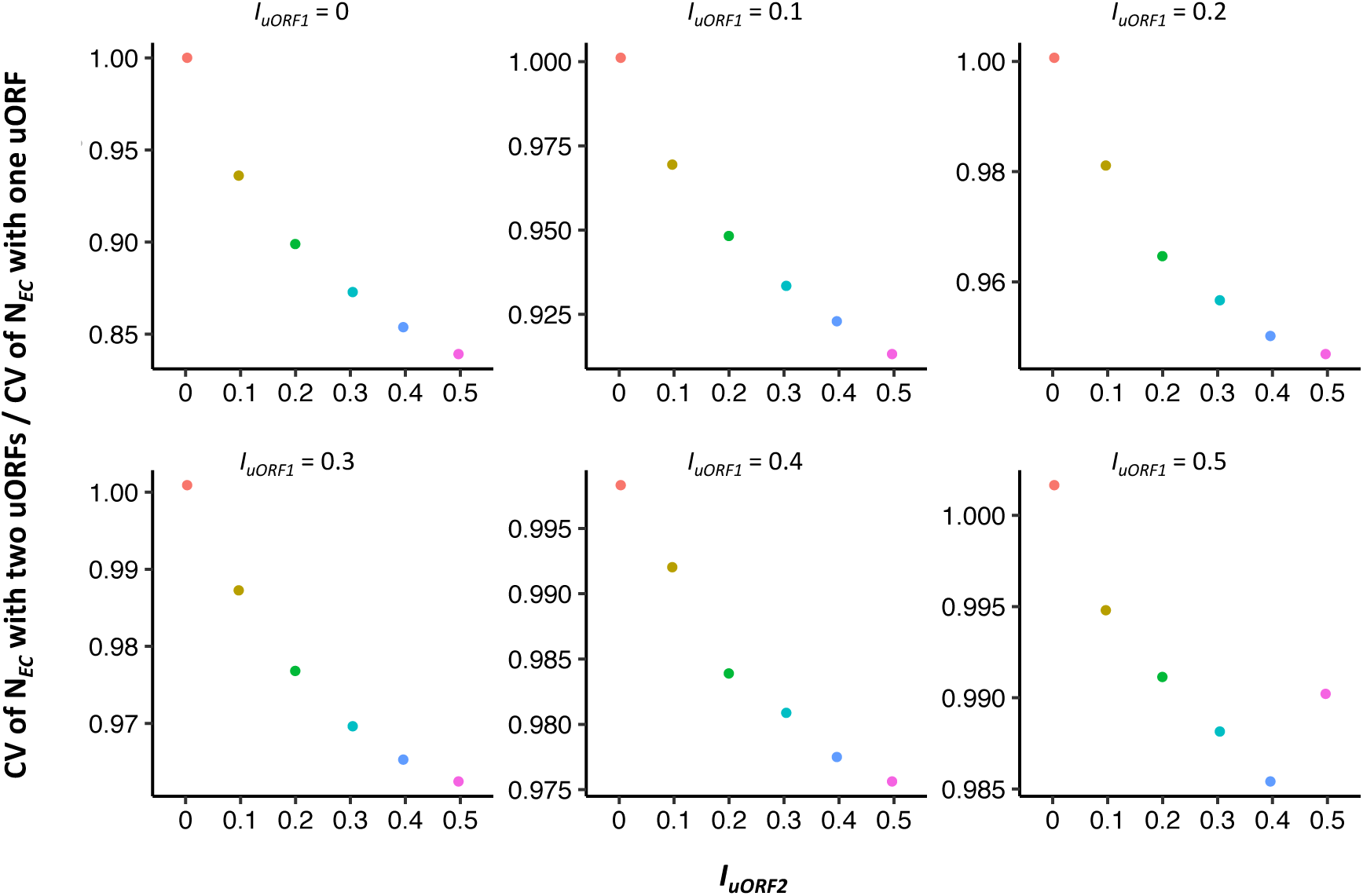
Comparison of the buffering effects between single uORF and two uORFs. In each panel, the *I*_*uORF*_ of single-uORF model equals to *I*_*uORF*1_of two-uORF model, with both values ranging from 0 to 0.5. The x-axis in each panel denotes the values of different *I*_*uORF*2_ in the two-uORF model, ranging from 0 to 0.5. The y-axis in each panel represents the ratio of the CV of *N*_*EC*_ with two uORFs to that with a single uORF.

**Figure S14.**
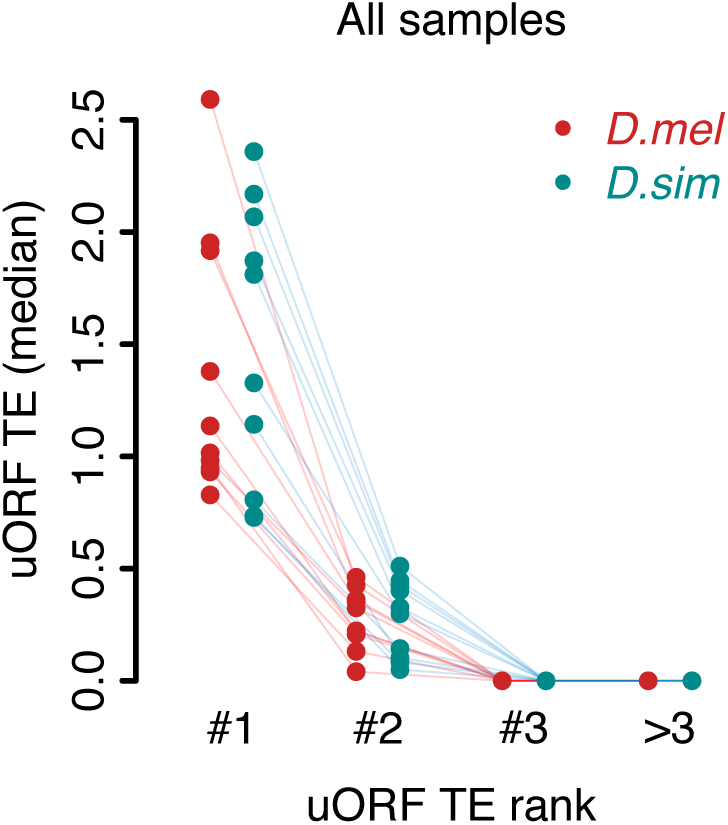
The dominant uORF showed highest TE than other uORFs with a same gene. uORFs were ranked by decreasing TEs within each gene. The uORF with the highest TE within each gene was defined as the dominant uORF (#1). “#2” represents the second highest uORF TE and the same goes for “#3” and “>3”. Each dot represents the median TE of a sample.

**Figure S15.**
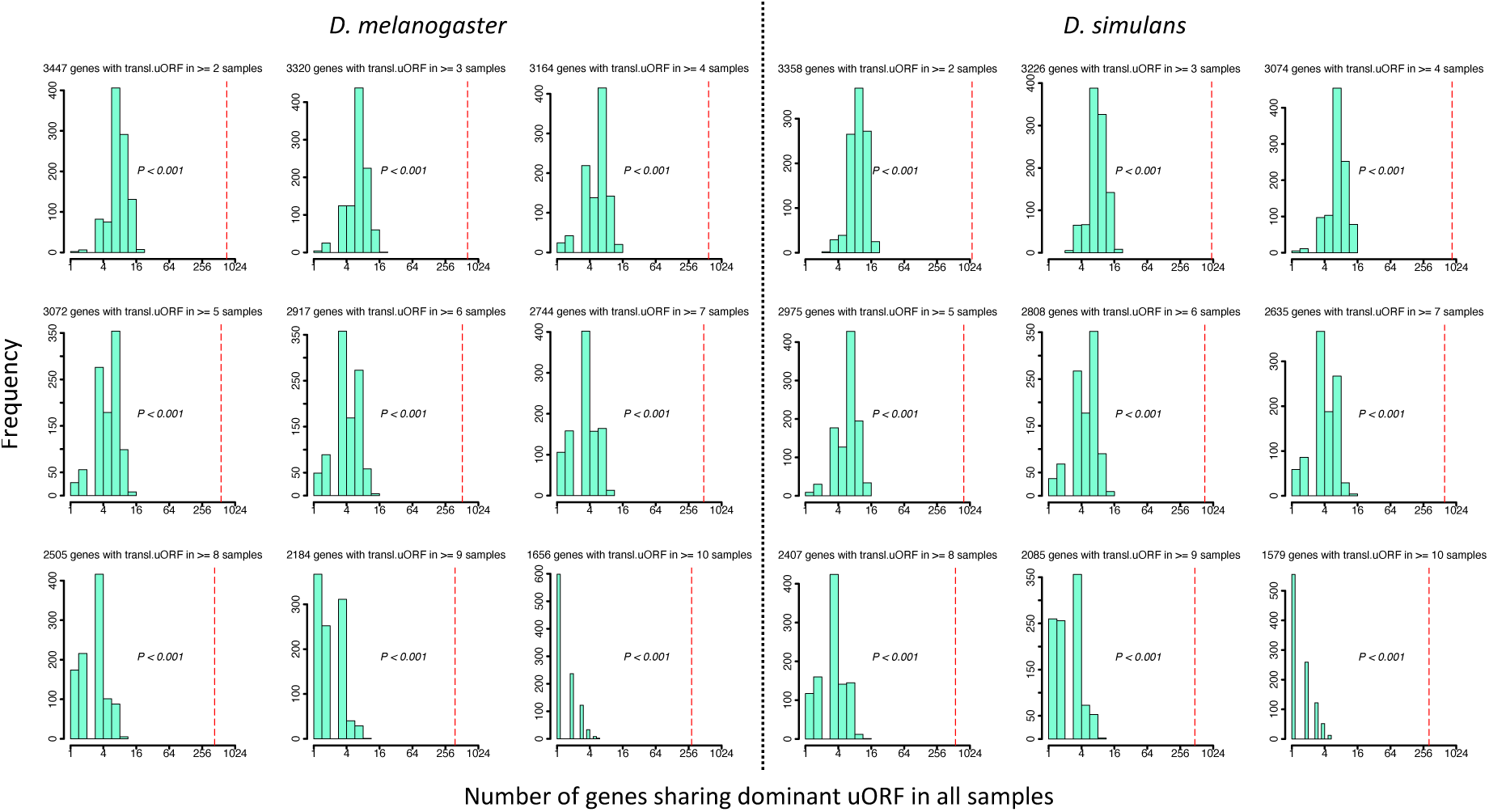
Observed and expected numbers of genes sharing dominant uORFs in all samples. Among the genes of *D. melanogaster* (where only the longest transcripts were considered), 7,259 (52.2%) genes had no uORF (“no-uORF” genes), 2,687 (19.3%) genes had one uORF (“one-uORF” genes), and 3,961 (28.5%) genes had multiple (≥ 2) uORFs (“multiple-uORF” genes). The numbers in the header of each panel represent genes with translated uORFs (TE > 0.1) in ≥ N samples (2 ≤ N ≤ 10). The TEs of all uORFs were shuffled 1,000 times. *P* values were obtained from randomization tests.

**Figure S16.**
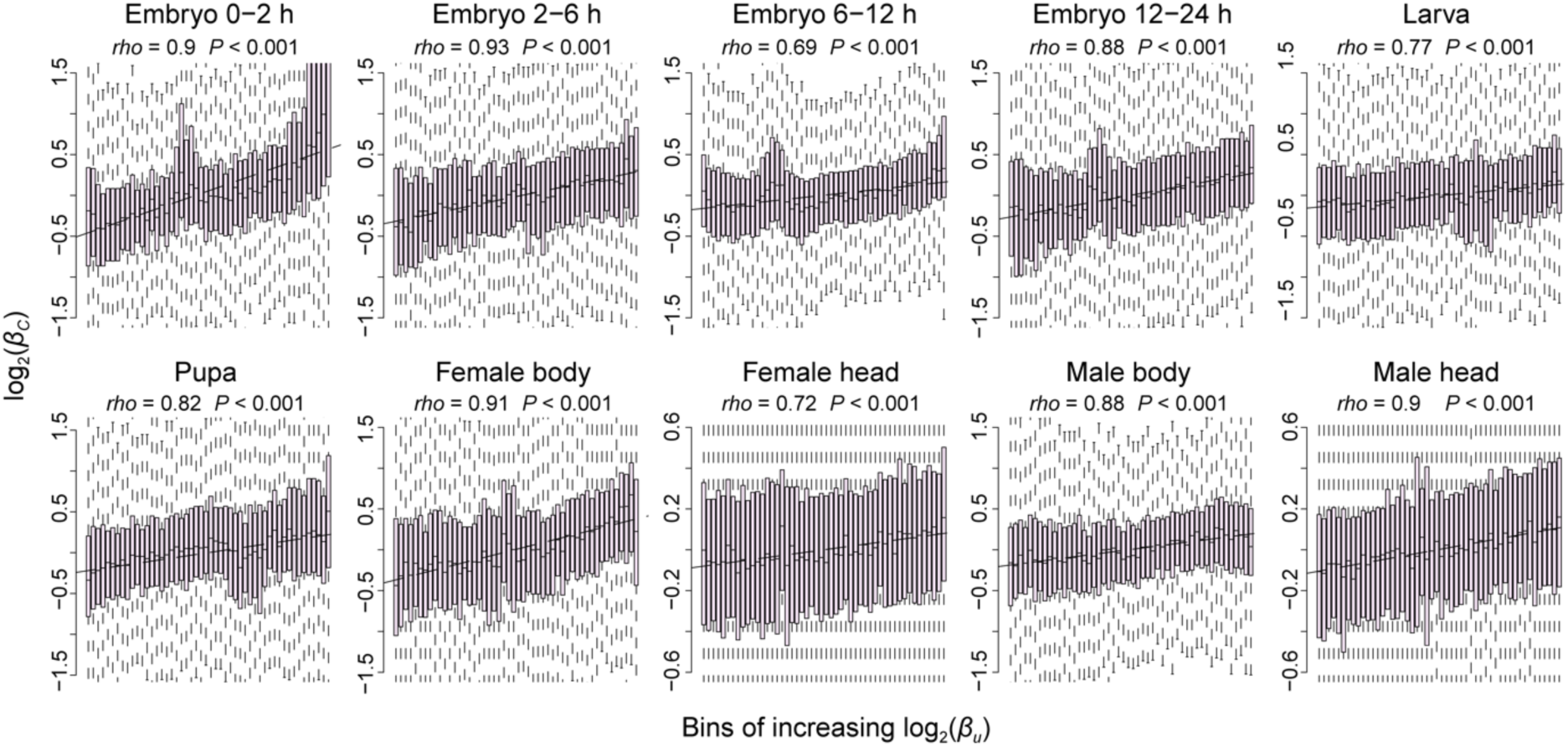
The positive correlation of interspecific TE changes between uORFs and CDSs. Correlations between interspecific uORF TE changes (log_2_*β*_*u*_) and CDS TE changes (log_2_*β*_*C*_) in 10 samples. The x-axis was divided into 50 equal bins with increasing *β*_*u*_. Spearman’s correlation coefficients (*rho*) are shown at the top left. ***, *P* < 0.001 in the correlation test.

**Figure S17.**
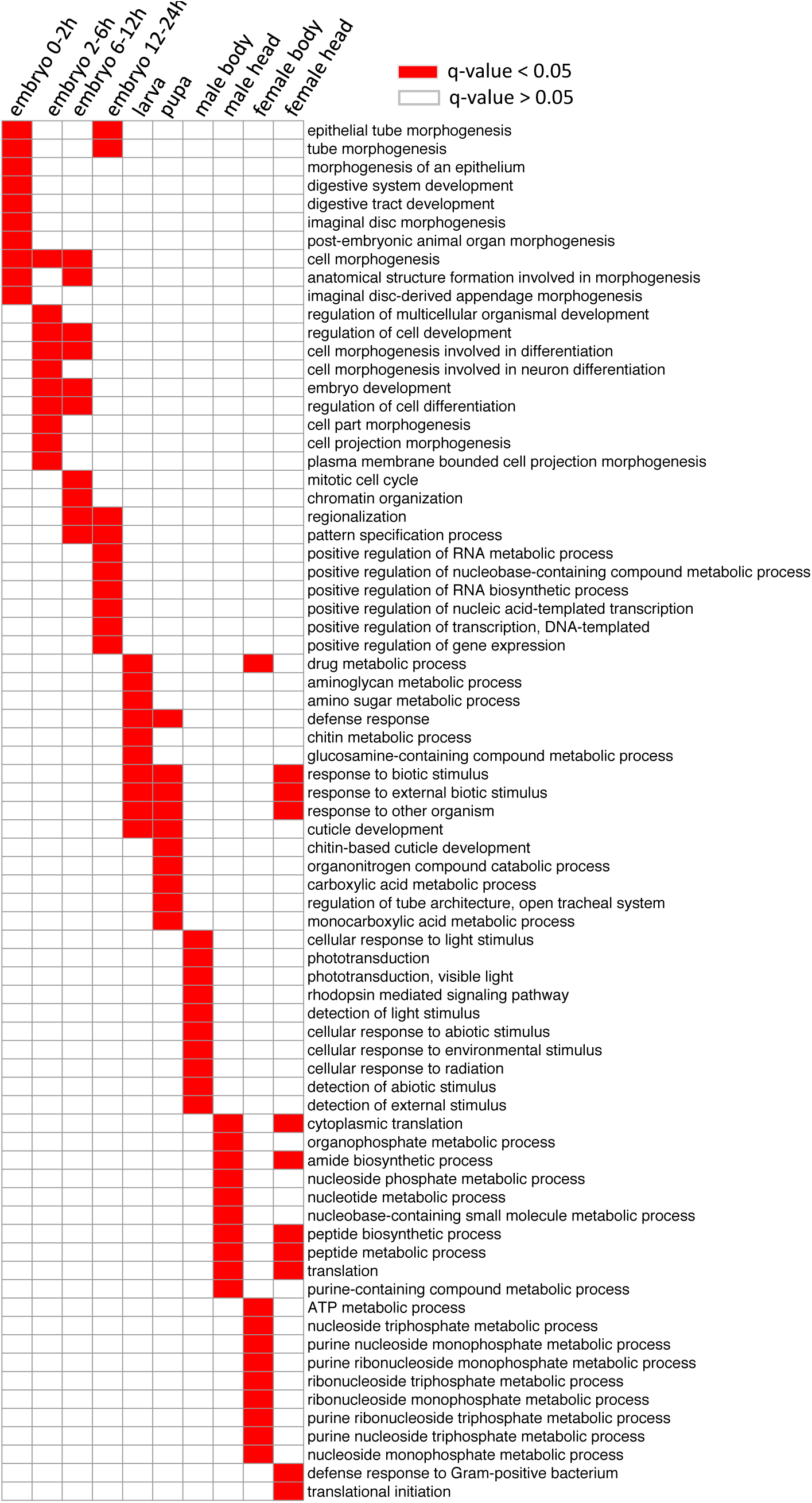
Gene ontology analysis of the genes with *log*_2_(*β*_*c*_) ≠ 0 in each stage and tissue. The biological process (BP) terms with q-values < 0.05 in each sample type are indicated with red, and others are indicated with white.

**Figure S18.**
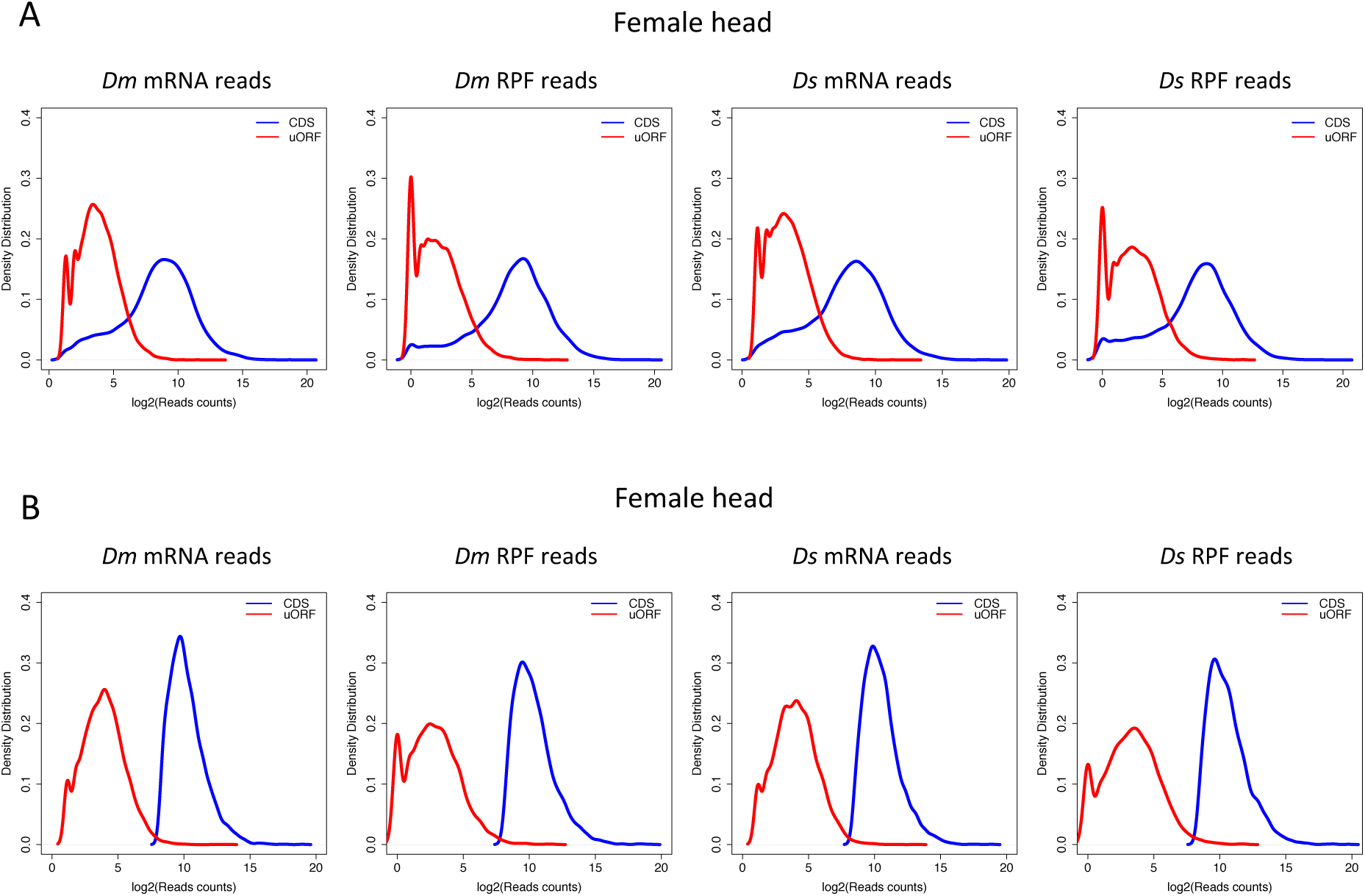
Reads count distribution of uORFs and CDSs. Distribution of mRNA reads counts and RPF counts mapped to uORFs and CDSs for all expressed uORFs (A) or only highly expressed genes (B) in female head sample. The distribution patterns were similar across other samples, and the data are not shown here. *Dm*, *D. melanogaster*; *Ds*, *D. simulans*. The reads counts were log_2_ transformed.

**Figure S19.**
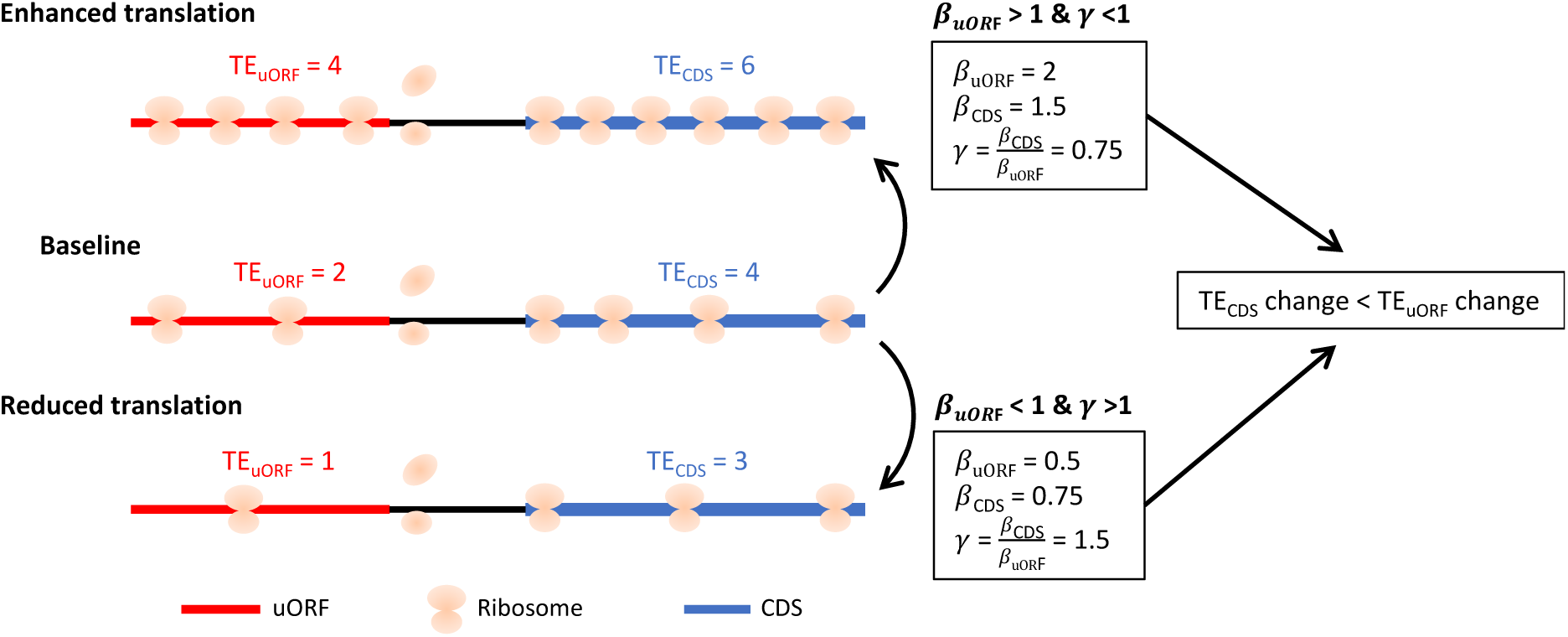
The scheme illustrating the calculation of *β*_*u*_, **c**_*c*_ and *γ*. When the cellular environment causes mRNA translation to be enhanced from baseline level, the TE_uORF_ changed from 2 to 4 (**c**_*u*_ = 2), while the TE_uORF_ changed with a smaller degree due to uORF’s buffering, from 4 to 6 (**c**_*C*_ = 1.5). This resulted in γ <1. Conversely, When the cellular environment causes mRNA translation to be reduced from the baseline level, the TE_uORF_ changed from 2 to 1 (**c**_*u*_ = 0.5), while the TE_uORF_ changed with a smaller degree due to uORF’s buffering, from 4 to 3 (**c**_*C*_ = 0.76). This resulted in γ >1. Overall, both **c**_*u*_ >1 &γ <1, and **c**_*u*_ <1 &γ >1, indicated the existence of uORFs’ buffering.

**Figure S20.**
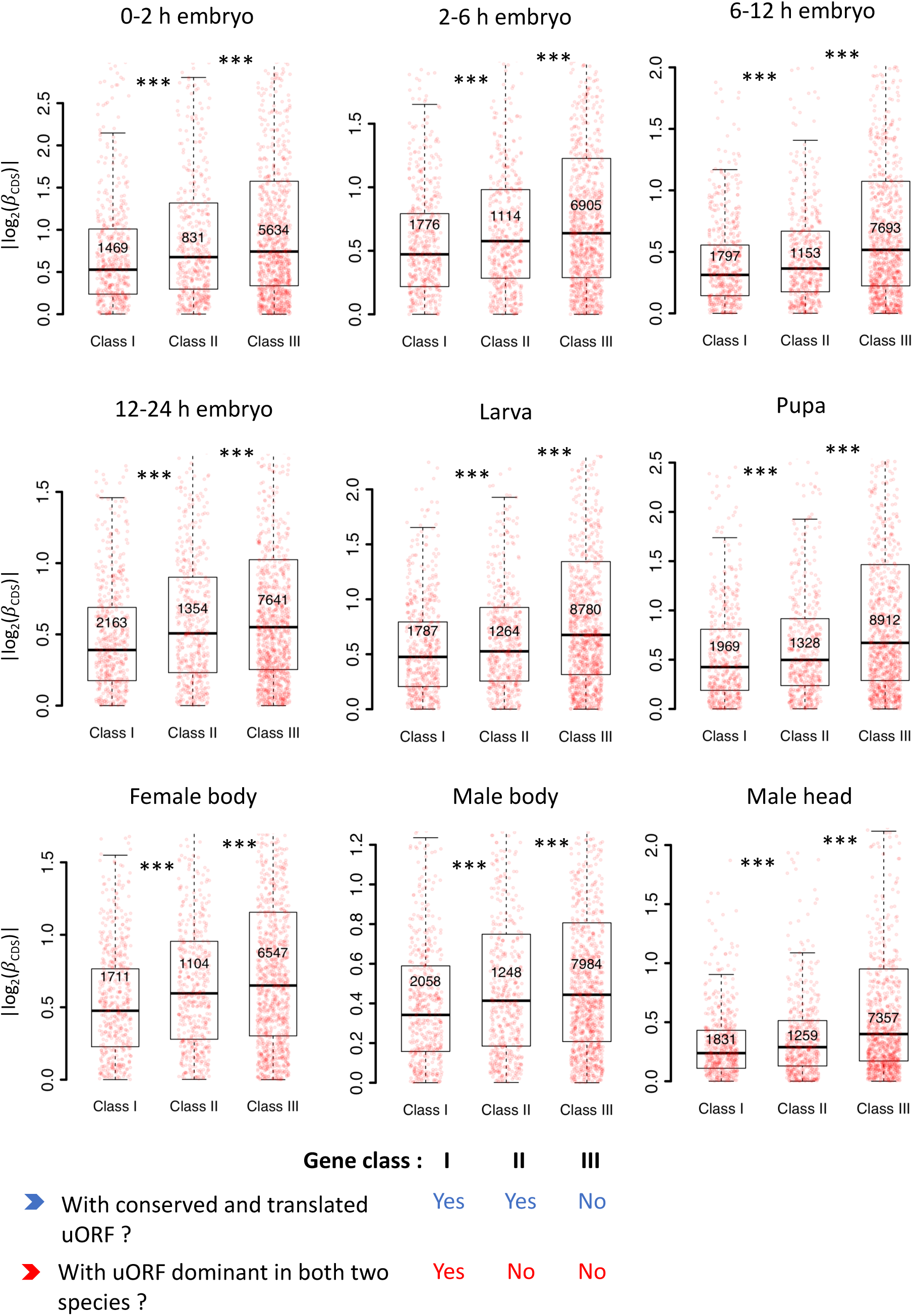
Conserved and dominantly translated uORFs showed the stronger buffering effect. Genes expressed in different stages/tissues (mRNA RPKM > 0.1 in both species) were classified into three classes according to whether a gene had a conserved and dominantly translated uORF or not. Boxplots showing interspecific CDS TE variability |*log*_2_(**c**_*c*_)| of different gene classes. *P* values were calculated using Wilcoxon rank-sum tests between the neighboring groups. ***, *P* < 0.001.

**Figure S21.**
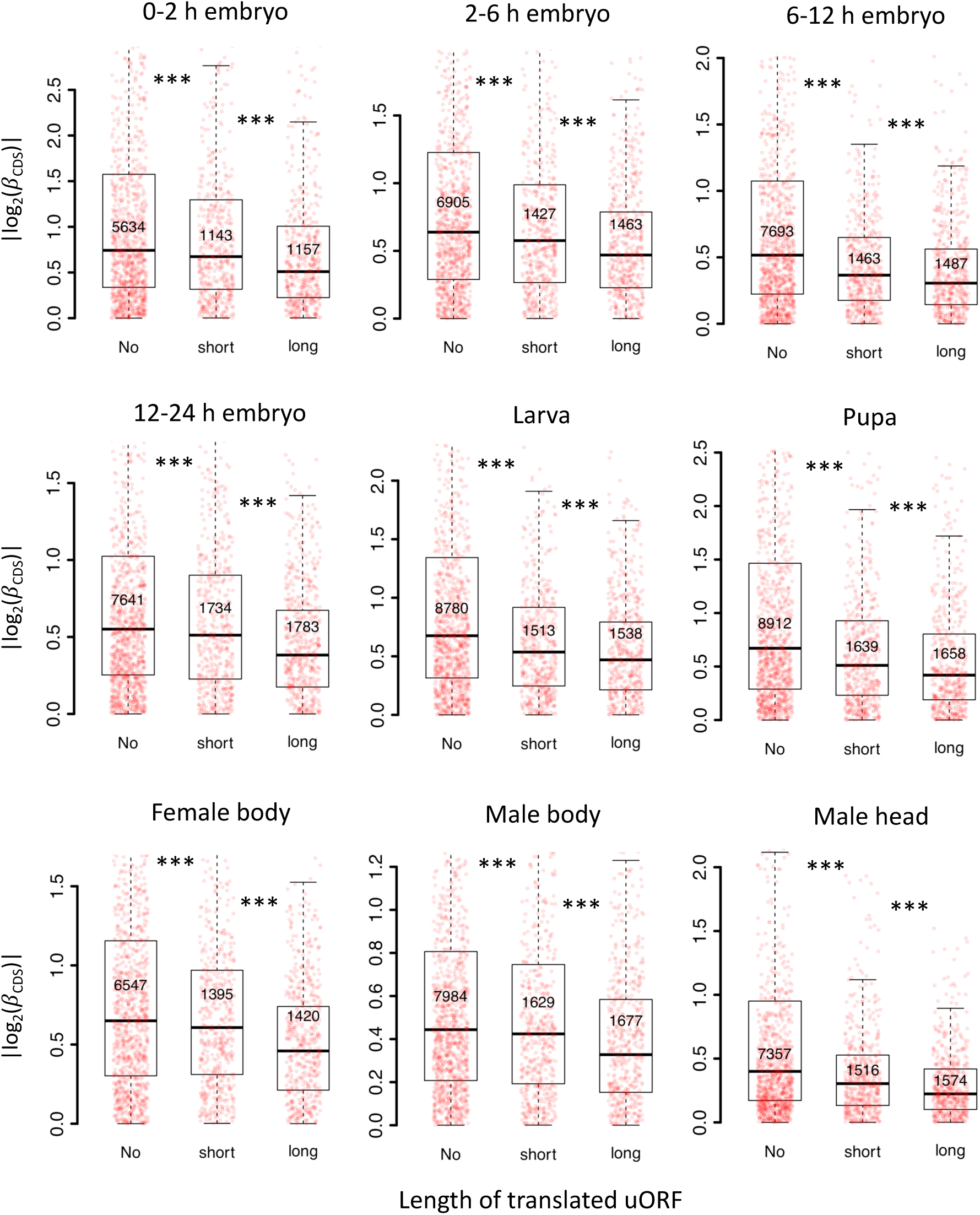
Longer uORFs showed the stronger buffering effect. Genes expressed in different stages/tissues (mRNA RPKM > 0.1 in both species) were classified into three classes according to the length of translated uORFs. Boxplots showing interspecific CDS TE variability |*log*_2_(**c**_*c*_)| of different gene classes. *P* values were calculated using Wilcoxon rank-sum tests between the neighboring groups. ***, *P* < 0.001.

**Figure S22.**
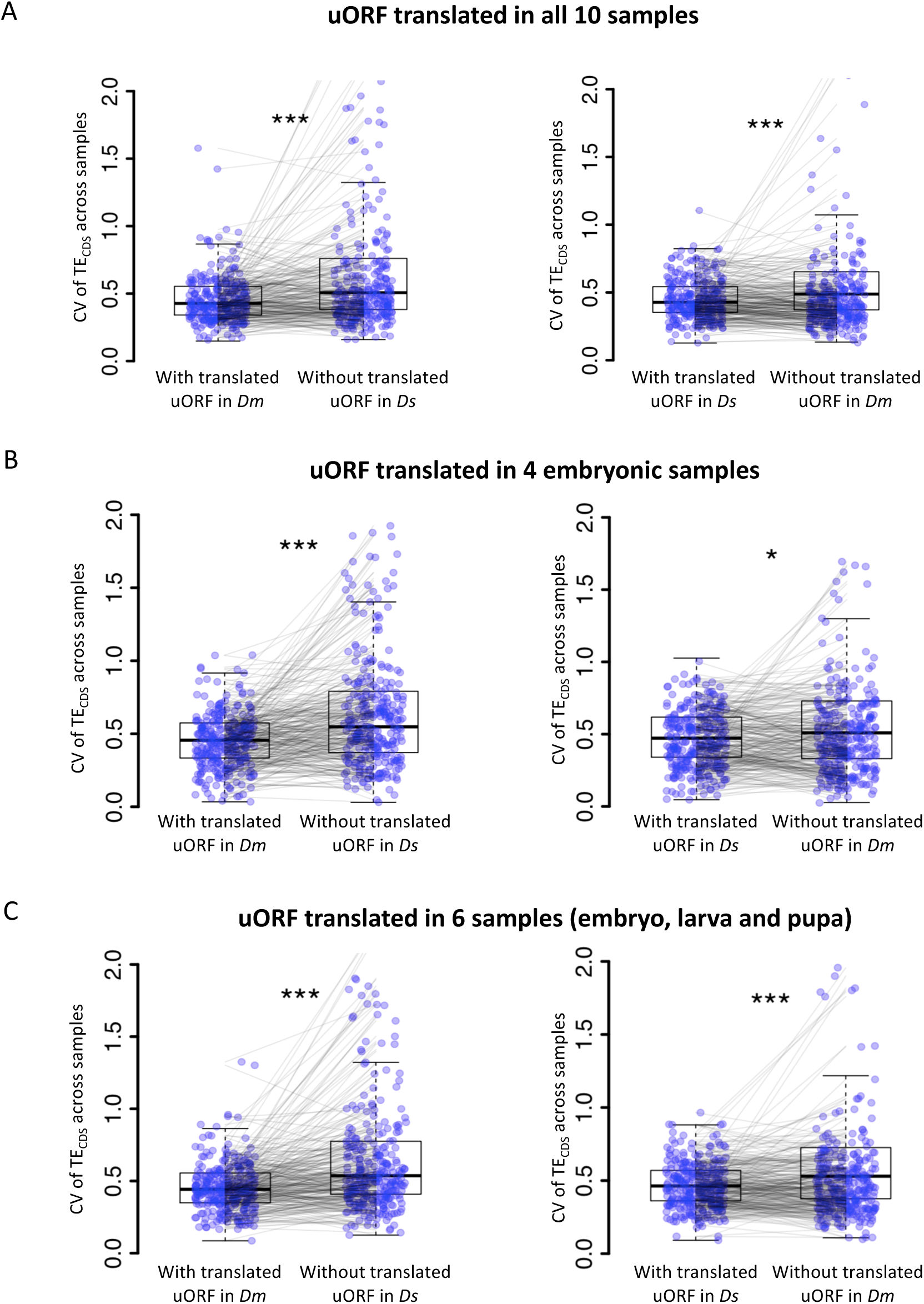
uORFs reduce CDS translational fluctuation during *Drosophila* development under different cutoffs on defining “translated uORFs”. The CV of TE_CDS_ across *Dm* (*D. melanogaster*) samples and *Ds* (*D. simulans*) samples was shown as boxplots. The selected genes harbor the translated uORFs in one of *Drosophila* species but its homologous gene without translated uORFs in another *Drosophila* species. Each pair of dots linked by a gray line represents a pair of homologous genes in *Dm* and *Ds*. (A) Translated uORF was defined as uORF translated (TE > 0.1) in all 10 samples. (B) Translated uORF was defined as uORF translated (TE > 0.1) in 4 embryonic stages. The CV of TE_CDS_ across these 4 stages was calculated. (C) Translated uORF was defined as uORF translated (TE > 0.1) in the 6 developmental stages including 4 embryonic stages, larva, and pupa. The CV of TE_CDS_ across these 6 stages was calculated. Wilcoxon signed-rank sum test. *, *P* < 0.05; ***, *P* < 0.001.

**Figure S23.**
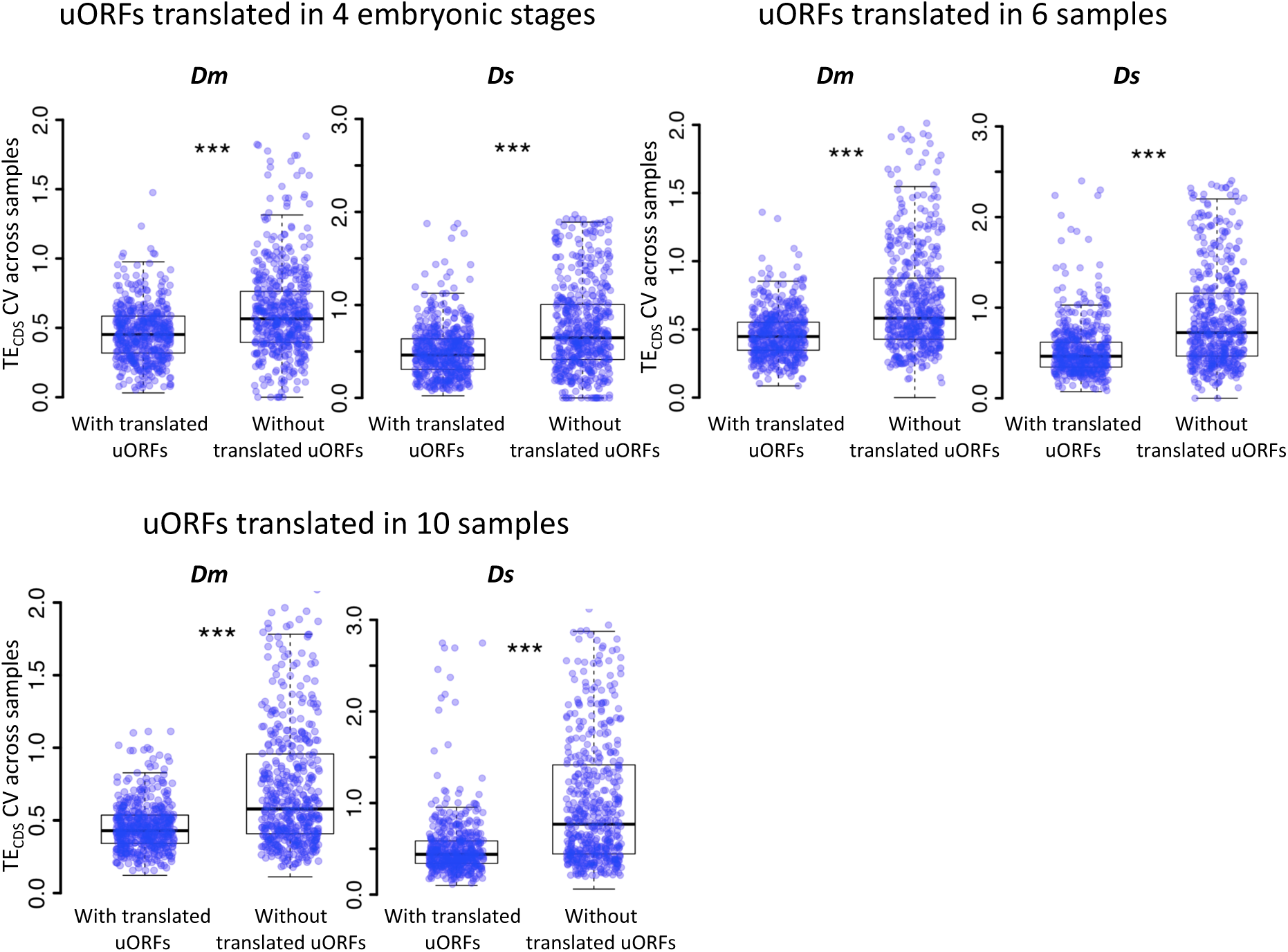
uORFs could reduce CDS translational variation during *Drosophila* development under different cutoffs on defining “translated uORFs”. Within each species, the CV of TE_CDS_ across different samples was measured for genes with translated uORFs and genes without translated uORFs, respectively. Each dot in the boxplot represents one gene. Translated uORF was defined as uORF translated (TE > 0.1) in at least 4 embryonic stages, six developmental stages (left) or translated or in all 10 samples. Wilcoxon rank-sum tests. ***, *P* < 0.001. *Dm*, *D. melanogaster*; *Ds*, *D. simulans*.

**Figure S24.**
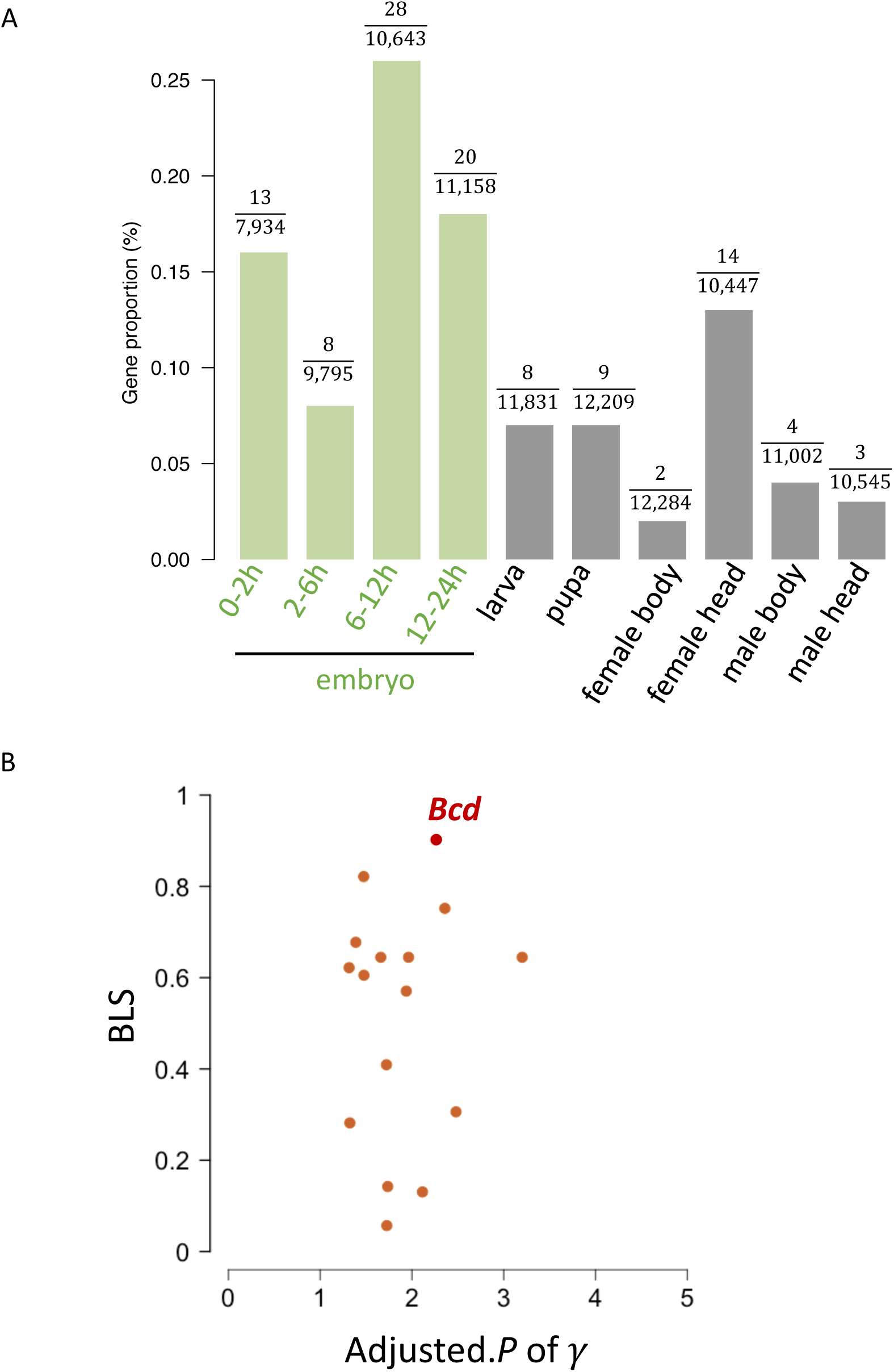
(A) The proportion (%) of genes with buffering uORFs in all expressed genes across developmental stages/tissues. (B) The distribution of adjusted. *P* of γ (x-axis) and BLS (y-axis) of dominant uORFs in embryonic stages. The *bcd* was highlighted by red color.

**Figure S25.**
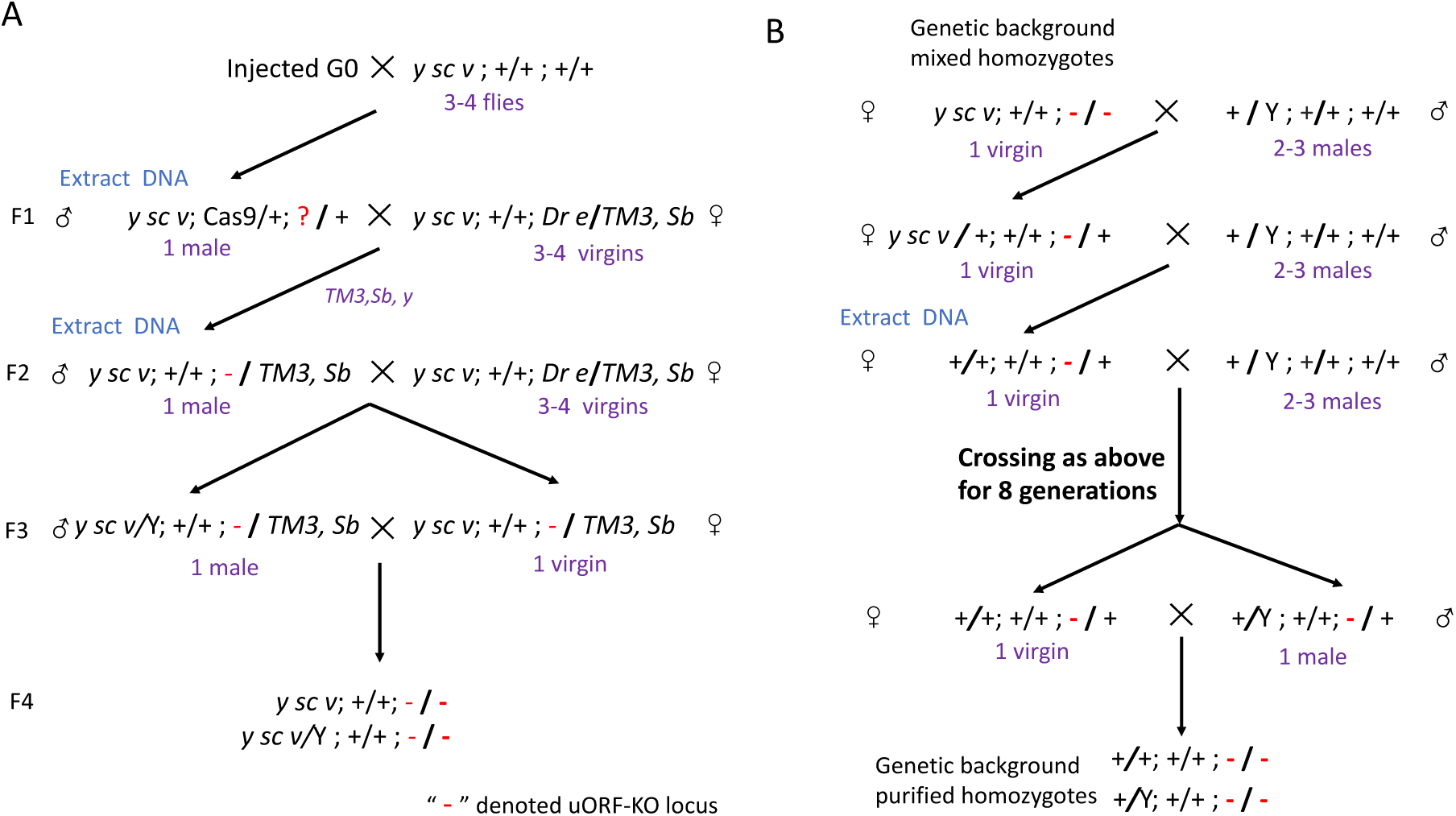
Overview of homozygous uORF-KO mutant screening. (A) Procedures for the primary screening of CRISPR/Cas9-induced mutants. Red “-” denotes the *bcd*-uORF-KO allele. (B) Purification of the genetic background of homozygous mutants by back-crossing with WT lines for nine generations.

**Figure S26.**
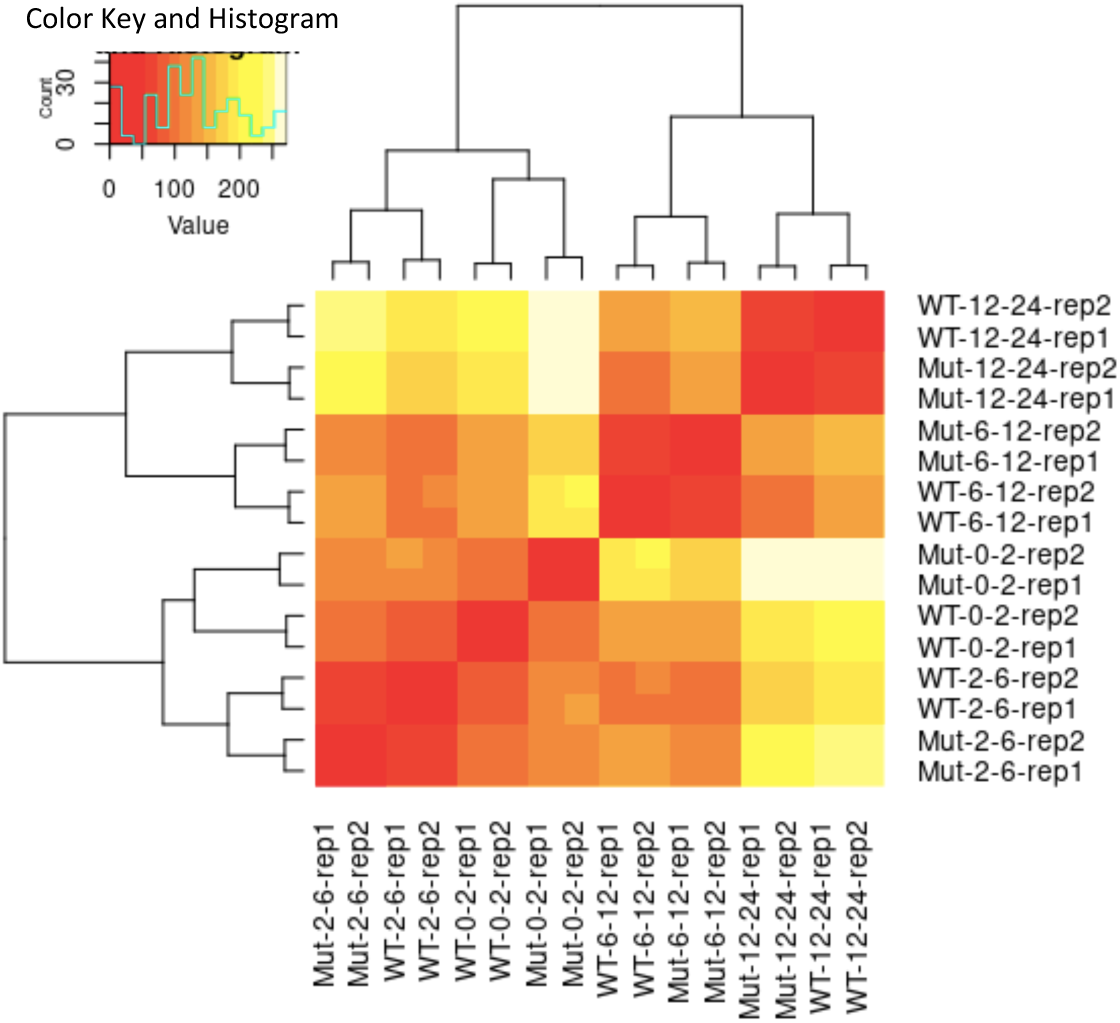
Heatmap of sample-to-sample distances of RNA-Seq libraries. Mut, mutant; WT, wild-type; rep1/rep2, biological replicate 1/biological replicate 2.

**Figure S27.**
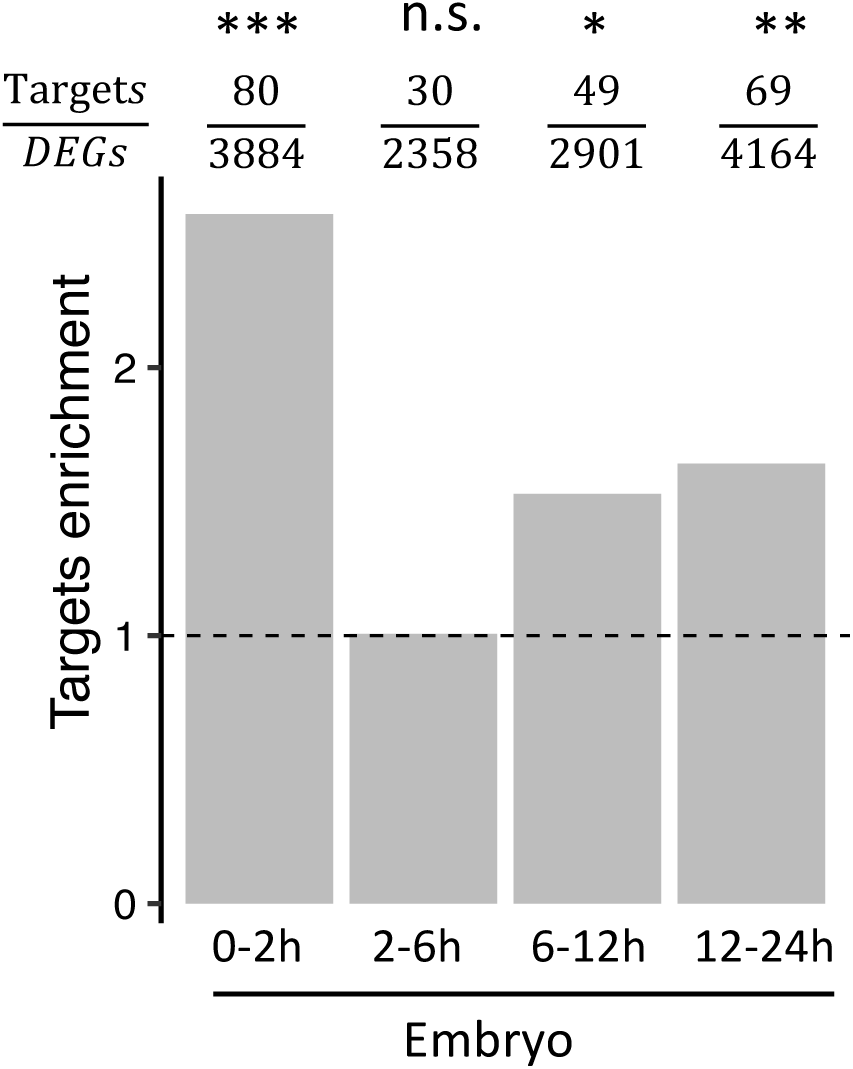
(A) The enrichment of Bcd targets in each embryo stage at 29°C. Enrichment was calculated as the ratio of the Bcd target proportion in DEGs to the Bcd target proportion in non-DEGs. *χ*^2^ tests were performed to compare the difference in Bcd target distribution in DEGs and non-DEGs. The number of DEGs and the number of Bcd targets in DEGs for each stage are provided above the bar. Asterisks indicate statistical significance (*χ*^2^ tests; *, *P* < 0.05; **, *P* < 0.01; ***, *P* < 0.001; n.s., *P* > 0.05). (B) The expression [log_2_(fold change)] levels of *bcd* and 3 of its targets, *Kr*, *btd*, and *otd,* in mutants compared to those in the WT in 4 embryonic stages. Asterisks are present at the top of bars only for |fold-changes| > 1.5 and adjusted *P* values < 0.05 (*, *P* < 0.05; **, *P* < 0.01; ***, *P* < 0.001). (C) Expression of *bcd* and its targets (*Kr*, *btd*, *otd*) measured by RNA-Seq in uKO2/uKO2 mutant (red line) and WT (blue line) of *D. melanogaster*. The expression of each gene in each stage is shown as the RPKM value.

**Figure S28.**
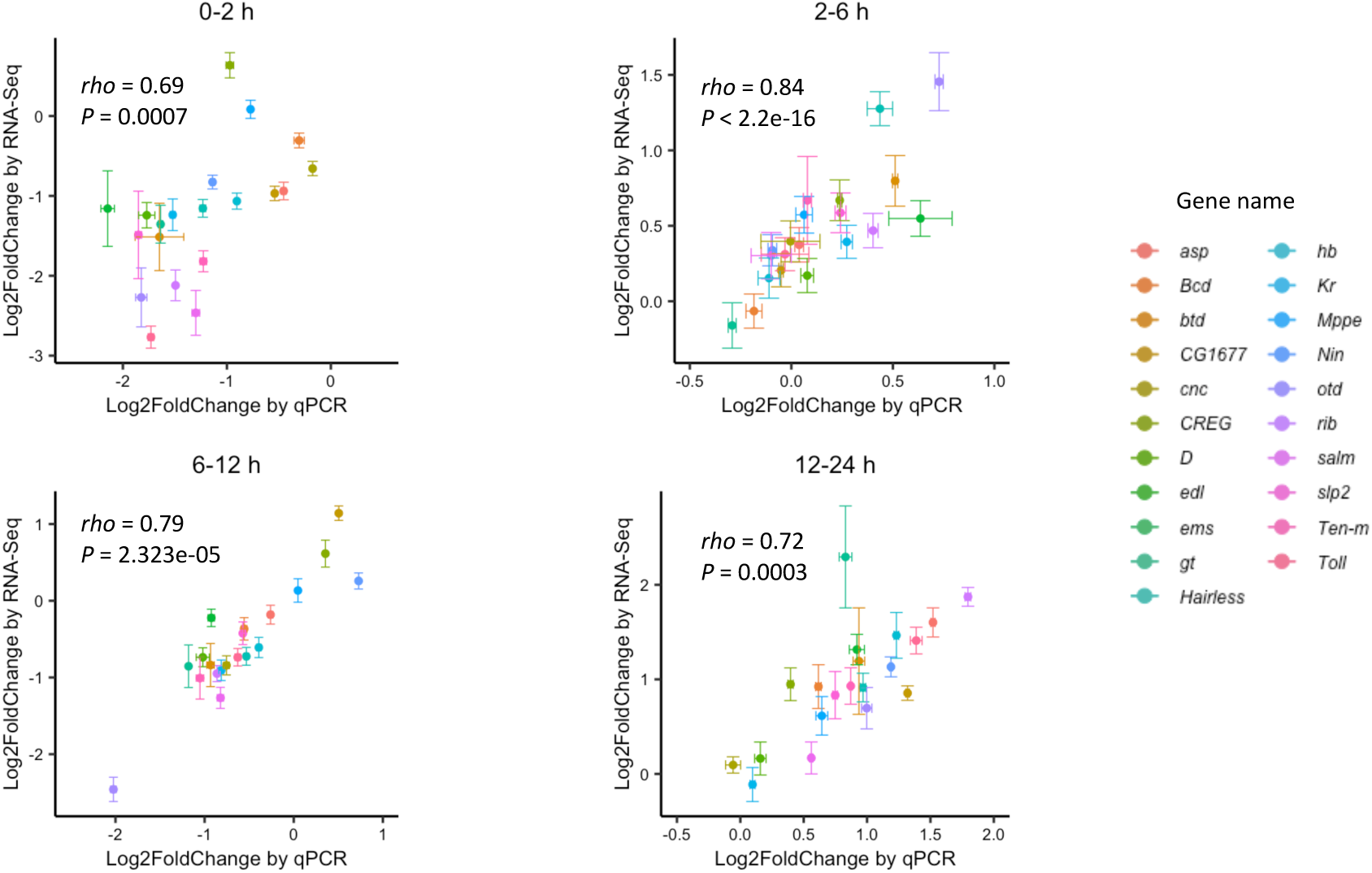
The correlation of expression changes (log_2_FoldChange) between RNA-Seq (y-axis) and RT-qPCR (x-axis) of 20 *bcd* targets in four embryo stages in uKO2/uKO2 mutant compared to WT.

**Figure S29.**
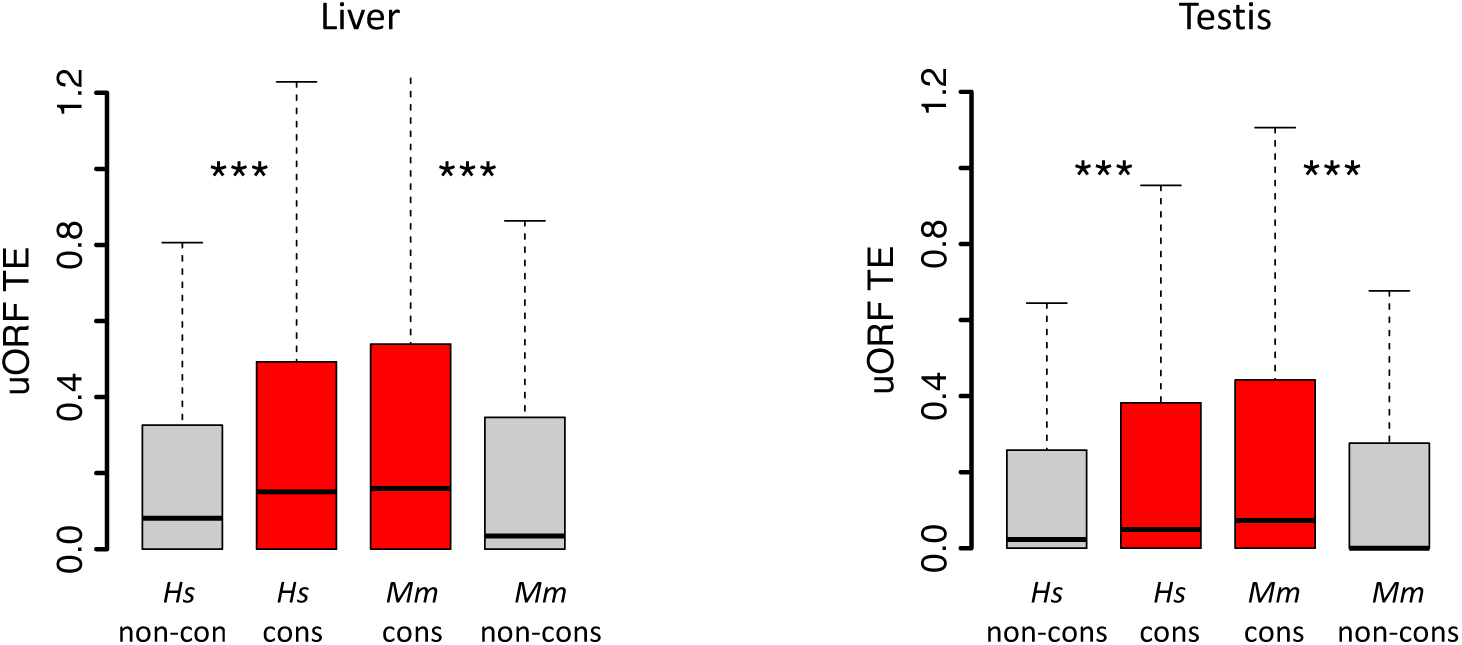
Boxplots showing the TEs of conserved and species-specific uORFs in the liver and testis of *H. sapiens* and *M. mulatta*. Wilcoxon rank-sum tests. ***, *P* < 0.001.

**Figure S30.**
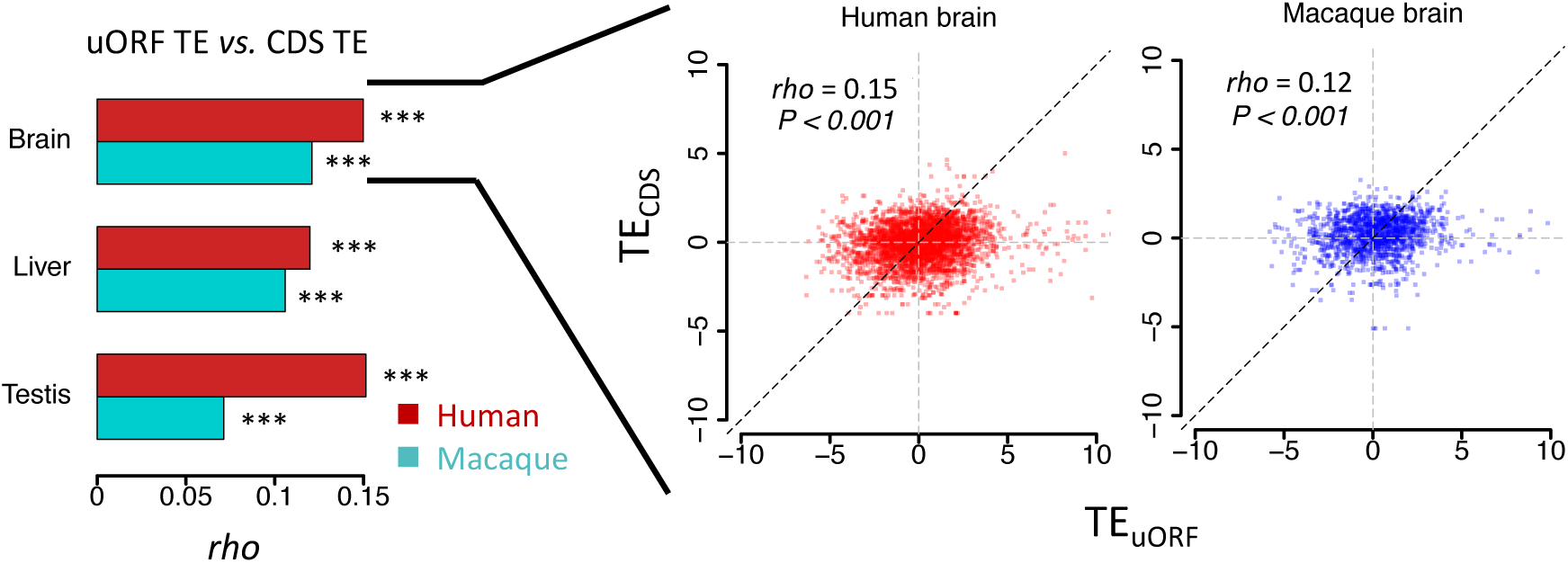
Correlations of uORF TE and corresponding CDS TE in 3 tissues of *H. sapiens* and *M. mulatta.* The bars represent Spearman’s correlation coefficient (*rho*). In all samples, we obtained *P* values < 0.001. Data from brains are shown as examples in the right panel.

**Figure S31.**
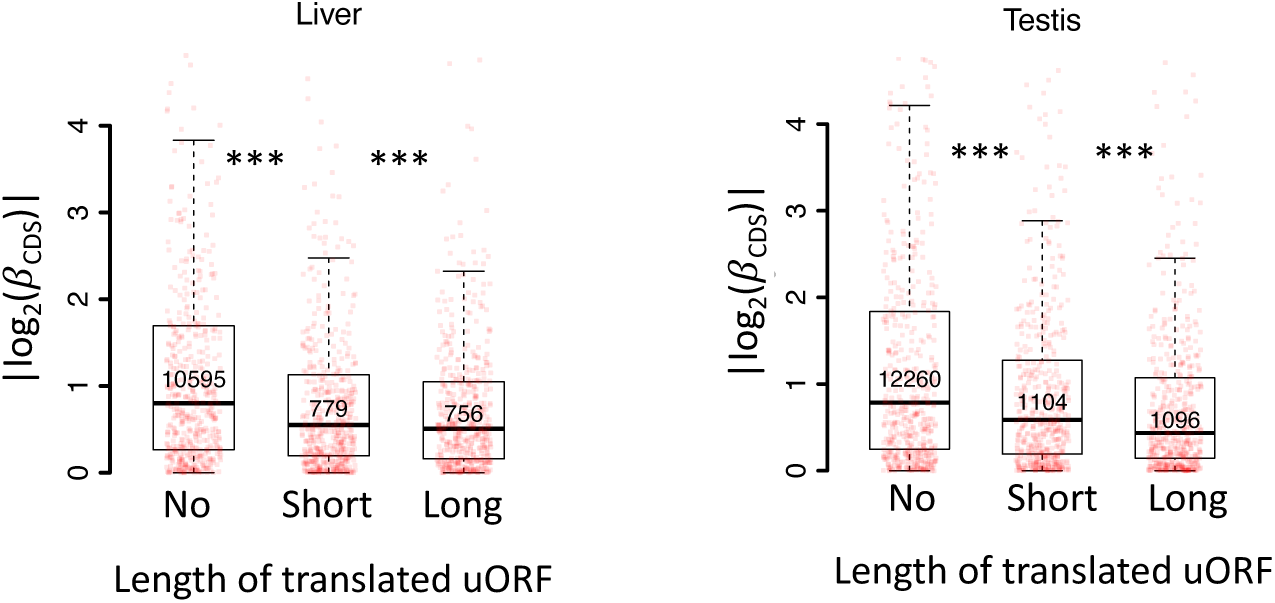
Boxplots showing interspecific CDS TE variability |*log*_2_(**c**_*CCS*_)| for three gene groups according to the length of their translated uORFs (No, short, long) in the liver (left) and testis (testis). Wilcoxon rank-sum tests. ***, *P* < 0.001.

**Figure S32.**
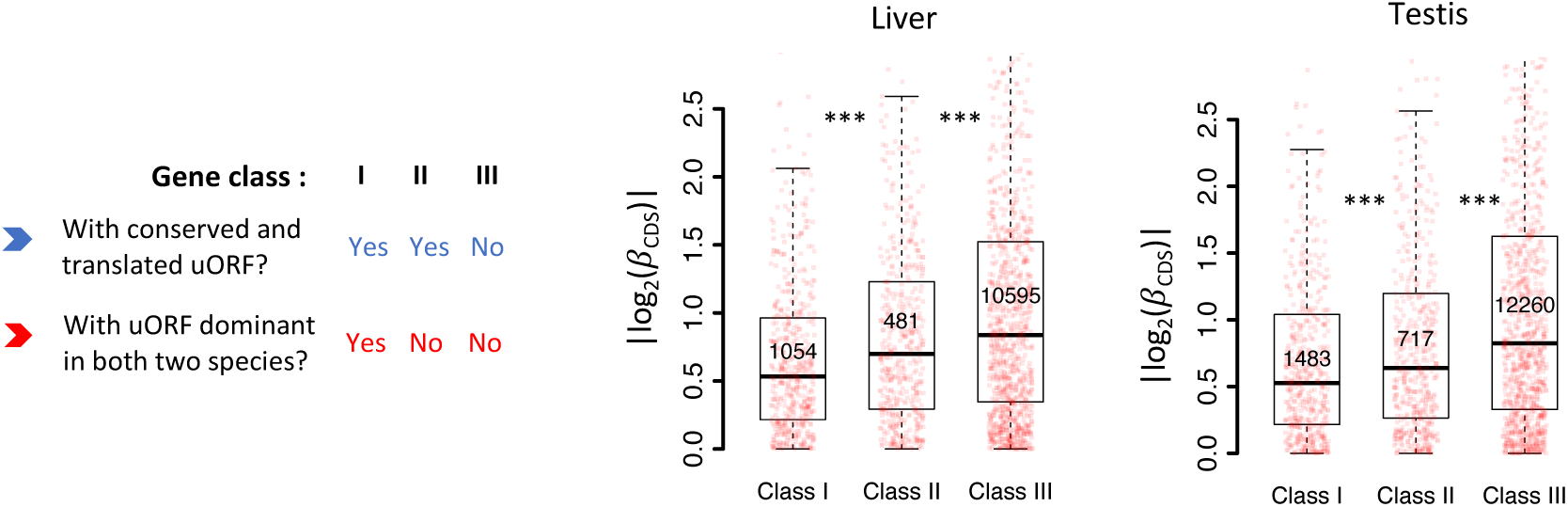
Boxplots showing interspecific CDS TE variability |*log*_2_(**c**_*CCS*_)| for different gene classes in the liver and testis. Genes expressed in liver or testis (mRNA RPKM > 0.1 in both species) were classified into three classes according to whether a gene had a conserved and dominant translated uORF (TE > 0.1) in both species or not. Wilcoxon rank-sum tests. ***, *P* < 0.001.

**Figure S33.**
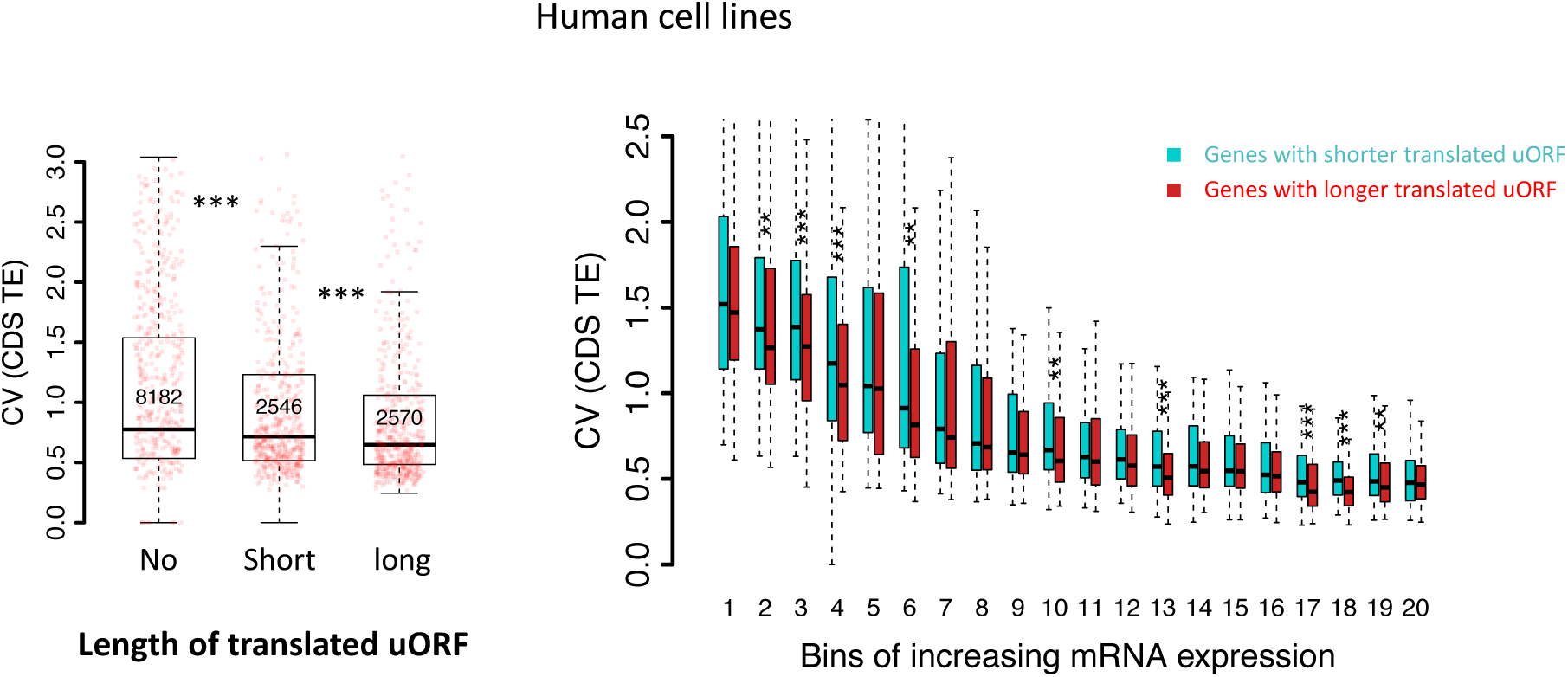
Left panel: Boxplot showing the coefficients of variation (CVs) of CDS TEs among 69 human lymphoblastoid cell lines (LCLs) for three gene groups. Genes were classified into 3 groups according to the total length of translated uORFs. Right panel: boxplot showing the coefficients of variation (CVs) of CDS TEs among the human cell lines. Genes with translated uORFs were divided into 20 bins with increasing mRNA expression levels. In each bin, genes were divided into two fractions according to the total length of translated uORFs.

**Table S1.**
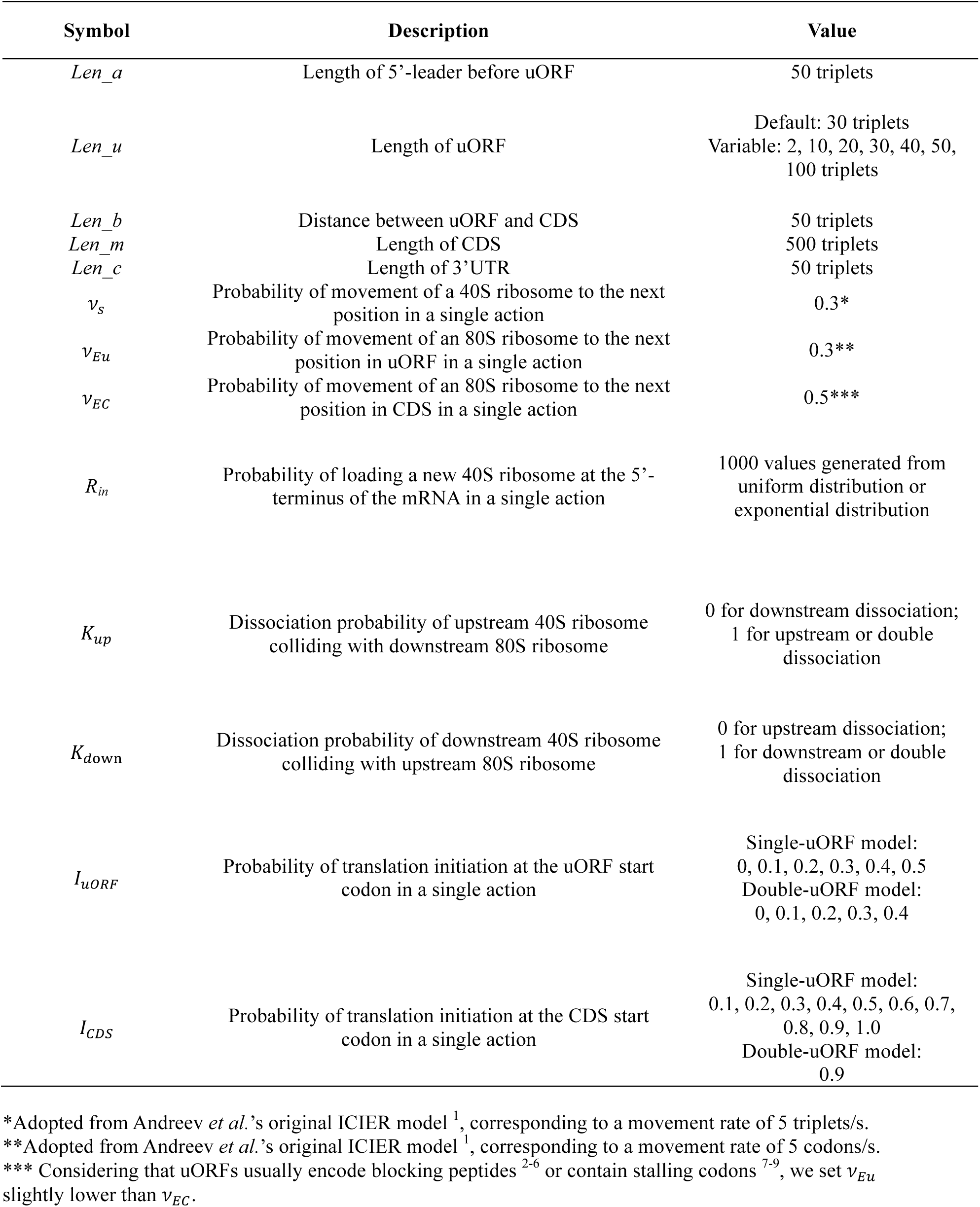
Parameters used in our simulation.

**Table S2.**
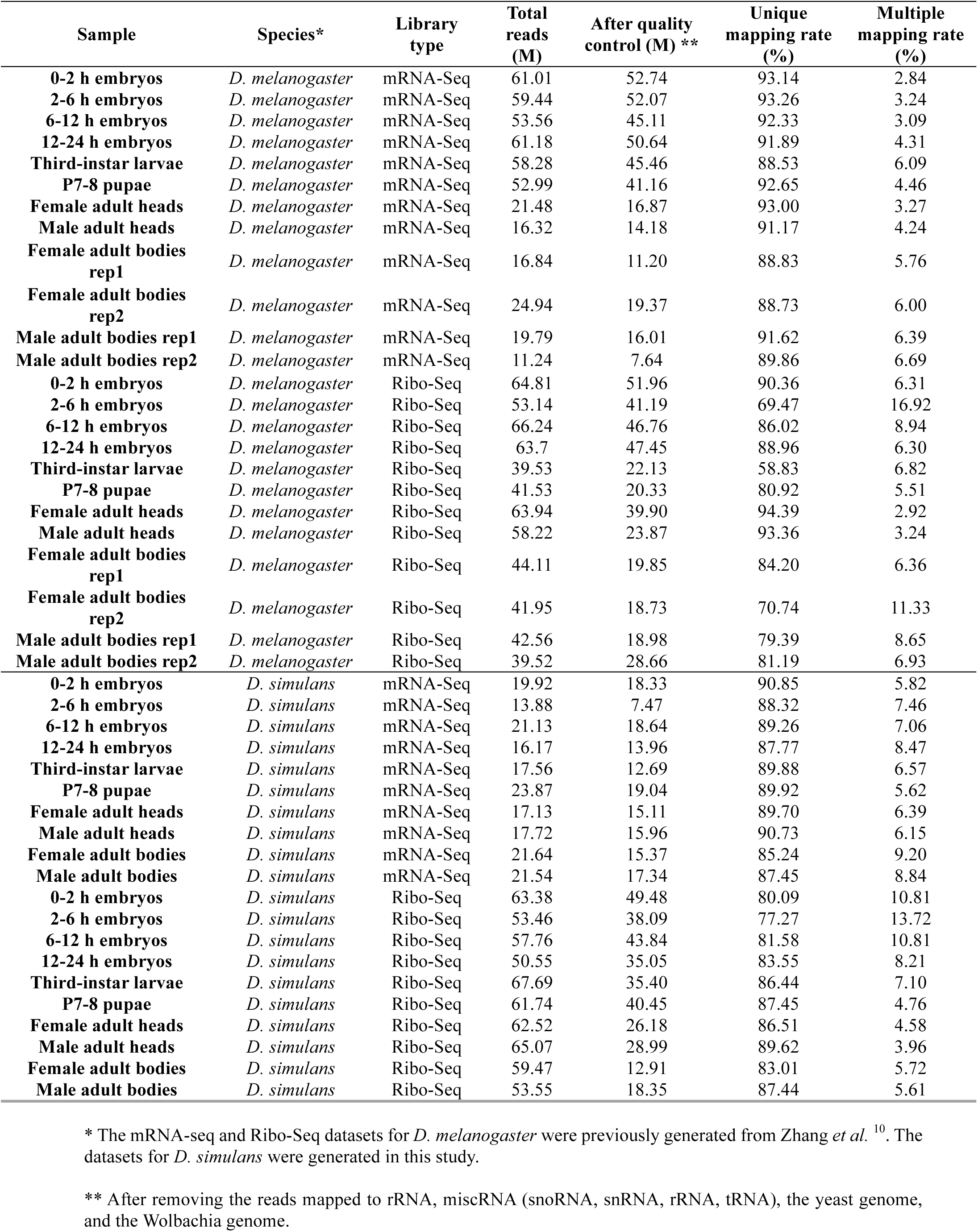
Mapping statistics of Ribo-Seq and matched mRNA-Seq libraries.

**Table S3.** Genes with uORFs showing strong evidence of translational buffering. (see TableS3_strong_buffer_uORFs.csv).

**Table S4.**
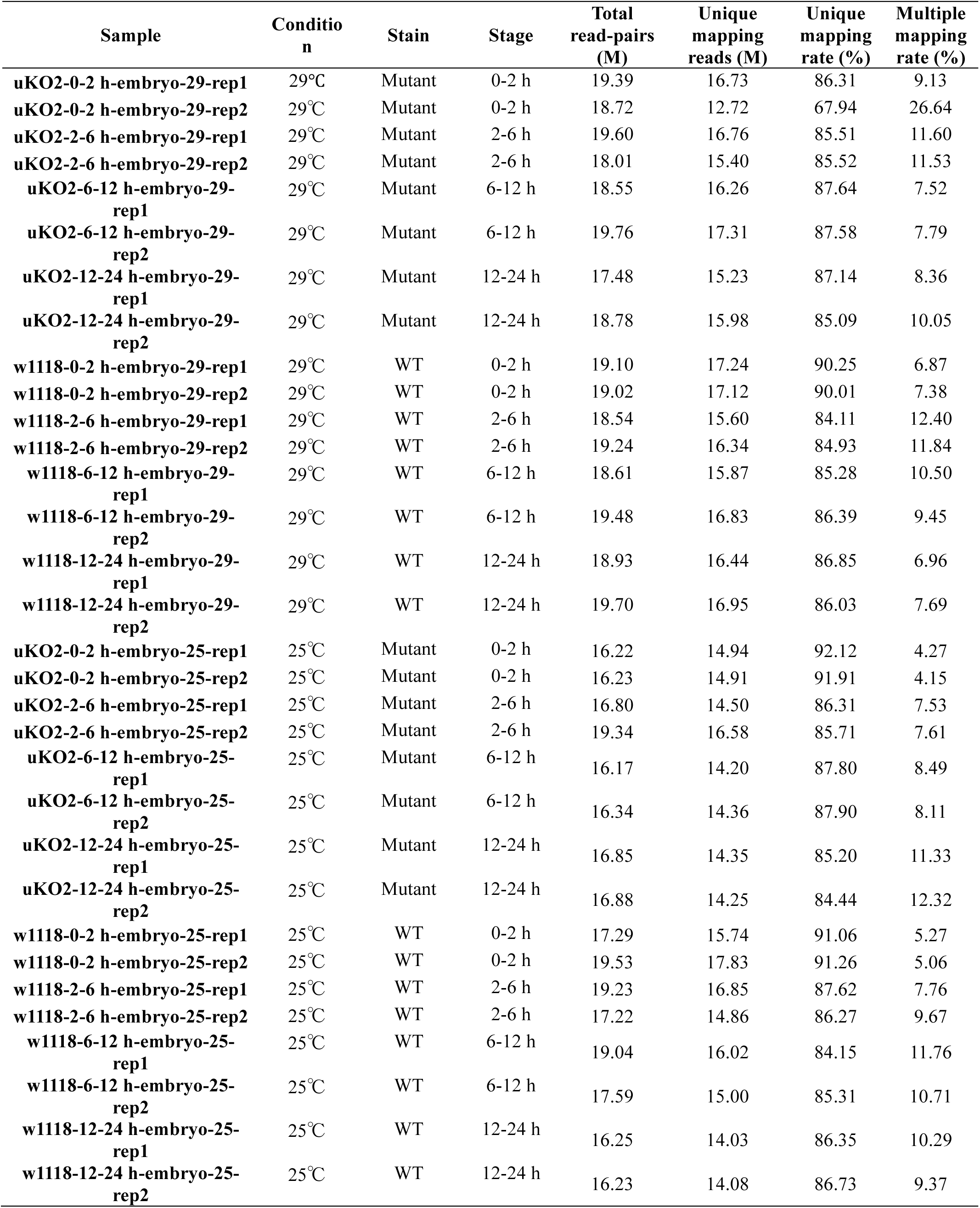
Mapping statistics of mRNA-Seq libraries for *bcd* uKO2/uKO2 mutant and WT flies.

**Table S5.**
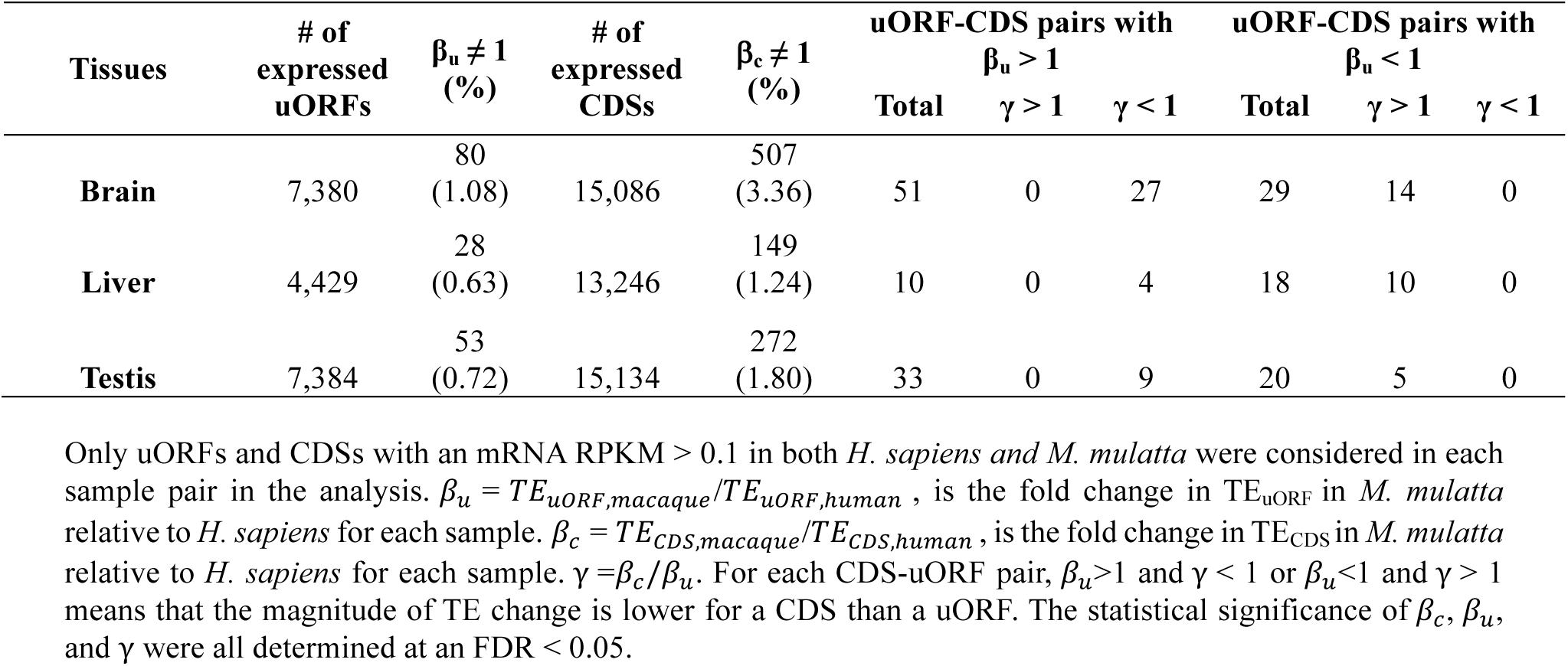
Numbers of genes showing different magnitudes of TE changes between uORFs and CDS at the interspecific level, *H. sapiens* and *M. mulatta*.

**Table S6.**
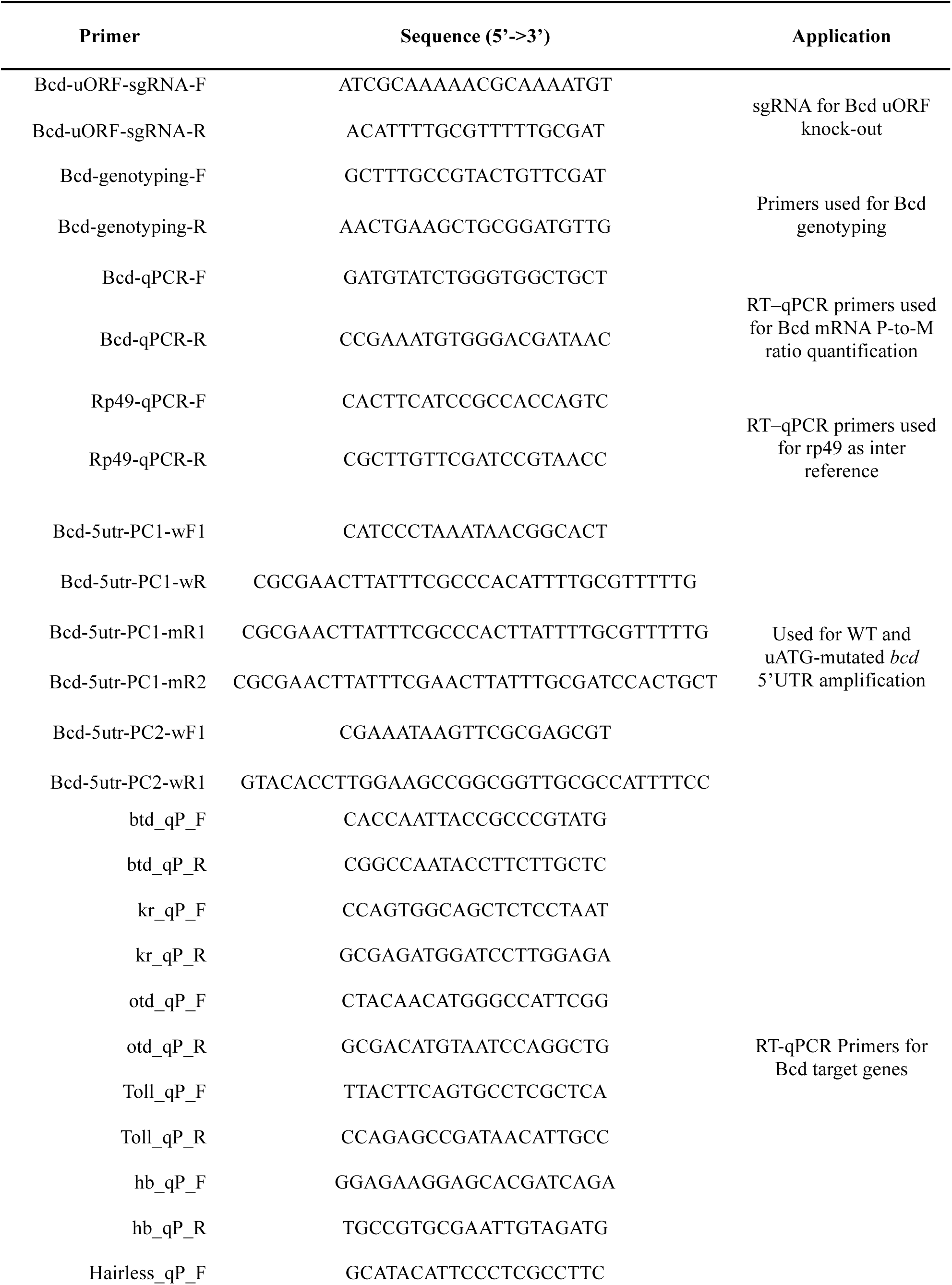
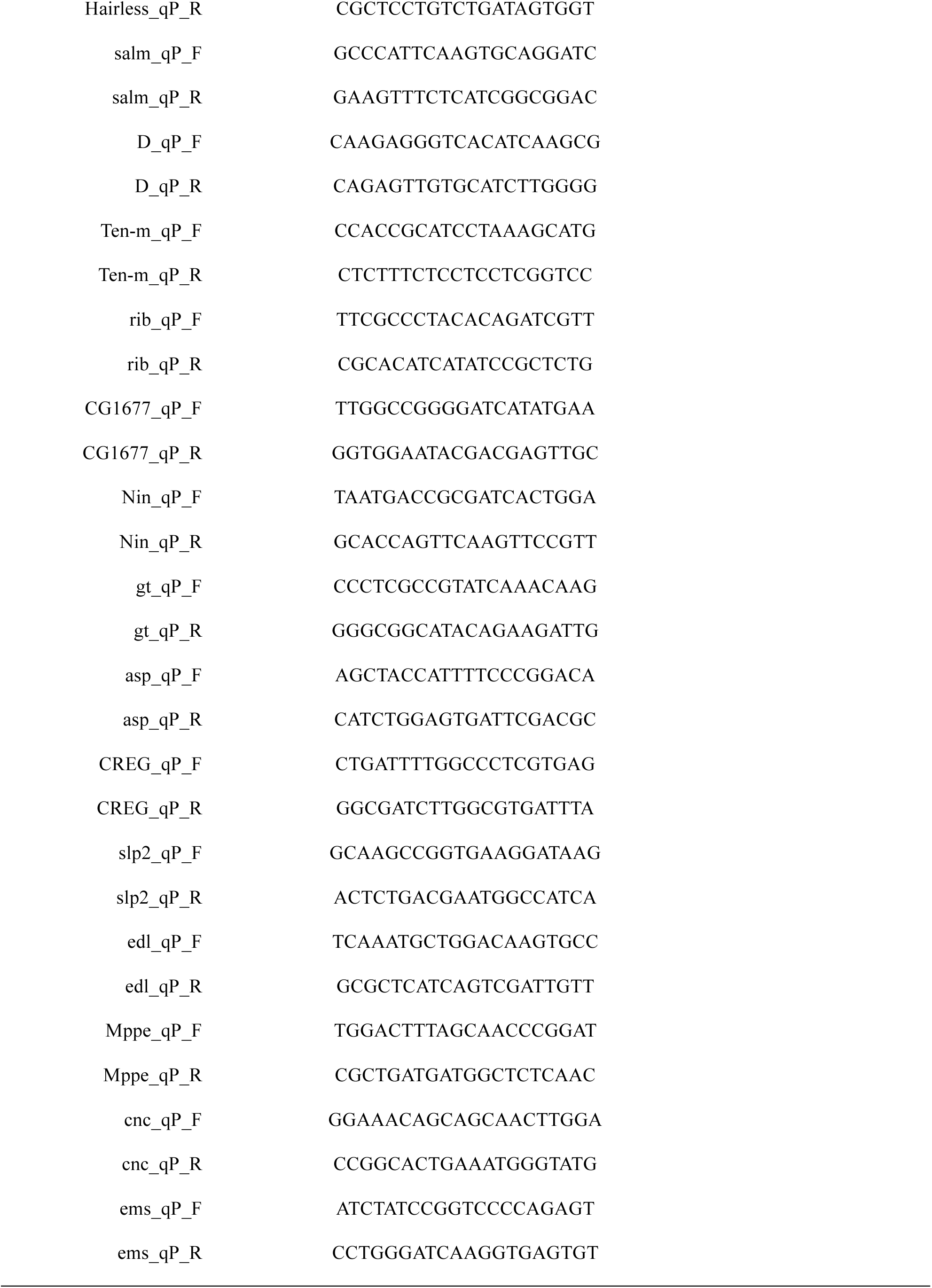
The primer sequences used in this study.

## Notes

https://github.com/lujlab/Buffer_eLife2025

### Competing Interest Statement

The authors have declared no competing interest.

### Summary of Updates

Discussion Section have been expanded; Figure 3 revised; Figure 9 added; Supplemental files updated.

## References

1. Buccitelli, C., and Selbach, M. (2020). mRNAs, proteins and the emerging principles of gene expression control. Nature Reviews Genetics 21, 630–644. 10.1038/s41576-020-0258-4.

2. Lee, T.I., and Young, R.A. (2013). Transcriptional regulation and its misregulation in disease. Cell 152, 1237–1251. 10.1016/j.cell.2013.02.014.

3. MacNeil, L.T., and Walhout, A.J.M. (2011). Gene regulatory networks and the role of robustness and stochasticity in the control of gene expression. Genome Research 21, 645–657. 10.1101/gr.097378.109.

4. Hill, M.S., Vande Zande, P., and Wittkopp, P.J. (2021). Molecular and evolutionary processes generating variation in gene expression. Nat Rev Genet 22, 203–215. 10.1038/s41576-020-00304-w.

5. Vogel, C. (2013). Protein Expression Under Pressure. Science 342, 1052–1053. 10.1126/science.1247833.

6. Signor, S.A., and Nuzhdin, S.V. (2018). The Evolution of Gene Expression in cis and trans. Trends Genet 34, 532–544. 10.1016/j.tig.2018.03.007.

7. Artieri, C.G., and Fraser, H.B. (2014). Evolution at two levels of gene expression in yeast. Genome Res 24, 411–421. 10.1101/gr.165522.113.

8. McManus, C.J., May, G.E., Spealman, P., and Shteyman, A. (2014). Ribosome profiling reveals post-transcriptional buffering of divergent gene expression in yeast. Genome Res 24, 422–430. 10.1101/gr.164996.113.

9. Wang, Z., Sun, X., Zhao, Y., Guo, X., Jiang, H., Li, H., and Gu, Z. (2015). Evolution of gene regulation during transcription and translation. Genome Biol Evol 7, 1155–1167. 10.1093/gbe/evv059.

10. Khan, Z., Ford, M.J., Cusanovich, D.A., Mitrano, A., Pritchard, J.K., and Gilad, Y. (2013). Primate transcript and protein expression levels evolve under compensatory selection pressures. Science 342, 1100–1104. 10.1126/science.1242379.

11. Wang, S.H., Hsiao, C.J., Khan, Z., and Pritchard, J.K. (2018). Post-translational buffering leads to convergent protein expression levels between primates. Genome Biol 19, 83. 10.1186/s13059-018-1451-z.

12. Schrimpf, S.P., Weiss, M., Reiter, L., Ahrens, C.H., Jovanovic, M., Malmström, J., Brunner, E., Mohanty, S., Lercher, M.J., Hunziker, P.E., et al. (2009). Comparative functional analysis of the *Caenorhabditis elegans* and *Drosophila melanogaster* proteomes. PLoS Biol 7, e48. 10.1371/journal.pbio.1000048.

13. Kusnadi, E.P., Timpone, C., Topisirovic, I., Larsson, O., and Furic, L. (2022). Regulation of gene expression via translational buffering. Biochim Biophys Acta Mol Cell Res 1869, 119140. 10.1016/j.bbamcr.2021.119140.

14. Laurent, J.M., Vogel, C., Kwon, T., Craig, S.A., Boutz, D.R., Huse, H.K., Nozue, K., Walia, H., Whiteley, M., Ronald, P.C., and Marcotte, E.M. (2010). Protein abundances are more conserved than mRNA abundances across diverse taxa. Proteomics 10, 4209–4212. 10.1002/pmic.201000327.

15. Teixeira, F.K., and Lehmann, R. (2019). Translational Control during Developmental Transitions. Cold Spring Harb Perspect Biol 11. 10.1101/cshperspect.a032987.

16. Jackson, R.J., Hellen, C.U., and Pestova, T.V. (2010). The mechanism of eukaryotic translation initiation and principles of its regulation. Nat Rev Mol Cell Biol 11, 113–127. 10.1038/nrm2838.

17. Sonenberg, N., and Hinnebusch, A.G. (2009). Regulation of translation initiation in eukaryotes: mechanisms and biological targets. Cell 136, 731–745. 10.1016/j.cell.2009.01.042.

18. Zhang, H., Wang, Y., Wu, X., Tang, X., Wu, C., and Lu, J. (2021). Determinants of genome-wide distribution and evolution of uORFs in eukaryotes. Nature Communications 12, 1076. 10.1038/s41467-021-21394-y.

19. Zhang, H., Wang, Y., and Lu, J. (2019). Function and Evolution of Upstream ORFs in Eukaryotes. Trends Biochem Sci 44, 782–794. 10.1016/j.tibs.2019.03.002.

20. Neafsey, D.E., and Galagan, J.E. (2007). Dual modes of natural selection on upstream open reading frames. Mol Biol Evol 24, 1744–1751. 10.1093/molbev/msm093.

21. Churbanov, A., Rogozin, I.B., Babenko, V.N., Ali, H., and Koonin, E.V. (2005). Evolutionary conservation suggests a regulatory function of AUG triplets in 5’-UTRs of eukaryotic genes. Nucleic Acids Res 33, 5512–5520. 10.1093/nar/gki847.

22. Resch, A.M., Ogurtsov, A.Y., Rogozin, I.B., Shabalina, S.A., and Koonin, E.V. (2009). Evolution of alternative and constitutive regions of mammalian 5’UTRs. BMC Genomics 10, 162. 10.1186/1471-2164-10-162.

23. Zhang, H., Dou, S., He, F., Luo, J., Wei, L., and Lu, J. (2018). Genome-wide maps of ribosomal occupancy provide insights into adaptive evolution and regulatory roles of uORFs during *Drosophila* development. PLoS biology 16, e2003903.

24. Malzer, E., Szajewska-Skuta, M., Dalton, L.E., Thomas, S.E., Hu, N., Skaer, H., Lomas, D.A., Crowther, D.C., and Marciniak, S.J. (2013). Coordinate regulation of eIF2alpha phosphorylation by PPP1R15 and GCN2 is required during *Drosophila* development. J Cell Sci 126, 1406–1415. 10.1242/jcs.117614.

25. Komonyi, O., Papai, G., Enunlu, I., Muratoglu, S., Pankotai, T., Kopitova, D., Maroy, P., Udvardy, A., and Boros, I. (2005). DTL, the *Drosophila* homolog of PIMT/Tgs1 nuclear receptor coactivator-interacting protein/RNA methyltransferase, has an essential role in development. J Biol Chem 280, 12397–12404. 10.1074/jbc.M409251200.

26. Medenbach, J., Seiler, M., and Hentze, M.W. (2011). Translational control via protein-regulated upstream open reading frames. Cell 145, 902–913. 10.1016/j.cell.2011.05.005.

27. Chen, J., Tresenrider, A., Chia, M., McSwiggen, D.T., Spedale, G., Jorgensen, V., Liao, H., van Werven, F.J., and Unal, E. (2017). Kinetochore inactivation by expression of a repressive mRNA. Elife 6. 10.7554/eLife.27417.

28. Cheng, Z., Otto, G.M., Powers, E.N., Keskin, A., Mertins, P., Carr, S.A., Jovanovic, M., and Brar, G.A. (2018). Pervasive, Coordinated Protein-Level Changes Driven by Transcript Isoform Switching during Meiosis. Cell 172, 910–923 e916. 10.1016/j.cell.2018.01.035.

29. Kurihara, Y., Makita, Y., Kawashima, M., Fujita, T., Iwasaki, S., and Matsui, M. (2018). Transcripts from downstream alternative transcription start sites evade uORF-mediated inhibition of gene expression in Arabidopsis. Proc Natl Acad Sci U S A 115, 7831–7836. 10.1073/pnas.1804971115.

30. Yang, Y.F., Zhang, X., Ma, X., Zhao, T., Sun, Q., Huan, Q., Wu, S., Du, Z., and Qian, W. (2017). Trans-splicing enhances translational efficiency in C. elegans. Genome Res 27, 1525–1535. 10.1101/gr.202150.115.

31. Wiestner, A., Schlemper, R.J., van der Maas, A.P., and Skoda, R.C. (1998). An activating splice donor mutation in the thrombopoietin gene causes hereditary thrombocythaemia. Nat Genet 18, 49–52. 10.1038/ng0198-49.

32. Liu, L., Dilworth, D., Gao, L., Monzon, J., Summers, A., Lassam, N., and Hogg, D. (1999). Mutation of the CDKN2A 5’ UTR creates an aberrant initiation codon and predisposes to melanoma. Nat Genet 21, 128–132. 10.1038/5082.

33. Wen, Y., Liu, Y., Xu, Y., Zhao, Y., Hua, R., Wang, K., Sun, M., Li, Y., Yang, S., Zhang, X.J., et al. (2009). Loss-of-function mutations of an inhibitory upstream ORF in the human hairless transcript cause Marie Unna hereditary hypotrichosis. Nat Genet 41, 228–233. 10.1038/ng.276.

34. Calvo, S.E., Pagliarini, D.J., and Mootha, V.K. (2009). Upstream open reading frames cause widespread reduction of protein expression and are polymorphic among humans. Proc Natl Acad Sci U S A 106, 7507–7512. 10.1073/pnas.0810916106.

35. Lee, D.S.M., Park, J., Kromer, A., Baras, A., Rader, D.J., Ritchie, M.D., Ghanem, L.R., and Barash, Y. (2021). Disrupting upstream translation in mRNAs is associated with human disease. Nat Commun 12, 1515. 10.1038/s41467-021-21812-1.

36. Costa-Mattioli, M., and Walter, P. (2020). The integrated stress response: From mechanism to disease. Science 368. 10.1126/science.aat5314.

37. Hinnebusch, A.G. (2005). Translational regulation of GCN4 and the general amino acid control of yeast. Annu Rev Microbiol 59, 407–450. 10.1146/annurev.micro.59.031805.133833.

38. Lu, P.D., Harding, H.P., and Ron, D. (2004). Translation reinitiation at alternative open reading frames regulates gene expression in an integrated stress response. J Cell Biol 167, 27–33. 10.1083/jcb.200408003.

39. Bohlen, J., Harbrecht, L., Blanco, S., Clemm von Hohenberg, K., Fenzl, K., Kramer, G., Bukau, B., and Teleman, A.A. (2020). DENR promotes translation reinitiation via ribosome recycling to drive expression of oncogenes including ATF4. Nat Commun 11, 4676. 10.1038/s41467-020-18452-2.

40. Vasudevan, D., Neuman, S.D., Yang, A., Lough, L., Brown, B., Bashirullah, A., Cardozo, T., and Ryoo, H.D. (2020). Translational induction of ATF4 during integrated stress response requires noncanonical initiation factors eIF2D and DENR. Nat Commun 11, 4677. 10.1038/s41467-020-18453-1.

41. Vattem, K.M., and Wek, R.C. (2004). Reinitiation involving upstream ORFs regulates ATF4 mRNA translation in mammalian cells. Proc Natl Acad Sci U S A 101, 11269–11274. 10.1073/pnas.0400541101.

42. Andreev, D.E., O’Connor, P.B., Fahey, C., Kenny, E.M., Terenin, I.M., Dmitriev, S.E., Cormican, P., Morris, D.W., Shatsky, I.N., and Baranov, P.V. (2015). Translation of 5’ leaders is pervasive in genes resistant to eIF2 repression. Elife 4, e03971. 10.7554/eLife.03971.

43. Andreev, D.E., Arnold, M., Kiniry, S.J., Loughran, G., Michel, A.M., Rachinskii, D., and Baranov, P.V. (2018). TASEP modelling provides a parsimonious explanation for the ability of a single uORF to derepress translation during the integrated stress response. Elife 7. 10.7554/eLife.32563.

44. Kozak, M. (1989). The scanning model for translation: an update. J Cell Biol 108, 229–241. 10.1083/jcb.108.2.229.

45. Johnstone, T.G., Bazzini, A.A., and Giraldez, A.J. (2016). Upstream ORFs are prevalent translational repressors in vertebrates. EMBO J 35, 706–723. 10.15252/embj.201592759.

46. Hinnebusch, A.G., Ivanov, I.P., and Sonenberg, N. (2016). Translational control by 5’-untranslated regions of eukaryotic mRNAs. Science 352, 1413–1416. 10.1126/science.aad9868.

47. Morris, D.R., and Geballe, A.P. (2000). Upstream open reading frames as regulators of mRNA translation. Mol Cell Biol 20, 8635–8642. 10.1128/MCB.20.23.8635-8642.2000.

48. Young, S.K., and Wek, R.C. (2016). Upstream Open Reading Frames Differentially Regulate Gene-specific Translation in the Integrated Stress Response. J Biol Chem 291, 16927–16935. 10.1074/jbc.R116.733899.

49. Dever, T.E., Feng, L., Wek, R.C., Cigan, A.M., Donahue, T.F., and Hinnebusch, A.G. (1992). Phosphorylation of initiation factor 2 alpha by protein kinase GCN2 mediates gene-specific translational control of GCN4 in yeast. Cell 68, 585–596. 10.1016/0092-8674(92)90193-g.

50. Palam, L.R., Baird, T.D., and Wek, R.C. (2011). Phosphorylation of eIF2 facilitates ribosomal bypass of an inhibitory upstream ORF to enhance CHOP translation. J Biol Chem 286, 10939–10949. 10.1074/jbc.M110.216093.

51. Baird, T.D., Palam, L.R., Fusakio, M.E., Willy, J.A., Davis, C.M., McClintick, J.N., Anthony, T.G., and Wek, R.C. (2014). Selective mRNA translation during eIF2 phosphorylation induces expression of IBTKalpha. Mol Biol Cell 25, 1686–1697. 10.1091/mbc.E14-02-0704.

52. Chen, Y.J., Tan, B.C., Cheng, Y.Y., Chen, J.S., and Lee, S.C. (2010). Differential regulation of CHOP translation by phosphorylated eIF4E under stress conditions. Nucleic Acids Res 38, 764–777. 10.1093/nar/gkp1034.

53. Fraser, H.B., Hirsh, A.E., Giaever, G., Kumm, J., and Eisen, M.B. (2004). Noise minimization in eukaryotic gene expression. PLoS Biol 2, e137. 10.1371/journal.pbio.0020137.

54. Thattai, M., and van Oudenaarden, A. (2001). Intrinsic noise in gene regulatory networks. Proc Natl Acad Sci U S A 98, 8614–8619. 10.1073/pnas.151588598.

55. Wu, H.W., Fajiculay, E., Wu, J.F., Yan, C.S., Hsu, C.P., and Wu, S.H. (2022). Noise reduction by upstream open reading frames. Nat Plants 8, 474–480. 10.1038/s41477-022-01136-8.

56. Bottorff, T.A., Park, H., Geballe, A.P., and Subramaniam, A.R. (2022). Translational buffering by ribosome stalling in upstream open reading frames. PLoS Genet 18, e1010460. 10.1371/journal.pgen.1010460.

57. Ciandrini, L., Stansfield, I., and Romano, M.C. (2010). Role of the particle’s stepping cycle in an asymmetric exclusion process: a model of mRNA translation. Phys Rev E Stat Nonlin Soft Matter Phys 81, 051904. 10.1103/PhysRevE.81.051904.

58. Reuveni, S., Meilijson, I., Kupiec, M., Ruppin, E., and Tuller, T. (2011). Genome-scale analysis of translation elongation with a ribosome flow model. PLoS Comput Biol 7, e1002127. 10.1371/journal.pcbi.1002127.

59. von der Haar, T. (2012). Mathematical and Computational Modelling of Ribosomal Movement and Protein Synthesis: an overview. Comput Struct Biotechnol J 1, e201204002. 10.5936/csbj.201204002.

60. Zhao, Y.B., and Krishnan, J. (2014). mRNA translation and protein synthesis: an analysis of different modelling methodologies and a new PBN based approach. BMC Syst Biol 8, 25. 10.1186/1752-0509-8-25.

61. Tamura, K., Subramanian, S., and Kumar, S. (2004). Temporal Patterns of Fruit Fly (*Drosophila*) Evolution Revealed by Mutation Clocks. Molecular Biology and Evolution 21, 36–44. 10.1093/molbev/msg236.

62. Ingolia, N.T., Ghaemmaghami, S., Newman, J.R., and Weissman, J.S. (2009). Genome-wide analysis in vivo of translation with nucleotide resolution using ribosome profiling. Science 324, 218–223. 10.1126/science.1168978.

63. Zhang, H., Dou, S., He, F., Luo, J., Wei, L., and Lu, J. (2018). Genome-wide maps of ribosomal occupancy provide insights into adaptive evolution and regulatory roles of uORFs during *Drosophila* development. PLoS Biol 16, e2003903. 10.1371/journal.pbio.2003903.

64. Chothani, S.P., Adami, E., Widjaja, A.A., Langley, S.R., Viswanathan, S., Pua, C.J., Zhihao, N.T., Harmston, N., D’Agostino, G., Whiffin, N., et al. (2022). A high-resolution map of human RNA translation. Mol Cell 82, 2885–2899 e2888. 10.1016/j.molcel.2022.06.023.

65. Ingolia, N.T. (2016). Ribosome Footprint Profiling of Translation throughout the Genome. Cell 165, 22–33. 10.1016/j.cell.2016.02.066.

66. Ingolia, N.T., Lareau, L.F., and Weissman, J.S. (2011). Ribosome profiling of mouse embryonic stem cells reveals the complexity and dynamics of mammalian proteomes. Cell 147, 789–802. 10.1016/j.cell.2011.10.002.

67. Zhang, H., Wang, Y., Wu, X., Tang, X., Wu, C., and Lu, J. (2021). Determinants of genome-wide distribution and evolution of uORFs in eukaryotes. Nat Commun 12, 1076. 10.1038/s41467-021-21394-y.

68. Kronja, I., Yuan, B., Eichhorn, S.W., Dzeyk, K., Krijgsveld, J., Bartel, D.P., and Orr-Weaver, T.L. (2014). Widespread changes in the posttranscriptional landscape at the *Drosophila* oocyte-to-embryo transition. Cell Rep 7, 1495–1508. 10.1016/j.celrep.2014.05.002.

69. Qin, X., Ahn, S., Speed, T.P., and Rubin, G.M. (2007). Global analyses of mRNA translational control during early *Drosophila* embryogenesis. Genome Biol 8, R63. 10.1186/gb-2007-8-4-r63.

70. Berleth, T., Burri, M., Thoma, G., Bopp, D., Richstein, S., Frigerio, G., Noll, M., and Nusslein-Volhard, C. (1988). The role of localization of bicoid RNA in organizing the anterior pattern of the *Drosophila* embryo. EMBO J 7, 1749–1756.

71. Driever, W., and Nusslein-Volhard, C. (1988). A gradient of bicoid protein in *Drosophila* embryos. Cell 54, 83–93. 10.1016/0092-8674(88)90182-1.

72. Driever, W., and Nusslein-Volhard, C. (1988). The bicoid protein determines position in the *Drosophila* embryo in a concentration-dependent manner. Cell 54, 95–104. 10.1016/0092-8674(88)90183-3.

73. Abdelmohsen, K., Panda, A.C., Kang, M.J., Guo, R., Kim, J., Grammatikakis, I., Yoon, J.H., Dudekula, D.B., Noh, J.H., Yang, X., et al. (2014). 7SL RNA represses p53 translation by competing with HuR. Nucleic Acids Res 42, 10099–10111. 10.1093/nar/gku686.

74. Panda, A.C., Abdelmohsen, K., Martindale, J.L., Di Germanio, C., Yang, X., Grammatikakis, I., Noh, J.H., Zhang, Y., Lehrmann, E., Dudekula, D.B., et al. (2016). Novel RNA-binding activity of MYF5 enhances Ccnd1/Cyclin D1 mRNA translation during myogenesis. Nucleic Acids Res 44, 2393–2408. 10.1093/nar/gkw023.

75. Panda, A.C., Martindale, J.L., and Gorospe, M. (2017). Polysome Fractionation to Analyze mRNA Distribution Profiles. Bio Protoc 7. 10.21769/BioProtoc.2126.

76. Amourda, C., Chong, J., and Saunders, T.E. (2018). MicroRNAs buffer genetic variation at specific temperatures during embryonic development. bioRxiv, 444810. 10.1101/444810.

77. Lu, G., Zhao, Y., Chen, Q., Lin, P., Tang, T., Tang, Z., Liufu, Z., and Wu, C.-I. (2021). When development is constantly but weakly perturbed - Canalization by microRNAs. bioRxiv, 2021.2009.2004.458966. 10.1101/2021.09.04.458966.

78. Li, X.Y., MacArthur, S., Bourgon, R., Nix, D., Pollard, D.A., Iyer, V.N., Hechmer, A., Simirenko, L., Stapleton, M., Luengo Hendriks, C.L., et al. (2008). Transcription factors bind thousands of active and inactive regions in the *Drosophila* blastoderm. PLoS Biol 6, e27. 10.1371/journal.pbio.0060027.

79. Wang, Z.Y., Leushkin, E., Liechti, A., Ovchinnikova, S., Mößinger, K., Brüning, T., Rummel, C., Grützner, F., Cardoso-Moreira, M., Janich, P., et al. (2020). Transcriptome and translatome co-evolution in mammals. Nature 588, 642–647. 10.1038/s41586-020-2899-z.

80. Hedges, S.B., Dudley, J., and Kumar, S. (2006). TimeTree: a public knowledge-base of divergence times among organisms. Bioinformatics 22, 2971–2972. 10.1093/bioinformatics/btl505.

81. Battle, A., Khan, Z., Wang, S.H., Mitrano, A., Ford, M.J., Pritchard, J.K., and Gilad, Y. (2015). Genomic variation. Impact of regulatory variation from RNA to protein. Science 347, 664–667. 10.1126/science.1260793.

82. Lappalainen, T., Sammeth, M., Friedlander, M.R., t Hoen, P.A., Monlong, J., Rivas, M.A., Gonzalez-Porta, M., Kurbatova, N., Griebel, T., Ferreira, P.G., et al. (2013). Transcriptome and genome sequencing uncovers functional variation in humans. Nature 501, 506–511. 10.1038/nature12531.

83. Stein, K.C., and Frydman, J. (2019). The stop-and-go traffic regulating protein biogenesis: How translation kinetics controls proteostasis. Journal of Biological Chemistry 294, 2076–2084.

84. Sherman, M.Y., and Qian, S.B. (2013). Less is more: improving proteostasis by translation slow down. Trends Biochem Sci 38, 585–591. 10.1016/j.tibs.2013.09.003.

85. Waddington, C.H. (1942). CANALIZATION OF DEVELOPMENT AND THE INHERITANCE OF ACQUIRED CHARACTERS. Nature 150, 563–565 10.1038/150563a0.

86. Payne, J.L., and Wagner, A. (2015). Mechanisms of mutational robustness in transcriptional regulation. Front Genet 6, 322. 10.3389/fgene.2015.00322.

87. Denby, C.M., Im, J.H., Yu, R.C., Pesce, C.G., and Brem, R.B. (2012). Negative feedback confers mutational robustness in yeast transcription factor regulation. Proc Natl Acad Sci U S A 109, 3874–3878. 10.1073/pnas.1116360109.

88. Jarosz, D.F., Taipale, M., and Lindquist, S. (2010). Protein homeostasis and the phenotypic manifestation of genetic diversity: principles and mechanisms. Annu Rev Genet 44, 189–216. 10.1146/annurev.genet.40.110405.090412.

89. Ebert, M.S., and Sharp, P.A. (2012). Roles for microRNAs in conferring robustness to biological processes. Cell 149, 515–524. 10.1016/j.cell.2012.04.005.

90. Lu, G.A., Zhang, J., Zhao, Y., Chen, Q., Lin, P., Tang, T., Tang, Z., Wen, H., Liufu, Z., and Wu, C.I. (2023). Canalization of Phenotypes-When the Transcriptome is Constantly but Weakly Perturbed. Mol Biol Evol 40 10.1093/molbev/msad005.

91. Alon, U. (2007). Network motifs: theory and experimental approaches. Nat Rev Genet 8, 450–461 10.1038/nrg2102.

92. Somogyvari, M., Khatatneh, S., and Soti, C. (2022). Hsp90: From Cellular to Organismal Proteostasis. Cells 11 10.3390/cells11162479.

93. Zabinsky, R.A., Mason, G.A., Queitsch, C., and Jarosz, D.F. (2019). It’s not magic - Hsp90 and its effects on genetic and epigenetic variation. Semin Cell Dev Biol 88, 21–35. 10.1016/j.semcdb.2018.05.015.

94. Lawless, C., Pearson, R.D., Selley, J.N., Smirnova, J.B., Grant, C.M., Ashe, M.P., Pavitt, G.D., and Hubbard, S.J. (2009). Upstream sequence elements direct post-transcriptional regulation of gene expression under stress conditions in yeast. BMC Genomics 10, 7 10.1186/1471-2164-10-7.

95. Mueller, P.P., and Hinnebusch, A.G. (1986). Multiple upstream AUG codons mediate translational control of GCN4. Cell 45, 201–207. 10.1016/0092-8674(86)90384-3.

96. Meijer, H.A., and Thomas, A.A. (2003). Ribosomes stalling on uORF1 in the Xenopus Cx41 5’ UTR inhibit downstream translation initiation. Nucleic Acids Res 31, 3174–3184. 10.1093/nar/gkg429.

97. Ivanov, I.P., Shin, B.S., Loughran, G., Tzani, I., Young-Baird, S.K., Cao, C., Atkins, J.F., and Dever, T.E. (2018). Polyamine Control of Translation Elongation Regulates Start Site Selection on Antizyme Inhibitor mRNA via Ribosome Queuing. Mol Cell 70, 254–264 e256. 10.1016/j.molcel.2018.03.015.

98. Lin, Y., May, G.E., Kready, H., Nazzaro, L., Mao, M., Spealman, P., Creeger, Y., and McManus, C.J. (2019). Impacts of uORF codon identity and position on translation regulation. Nucleic Acids Res 47, 9358–9367. 10.1093/nar/gkz681.

99. Wei, J., Wu, C., and Sachs, M.S. (2012). The arginine attenuator peptide interferes with the ribosome peptidyl transferase center. Mol Cell Biol 32, 2396–2406. 10.1128/MCB.00136-12.

100. Ito, K., and Chiba, S. (2013). Arrest peptides: cis-acting modulators of translation. Annu Rev Biochem 82, 171–202. 10.1146/annurev-biochem-080211-105026.

101. Vilela, C., and McCarthy, J.E. (2003). Regulation of fungal gene expression via short open reading frames in the mRNA 5’untranslated region. Mol Microbiol 49, 859–867. 10.1046/j.1365-2958.2003.03622.x.

102. Lovett, P.S., and Rogers, E.J. (1996). Ribosome regulation by the nascent peptide. Microbiol Rev 60, 366–385. 10.1128/mr.60.2.366-385.1996.

103. Zhao, T., Chen, Y.-M., Li, Y., Wang, J., Chen, S., Gao, N., and Qian, W. (2021). Disome-seq reveals widespread ribosome collisions that promote cotranslational protein folding. Genome Biology 22, 16. 10.1186/s13059-020-02256-0.

104. Meydan, S., and Guydosh, N.R. (2020). Disome and Trisome Profiling Reveal Genome-wide Targets of Ribosome Quality Control. Mol Cell 79, 588–602.e586.

105. Han, P., Shichino, Y., Schneider-Poetsch, T., Mito, M., Hashimoto, S., Udagawa, T., Kohno, K., Yoshida, M., Mishima, Y., Inada, T., and Iwasaki, S. (2020). Genome-wide Survey of Ribosome Collision. Cell Rep 31, 107610. 10.1016/j.celrep.2020.107610.

106. Juszkiewicz, S., Slodkowicz, G., Lin, Z., Freire-Pritchett, P., Peak-Chew, S.Y., and Hegde, R.S. (2020). Ribosome collisions trigger cis-acting feedback inhibition of translation initiation. Elife 9. 10.7554/eLife.60038.

107. Simms, C.L., Yan, L.L., Qiu, J.K., and Zaher, H.S. (2019). Ribosome Collisions Result in +1 Frameshifting in the Absence of No-Go Decay. Cell Reports 28, 1679–1689.e1674.

108. Goldman, D.H., Livingston, N.M., Movsik, J., Wu, B., and Green, R. (2021). Live-cell imaging reveals kinetic determinants of quality control triggered by ribosome stalling. Mol Cell 81, 1830–1840.e1838.

109. Sundaramoorthy, E., Leonard, M., Mak, R., Liao, J., Fulzele, A., and Bennett, E.J. (2017). ZNF598 and RACK1 Regulate Mammalian Ribosome-Associated Quality Control Function by Mediating Regulatory 40S Ribosomal Ubiquitylation. Mol Cell 65, 751–760.e754. 10.1016/j.molcel.2016.12.026.

110. Juszkiewicz, S., Chandrasekaran, V., Lin, Z., Kraatz, S., Ramakrishnan, V., and Hegde, R.S. (2018). ZNF598 Is a Quality Control Sensor of Collided Ribosomes. Mol Cell 72, 469–481.e467. 10.1016/j.molcel.2018.08.037.

111. Ugajin, N., Imami, K., Takada, H., Ishihama, Y., Chiba, S., and Mishima, Y. (2023). Znf598-mediated Rps10/eS10 ubiquitination contributes to the ribosome ubiquitination dynamics during zebrafish development. Rna 29, 1910–1927. 10.1261/rna.079633.123.

112. Geng, J., Li, S., Li, Y., Wu, Z., Bhurtel, S., Rimal, S., Khan, D., Ohja, R., Brandman, O., and Lu, B. (2024). Stalled translation by mitochondrial stress upregulates a CNOT4-ZNF598 ribosomal quality control pathway important for tissue homeostasis. Nat Commun 15, 1637. 10.1038/s41467-024-45525-3.

113. Narita, M., Denk, T., Matsuo, Y., Sugiyama, T., Kikuguchi, C., Ito, S., Sato, N., Suzuki, T., Hashimoto, S., Machova, I., et al. (2022). A distinct mammalian disome collision interface harbors K63-linked polyubiquitination of uS10 to trigger hRQT-mediated subunit dissociation. Nat Commun 13, 6411. 10.1038/s41467-022-34097-9.

114. Best, K., Ikeuchi, K., Kater, L., Best, D., Musial, J., Matsuo, Y., Berninghausen, O., Becker, T., Inada, T., and Beckmann, R. (2023). Structural basis for clearing of ribosome collisions by the RQT complex. Nat Commun 14, 921. 10.1038/s41467-023-36230-8.

115. Hashimoto, S., Sugiyama, T., Yamazaki, R., Nobuta, R., and Inada, T. (2020). Identification of a novel trigger complex that facilitates ribosome-associated quality control in mammalian cells. Sci Rep 10, 3422. 10.1038/s41598-020-60241-w.

116. Madern, M.F., Yang, S., Witteveen, O., Segeren, H.A., Bauer, M., and Tanenbaum, M.E. (2025). Long-term imaging of individual ribosomes reveals ribosome cooperativity in mRNA translation. Cell. 10.1016/j.cell.2025.01.016.

117. Garshott, D.M., An, H., Sundaramoorthy, E., Leonard, M., Vicary, A., Harper, J.W., and Bennett, E.J. (2021). iRQC, a surveillance pathway for 40S ribosomal quality control during mRNA translation initiation. Cell Rep 36, 109642. 10.1016/j.celrep.2021.109642.

118. Ford, P.W., Garshott, D.M., Narasimhan, M., Ge, X., Jordahl, E.M., Subramanya, S., and Bennett, E.J. (2025). RNF10 and RIOK3 facilitate 40S ribosomal subunit degradation upon 60S biogenesis disruption or amino acid starvation. Cell Rep 44, 115371. 10.1016/j.celrep.2025.115371.

119. Cho, P.F., Poulin, F., Cho-Park, Y.A., Cho-Park, I.B., Chicoine, J.D., Lasko, P., and Sonenberg, N. (2005). A new paradigm for translational control: inhibition via 5’-3’ mRNA tethering by Bicoid and the eIF4E cognate 4EHP. Cell 121, 411–423. 10.1016/j.cell.2005.02.024.

120. Singh, A.P., Wu, P., Ryabichko, S., Raimundo, J., Swan, M., Wieschaus, E., Gregor, T., and Toettcher, J.E. (2022). Optogenetic control of the Bicoid morphogen reveals fast and slow modes of gap gene regulation. Cell Rep 38, 110543. 10.1016/j.celrep.2022.110543.

121. Hannon, C.E., Blythe, S.A., and Wieschaus, E.F. (2017). Concentration dependent chromatin states induced by the bicoid morphogen gradient. Elife 6. 10.7554/eLife.28275.

122. Struhl, G., Struhl, K., and Macdonald, P.M. (1989). The gradient morphogen bicoid is a concentration-dependent transcriptional activator. Cell 57, 1259–1273. 10.1016/0092-8674(89)90062-7.

123. Dubnau, J., and Struhl, G. (1996). RNA recognition and translational regulation by a homeodomain protein. Nature 379, 694–699. 10.1038/379694a0.

124. Wethmar, K., Begay, V., Smink, J.J., Zaragoza, K., Wiesenthal, V., Dorken, B., Calkhoven, C.F., and Leutz, A. (2010). C/EBPbetaDeltauORF mice--a genetic model for uORF-mediated translational control in mammals. Genes Dev 24, 15–20. 10.1101/gad.557910.

125. Miyake, T., Inoue, Y., Shao, X., Seta, T., Aoki, Y., Nguyen Pham, K.T., Shichino, Y., Sasaki, J., Sasaki, T., Ikawa, M., et al. (2023). Minimal upstream open reading frame of Per2 mediates phase fitness of the circadian clock to day/night physiological body temperature rhythm. Cell Rep 42, 112157. 10.1016/j.celrep.2023.112157.

126. Xing, S., Chen, K., Zhu, H., Zhang, R., Zhang, H., Li, B., and Gao, C. (2020). Fine-tuning sugar content in strawberry. Genome Biol 21, 230. 10.1186/s13059-020-02146-5.

127. Zhang, H., Si, X., Ji, X., Fan, R., Liu, J., Chen, K., Wang, D., and Gao, C. (2018). Genome editing of upstream open reading frames enables translational control in plants. Nat Biotechnol 36, 894–898. 10.1038/nbt.4202.

128. Si, X., Zhang, H., Wang, Y., Chen, K., and Gao, C. (2020). Manipulating gene translation in plants by CRISPR-Cas9-mediated genome editing of upstream open reading frames. Nat Protoc 15, 338–363. 10.1038/s41596-019-0238-3.

129. Xue, C., Qiu, F., Wang, Y., Li, B., Zhao, K.T., Chen, K., and Gao, C. (2023). Tuning plant phenotypes by precise, graded downregulation of gene expression. Nat Biotechnol. 10.1038/s41587-023-01707-w.

130. Luo, Z., and Sachs, M.S. (1996). Role of an upstream open reading frame in mediating arginine-specific translational control in Neurospora crassa. J Bacteriol 178, 2172–2177. 10.1128/jb.178.8.2172-2177.1996.

131. Rosenbloom, K.R., Armstrong, J., Barber, G.P., Casper, J., Clawson, H., Diekhans, M., Dreszer, T.R., Fujita, P.A., Guruvadoo, L., Haeussler, M., et al. (2015). The UCSC Genome Browser database: 2015 update. Nucleic Acids Res 43, D670–681. 10.1093/nar/gku1177.

132. Chakraborty, M., Chang, C.H., Khost, D.E., Vedanayagam, J., Adrion, J.R., Liao, Y., Montooth, K.L., Meiklejohn, C.D., Larracuente, A.M., and Emerson, J.J. (2021). Evolution of genome structure in the *Drosophila simulans* species complex. Genome Res 31, 380–396. 10.1101/gr.263442.120.

133. Chiaromonte, F., Yap, V.B., and Miller, W. (2002). Scoring pairwise genomic sequence alignments. Pac Symp Biocomput, 115-126. 10.1142/9789812799623_0012.

134. Blanchette, M., Kent, W.J., Riemer, C., Elnitski, L., Smit, A.F., Roskin, K.M., Baertsch, R., Rosenbloom, K., Clawson, H., Green, E.D., et al. (2004). Aligning multiple genomic sequences with the threaded blockset aligner. Genome Res 14, 708–715. 10.1101/gr.1933104.

135. Martin, M. (2011). Cutadapt removes adapter sequences from high-throughput sequencing reads. EMBnet. journal 17, 10–12.

136. Langmead, B., and Salzberg, S.L. (2012). Fast gapped-read alignment with Bowtie 2. Nat Methods 9, 357–359. 10.1038/nmeth.1923.

137. Dobin, A., Davis, C.A., Schlesinger, F., Drenkow, J., Zaleski, C., Jha, S., Batut, P., Chaisson, M., and Gingeras, T.R. (2013). STAR: ultrafast universal RNA-seq aligner. Bioinformatics 29, 15–21.

138. Dunn, J.G., and Weissman, J.S. (2016). Plastid: nucleotide-resolution analysis of next-generation sequencing and genomics data. BMC Genomics 17, 958. 10.1186/s12864-016-3278-x.

139. Ni, J.Q., Zhou, R., Czech, B., Liu, L.P., Holderbaum, L., Yang-Zhou, D., Shim, H.S., Tao, R., Handler, D., Karpowicz, P., et al. (2011). A genome-scale shRNA resource for transgenic RNAi in *Drosophila*. Nat Methods 8, 405–407. 10.1038/nmeth.1592.

140. Anders, S., Pyl, P.T., and Huber, W. (2015). HTSeq--a Python framework to work with high-throughput sequencing data. Bioinformatics 31, 166–169. 10.1093/bioinformatics/btu638.

141. McCarthy, D.J., Chen, Y., and Smyth, G.K. (2012). Differential expression analysis of multifactor RNA-Seq experiments with respect to biological variation. Nucleic Acids Res 40, 4288–4297. 10.1093/nar/gks042.

142. Yu, G., Wang, L.G., Han, Y., and He, Q.Y. (2012). clusterProfiler: an R package for comparing biological themes among gene clusters. OMICS 16, 284–287. 10.1089/omi.2011.0118.

143. Xiao, Z., Zou, Q., Liu, Y., and Yang, X. (2016). Genome-wide assessment of differential translations with ribosome profiling data. Nat Commun 7, 11194. 10.1038/ncomms11194.

## SOM Reference

1. Andreev, D.E., Arnold, M., Kiniry, S.J., Loughran, G., Michel, A.M., Rachinskii, D., and Baranov, P.V. (2018). TASEP modelling provides a parsimonious explanation for the ability of a single uORF to derepress translation during the integrated stress response. Elife 7. 10.7554/eLife.32563.

2. Ivanov, I.P., Shin, B.S., Loughran, G., Tzani, I., Young-Baird, S.K., Cao, C., Atkins, J.F., and Dever, T.E. (2018). Polyamine Control of Translation Elongation Regulates Start Site Selection on Antizyme Inhibitor mRNA via Ribosome Queuing. Mol Cell 70, 254–264 e256. 10.1016/j.molcel.2018.03.015.

3. Luo, Z., and Sachs, M.S. (1996). Role of an upstream open reading frame in mediating arginine-specific translational control in Neurospora crassa. J Bacteriol 178, 2172–2177. 10.1128/jb.178.8.2172-2177.1996.

4. Lovett, P.S., and Rogers, E.J. (1996). Ribosome regulation by the nascent peptide. Microbiol Rev 60, 366–385. 10.1128/mr.60.2.366-385.1996.

5. Vilela, C., and McCarthy, J.E. (2003). Regulation of fungal gene expression via short open reading frames in the mRNA 5’untranslated region. Mol Microbiol 49, 859–867. 10.1046/j.1365-2958.2003.03622.x.

6. Raney, A., Law, G.L., Mize, G.J., and Morris, D.R. (2002). Regulated translation termination at the upstream open reading frame in s-adenosylmethionine decarboxylase mRNA. J Biol Chem 277, 5988–5994. 10.1074/jbc.M108375200.

7. Lin, Y., May, G.E., Kready, H., Nazzaro, L., Mao, M., Spealman, P., Creeger, Y., and McManus, C.J. (2019). Impacts of uORF codon identity and position on translation regulation. Nucleic Acids Res 47, 9358–9367. 10.1093/nar/gkz681.

8. Meijer, H.A., and Thomas, A.A. (2003). Ribosomes stalling on uORF1 in the Xenopus Cx41 5’ UTR inhibit downstream translation initiation. Nucleic Acids Res 31, 3174–3184. 10.1093/nar/gkg429.

9. Bottorff, T.A., Park, H., Geballe, A.P., and Subramaniam, A.R. (2022). Translational buffering by ribosome stalling in upstream open reading frames. PLoS Genet 18, e1010460. 10.1371/journal.pgen.1010460.

10. Zhang, H., Dou, S., He, F., Luo, J., Wei, L., and Lu, J. (2018). Genome-wide maps of ribosomal occupancy provide insights into adaptive evolution and regulatory roles of uORFs during Drosophila development. PLoS biology 16, e2003903.

