## Supplemental figures and tables for "Upstream open reading frames buffer translational variability during *Drosophila* evolution and development"

These above operations are executed repeatedly until the action count reaches the limit given by the user (1,000,000 in this study).

### Supplementary Figures and Tables

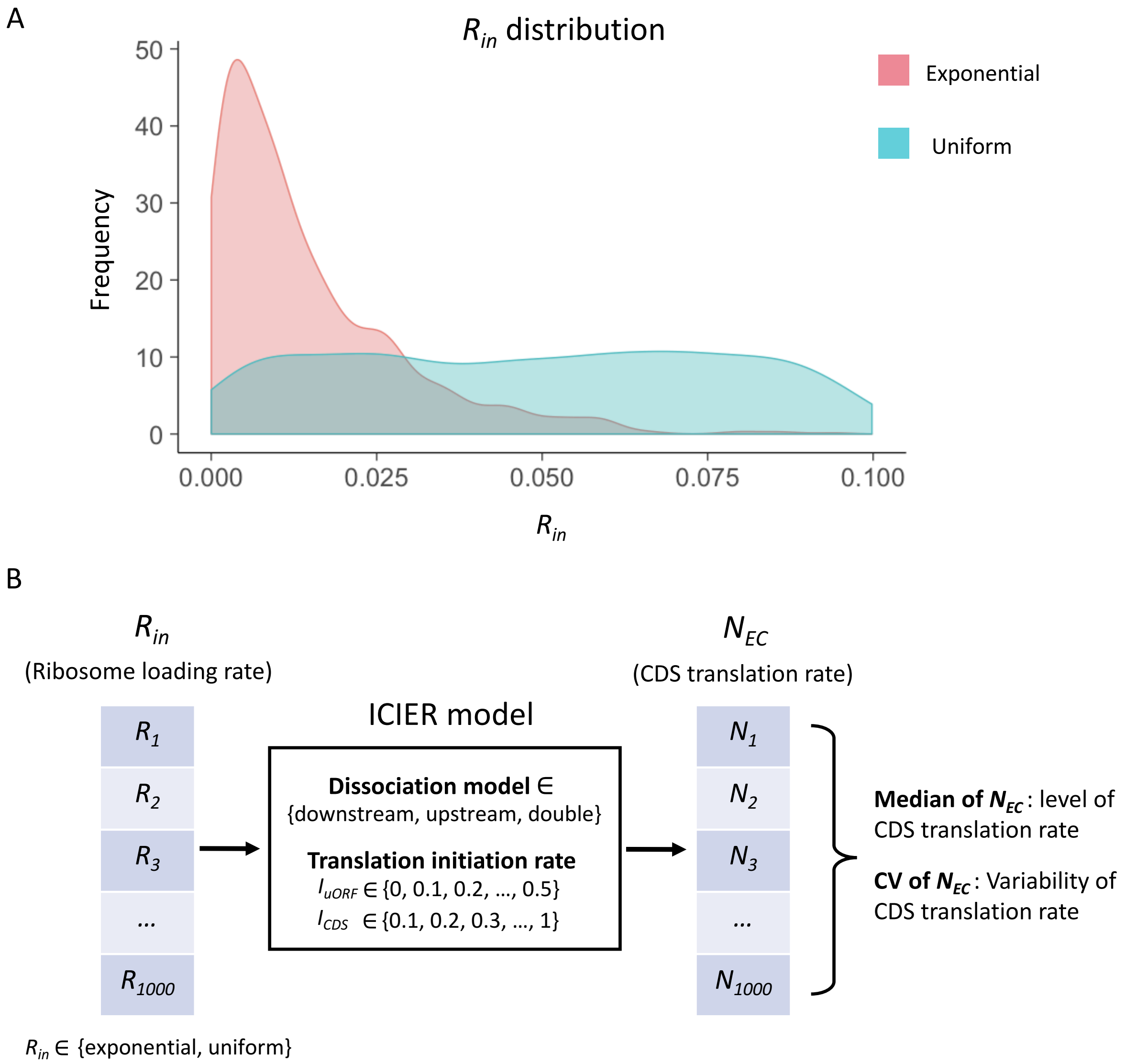


**Figure S1. The input distribution and model simulation flow.** (A) Two distributions of the $R_{in}$ input were used in our simulation: an exponential distribution and a uniform distribution. A total of 1,000 $R_{in}$ values (ranging from 0 to 0.1) were randomly generated following either distribution. (B) Under a fixed combination of parameters ($R_{in}$ distribution, dissociation model, $I_{uORF}$ and $I_{CDS}$), the model simulation produces 1000 $N_{EC}$ values from 1000 $R_{in}$ input. The median of these 1000 $N_{EC}$ represented the median of translation rate of CDS under the 1000 varying $R_{in}$ inputs and the other fixed parameters. The the coefficient of variation (CV) of these 1000 $N_{EC}$ reflected the variability of translation rate of CDS under the 1000 varying $R_{in}$ inputs and the other fixed parameters.

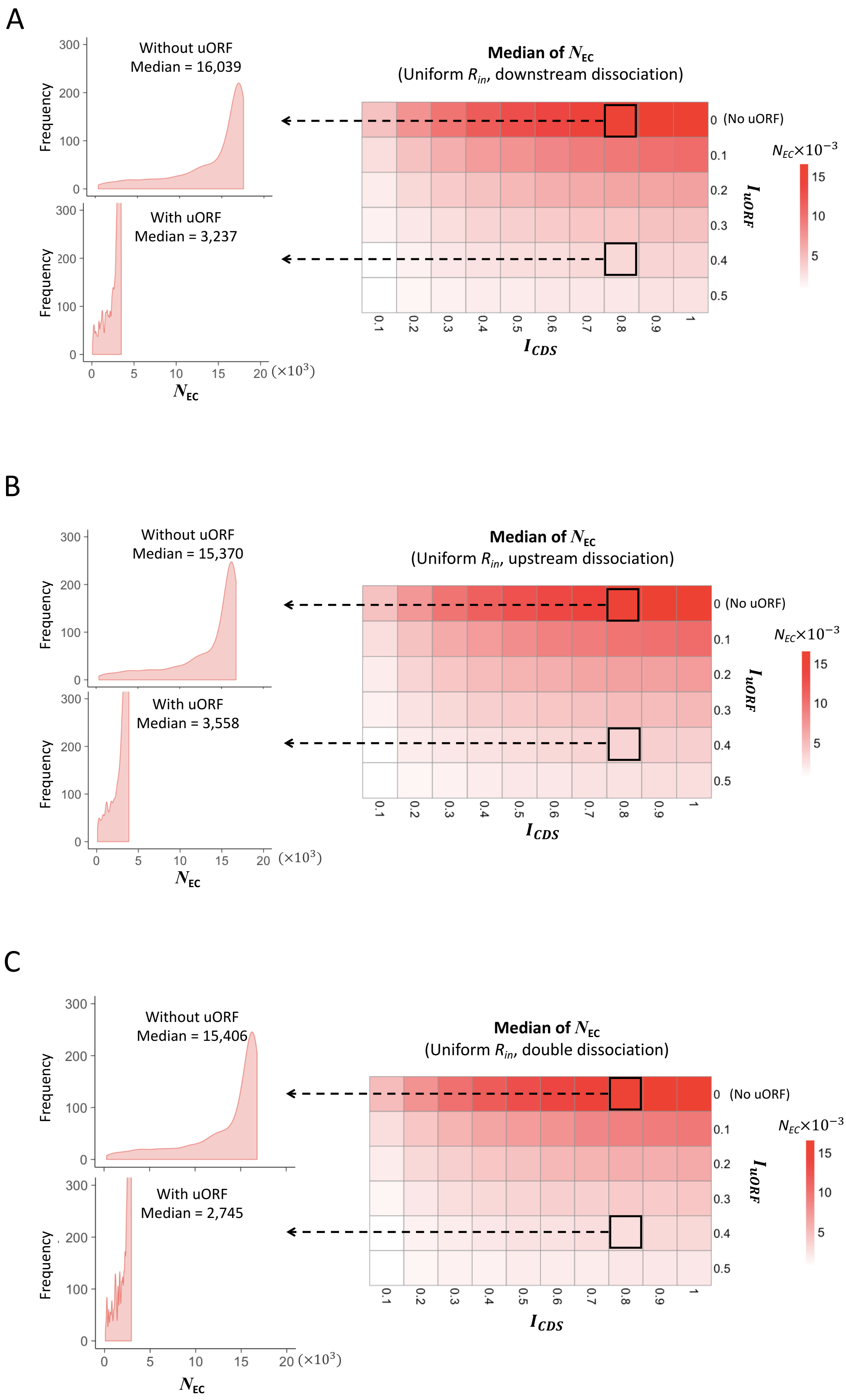

A
Without uORF
Median = 16,039
With uORF
Median = 3,237
Frequency
$N_{EC}$
(×10³)
Median of $N_{EC}$
(Uniform $R_{in}$, downstream dissociation)
0 (No uORF)
$I_{uORF}$
$I_{CDS}$
$N_{EC}\times10^{-3}$
B
Without uORF
Median = 15,370
With uORF
Median = 3,558
Median of $N_{EC}$
(Uniform $R_{in}$, upstream dissociation)
C
Without uORF
Median = 15,406
With uORF
Median = 2,745
Median of $N_{EC}$
(Uniform $R_{in}$, double dissociation)

**Figure S2.** Heatmaps showing the median CDS translation rate ($N_{EC}$) under different $I_{CDS}$ (x-axis) and $I_{uORF}$ (y-axis) combinations with a uniform distribution of $R_{in}$ input and the downstream dissociation model (A), a uniform distribution of $R_{in}$ input and the upstream dissociation model (B), a uniform distribution of $R_{in}$ input and the double dissociation model (C). For each heatmap, the left panels elicited by the dotted lines from specific squares of right heatmap were two examples showing the distribution of $N_{EC}$ under $I_{CDS}$ = 0.8 & $I_{uORF}$ = 0 (top panel, without uORF) and $I_{CDS}$ = 0.8 & $I_{uORF}$ = 0.4 (bottom panel, with uORF).

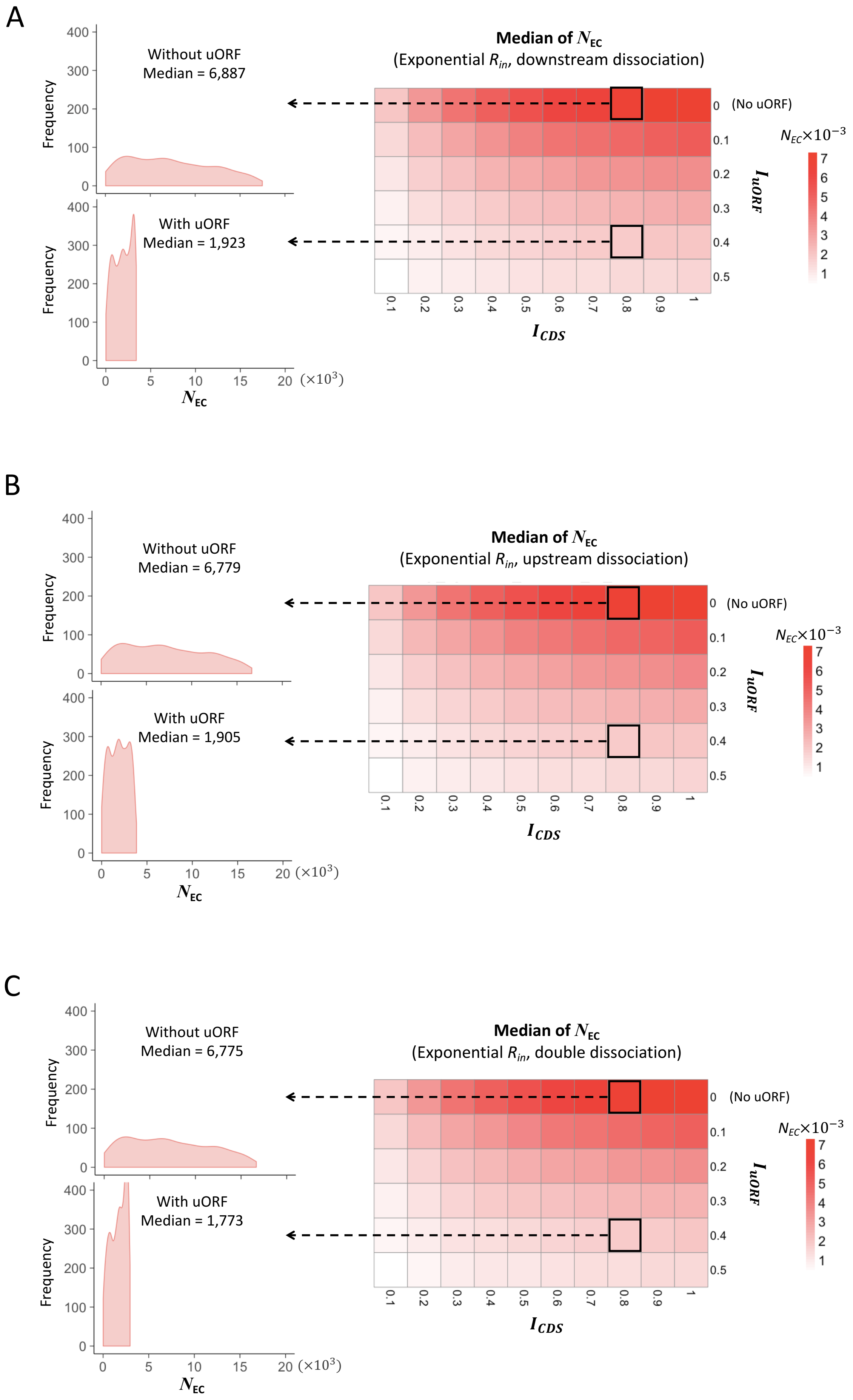

A
Without uORF
Median = 6,887
With uORF
Median = 1,923
Frequency
Median of $N_{EC}$
(Exponential $R_{in}$, downstream dissociation)
(No uORF)
$N_{EC}\times10^{-3}$
$I_{uORF}$
$I_{CDS}$
$N_{EC}$
($\times10^3$)
B
Without uORF
Median = 6,779
With uORF
Median = 1,905
Frequency
Median of $N_{EC}$
(Exponential $R_{in}$, upstream dissociation)
(No uORF)
$N_{EC}\times10^{-3}$
$I_{uORF}$
$I_{CDS}$
$N_{EC}$
($\times10^3$)
C
Without uORF
Median = 6,775
With uORF
Median = 1,773
Frequency
Median of $N_{EC}$
(Exponential $R_{in}$, double dissociation)
(No uORF)
$N_{EC}\times10^{-3}$
$I_{uORF}$
$I_{CDS}$
$N_{EC}$
($\times10^3$)

**Figure S3.** Heatmaps showing the median CDS translation rate ($N_{EC}$) under different $I_{CDS}$ (x-axis) and $I_{uORF}$ (y-axis) combinations with an exponential distribution of $R_{in}$ input and the downstream dissociation model (A), an exponential distribution of $R_{in}$ input and the upstream dissociation model (B), an exponential distribution of $R_{in}$ input and the double dissociation model (C). For each heatmap, the left panels elicited by the dotted lines from specific squares of right heatmap were two examples showing the distribution of $N_{EC}$ under $I_{CDS}$ = 0.8 & $I_{uORF}$ = 0 (top panel, without uORF) and $I_{CDS}$ = 0.8 & $I_{uORF}$ = 0.4 (bottom panel, with uORF).

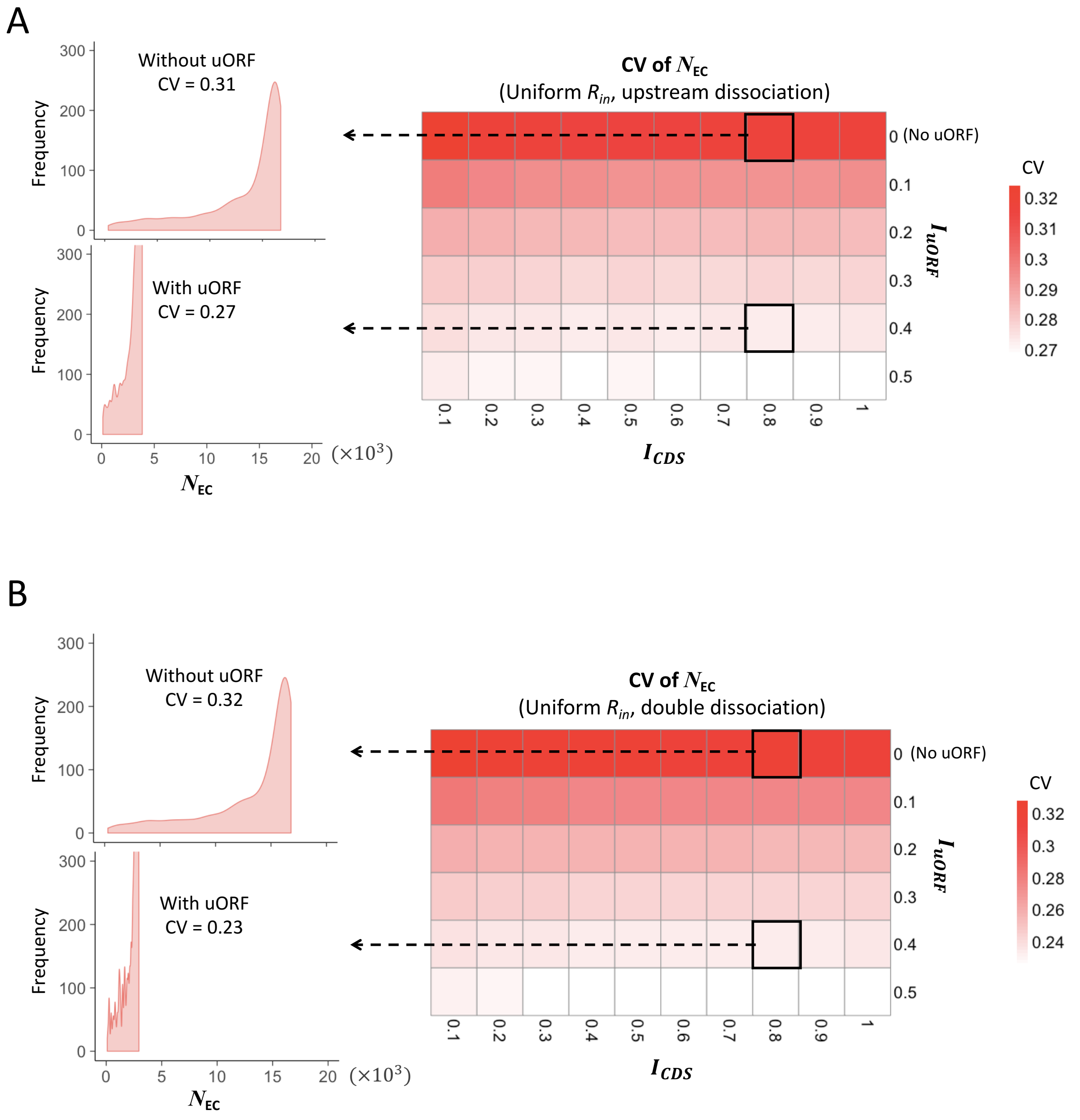


**Figure S4.** Heatmaps showing the CVs of CDS translation rate ($N_{EC}$) under different $I_{CDS}$ (x-axis) and $I_{uORF}$ (y-axis) combinations with a uniform distribution of $R_{in}$ input and the upstream dissociation model (A), a uniform distribution of $R_{in}$ input and the double dissociation model (B). For each heatmap, the left panels elicited by the dotted lines from specific squares of right heatmap were two examples showing the distribution of $N_{EC}$ under $I_{CDS}$ = 0.8 & $I_{uORF}$ = 0 (top panel, without uORF) and $I_{CDS}$ = 0.8 & $I_{uORF}$ = 0.4 (bottom panel, with uORF).

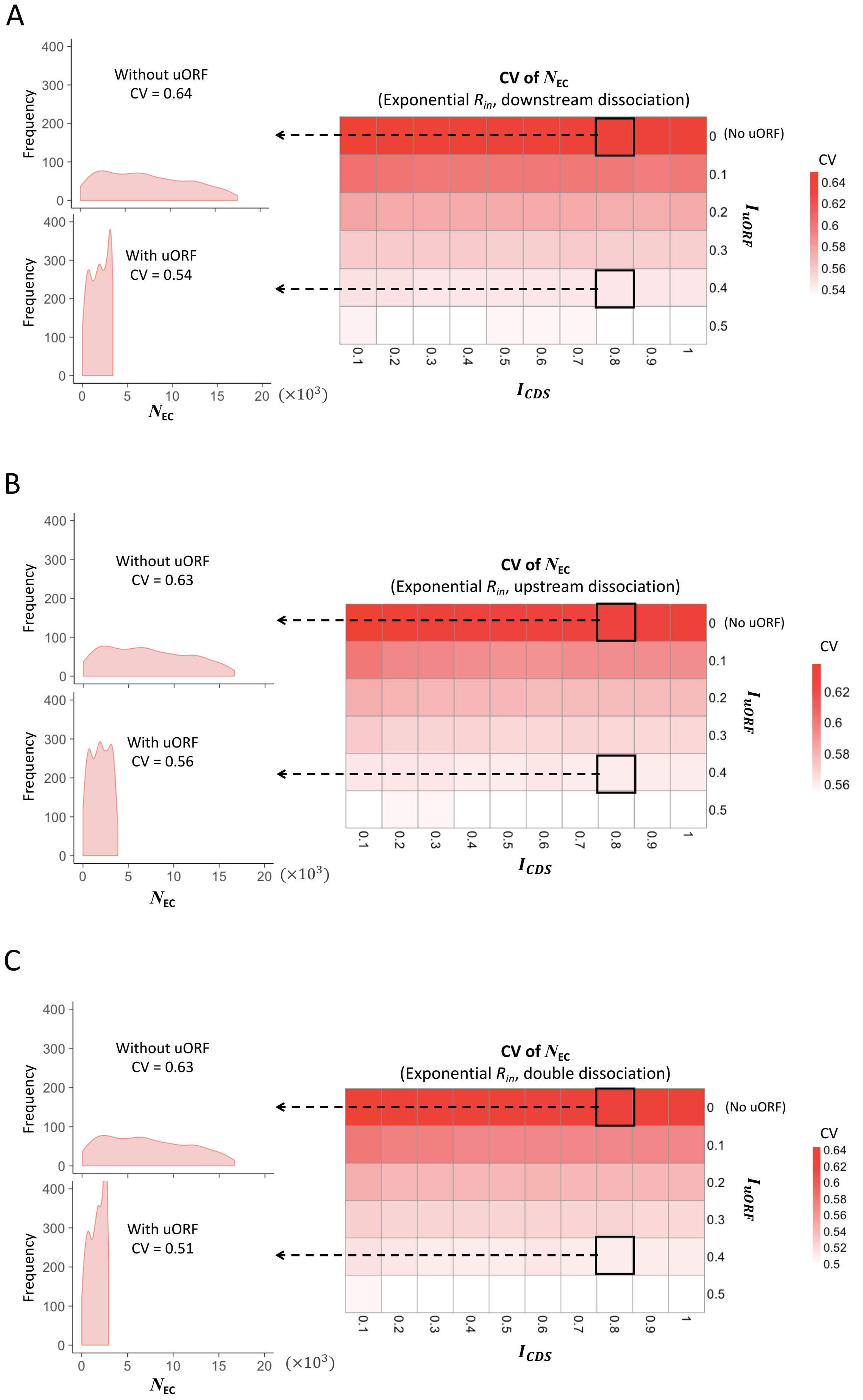
A
Without uORF
CV = 0.64
With uORF
CV = 0.54
Frequency
$N_{EC}$
(×10³)
CV of $N_{EC}$
(Exponential $R_{in}$, downstream dissociation)
0 (No uORF)
$I_{uORF}$
$I_{CDS}$
CV
B
Without uORF
CV = 0.63
With uORF
CV = 0.56
CV of $N_{EC}$
(Exponential $R_{in}$, upstream dissociation)
C
Without uORF
CV = 0.63
With uORF
CV = 0.51
CV of $N_{EC}$
(Exponential $R_{in}$, double dissociation)

72

**Figure S5.** Heatmaps showing the CVs of CDS translation rate ($N_{EC}$) under different $I_{CDS}$ (x-axis) and $I_{uORF}$ (y-axis) combinations with an exponential distribution of $R_{in}$ input and the downstream dissociation model (A), an exponential distribution of $R_{in}$ input and the upstream dissociation model (B), an exponential distribution of $R_{in}$ input and the double dissociation model (C). For each heatmap, the left panels elicited by the dotted lines from specific squares of right heatmap were two examples showing the distribution of $N_{EC}$ under $I_{CDS}$ = 0.8 & $I_{uORF}$ = 0 (top panel, without uORF) and $I_{CDS}$ = 0.8 & $I_{uORF}$ = 0.4 (bottom panel, with uORF).

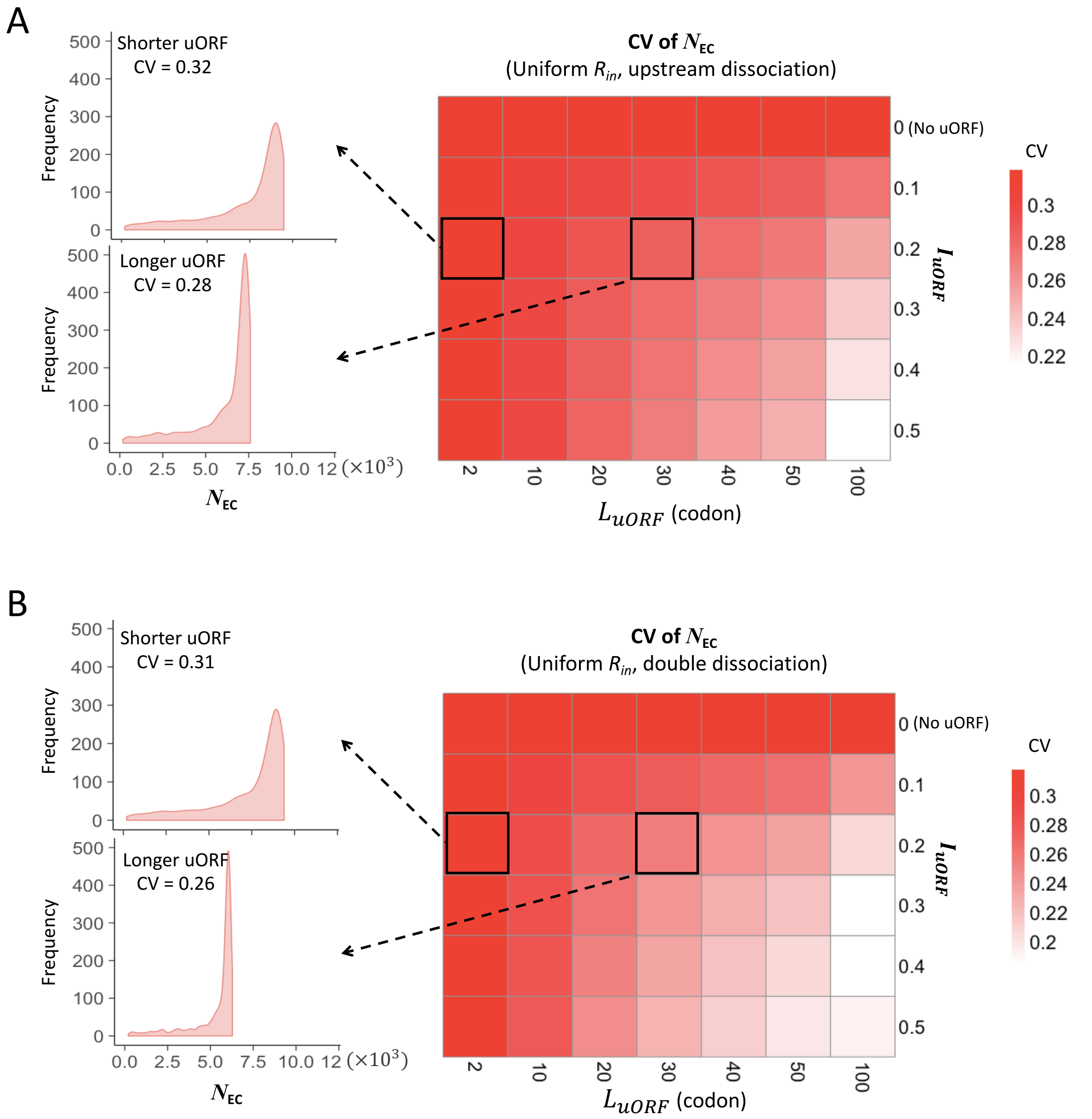


**Figure S6.** Heatmaps showing the CVs of CDS translation rate ($N_{EC}$) under different $L_{uORF}$ (x-axis) and $I_{uORF}$ (y-axis) combinations with a uniform distribution of $R_{in}$ input and the upstream dissociation model (A), a uniform distribution of $R_{in}$ input and the double dissociation model (B). For each heatmap, the left panels elicited by the dotted lines from specific squares of right heatmap were two examples showing the distribution of $N_{EC}$ under $L_{uORF} = 2$ & $I_{uORF} = 0.2$ (top panel, shorter uORF) and $L_{uORF} = 30$ & $I_{uORF} = 0.2$ (bottom panel, longer uORF).

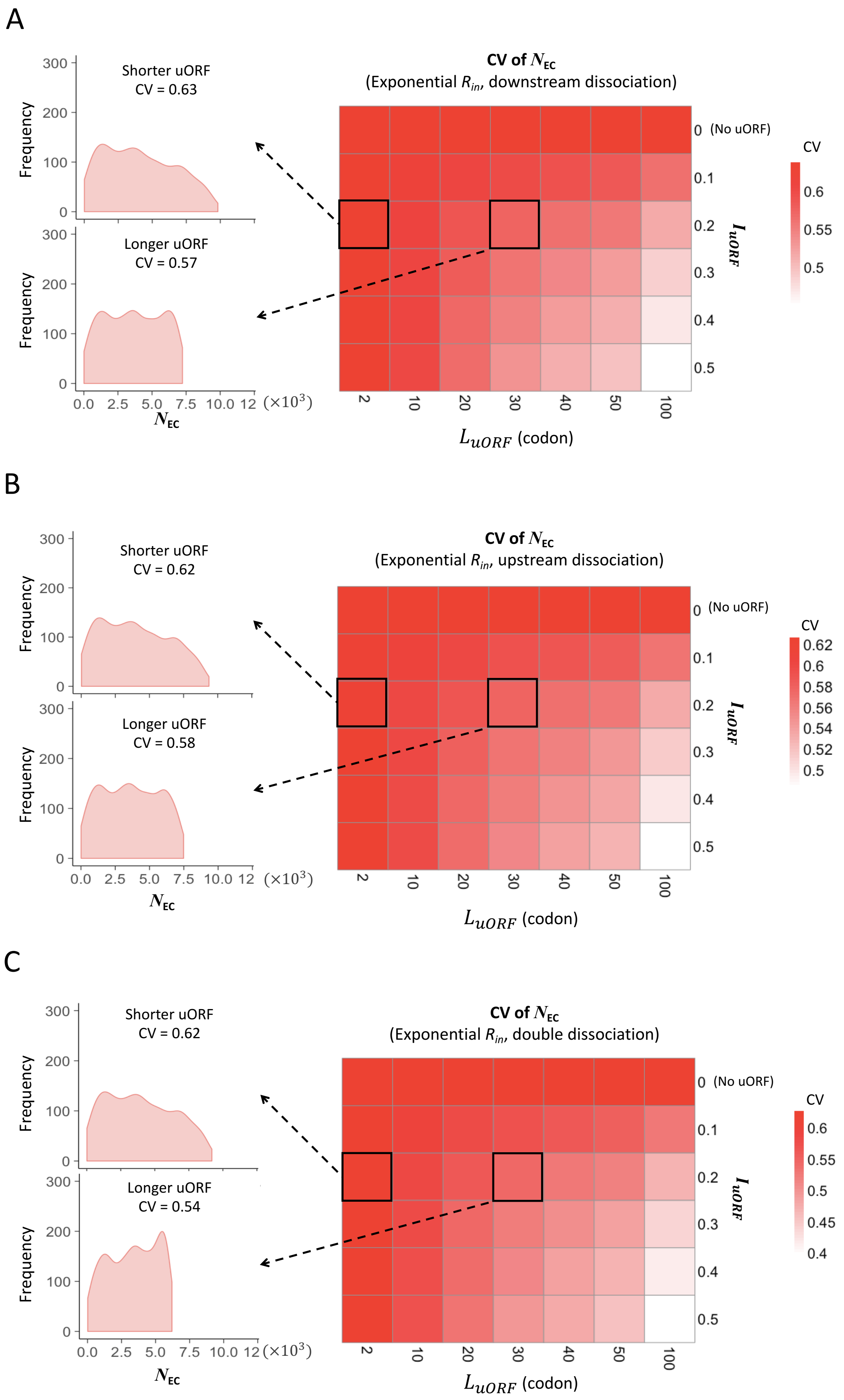
A
Shorter uORF
CV = 0.63
Longer uORF
CV = 0.57
Frequency
0.0 2.5 5.0 7.5 10.0 12 (×10³)
N_EC
CV of N_EC
(Exponential R_in, downstream dissociation)
0 (No uORF)
0.1
0.2
0.3
0.4
0.5
I_uORF
L_uORF (codon)
2 10 20 30 40 50 100
CV
0.6
0.55
0.5
B
Shorter uORF
CV = 0.62
Longer uORF
CV = 0.58
CV of N_EC
(Exponential R_in, upstream dissociation)
0.62
0.6
0.58
0.56
0.54
0.52
0.5
C
Shorter uORF
CV = 0.62
Longer uORF
CV = 0.54
CV of N_EC
(Exponential R_in, double dissociation)
0.6
0.55
0.5
0.45
0.4

89

**Figure S7.** Heatmaps showing the CVs of CDS translation rate ($N_{EC}$) under different $L_{uORF}$ (x-axis) and $I_{uORF}$ (y-axis) combinations with an exponential distribution of $R_{in}$ input and the downstream dissociation model (A), an exponential distribution of $R_{in}$ input and the upstream dissociation model (B), an exponential distribution of $R_{in}$ input and the double dissociation model (C). For each heatmap, the left panels elicited by the dotted lines from specific squares of right heatmap were two examples showing the distribution of $N_{EC}$ under $L_{uORF}$ = 2 & $I_{uORF}$ = 0.2 (top panel, shorter uORF) and $L_{uORF}$ = 30 & $I_{uORF}$ = 0.2 (bottom panel, longer uORF).

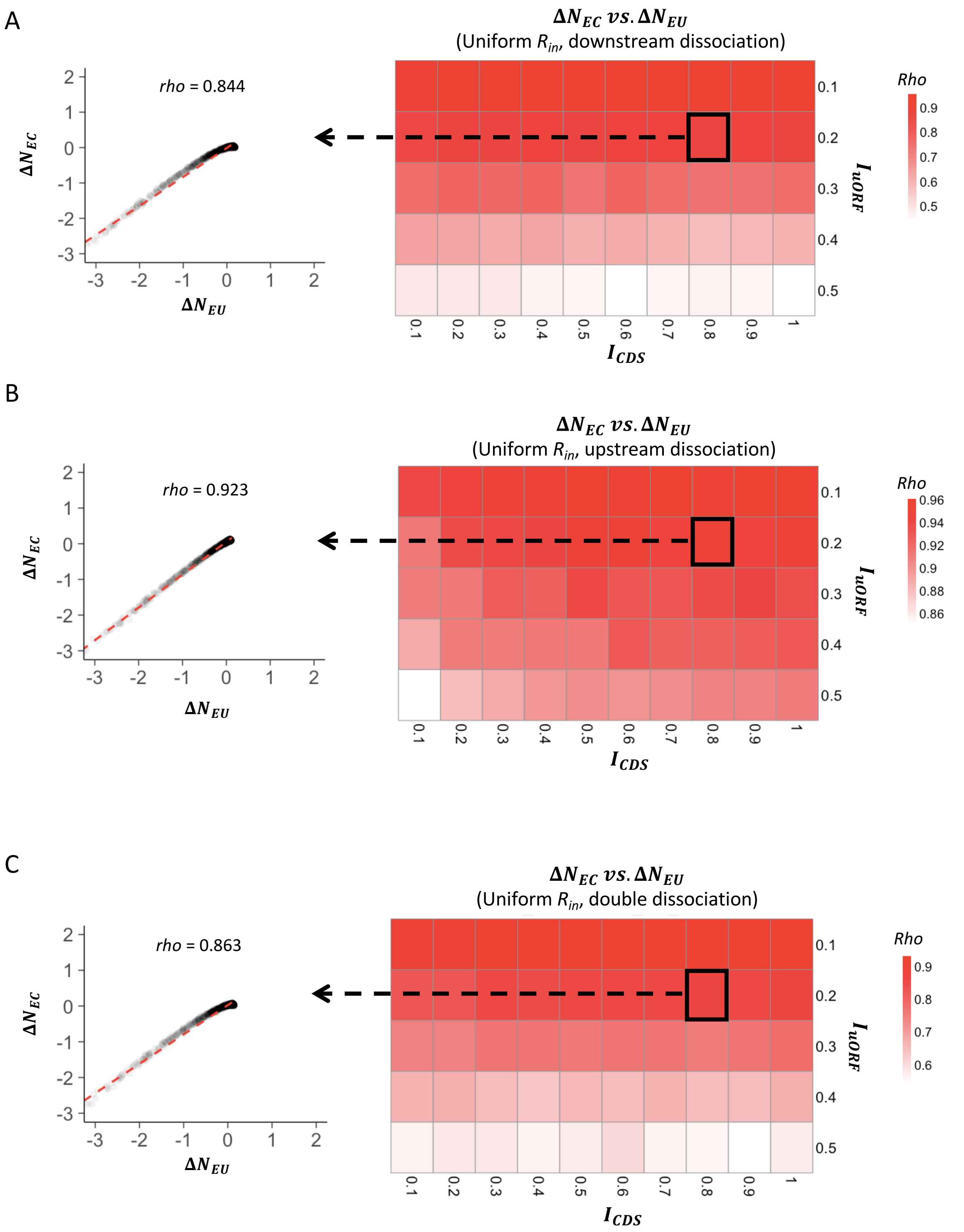


**Figure S8.** Heatmaps showing the Spearman's correlations between changes of uORF translation rate ($\Delta N_{EU}$) and downstream CDS translation rate ($\Delta N_{EC}$) under different $I_{CDS}$ (x-axis) and $I_{uORF}$ (y-axis) combinations with a uniform distribution of $R_{in}$ input and the downstream dissociation model (A), a uniform distribution of $R_{in}$ input and the upstream dissociation model (B), a uniform distribution of $R_{in}$ input and the double dissociation model (C). For each heatmap, the left panel elicited by the dotted line from the specific square of right heatmap was an example showing the Spearman's correlations between changes of uORF translation rate ($\Delta N_{EU}$) and downstream CDS translation rate ($\Delta N_{EC}$) under $I_{CDS}$ = 0.8 & $I_{uORF}$ = 0.2. All $P$ values < 0.001.

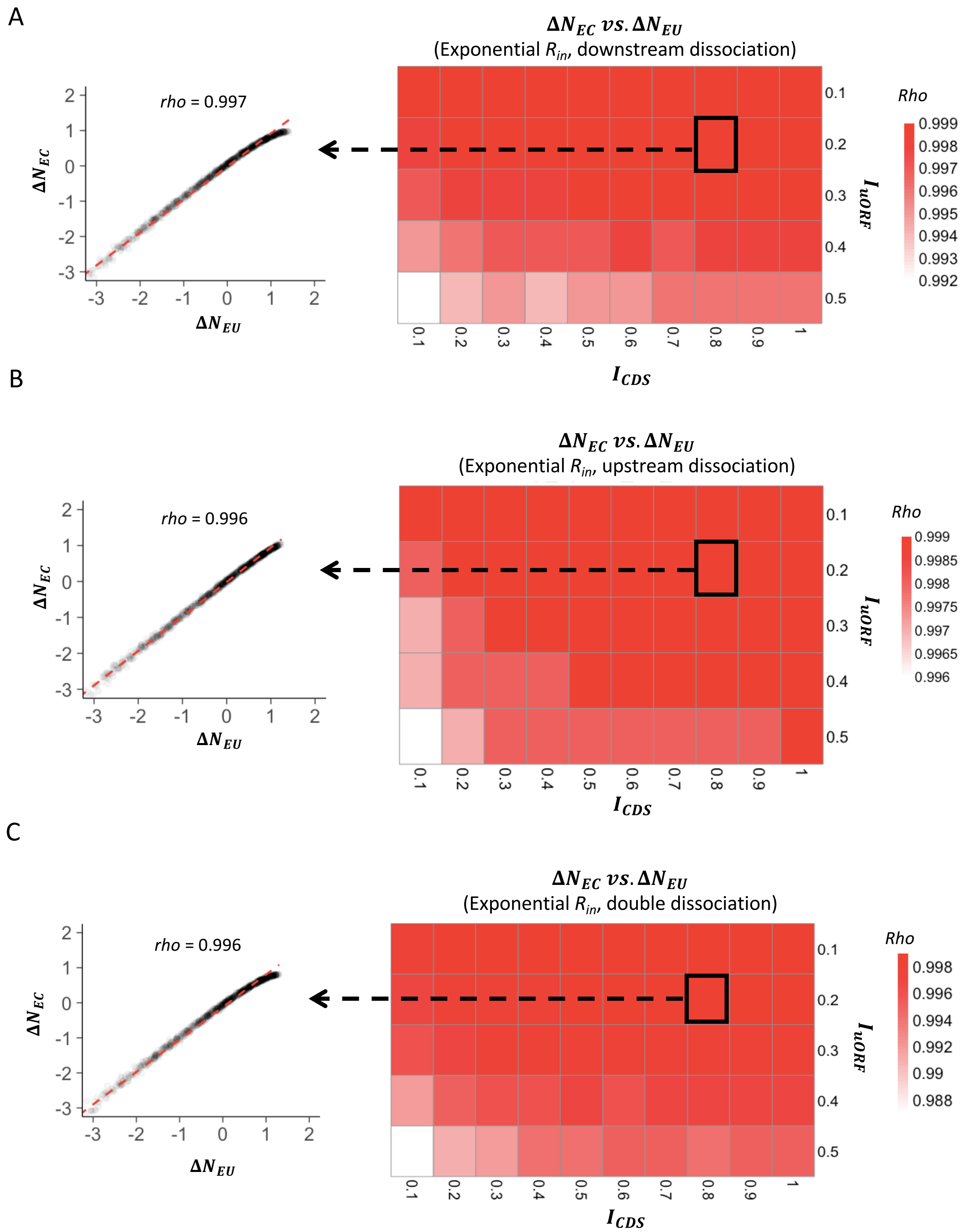


**Figure S9.** Heatmaps showing the Spearman's correlations between changes of uORF translation rate ($\Delta N_{EU}$) and downstream CDS translation rate ($\Delta N_{EC}$) under different $I_{CDS}$ (x-axis) and $I_{uORF}$ (y-axis) combinations with an exponential distribution of $R_{in}$ input and the downstream dissociation model (A), an exponential distribution of $R_{in}$ input and the upstream dissociation model (B), an exponential distribution of $R_{in}$ input and the double dissociation model (C). For each heatmap, the left panel elicited by the dotted line from the specific square of right heatmap was an example showing the Spearman's correlations between changes of uORF translation rate ($\Delta N_{EU}$) and downstream CDS translation rate ($\Delta N_{EC}$) under $I_{CDS}$ = 0.8 & $I_{uORF}$ = 0.2. All $P$ values < 0.001.

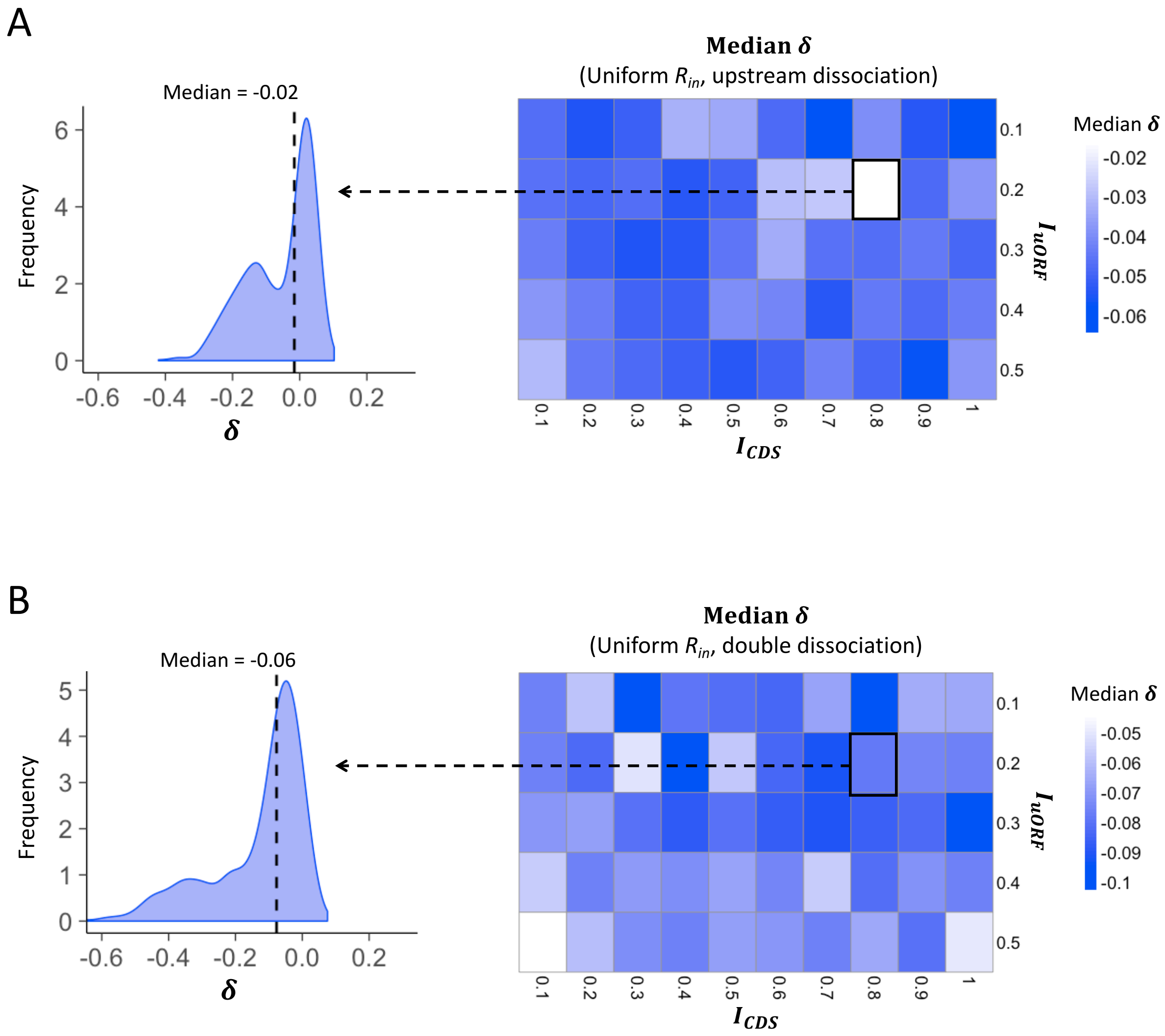


**Figure S10.** Heatmaps showing the median $\delta$ under different $I_{CDS}$ (x-axis) and $I_{uORF}$ (y-axis) combinations with a uniform distribution of $R_{in}$ input and the upstream dissociation model (A), a uniform distribution of $R_{in}$ input, and the double dissociation model (B). For each heatmap, the left panel elicited by the dotted line from the specific square of right heatmap was an example showing the distribution of $\delta$ under $I_{CDS}$ = 0.8 & $I_{uORF}$ = 0.2. The vertical dashed line indicated the median value of $\delta$.

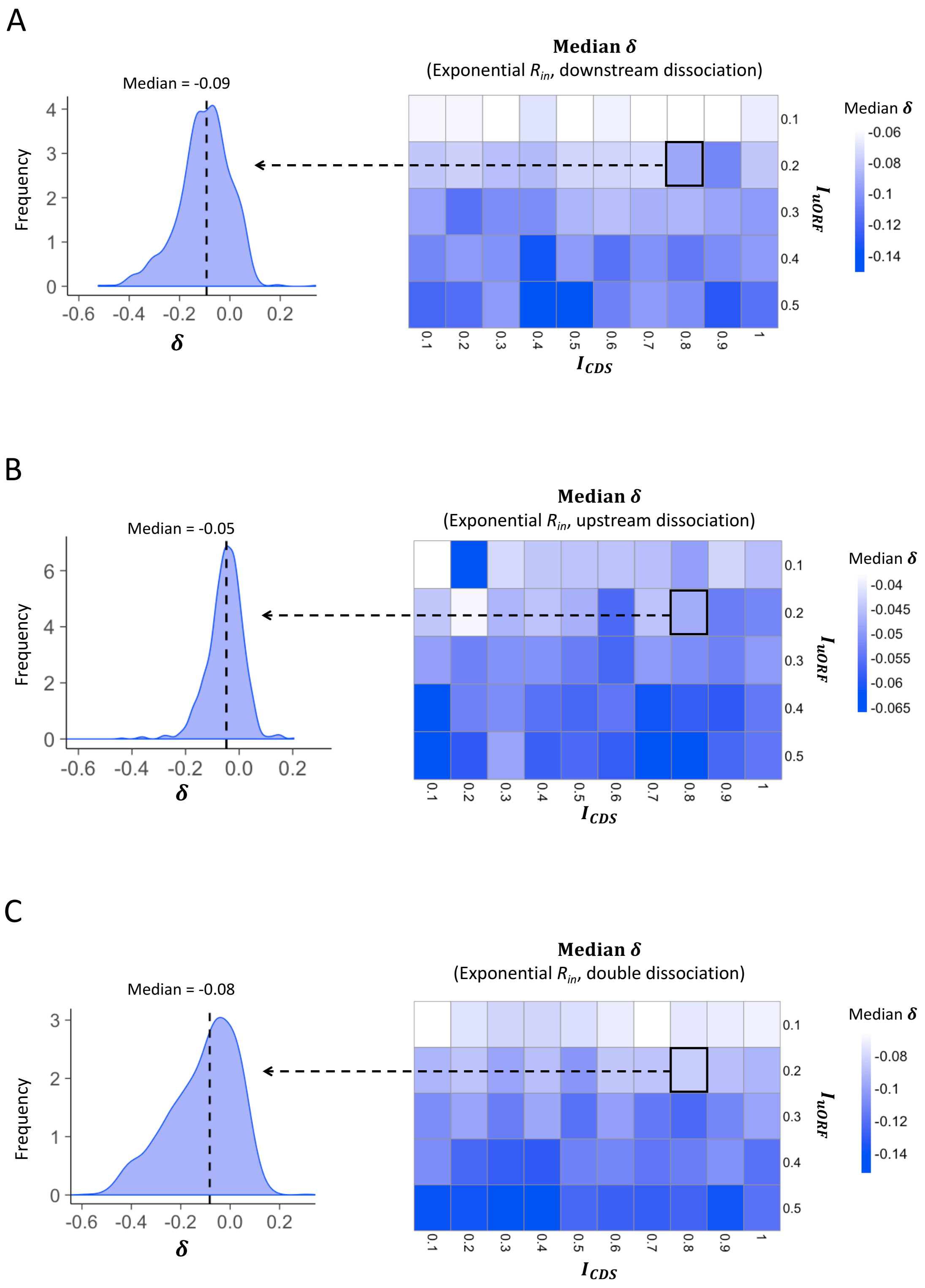


**Figure S11.** Heatmaps showing the median $\delta$ under different $I_{CDS}$ (x-axis) and $I_{uORF}$ (y-axis) combinations with an exponential distribution of $R_{in}$ input and the downstream dissociation model (A), an exponential distribution of $R_{in}$ input and the upstream dissociation model (B), an exponential distribution of $R_{in}$ input and the double dissociation model (C). For each heatmap, the left panel elicited by the dotted line from the specific square of right heatmap was an example showing the distribution of $\delta$ under $I_{CDS} = 0.8$ & $I_{uORF} = 0.2$. The vertical dashed line indicated the median value of $\delta$.

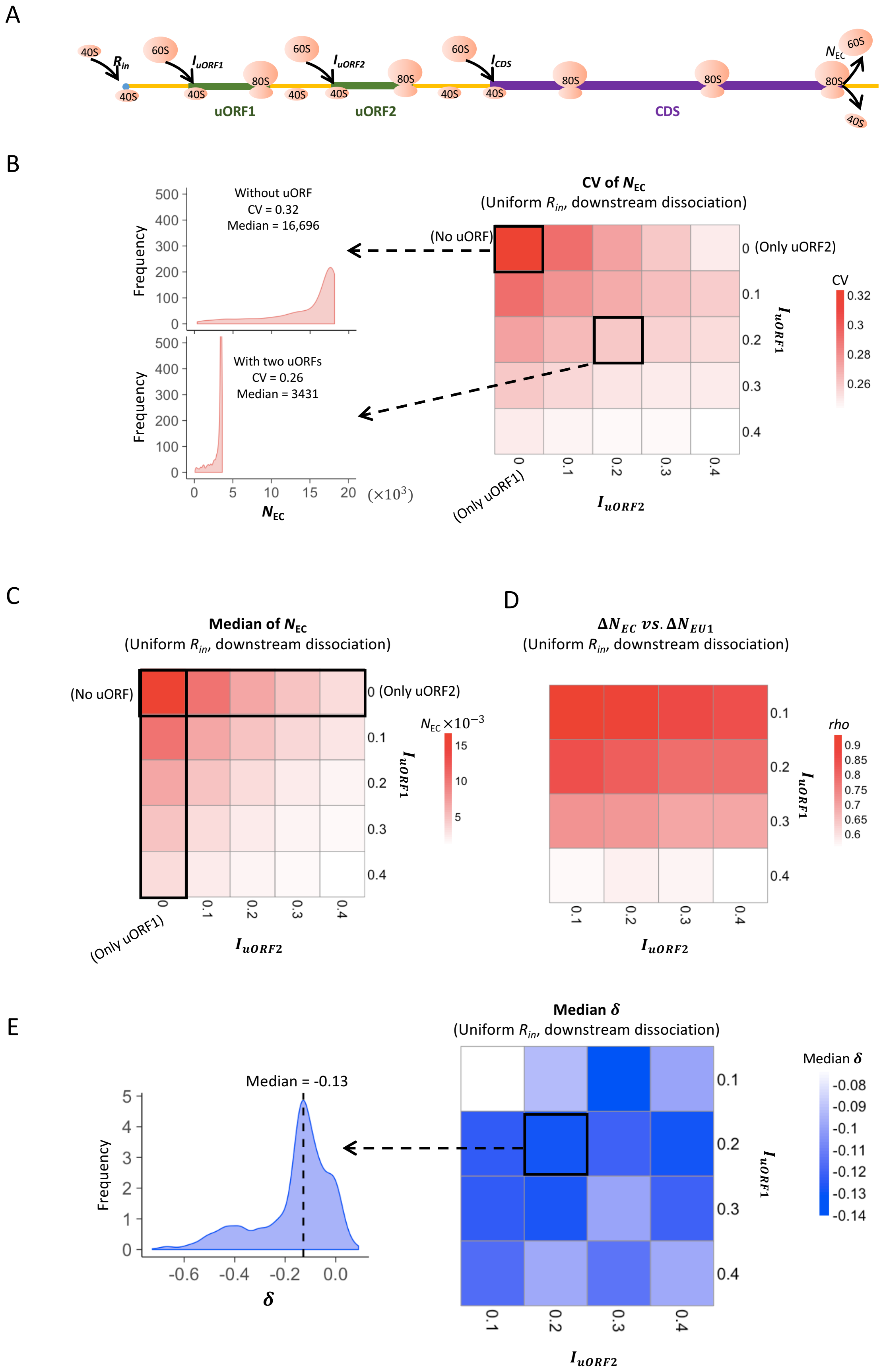


**Figure S12. Two-uORF model simulation measurements.** (A) The two-uORF model schema. The factors and parameters were the same as those illustrated in Fig. 1A except where specifically indicated. (B) Heatmap showing the CVs of CDS translation rate ($N_{EC}$) under different $I_{uORF2}$ (x-axis) and $I_{uORF1}$ (y-axis) combinations with a uniform distribution of $R_{in}$ input and the downstream dissociation model. The left panels elicited by the dotted lines from specific squares of right heatmap were two examples showing the distribution of $N_{EC}$ under $I_{uORF1}$ = 0 & $I_{uORF2}$ = 0 (top panel, without uORF) and $I_{uORF1}$ = 0.2 & $I_{uORF2}$ = 0.2 (bottom panel, with two uORFs). (C) Heatmap showing the median CDS translation rate ($N_{EC}$) under different $I_{uORF2}$ (x-axis) and $I_{uORF1}$ (y-axis) combinations with a uniform distribution of $R_{in}$ input and the downstream dissociation model. (D) Spearman's correlations between changes of uORF1 translation rate ($\Delta N_{EU1}$) and downstream CDS translation rate ($\Delta N_{EC}$) under different $I_{uORF2}$ (x-axis) and $I_{uORF1}$ (y-axis) combinations with a uniform distribution of $R_{in}$ input and the downstream dissociation model. (E) Heatmap showing the median $\delta$ [$log_2(\Delta N_{EC}/\Delta N_{EU1})$] under different $I_{uORF2}$ (x-axis) and $I_{uORF1}$ (y-axis) combinations with a uniform distribution of $R_{in}$ input and the downstream dissociation model. The left panel elicited by the dotted line from the specific square of right heatmap was an example showing the distribution of $\delta$ under $I_{uORF1}$ = 0.2 & $I_{uORF2}$ = 0.2. The vertical dashed line indicated the median value of $\delta$.

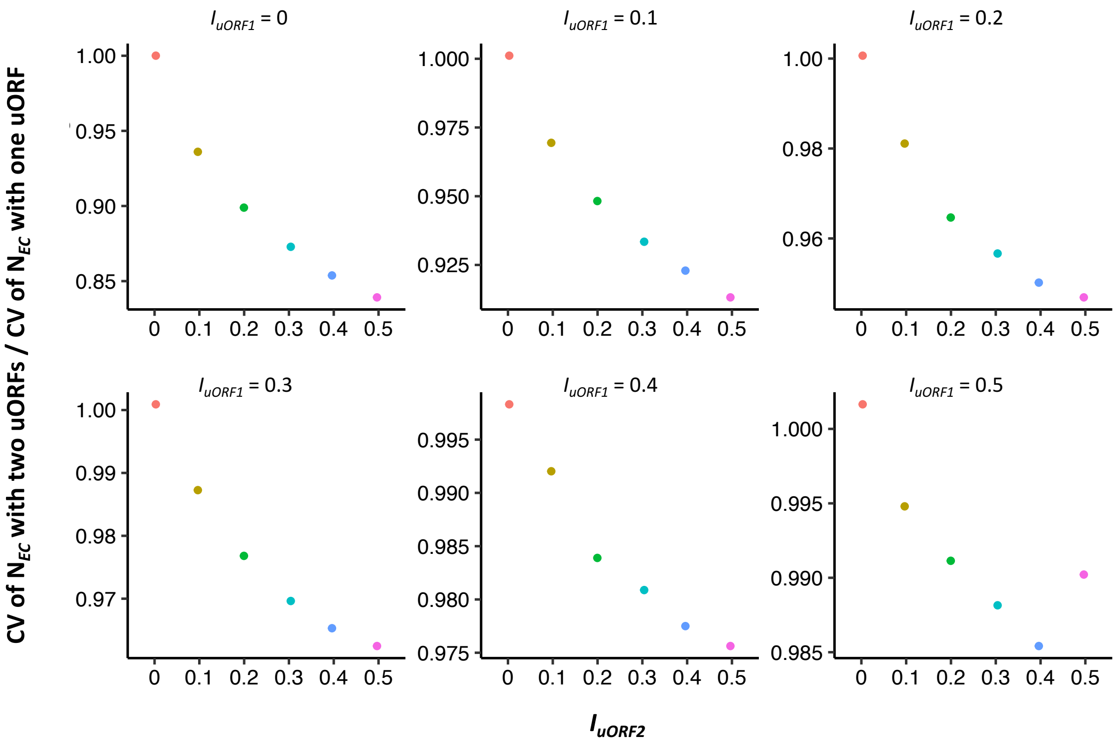


**Figure S13. Comparison of the buffering effects between single uORF and two uORFs.** In each panel, the $I_{uORF}$ of single-uORF model equals to $I_{uORF1}$ of two-uORF model, with both values ranging from 0 to 0.5. The x-axis in each panel denotes the values of different $I_{uORF2}$ in the two-uORF model, ranging from 0 to 0.5. The y-axis in each panel represents the ratio of the CV of $N_{EC}$ with two uORFs to that with a single uORF.

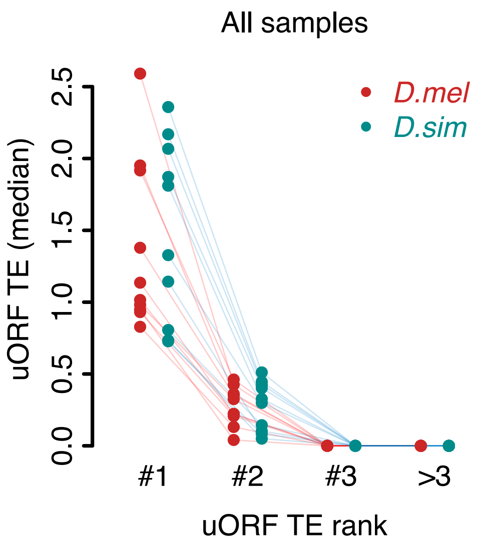


**Figure S14**. **The dominant uORF showed highest TE than other uORFs with a same gene.** uORFs were ranked by decreasing TEs within each gene. The uORF with the highest TE within each gene was defined as the dominant uORF (#1). "#2" represents the second highest uORF TE and the same goes for "#3" and ">3". Each dot represents the median TE of a sample.

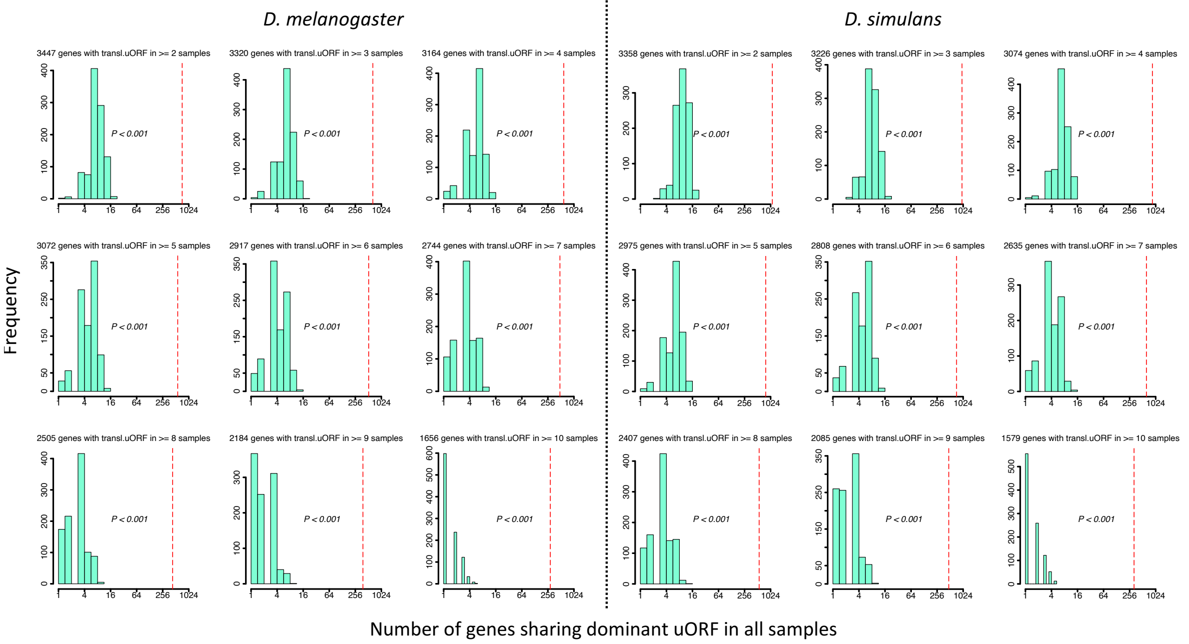


**Figure S15. Observed and expected numbers of genes sharing dominant uORFs in all samples.** Among the genes of *D. melanogaster* (where only the longest transcripts were considered), 7,259 (52.2%) genes had no uORF (“no-uORF” genes), 2,687 (19.3%) genes had one uORF (“one-uORF” genes), and 3,961 (28.5%) genes had multiple (≥ 2) uORFs (“multiple-uORF” genes). The numbers in the header of each panel represent genes with translated uORFs (TE > 0.1) in ≥ N samples (2 ≤ N ≤ 10). The TEs of all uORFs were shuffled 1,000 times. *P* values were obtained from randomization tests.

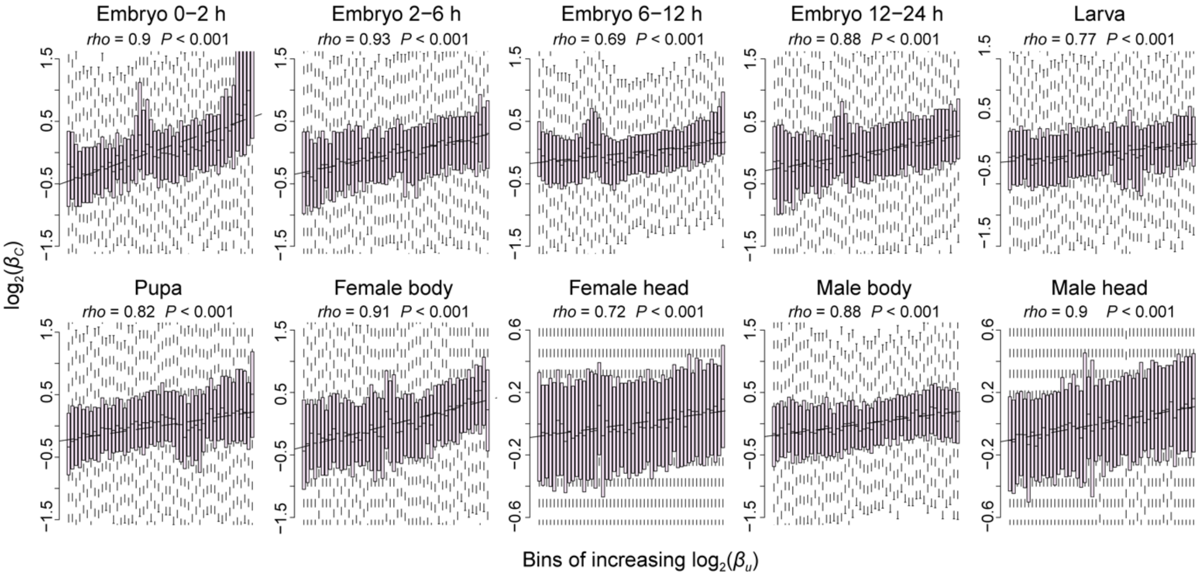


**Figure S16.The positive correlation of interspecific TE changes between uORFs and CDSs.** Correlations between interspecific uORF TE changes ($\log_2\beta_u$) and CDS TE changes ($\log_2\beta_C$) in 10 samples. The x-axis was divided into 50 equal bins with increasing $\beta_u$. Spearman's correlation coefficients (*rho*) are shown at the top left. ***, $P < 0.001$ in the correlation test.

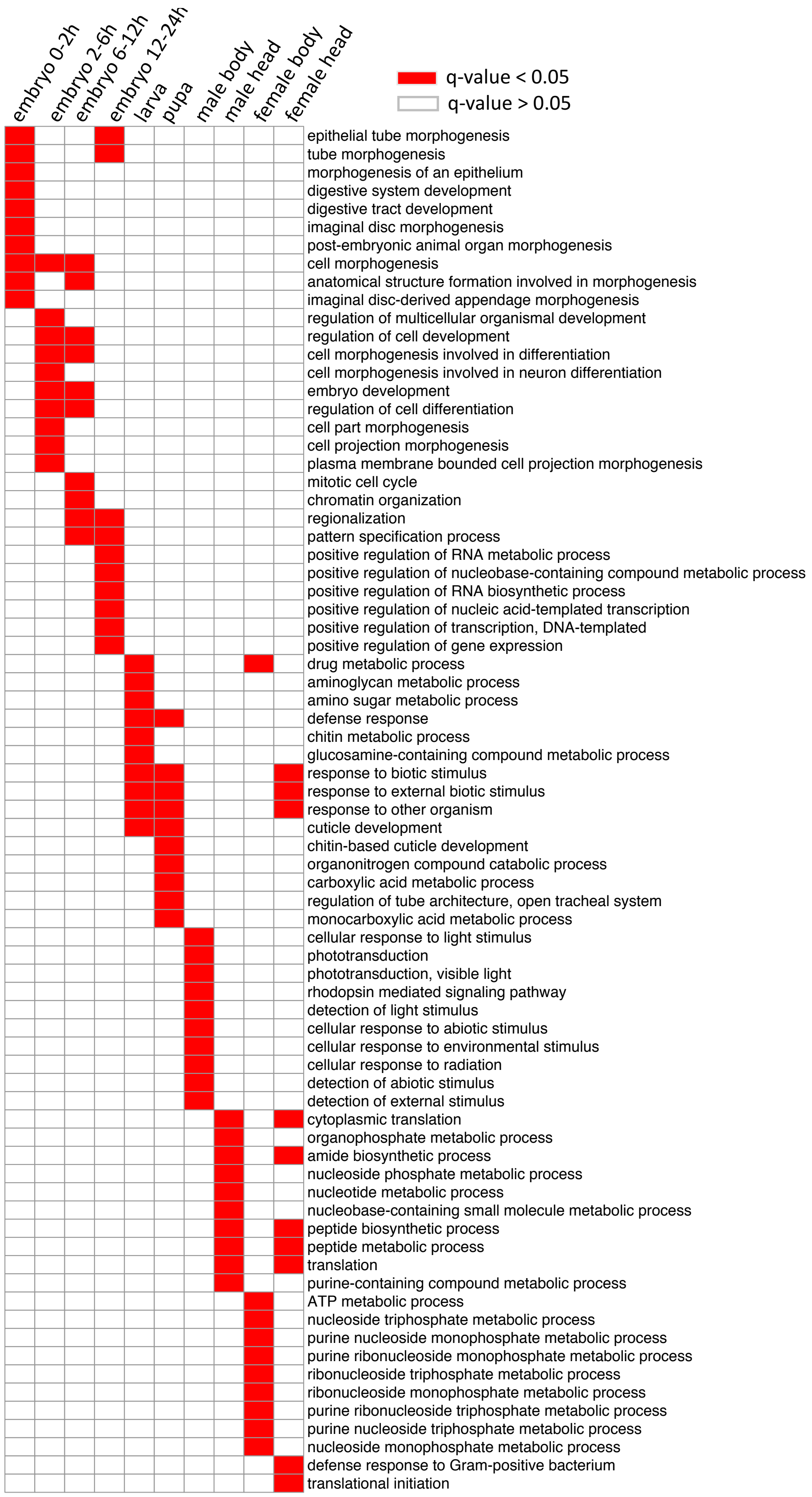


**Figure S17. Gene ontology analysis of the genes with $log_2(\beta_c) \neq 0$ in each stage and tissue.** The biological process (BP) terms with q-values < 0.05 in each sample type are indicated with red, and others are indicated with white.

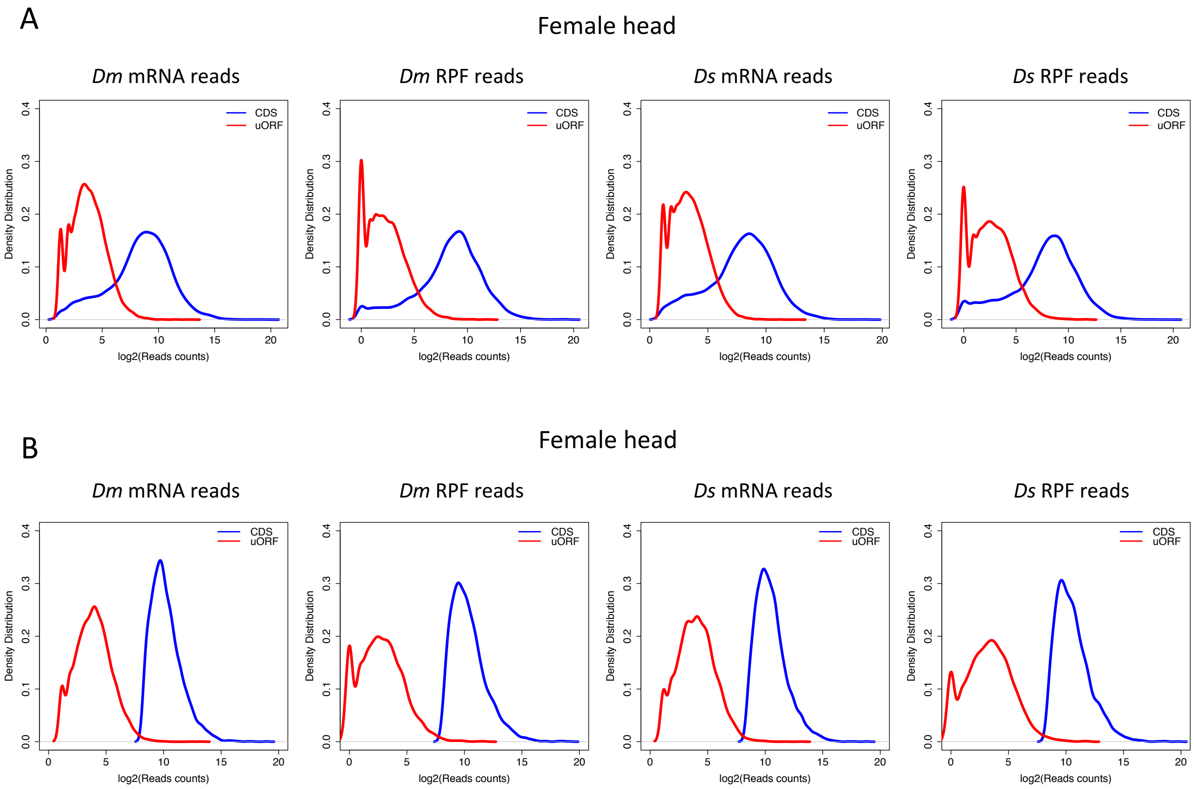


**Figure S18. Reads count distribution of uORFs and CDSs.** Distribution of mRNA reads counts and RPF counts mapped to uORFs and CDSs for all expressed uORFs (A) or only highly expressed genes (B) in female head sample. The distribution patterns were similar across other samples, and the data are not shown here. *Dm*, *D. melanogaster*; *Ds*, *D. simulans*. The reads counts were $\log_2$ transformed.

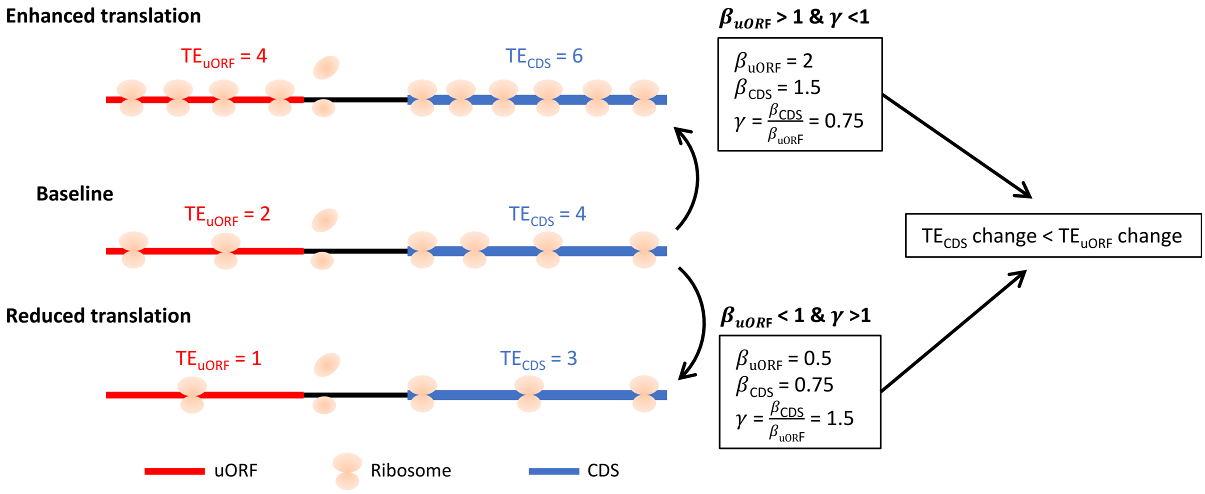


**Figure S19. The scheme illustrating the calculation of $\beta_u$, $\beta_c$ and $\gamma$.** When the cellular environment causes mRNA translation to be enhanced from baseline level, the TE$_{uORF}$ changed from 2 to 4 ($\beta_u$ = 2), while the TE$_{uORF}$ changed with a smaller degree due to uORF's buffering, from 4 to 6 ($\beta_C$ = 1.5). This resulted in $\gamma$ <1. Conversely, When the cellular environment causes mRNA translation to be reduced from the baseline level, the TE$_{uORF}$ changed from 2 to 1 ($\beta_u$ = 0.5), while the TE$_{uORF}$ changed with a smaller degree due to uORF's buffering, from 4 to 3 ($\beta_C$ = 0.76). This resulted in $\gamma$ >1. Overall, both $\beta_u$ >1 &$\gamma$ <1, and $\beta_u$ <1 &$\gamma$ >1, indicated the existence of uORFs' buffering.

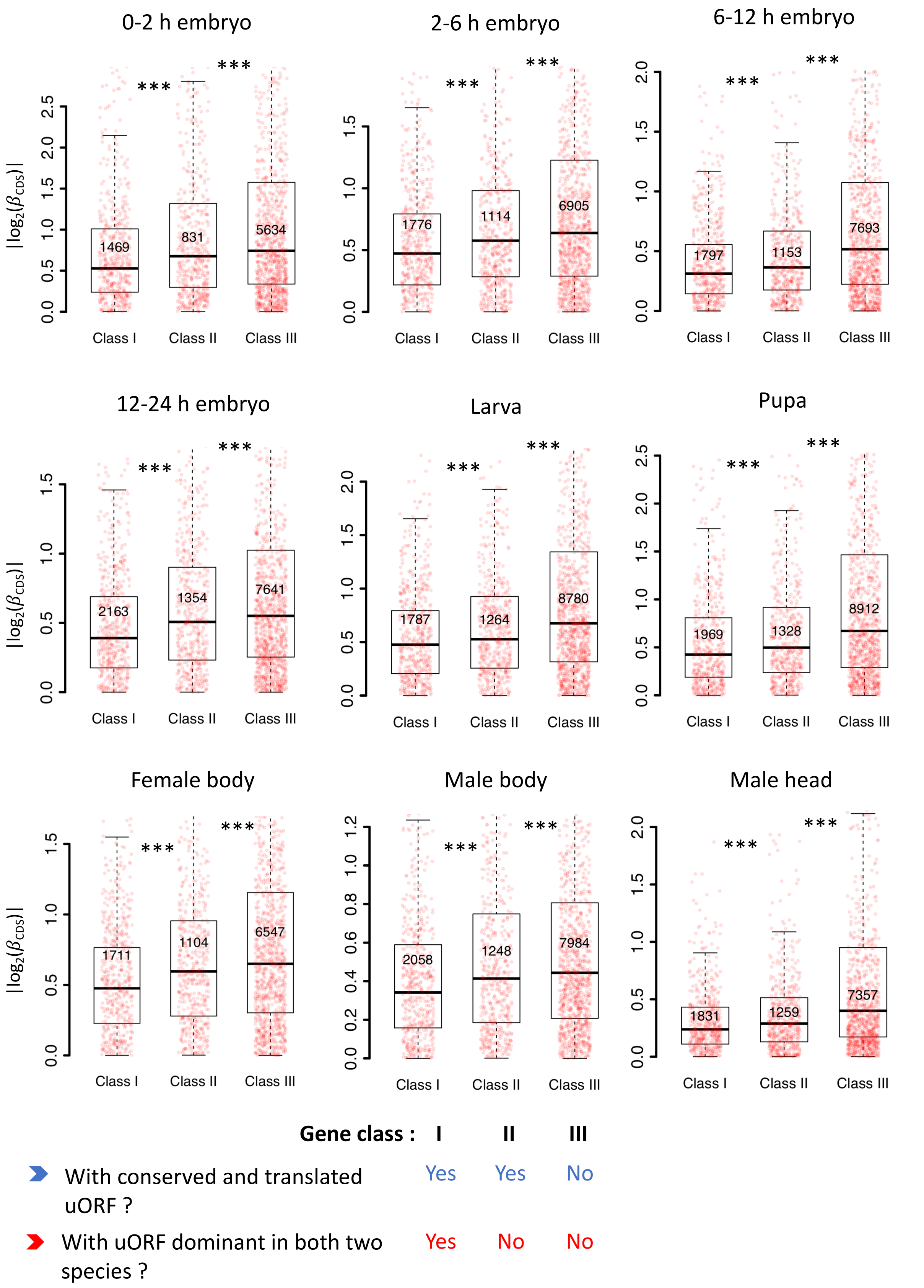


**Figure S20. Conserved and dominantly translated uORFs showed the stronger buffering effect.** Genes expressed in different stages/tissues (mRNA RPKM > 0.1 in both species) were classified into three classes according to whether a gene had a conserved and dominantly translated uORF or not.

Boxplots showing interspecific CDS TE variability $|log_2(\beta_c)|$ of different gene classes. $P$ values were calculated using Wilcoxon rank-sum tests between the neighboring groups. ***, $P < 0.001$.

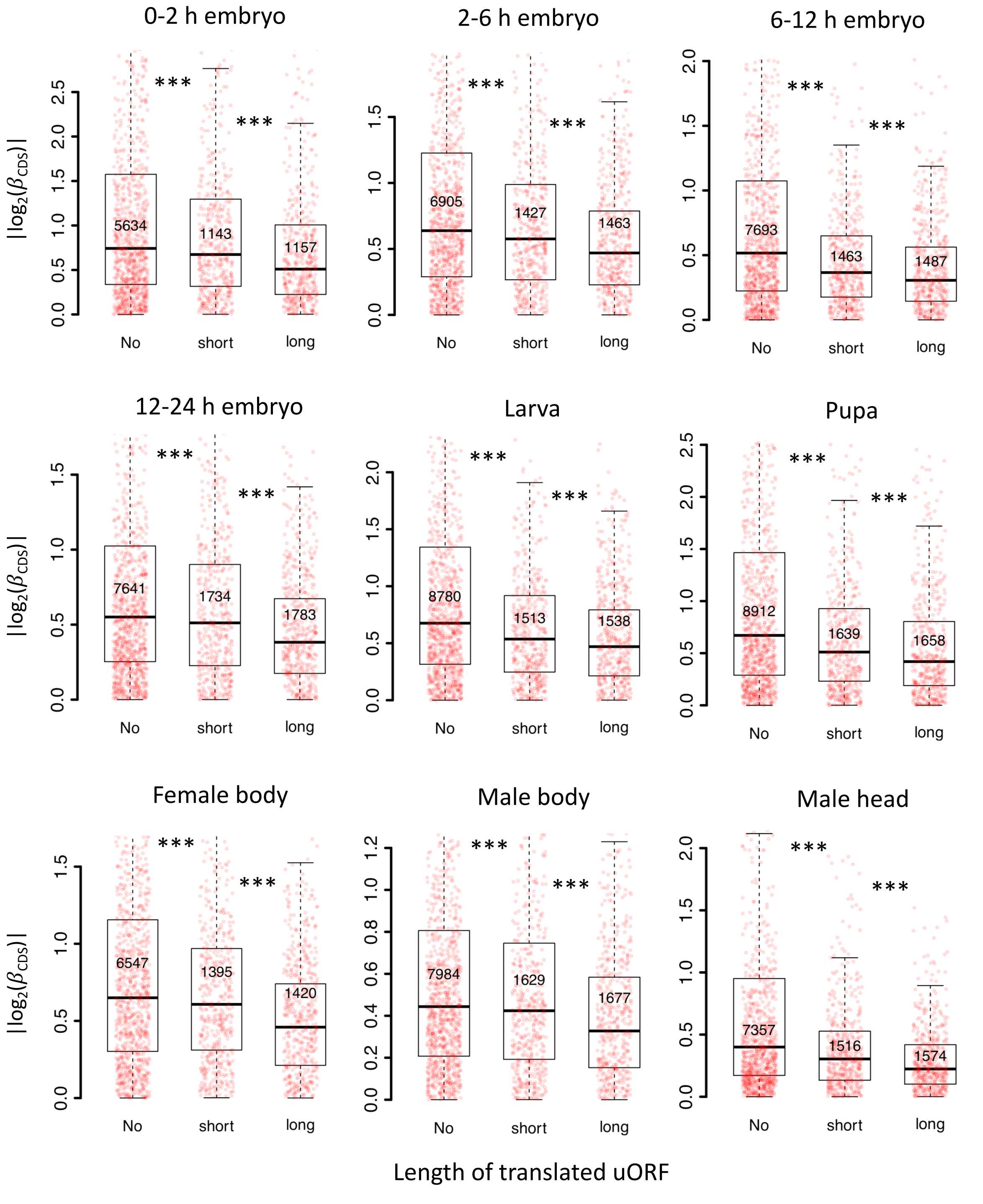


**Figure S21. Longer uORFs showed the stronger buffering effect.** Genes expressed in different stages/tissues (mRNA RPKM > 0.1 in both species) were classified into three classes according to the length of translated uORFs. Boxplots showing interspecific CDS TE variability $|log_2(\beta_c)|$ of different gene classes. *P* values were calculated using Wilcoxon rank-sum tests between the neighboring groups. ***, $P < 0.001$.

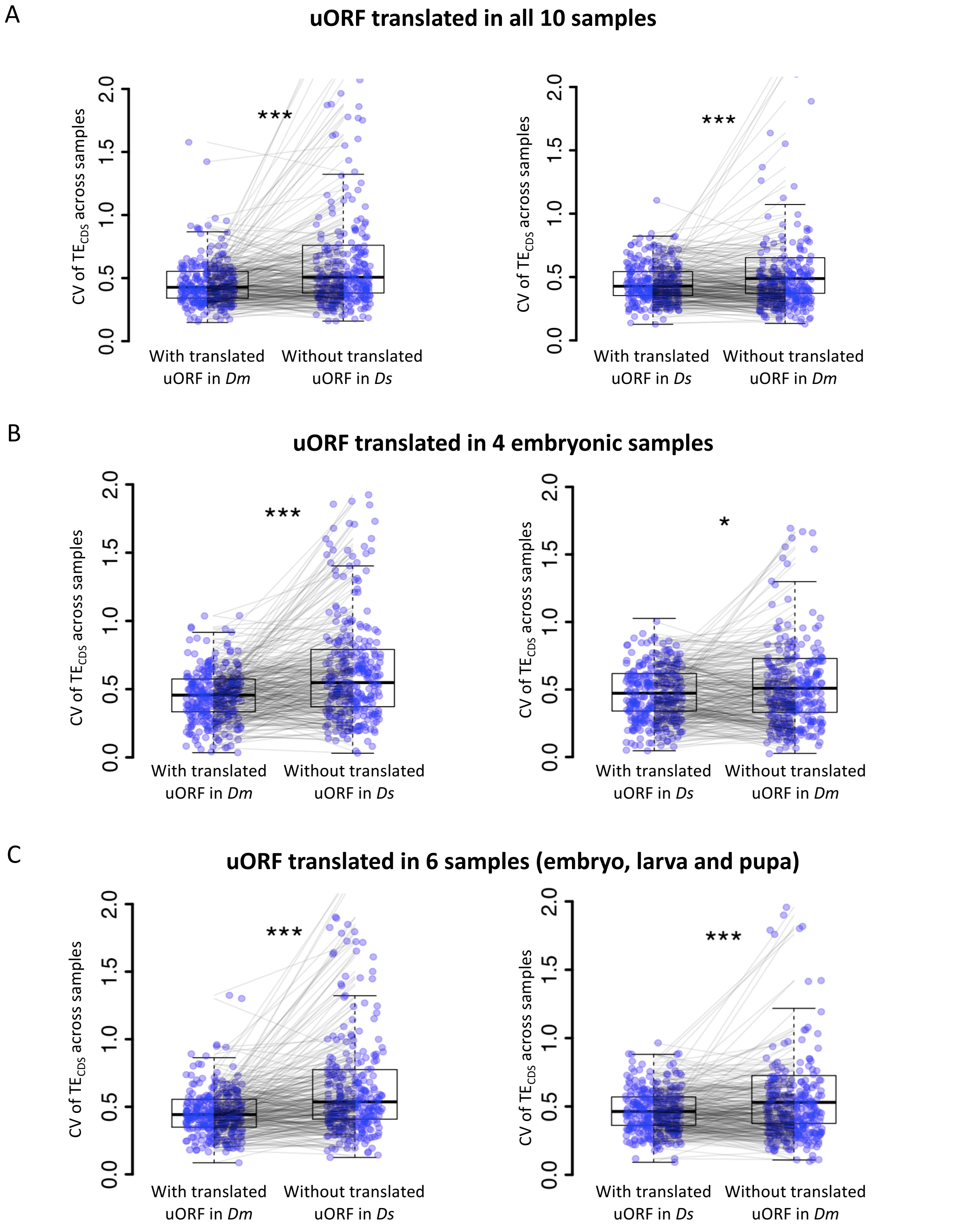


**Figure S22. uORFs reduce CDS translational fluctuation during *Drosophila* development under different cutoffs on defining "translated uORFs".** The CV of $TE_{CDS}$ across *Dm* (*D. melanogaster*) samples and *Ds* (*D. simulans*) samples was shown as boxplots. The selected genes harbor the translated uORFs in one of *Drosophila* species but its homologous gene without translated uORFs in another *Drosophila* species. Each pair of dots linked by a gray line represents a pair of homologous genes in *Dm* and *Ds*. (A) Translated uORF was defined as uORF translated (TE > 0.1) in all 10 samples. (B) Translated uORF was defined as uORF translated (TE > 0.1) in 4 embryonic stages. The CV of $TE_{CDS}$ across these 4 stages was calculated. (C) Translated uORF was defined as uORF translated (TE > 0.1) in the 6 developmental stages including 4 embryonic stages, larva, and pupa.

The CV of $TE_{CDS}$ across these 6 stages was calculated. Wilcoxon signed-rank sum test. *, $P < 0.05$; ***, $P < 0.001$.

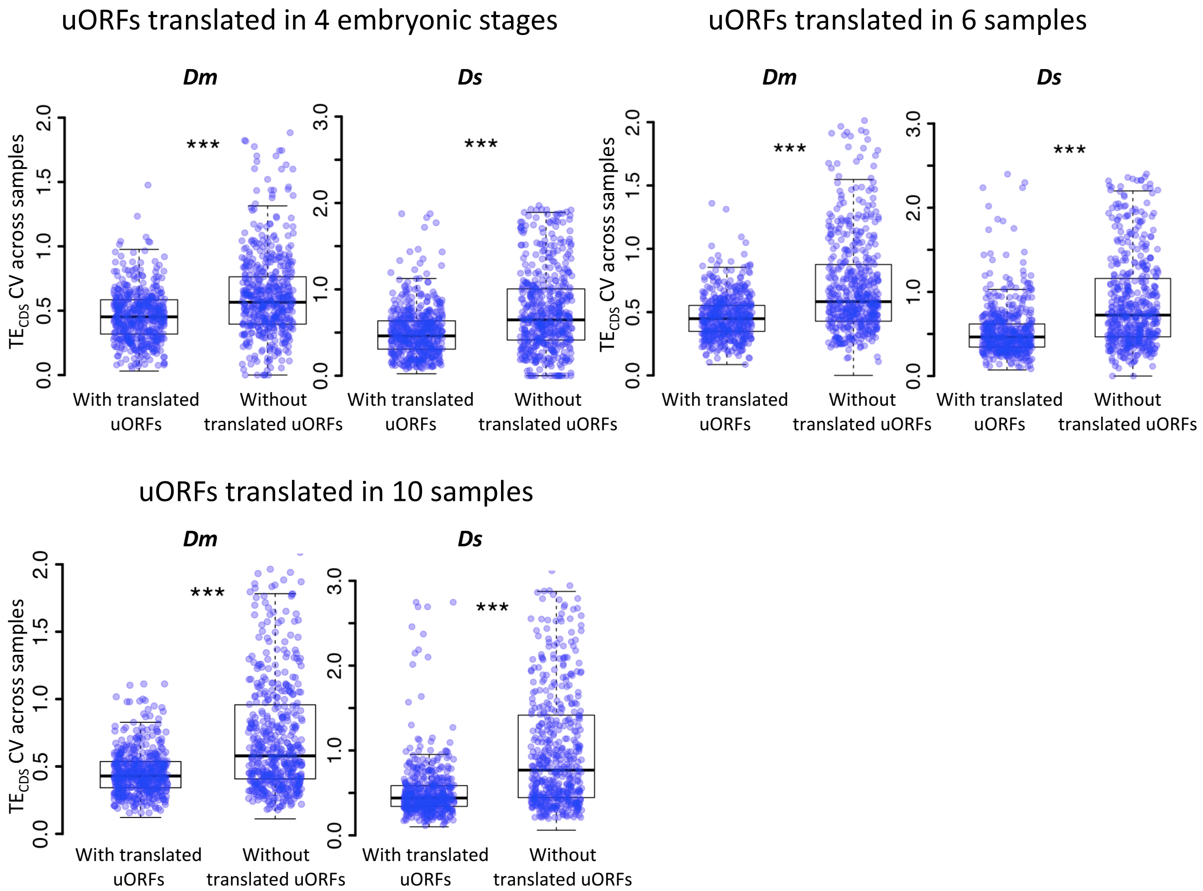


**Figure S23. uORFs could reduce CDS translational variation during *Drosophila* development under different cutoffs on defining "translated uORFs".** Within each species, the CV of $TE_{CDS}$ across different samples was measured for genes with translated uORFs and genes without translated uORFs, respectively. Each dot in the boxplot represents one gene. Translated uORF was defined as uORF translated (TE > 0.1) in at least 4 embryonic stages, six developmental stages (left) or translated or in all 10 samples. Wilcoxon rank-sum tests. ***, $P < 0.001$. *Dm*, *D. melanogaster*; *Ds*, *D. simulans*.

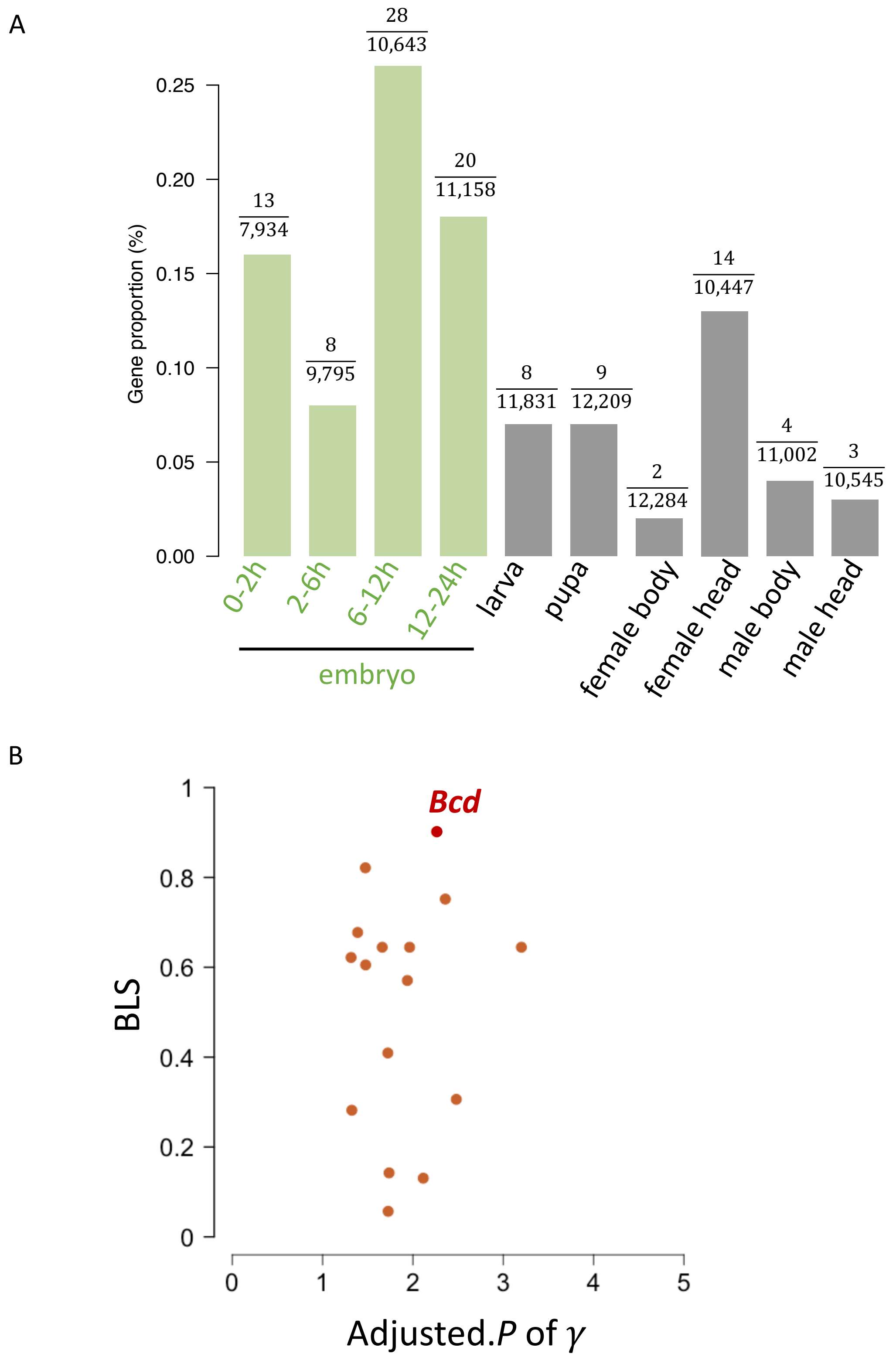


**Figure S24**. (A) The proportion (%) of genes with buffering uORFs in all expressed genes across developmental stages/tissues. (B) The distribution of adjusted. *P* of γ (x-axis) and BLS (y-axis) of dominant uORFs in embryonic stages. The *bcd* was highlighted by red color.

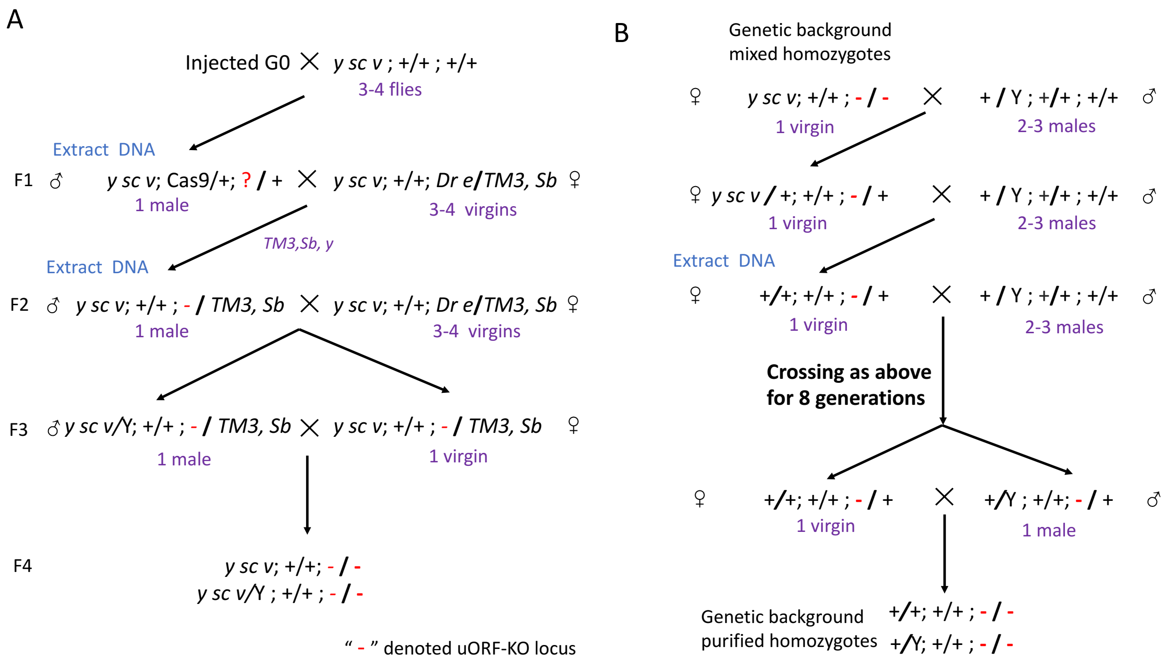


**Figure S25. Overview of homozygous uORF-KO mutant screening.** (A) Procedures for the primary screening of CRISPR/Cas9-induced mutants. Red "-" denotes the *bcd*-uORF-KO allele. (B) Purification of the genetic background of homozygous mutants by back-crossing with WT lines for nine generations.

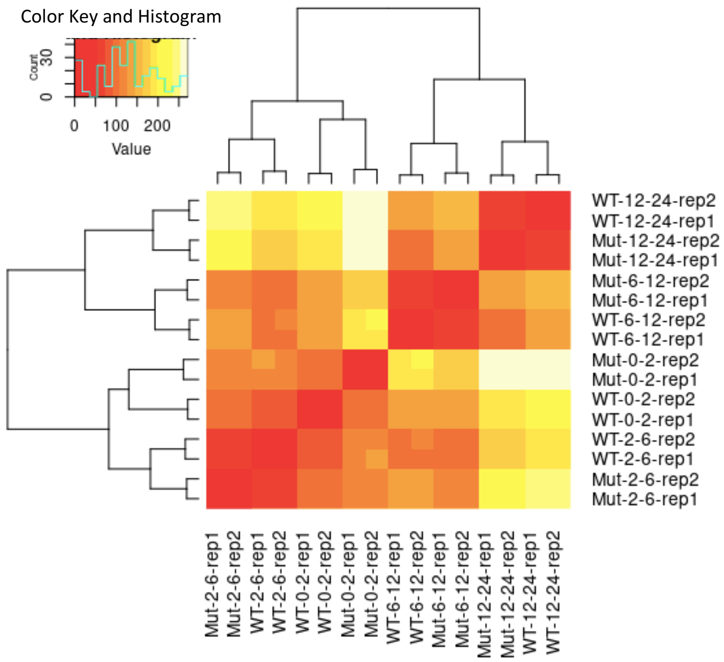


**Figure S26. Heatmap of sample-to-sample distances of RNA-Seq libraries. Mut, mutant; WT, wild-type; rep1/rep2, biological replicate 1/biological replicate 2.**

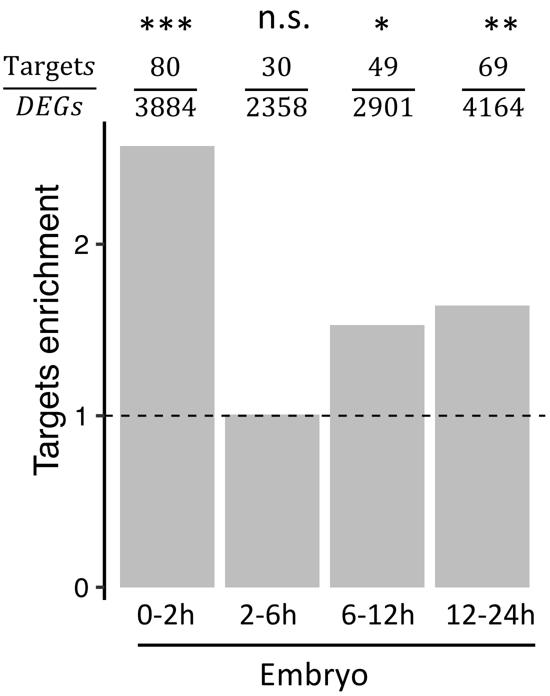


**Figure S27.** (A) The enrichment of Bcd targets in each embryo stage at 29°C. Enrichment was calculated as the ratio of the Bcd target proportion in DEGs to the Bcd target proportion in non-DEGs. $\chi^2$ tests were performed to compare the difference in Bcd target distribution in DEGs and non-DEGs. The number of DEGs and the number of Bcd targets in DEGs for each stage are provided above the bar. Asterisks indicate statistical significance ($\chi^2$ tests; *, $P < 0.05$; **, $P < 0.01$; ***, $P < 0.001$; n.s., $P > 0.05$). (B) The expression [$\log_2$(fold change)] levels of *bcd* and 3 of its targets, *Kr*, *btd*, and *otd,* in mutants compared to those in the WT in 4 embryonic stages. Asterisks are present at the top of bars only for |fold-changes| > 1.5 and adjusted *P* values < 0.05 (*, $P < 0.05$; **, $P < 0.01$; ***, $P < 0.001$). (C) Expression of *bcd* and its targets (*Kr*, *btd*, *otd*) measured by RNA-Seq in uKO2/uKO2 mutant (red line) and WT (blue line) of *D. melanogaster*. The expression of each gene in each stage is shown as the RPKM value.

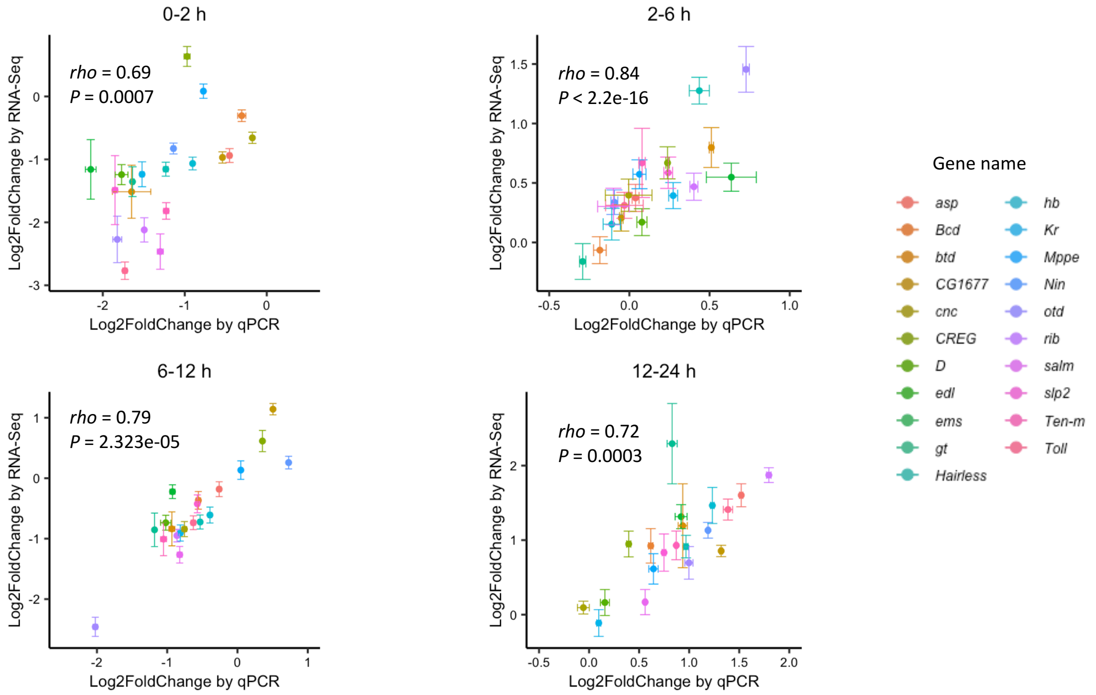


**Figure S28.** The correlation of expression changes (log$_2$FoldChange) between RNA-Seq (y-axis) and RT-qPCR (x-axis) of 20 *bcd* targets in four embryo stages in uKO2/uKO2 mutant compared to WT.

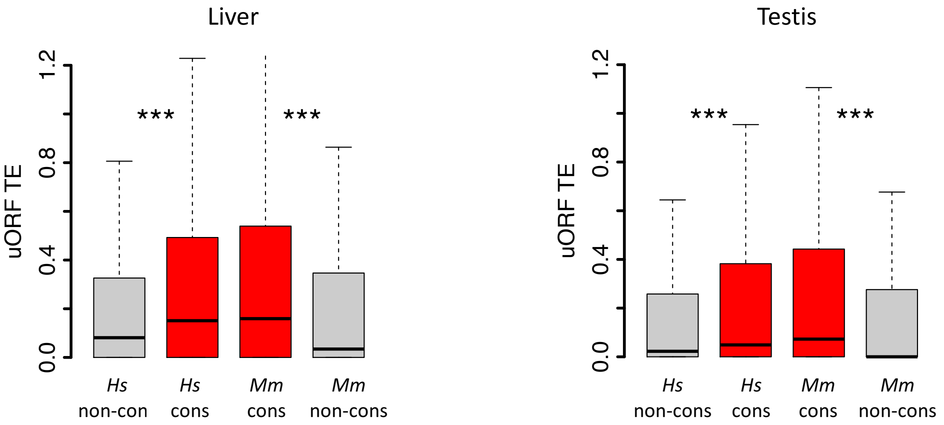


**Figure S29.** Boxplots showing the TEs of conserved and species-specific uORFs in the liver and testis of *H. sapiens* and *M. mulatta*. Wilcoxon rank-sum tests. ***, $P < 0.001$.

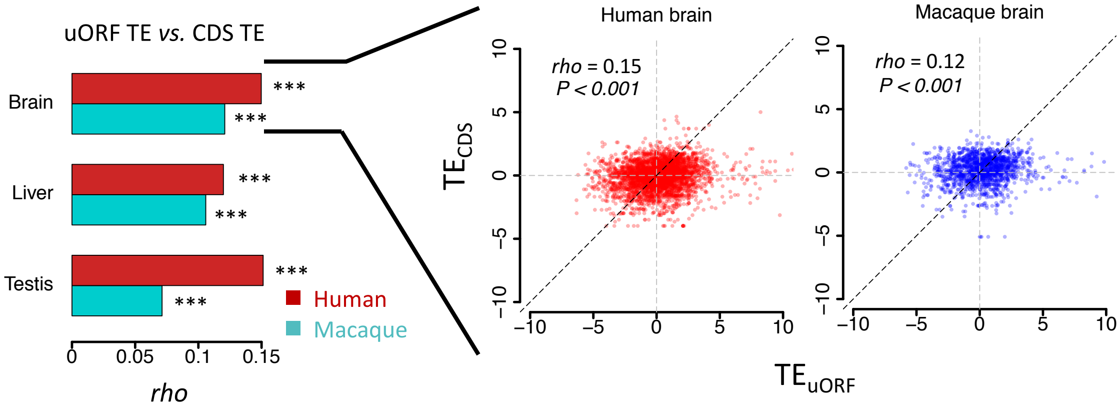


**Figure S30.** Correlations of uORF TE and corresponding CDS TE in 3 tissues of *H. sapiens* and *M. mulatta*. The bars represent Spearman's correlation coefficient (*rho*). In all samples, we obtained *P* values < 0.001. Data from brains are shown as examples in the right panel.

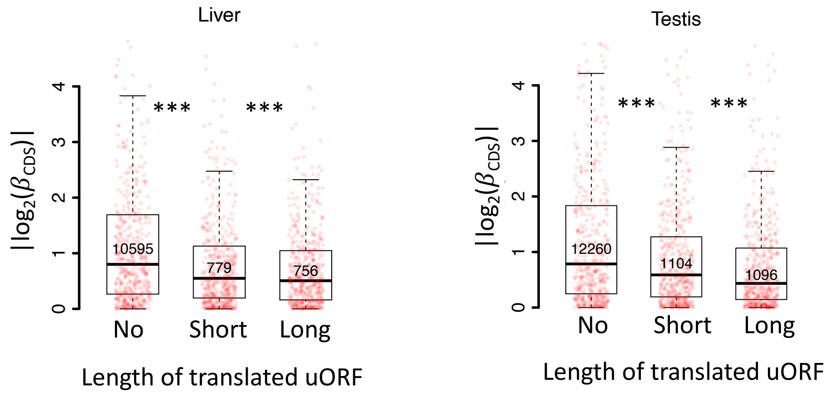


**Figure S31.** Boxplots showing interspecific CDS TE variability $|log_2(\beta_{CDS})|$ for three gene groups according to the length of their translated uORFs (No, short, long) in the liver (left) and testis (testis). Wilcoxon rank-sum tests. ***, $P < 0.001$.

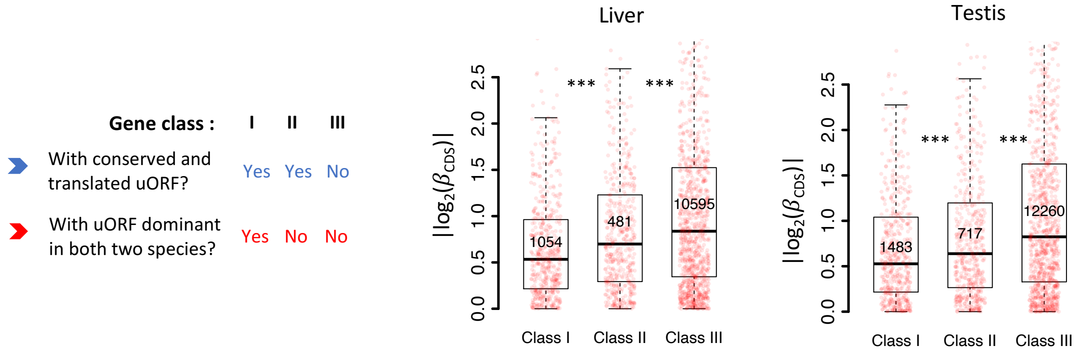


**Figure S32.** Boxplots showing interspecific CDS TE variability $|log_2(\beta_{CDS})|$ for different gene classes in the liver and testis. Genes expressed in liver or testis (mRNA RPKM > 0.1 in both species) were classified into three classes according to whether a gene had a conserved and dominant translated uORF (TE > 0.1) in both species or not. Wilcoxon rank-sum tests. ***, $P < 0.001$.

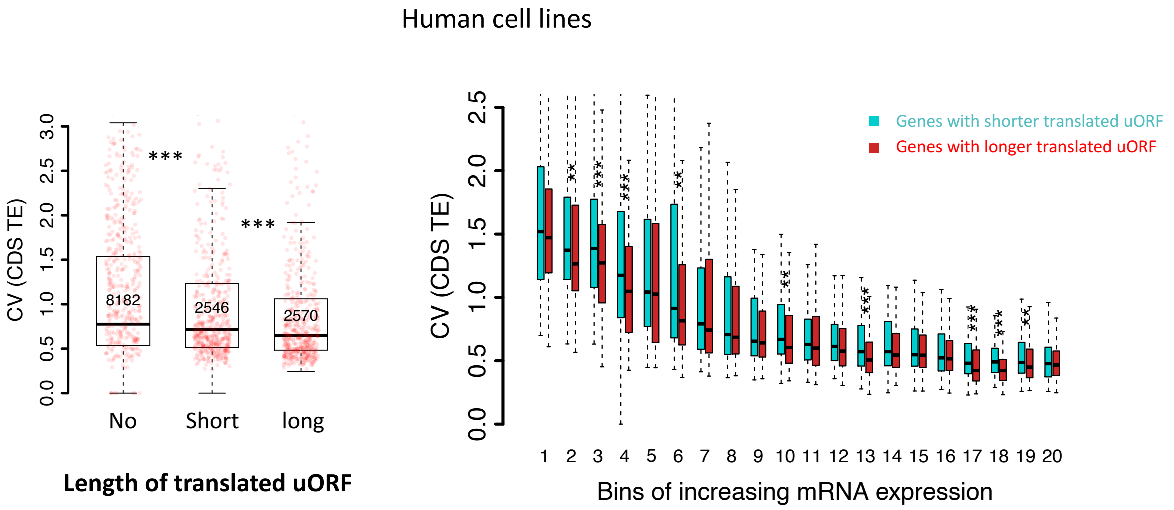


**Figure S33.** Left panel: Boxplot showing the coefficients of variation (CVs) of CDS TEs among 69 human lymphoblastoid cell lines (LCLs) for three gene groups. Genes were classified into 3 groups according to the total length of translated uORFs. Right panel: boxplot showing the coefficients of variation (CVs) of CDS TEs among the human cell lines. Genes with translated uORFs were divided into 20 bins with increasing mRNA expression levels. In each bin, genes were divided into two fractions according to the total length of translated uORFs.

**Table S1. Parameters used in our simulation**

| Symbol | Description | Value |
|---|---|---|
| $Len\_a$ | Length of 5'-leader before uORF | 50 triplets |
| $Len\_u$ | Length of uORF | Default: 30 triplets<br>Variable: 2, 10, 20, 30, 40, 50, 100 triplets |
| $Len\_b$ | Distance between uORF and CDS | 50 triplets |
| $Len\_m$ | Length of CDS | 500 triplets |
| $Len\_c$ | Length of 3'UTR | 50 triplets |
| $\nu_s$ | Probability of movement of a 40S ribosome to the next position in a single action | 0.3* |
| $\nu_{Eu}$ | Probability of movement of an 80S ribosome to the next position in uORF in a single action | 0.3** |
| $\nu_{EC}$ | Probability of movement of an 80S ribosome to the next position in CDS in a single action | 0.5*** |
| $R_{in}$ | Probability of loading a new 40S ribosome at the 5'-terminus of the mRNA in a single action | 1000 values generated from uniform distribution or exponential distribution |
| $K_{up}$ | Dissociation probability of upstream 40S ribosome colliding with downstream 80S ribosome | 0 for downstream dissociation;<br>1 for upstream or double dissociation |
| $K_{down}$ | Dissociation probability of downstream 40S ribosome colliding with upstream 80S ribosome | 0 for upstream dissociation;<br>1 for downstream or double dissociation |
| $I_{uORF}$ | Probability of translation initiation at the uORF start codon in a single action | Single-uORF model:<br>0, 0.1, 0.2, 0.3, 0.4, 0.5<br>Double-uORF model:<br>0, 0.1, 0.2, 0.3, 0.4 |
| $I_{CDS}$ | Probability of translation initiation at the CDS start codon in a single action | Single-uORF model:<br>0.1, 0.2, 0.3, 0.4, 0.5, 0.6, 0.7, 0.8, 0.9, 1.0<br>Double-uORF model:<br>0.9 |

*Adopted from Andreev *et al.*'s original ICIER model [1], corresponding to a movement rate of 5 triplets/s.
**Adopted from Andreev *et al.*'s original ICIER model [1], corresponding to a movement rate of 5 codons/s.
*** Considering that uORFs usually encode blocking peptides [2-6] or contain stalling codons [7-9], we set $\nu_{Eu}$ slightly lower than $\nu_{EC}$.

**Table S2.** Mapping statistics of Ribo-Seq and matched mRNA-Seq libraries

| Sample | Species* | Library type | Total reads (M) | After quality control (M) ** | Unique mapping rate (%) | Multiple mapping rate (%) |
|---|---|---|---|---|---|---|
| **0-2 h embryos** | *D. melanogaster* | mRNA-Seq | 61.01 | 52.74 | 93.14 | 2.84 |
| **2-6 h embryos** | *D. melanogaster* | mRNA-Seq | 59.44 | 52.07 | 93.26 | 3.24 |
| **6-12 h embryos** | *D. melanogaster* | mRNA-Seq | 53.56 | 45.11 | 92.33 | 3.09 |
| **12-24 h embryos** | *D. melanogaster* | mRNA-Seq | 61.18 | 50.64 | 91.89 | 4.31 |
| **Third-instar larvae** | *D. melanogaster* | mRNA-Seq | 58.28 | 45.46 | 88.53 | 6.09 |
| **P7-8 pupae** | *D. melanogaster* | mRNA-Seq | 52.99 | 41.16 | 92.65 | 4.46 |
| **Female adult heads** | *D. melanogaster* | mRNA-Seq | 21.48 | 16.87 | 93.00 | 3.27 |
| **Male adult heads** | *D. melanogaster* | mRNA-Seq | 16.32 | 14.18 | 91.17 | 4.24 |
| **Female adult bodies rep1** | *D. melanogaster* | mRNA-Seq | 16.84 | 11.20 | 88.83 | 5.76 |
| **Female adult bodies rep2** | *D. melanogaster* | mRNA-Seq | 24.94 | 19.37 | 88.73 | 6.00 |
| **Male adult bodies rep1** | *D. melanogaster* | mRNA-Seq | 19.79 | 16.01 | 91.62 | 6.39 |
| **Male adult bodies rep2** | *D. melanogaster* | mRNA-Seq | 11.24 | 7.64 | 89.86 | 6.69 |
| **0-2 h embryos** | *D. melanogaster* | Ribo-Seq | 64.81 | 51.96 | 90.36 | 6.31 |
| **2-6 h embryos** | *D. melanogaster* | Ribo-Seq | 53.14 | 41.19 | 69.47 | 16.92 |
| **6-12 h embryos** | *D. melanogaster* | Ribo-Seq | 66.24 | 46.76 | 86.02 | 8.94 |
| **12-24 h embryos** | *D. melanogaster* | Ribo-Seq | 63.7 | 47.45 | 88.96 | 6.30 |
| **Third-instar larvae** | *D. melanogaster* | Ribo-Seq | 39.53 | 22.13 | 58.83 | 6.82 |
| **P7-8 pupae** | *D. melanogaster* | Ribo-Seq | 41.53 | 20.33 | 80.92 | 5.51 |
| **Female adult heads** | *D. melanogaster* | Ribo-Seq | 63.94 | 39.90 | 94.39 | 2.92 |
| **Male adult heads** | *D. melanogaster* | Ribo-Seq | 58.22 | 23.87 | 93.36 | 3.24 |
| **Female adult bodies rep1** | *D. melanogaster* | Ribo-Seq | 44.11 | 19.85 | 84.20 | 6.36 |
| **Female adult bodies rep2** | *D. melanogaster* | Ribo-Seq | 41.95 | 18.73 | 70.74 | 11.33 |
| **Male adult bodies rep1** | *D. melanogaster* | Ribo-Seq | 42.56 | 18.98 | 79.39 | 8.65 |
| **Male adult bodies rep2** | *D. melanogaster* | Ribo-Seq | 39.52 | 28.66 | 81.19 | 6.93 |
| **0-2 h embryos** | *D. simulans* | mRNA-Seq | 19.92 | 18.33 | 90.85 | 5.82 |
| **2-6 h embryos** | *D. simulans* | mRNA-Seq | 13.88 | 7.47 | 88.32 | 7.46 |
| **6-12 h embryos** | *D. simulans* | mRNA-Seq | 21.13 | 18.64 | 89.26 | 7.06 |
| **12-24 h embryos** | *D. simulans* | mRNA-Seq | 16.17 | 13.96 | 87.77 | 8.47 |
| **Third-instar larvae** | *D. simulans* | mRNA-Seq | 17.56 | 12.69 | 89.88 | 6.57 |
| **P7-8 pupae** | *D. simulans* | mRNA-Seq | 23.87 | 19.04 | 89.92 | 5.62 |
| **Female adult heads** | *D. simulans* | mRNA-Seq | 17.13 | 15.11 | 89.70 | 6.39 |
| **Male adult heads** | *D. simulans* | mRNA-Seq | 17.72 | 15.96 | 90.73 | 6.15 |
| **Female adult bodies** | *D. simulans* | mRNA-Seq | 21.64 | 15.37 | 85.24 | 9.20 |
| **Male adult bodies** | *D. simulans* | mRNA-Seq | 21.54 | 17.34 | 87.45 | 8.84 |
| **0-2 h embryos** | *D. simulans* | Ribo-Seq | 63.38 | 49.48 | 80.09 | 10.81 |
| **2-6 h embryos** | *D. simulans* | Ribo-Seq | 53.46 | 38.09 | 77.27 | 13.72 |
| **6-12 h embryos** | *D. simulans* | Ribo-Seq | 57.76 | 43.84 | 81.58 | 10.81 |
| **12-24 h embryos** | *D. simulans* | Ribo-Seq | 50.55 | 35.05 | 83.55 | 8.21 |
| **Third-instar larvae** | *D. simulans* | Ribo-Seq | 67.69 | 35.40 | 86.44 | 7.10 |
| **P7-8 pupae** | *D. simulans* | Ribo-Seq | 61.74 | 40.45 | 87.45 | 4.76 |
| **Female adult heads** | *D. simulans* | Ribo-Seq | 62.52 | 26.18 | 86.51 | 4.58 |
| **Male adult heads** | *D. simulans* | Ribo-Seq | 65.07 | 28.99 | 89.62 | 3.96 |
| **Female adult bodies** | *D. simulans* | Ribo-Seq | 59.47 | 12.91 | 83.01 | 5.72 |
| **Male adult bodies** | *D. simulans* | Ribo-Seq | 53.55 | 18.35 | 87.44 | 5.61 |

* The mRNA-seq and Ribo-Seq datasets for *D. melanogaster* were previously generated from Zhang *et al.* [10]. The datasets for *D. simulans* were generated in this study.

** After removing the reads mapped to rRNA, miscRNA (snoRNA, snRNA, rRNA, tRNA), the yeast genome, and the Wolbachia genome.

**Table S3.** Genes with uORFs showing strong evidence of translational buffering.
(see TableS3_strong_buffer_uORFs.csv)

**Table S4**. Mapping statistics of mRNA-Seq libraries for *bcd* uKO2/uKO2 mutant and WT flies

| Sample | Condition | Stain | Stage | Total read-pairs (M) | Unique mapping reads (M) | Unique mapping rate (%) | Multiple mapping rate (%) |
|---|---|---|---|---|---|---|---|
| **uKO2-0-2 h-embryo-29-rep1** | 29℃ | Mutant | 0-2 h | 19.39 | 16.73 | 86.31 | 9.13 |
| **uKO2-0-2 h-embryo-29-rep2** | 29℃ | Mutant | 0-2 h | 18.72 | 12.72 | 67.94 | 26.64 |
| **uKO2-2-6 h-embryo-29-rep1** | 29℃ | Mutant | 2-6 h | 19.60 | 16.76 | 85.51 | 11.60 |
| **uKO2-2-6 h-embryo-29-rep2** | 29℃ | Mutant | 2-6 h | 18.01 | 15.40 | 85.52 | 11.53 |
| **uKO2-6-12 h-embryo-29-rep1** | 29℃ | Mutant | 6-12 h | 18.55 | 16.26 | 87.64 | 7.52 |
| **uKO2-6-12 h-embryo-29-rep2** | 29℃ | Mutant | 6-12 h | 19.76 | 17.31 | 87.58 | 7.79 |
| **uKO2-12-24 h-embryo-29-rep1** | 29℃ | Mutant | 12-24 h | 17.48 | 15.23 | 87.14 | 8.36 |
| **uKO2-12-24 h-embryo-29-rep2** | 29℃ | Mutant | 12-24 h | 18.78 | 15.98 | 85.09 | 10.05 |
| **w1118-0-2 h-embryo-29-rep1** | 29℃ | WT | 0-2 h | 19.10 | 17.24 | 90.25 | 6.87 |
| **w1118-0-2 h-embryo-29-rep2** | 29℃ | WT | 0-2 h | 19.02 | 17.12 | 90.01 | 7.38 |
| **w1118-2-6 h-embryo-29-rep1** | 29℃ | WT | 2-6 h | 18.54 | 15.60 | 84.11 | 12.40 |
| **w1118-2-6 h-embryo-29-rep2** | 29℃ | WT | 2-6 h | 19.24 | 16.34 | 84.93 | 11.84 |
| **w1118-6-12 h-embryo-29-rep1** | 29℃ | WT | 6-12 h | 18.61 | 15.87 | 85.28 | 10.50 |
| **w1118-6-12 h-embryo-29-rep2** | 29℃ | WT | 6-12 h | 19.48 | 16.83 | 86.39 | 9.45 |
| **w1118-12-24 h-embryo-29-rep1** | 29℃ | WT | 12-24 h | 18.93 | 16.44 | 86.85 | 6.96 |
| **w1118-12-24 h-embryo-29-rep2** | 29℃ | WT | 12-24 h | 19.70 | 16.95 | 86.03 | 7.69 |
| **uKO2-0-2 h-embryo-25-rep1** | 25℃ | Mutant | 0-2 h | 16.22 | 14.94 | 92.12 | 4.27 |
| **uKO2-0-2 h-embryo-25-rep2** | 25℃ | Mutant | 0-2 h | 16.23 | 14.91 | 91.91 | 4.15 |
| **uKO2-2-6 h-embryo-25-rep1** | 25℃ | Mutant | 2-6 h | 16.80 | 14.50 | 86.31 | 7.53 |
| **uKO2-2-6 h-embryo-25-rep2** | 25℃ | Mutant | 2-6 h | 19.34 | 16.58 | 85.71 | 7.61 |
| **uKO2-6-12 h-embryo-25-rep1** | 25℃ | Mutant | 6-12 h | 16.17 | 14.20 | 87.80 | 8.49 |
| **uKO2-6-12 h-embryo-25-rep2** | 25℃ | Mutant | 6-12 h | 16.34 | 14.36 | 87.90 | 8.11 |
| **uKO2-12-24 h-embryo-25-rep1** | 25℃ | Mutant | 12-24 h | 16.85 | 14.35 | 85.20 | 11.33 |
| **uKO2-12-24 h-embryo-25-rep2** | 25℃ | Mutant | 12-24 h | 16.88 | 14.25 | 84.44 | 12.32 |
| **w1118-0-2 h-embryo-25-rep1** | 25℃ | WT | 0-2 h | 17.29 | 15.74 | 91.06 | 5.27 |
| **w1118-0-2 h-embryo-25-rep2** | 25℃ | WT | 0-2 h | 19.53 | 17.83 | 91.26 | 5.06 |
| **w1118-2-6 h-embryo-25-rep1** | 25℃ | WT | 2-6 h | 19.23 | 16.85 | 87.62 | 7.76 |
| **w1118-2-6 h-embryo-25-rep2** | 25℃ | WT | 2-6 h | 17.22 | 14.86 | 86.27 | 9.67 |
| **w1118-6-12 h-embryo-25-rep1** | 25℃ | WT | 6-12 h | 19.04 | 16.02 | 84.15 | 11.76 |
| **w1118-6-12 h-embryo-25-rep2** | 25℃ | WT | 6-12 h | 17.59 | 15.00 | 85.31 | 10.71 |
| **w1118-12-24 h-embryo-25-rep1** | 25℃ | WT | 12-24 h | 16.25 | 14.03 | 86.35 | 10.29 |
| **w1118-12-24 h-embryo-25-rep2** | 25℃ | WT | 12-24 h | 16.23 | 14.08 | 86.73 | 9.37 |

**Table S5.** Numbers of genes showing different magnitudes of TE changes between uORFs and CDS at the interspecific level, *H. sapiens* and *M. mulatta*.

| Tissues | # of expressed uORFs | $\beta_u \neq 1$ (%) | # of expressed CDSs | $\beta_c \neq 1$ (%) | uORF-CDS pairs with $\beta_u > 1$ | | | uORF-CDS pairs with $\beta_u < 1$ | | |
|---|---|---|---|---|---|---|---|---|---|---|
| | | | | | Total | $\gamma > 1$ | $\gamma < 1$ | Total | $\gamma > 1$ | $\gamma < 1$ |
| **Brain** | 7,380 | 80 (1.08) | 15,086 | 507 (3.36) | 51 | 0 | 27 | 29 | 14 | 0 |
| **Liver** | 4,429 | 28 (0.63) | 13,246 | 149 (1.24) | 10 | 0 | 4 | 18 | 10 | 0 |
| **Testis** | 7,384 | 53 (0.72) | 15,134 | 272 (1.80) | 33 | 0 | 9 | 20 | 5 | 0 |

Only uORFs and CDSs with an mRNA RPKM > 0.1 in both *H. sapiens and M. mulatta* were considered in each sample pair in the analysis. $\beta_u = TE_{uORF,macaque}/TE_{uORF,human}$ , is the fold change in $TE_{uORF}$ in *M. mulatta* relative to *H. sapiens* for each sample. $\beta_c = TE_{CDS,macaque}/TE_{CDS,human}$ , is the fold change in $TE_{CDS}$ in *M. mulatta* relative to *H. sapiens* for each sample. $\gamma = \beta_c/\beta_u$. For each CDS-uORF pair, $\beta_u$>1 and $\gamma$ < 1 or $\beta_u$<1 and $\gamma$ > 1 means that the magnitude of TE change is lower for a CDS than a uORF. The statistical significance of $\beta_c$, $\beta_u$, and $\gamma$ were all determined at an FDR < 0.05.

**Table S6.** The primer sequences used in this study

| Primer | Sequence (5'->3') | Application |
|---|---|---|
| Bcd-uORF-sgRNA-F | ATCGCAAAAACGCAAAATGT | sgRNA for Bcd uORF knock-out |
| Bcd-uORF-sgRNA-R | ACATTTTGCGTTTTTGCGAT | |
| Bcd-genotyping-F | GCTTTGCCGTACTGTTCGAT | Primers used for Bcd genotyping |
| Bcd-genotyping-R | AACTGAAGCTGCGGATGTTG | |
| Bcd-qPCR-F | GATGTATCTGGGTGGCTGCT | RT–qPCR primers used for Bcd mRNA P-to-M ratio quantification |
| Bcd-qPCR-R | CCGAAATGTGGGACGATAAC | |
| Rp49-qPCR-F | CACTTCATCCGCCACCAGTC | RT–qPCR primers used for rp49 as inter reference |
| Rp49-qPCR-R | CGCTTGTTCGATCCGTAACC | |
| Bcd-5utr-PC1-wF1 | CATCCCTAAATAACGGCACT | Used for WT and uATG-mutated *bcd* 5'UTR amplification |
| Bcd-5utr-PC1-wR | CGCGAACTTATTTCGCCCACATTTTGCGTTTTTG | |
| Bcd-5utr-PC1-mR1 | CGCGAACTTATTTCGCCCACTTATTTTGCGTTTTTG | |
| Bcd-5utr-PC1-mR2 | CGCGAACTTATTTCGAACTTATTTGCGATCCACTGCT | |
| Bcd-5utr-PC2-wF1 | CGAAATAAGTTCGCGAGCGT | |
| Bcd-5utr-PC2-wR1 | GTACACCTTGGAAGCCGGCGGTTGCGCCATTTTCC | |
| btd_qP_F | CACCAATTACCGCCCGTATG | RT-qPCR Primers for Bcd target genes |
| btd_qP_R | CGGCCAATACCTTCTTGCTC | |
| kr_qP_F | CCAGTGGCAGCTCTCCTAAT | |
| kr_qP_R | GCGAGATGGATCCTTGGAGA | |
| otd_qP_F | CTACAACATGGGCCATTCGG | |
| otd_qP_R | GCGACATGTAATCCAGGCTG | |
| Toll_qP_F | TTACTTCAGTGCCTCGCTCA | |
| Toll_qP_R | CCAGAGCCGATAACATTGCC | |
| hb_qP_F | GGAGAAGGAGCACGATCAGA | |
| hb_qP_R | TGCCGTGCGAATTGTAGATG | |
| Hairless_qP_F | GCATACATTCCCTCGCCTTC | |
| Hairless_qP_R | CGCTCCTGTCTGATAGTGGT | |

| | |
|---|---|
| salm_qP_F | GCCCATTCAAGTGCAGGATC |
| salm_qP_R | GAAGTTTCTCATCGGCGGAC |
| D_qP_F | CAAGAGGGTCACATCAAGCG |
| D_qP_R | CAGAGTTGTGCATCTTGGGG |
| Ten-m_qP_F | CCACCGCATCCTAAAGCATG |
| Ten-m_qP_R | CTCTTTCTCCTCCTCGGTCC |
| rib_qP_F | TTCGCCCTACACAGATCGTT |
| rib_qP_R | CGCACATCATATCCGCTCTG |
| CG1677_qP_F | TTGGCCGGGGATCATATGAA |
| CG1677_qP_R | GGTGGAATACGACGAGTTGC |
| Nin_qP_F | TAATGACCGCGATCACTGGA |
| Nin_qP_R | GCACCAGTTCAAGTTCCGTT |
| gt_qP_F | CCCTCGCCGTATCAAACAAG |
| gt_qP_R | GGGCGGCATACAGAAGATTG |
| asp_qP_F | AGCTACCATTTTCCCGGACA |
| asp_qP_R | CATCTGGAGTGATTCGACGC |
| CREG_qP_F | CTGATTTTGGCCCTCGTGAG |
| CREG_qP_R | GGCGATCTTGGCGTGATTTA |
| slp2_qP_F | GCAAGCCGGTGAAGGATAAG |
| slp2_qP_R | ACTCTGACGAATGGCCATCA |
| edl_qP_F | TCAAATGCTGGACAAGTGCC |
| edl_qP_R | GCGCTCATCAGTCGATTGTT |
| Mppe_qP_F | TGGACTTTAGCAACCCGGAT |
| Mppe_qP_R | CGCTGATGATGGCTCTCAAC |
| cnc_qP_F | GGAAACAGCAGCAACTTGGA |
| cnc_qP_R | CCGGCACTGAAATGGGTATG |
| ems_qP_F | ATCTATCCGGTCCCCAGAGT |
| ems_qP_R | CCTGGGATCAAGGTGAGTGT |
